## Supplementary Materials for "Information, certainty, and learning"

### Contents

| Contents | Pages |
|---|---|
| **Figure S1** Mean response rate for each group across sessions or time since CS onset ………. | 3 |
| **Figure S2** and analyses of terminal response rates (last 5 sessions) ………………………………… | 4 – 5 |
| **Figure S3** Response rate and trials to criterion for an individual rat ………………………………… | 6 |
| Additional analyses of trials to criterion using *the methods adopted by Thrailkill, Todd and Bouton (2020) and Kirkpatrick and Church (2000).and* ***Figure S4***.……………. | 7– 9 |
| **Data plots for each individual rat** ……………………………………………………………………… | 10– 185 |

*Each 6-panel figure plots data from one of the 176 rats. The rat number and the CS informativeness (ι), based on the C/T ratio, are shown above the first panel in the top left of the figure.*

***Panel 1*** *plots the rat's mean response rate (number of pokes per second) during the CS and during the pre-CS period in the inter-trial interval (ITI) on each of the 42 conditioning sessions.*
***Panel 2*** *(top right) shows on each trial the rat's response rate during the CS (dark grey line) and during the pre-CS period (light grey line). On the same panel, the blue line shows the cumulative poke count during the CS plotted against the cumulative CS duration across trials. (For this, the cumulative poke count excluded the first response in each trial, and the cumulative CS duration excluded the latency to first poke and the time the rat was in the magazine.)*
***Panels 3 and 4*** *(middle row) plot the cumulative response rates during the CS (solid black line), during the ITI (dashed black line), and the overall response rate in the context (CS plus ITI; dotted black line) across trials. The thick red line plots the nDkl for the comparison between CS rate and overall (context) rate. In Panel 3 (left), response rates were calculated in the conventional manner, as the cumulative number of responses divided by total time. In Panel 4 (right), response rates were calculated as the cumulative number of responses excluding the first response in each CS divided by the remaining time out of the magazine (i.e., excluding the latency to 1st response in the CS and excluding the cumulative time in the magazine). The black vertical line marks the trial on which the cumulative CS response rate permanently exceeded the cumulative context response rate. The two red dashed vertical lines to the right of the black line mark the trial on which the nDkl reached 0.82 (Odds 4:1 that CS rate > Context rate) and 1.92 (p < .05 that CS rate = Context rate). Note that these latter two values for the nDkl were calculated starting from the trial on which the CS rate permanently exceeded the context rate (marked by the black vertical line) and therefore do not correspond to the nDkl values calculated from Trial 1 shown by the thick red plotted line. The unbroken red vertical line marks the trial when the nDkl was minimum. To make the initial changes in responding clearer, the x-axis of each plot is truncated at the trial number 1.5 times the trial number at which the p<.05 threshold was reached (e.g.,*

*if the nDkl reached p<.05 at Trial 100, the axis is truncated at trial 150), or after a minimum of 20 trials.*

***Panels 5 and 6*** *(bottom row) show the parsed estimates of the CS response rate and pre-CS response rate estimated from the response rates as described for Panel 4. In both panels, the vertical red lines mark estimates of acquisition based on the* $nD_{KL}$ *as the Earliest estimate (leftmost), and when the odds against the null hypothesis that the parsed CS and ITI rates were equal reached 4:1, 10:1, 20:1, and 100:1. (The x-axis in Panel 5 has been right-cropped 20 trials after the difference in parsed response rates reached Odds 20:1.)*

**Figure S1**. Mean response rate for each group across Sessions or across time since CS onset.

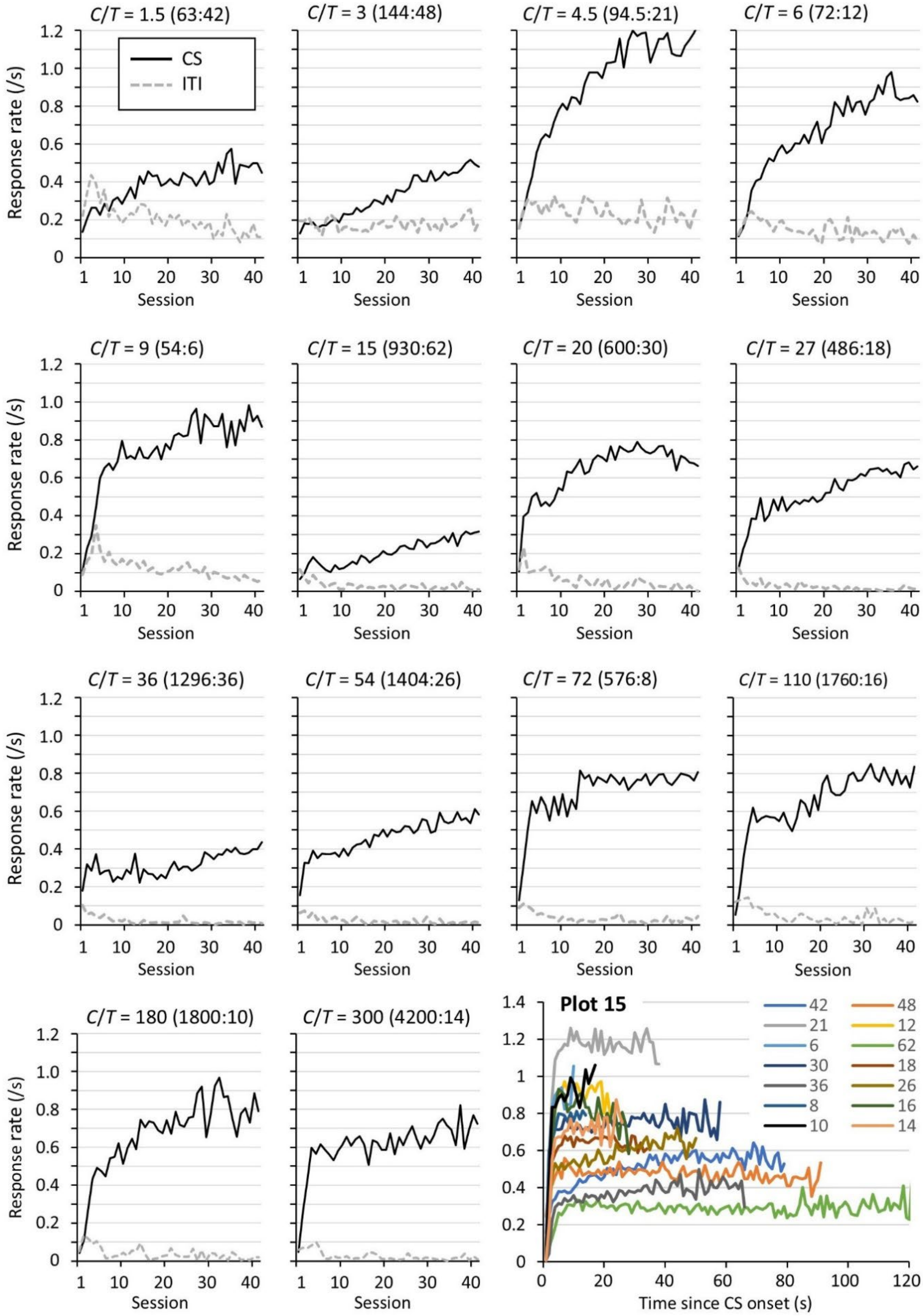


***Note***. *There were 10 (groups with C/T ≤ 72) or 3 (groups with C/T ≥ 110) trials per session. Plot 15 (bottom right) shows the mean response rate per second during the CS, averaged from the last 5 sessions. Each group is identified by the length of the mean CS-US interval (T).*

**Analyses of terminal response rates (last 5 sessions).**

As a measure of the terminal level of responding, the response rates during the CS were averaged over the last 5 sessions for each rat. We have conducted the same analysis described here using the data from the final 10 session, with very similar results. (Response rate data of individual rats averaged from both the last 5 and last 10 sessions can be downloaded from https://osf.io/vmwzr/) We first calculated CS response rate as total number of responses during the CS (summed across all trials over the 5 sessions) divided by the total CS duration (summed across all trials over the 5 sessions). We then compared this terminal response rate with the log of *T*, *C*, and *C/T* ($\alpha$ = 0.017 after correction for multiple comparisons). The rate of responding to the CS was marginally correlated with log(*T*) (see Figure S2A), $r$ = -0.60, $p$ = .025, but was not correlated with either log(*C*), $r$ = -0.38, $p$ = .181, or log(*C/T*), $r$ = -0.08, $p$ =.778.

**Figure S2**. Response metrics, over the final 5 conditioning sessions, plotted against *T.*

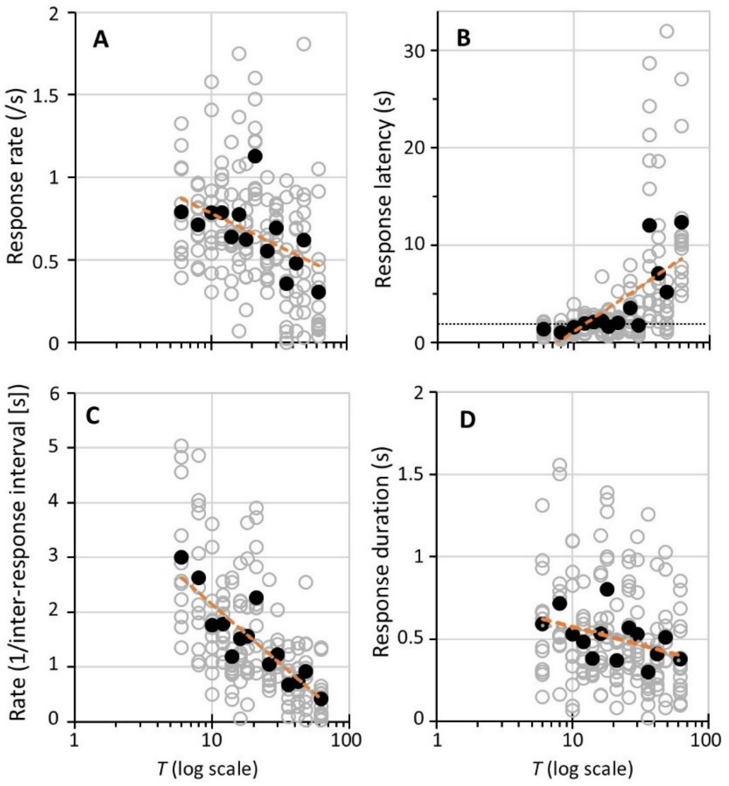


***Note****: Open grey circles show data for individual rats. Black filled circles show mean data for each group. The dashed orange lines are the best-fitting regression lines.*

More detailed analyses were conducted after segmenting the response rate into 3 separate components: (1) the latency to the first response in a trial; (2) the mean duration of each response (the time spent in the magazine); and (3) the inverse of the mean inter-response interval, 1/IRI, which equals the response rate after excluding the latency and response durations. The group medians for each of these indices was correlated with the log of *C*, *T*, and *C/T*. None correlated significantly with log(*C*), largest $r$ = -0.39, $p$ =.168, or with log(*C/T*), largest $r$ = -0.17, $p$ =.557. On the other hand, log(*T*) correlated significantly with latency, $r$ = 0.68, $p$ = .007 (Figure S2B), and with 1/IRI, $r$ = 0.87, $p$ < .001 (Figure S2C). Duration of responding did not correlate significantly with log(*T*), $r$ = -0.42, $p$ = .149 (Figure S2D). As is evident in Figure S2B, the positive correlation between latency and log(*T*) was largely confined to groups with long CS-US intervals ($T$ > 25 s), whereas latency varied little among groups with shorter CS-US intervals. This invariance at short CS-US intervals may have been due to a floor effect because mean latencies to first response did not decrease below 2 s (horizontal dotted line in Figure S2B). This is also consistent with the plots of response by time-in-CS, shown in plot 15 of Figure S1, where responding to the CS was low for the first few seconds after CS onset. This apparent floor effect might reflect a constraint on how quickly the rats can commence responding after CS onset or it might have arisen because 2 s was the minimum CS-US interval used in this experiment. Regardless, these analyses indicate that neither duration of responding nor latency to first response are good markers of what the rats learn about the rate of reinforcement of the CS. By contrast, the response rate, 1/IRI, varied systematically across the entire range of values of *T* (Figure S2C). This confirms earlier evidence that rats' response rates scale with the log of the reinforcement rate (Harris & Carpenter, 2011).

**Figure S3.** Response rate and trials to criterion for an individual rat.

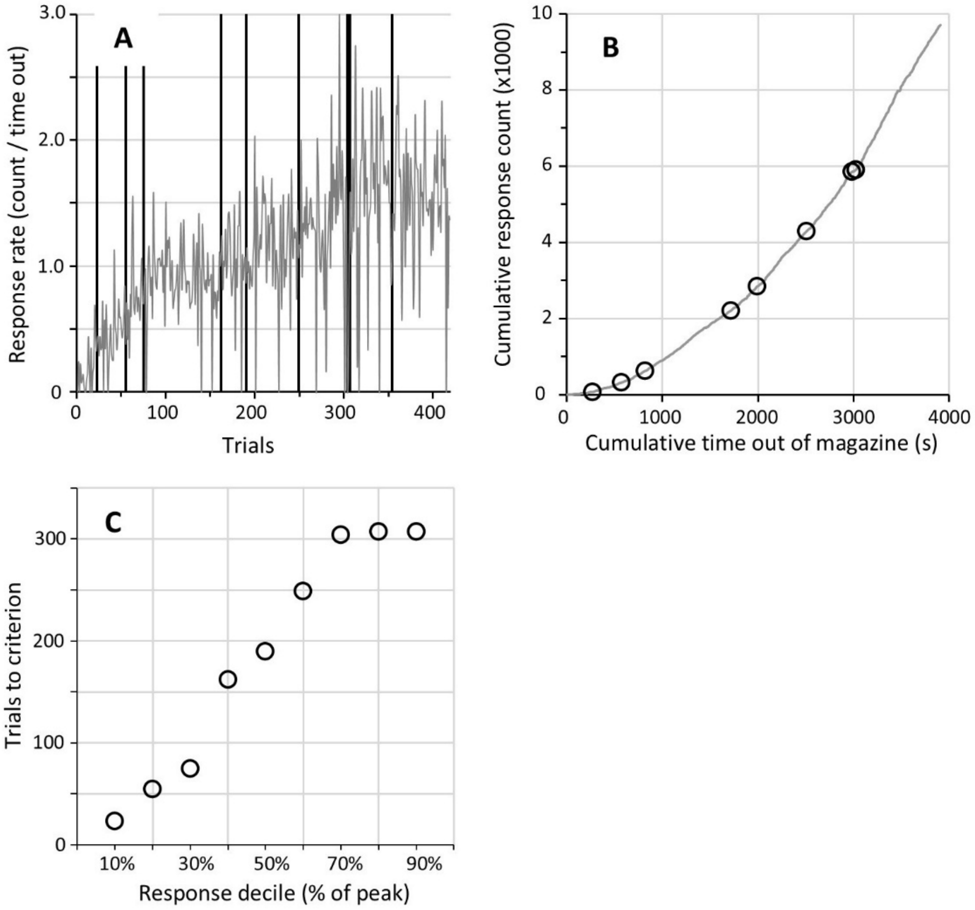


***Note.*** *The grey line in* ***A*** *shows the response rate on each trial for an individual rat (Rat 30, Group C/T = 4.5). The response rate was calculated as the number of responses during the CS divided by the total time out of the magazine but excluding the latency to first response (i.e., the total time across which the animal could respond). In* ***B****, the cumulative response count across trials for the same rat is plotted against the cumulative opportunity to respond (cumulative time out of the magazine). The slope of this cumulative function was used to identify the trial on which the rat's response rate reached each decile (from 10% to 90%) of its peak response rate, as shown in* ***C****. These trials are also marked as circles on the response plots in A and B.*

### Additional analyses of trials to criterion.

In addition to the analysis we described in the main article, we also conducted analyses of trials to criterion on our data following two approaches used previously by other researchers. One of these methods was used by **Thrailkill, Todd and Bouton (2020)**. Their method identified the point of acquisition using a criterion in which the mean response rate during the CS had to exceed the mean pre-CS response rate by at least 1.5 responses/10 s (9 responses/min) on 3 consecutive 4-trial blocks. We applied this method to our data but we multiplied the obtained value by 4 in order to convert the blocks-to-criterion results back into trials-to-criterion so that the results could be compared directly to those of our own analyses. The trials to criterion for each group is shown in Figure S5, plotted against the *C*/*T* ratio, which also plots for comparison the trials to criterion identified as the trial after which the $nD_{kl}$ became permanently positive (shown in Figure 3A of the main article). The group means of trials to criterion produced using the two methods correlated well with each other, r = .85, $p < .001$. However, the values obtained using the Thrailkill *et al*. method were on average 11 times higher than those identified using the $nD_{kl}$, $t(167) = 6.27$, $p < .001$. This difference indicates that the criterion used by Thrailkill *et al*. was sensitive to more advanced stages of response acquisition. Nonetheless, the scores from this method did scale with the log of the *C*/*T* ratio, as shown in Figure S5. The same correlational analyses described in the main article using the $nD_{kl}$ criterion were conducted from the results using the Thrailkill *et al*. method. The correlation between log of trials to criterion and log(*C*/*T*) was significant, $r = -0.83$, $p < .001$, whereas the correlations with log(*T*) and log(*C*) fell short of the corrected level of significance, $r = 0.61$ and -0.61, $p = .021$ and .020, respectively. Given that log(*C*/*T*) was correlated with both log(*C*) and log(*T*), partial correlations were calculated to assess the relationship between each variable and trials to criterion. The correlation between log(*C*/*T*) and trials to criterion remained significant after partialling out the effect of log(*T*), r = -.80, $p < .001$, or log(*C*), r = -.80, $p = .001$.

The analyses described above show that the relationship between learning rate (trials to an acquisition criterion) and informativeness still hold when using the acquisition criterion adopted by Thrailkill *et al*. (2020). Nonetheless, that criterion identifies a later point in the learning process than the point identified using the $nD_{kl}$. Therefore, their method is more likely to be affected by factors, such as *T*, that affect the subsequent strength of responding. This may explain why Thrailkill *et al*. observed an effect of *T* on the rate of learning even when *C*/*T* was held constant, whereas our analysis, using the $nD_{kl}$, did not.

**Figure S4.**

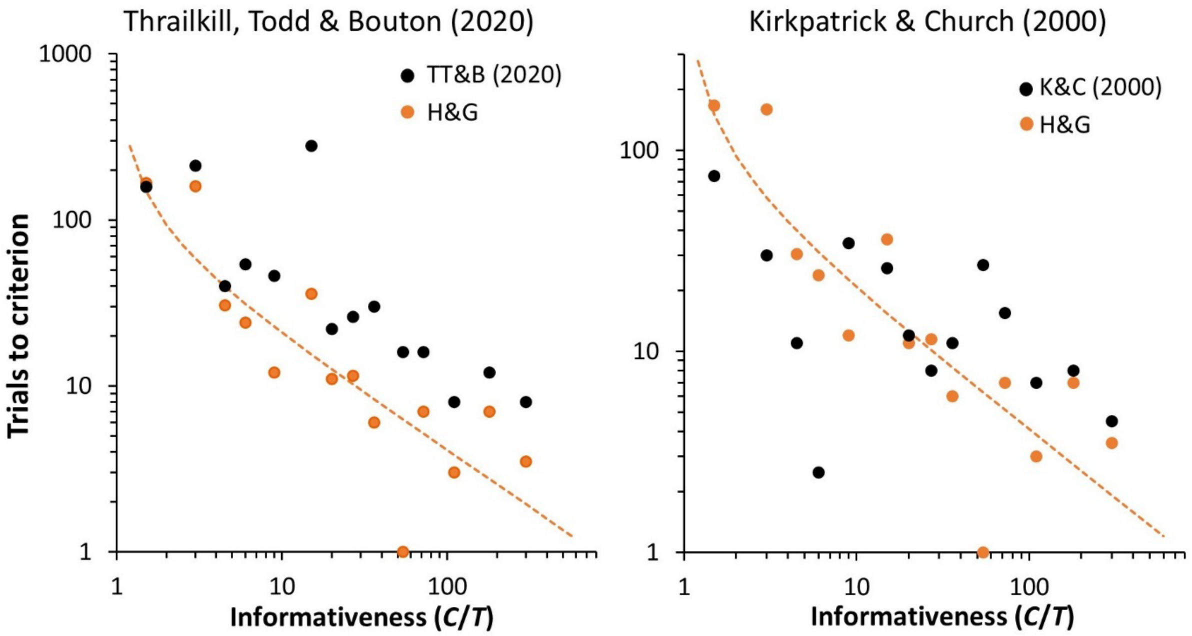


***Note*. The mean number of trials to criterion for each of the 14 groups of rats (with different C/T ratios) in the current experiment. In the plot on the left, the filled black circles show the number of trials for each group to reach the criterion used by Thrailkill, Todd and Bouton (2020). In the plot on the right, the filled black circles show the number of trials for each group to reach the criterion used by Kirkpatrick and Church (2000). For reference, the orange circles in both graphs show the number of trials for the difference between the cumulative CS response rate and cumulative ITI rate to become permanently positive (the criterion shown in Figure 2A of the main article).*

The final method we have used to analyse our data followed the method described by **Kirkpatrick and Church (2000)**. This was based on a discrimination ratio calculated as the number of responses ($R_{CS}$) made during a brief time window (of length 2/15$^{ths}$ of *T*) in the middle of the CS-US interval divided by the same response count plus a baseline response count ($R_b$) measured during a window of equivalent length in the middle of the pre-CS interval (i.e., $R_{CS}/[R_{CS}+R_b]$). To identify how quickly responding to the CS was acquired, Kirkpatrick and Church fitted a low-high step function to the discrimination ratios across trials to identify the trial, *t*, at which there was maximal change in the ratio averaged across trials before *t* versus trials after *t*. We have applied Kirkpatrick and Church's step-function method to discrimination ratios calculated from our data but using response rates across the full CS and pre-CS trial lengths rather than sampled from a small window (2/15$^{ths}$ of *T*) within those intervals. The trials to criterion identified with this method are shown in the right graph of Figure S5, plotted against the *C*/*T* ratio. As is evident in the scatter of values shown in the figure, this method for estimating trials to acquisition was less sensitive to differences in *C*/*T* than any of our criteria or the method used by Thrailkill *et al*. (2020). The log of trials to criterion was not significantly correlated with log(*C*/*T*), r = -.50, *p* = .071, or log(*C*), r = -.35, *p* = .221, or log(*T*), r = .40, *p* =

.157. The data obtained using Kirkpatrick and Church's method were not significantly correlated with the data produced by the criterion we used, r = .409, *p* = .147, and fell short of a significant correlation with the data produced using Thrailkill et al.'s method, r = .57, *p* = .034. One limitation with the Kirkpatrick and Church measure is that simple changes in response rates between the CS and baseline undergo a non-linear transformation when computed as a ratio. For example, as the pre-CS response count goes to zero, the ratio goes to a maximum of 1 regardless of how many responses are made during the CS. Because this algorithm uses response rates across the whole experiment to identify the acquisition trial, the trial selected will depend on when the discrimination ratio reached its peak, which will itself be strongly influenced by the decline in baseline responding, $R_b$.

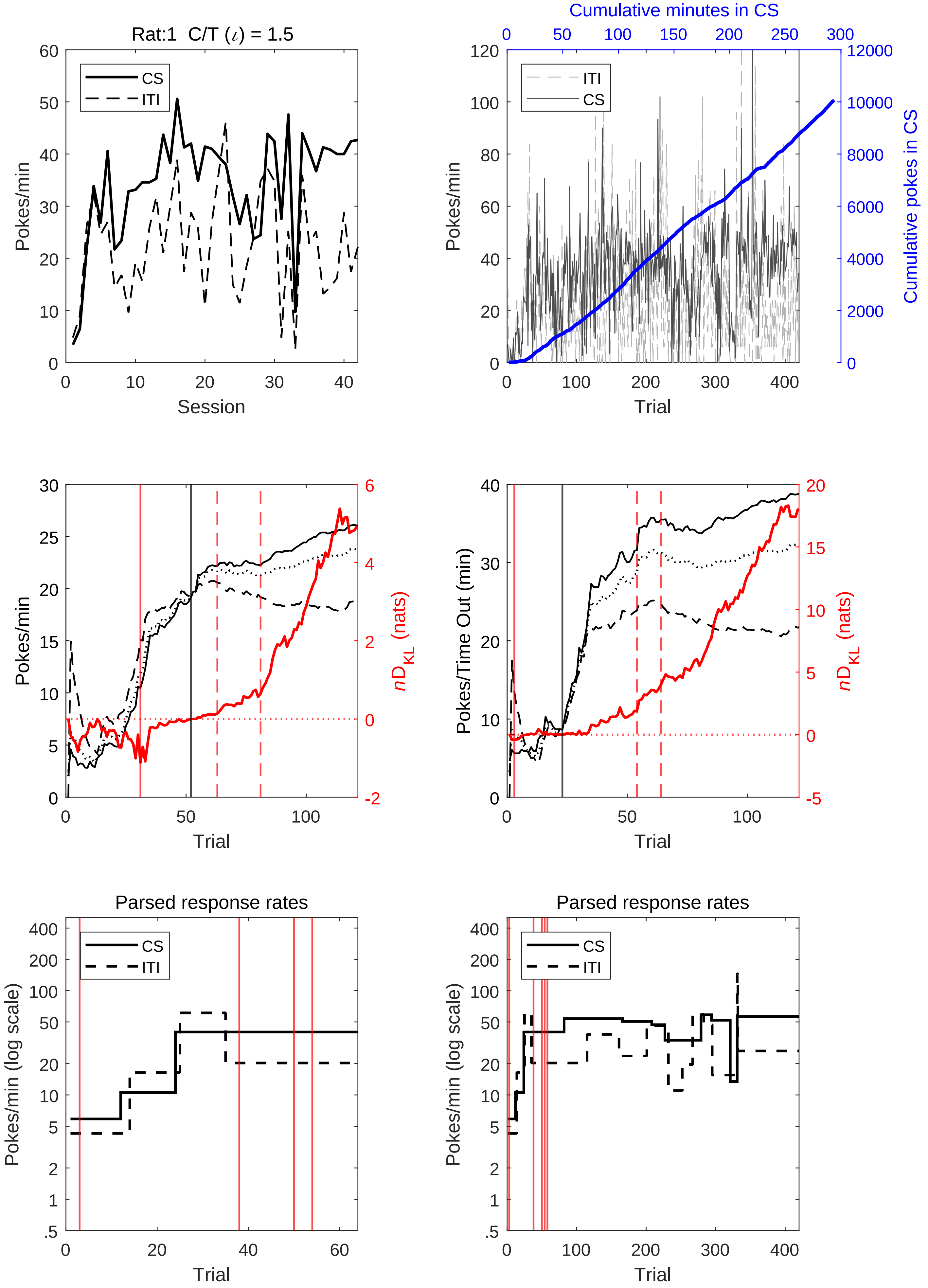

Rat:1 C/T (ι) = 1.5
CS
ITI
Pokes/min
Session
Cumulative minutes in CS
Cumulative pokes in CS
Trial
Pokes/Time Out (min)
nD_KL (nats)
Parsed response rates
Pokes/min (log scale)

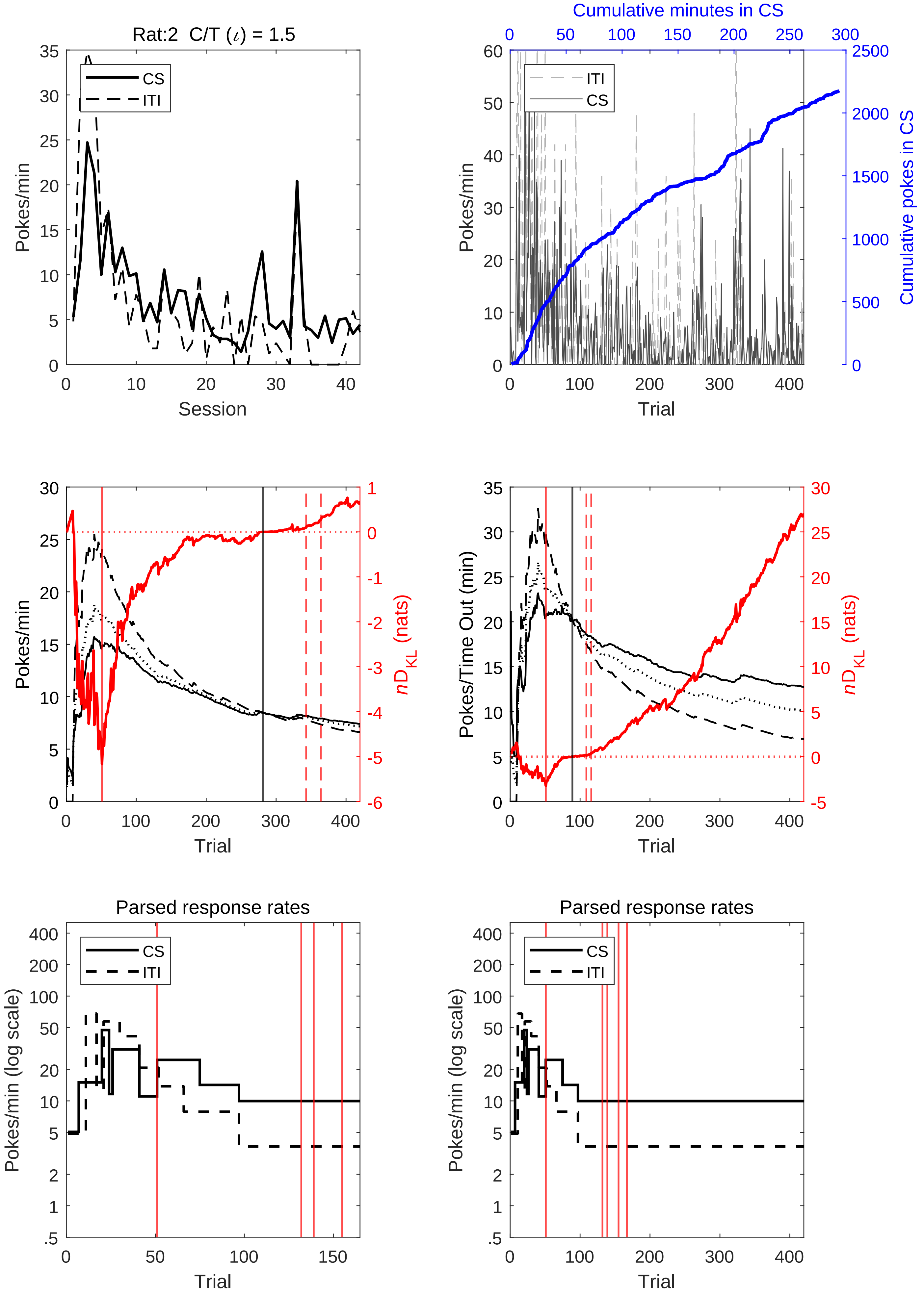
Rat:2 C/T (ι) = 1.5
CS
ITI
Pokes/min
Session
Cumulative minutes in CS
Cumulative pokes in CS
Trial
nDKL (nats)
Pokes/Time Out (min)
Parsed response rates
Pokes/min (log scale)

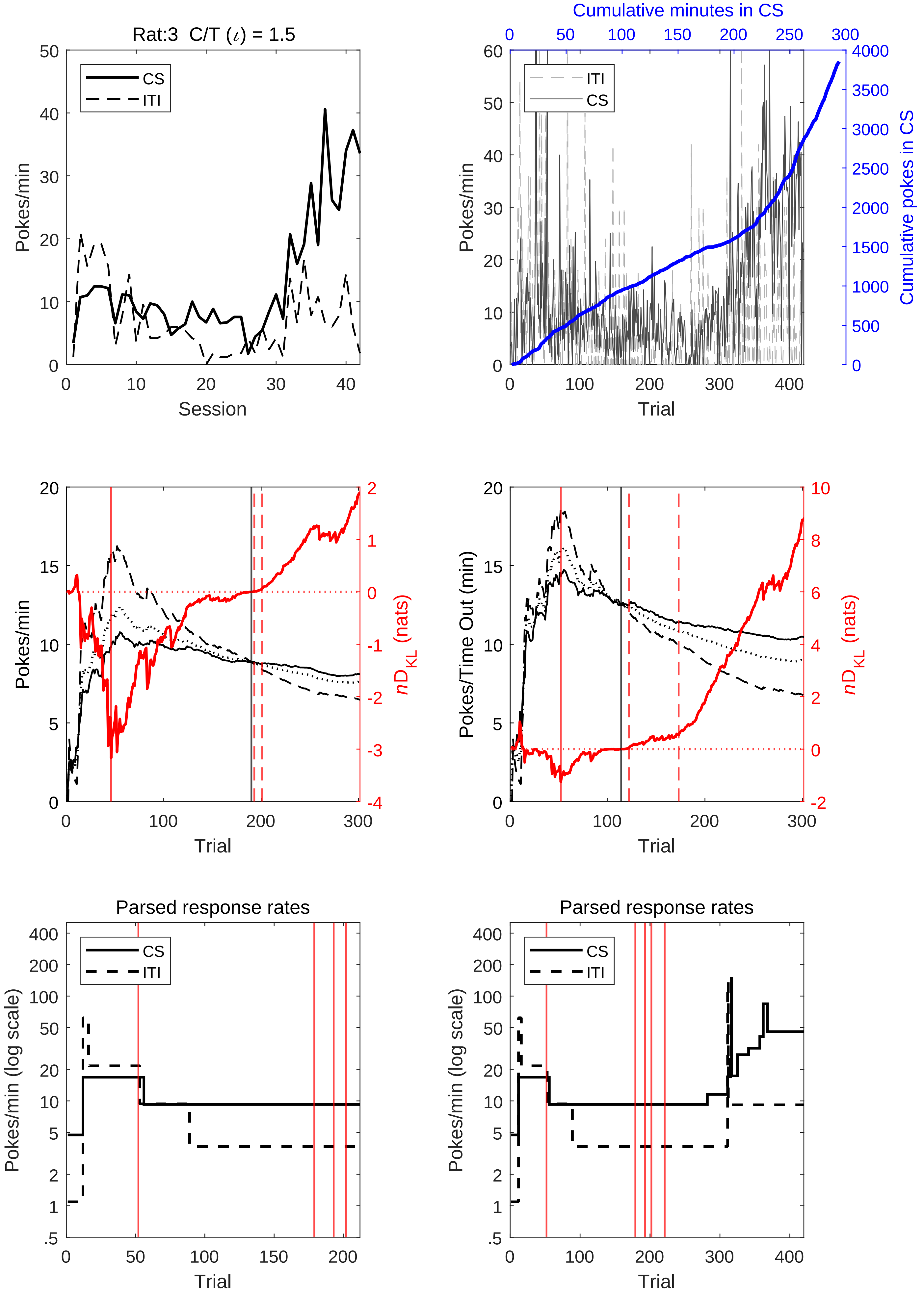
Rat:3 C/T (ι) = 1.5
Pokes/min
Session
CS
ITI
Cumulative minutes in CS
Pokes/min
Trial
Cumulative pokes in CS
Pokes/min
Trial
nD_KL (nats)
Pokes/Time Out (min)
Trial
nD_KL (nats)
Parsed response rates
Pokes/min (log scale)
Trial
Parsed response rates
Pokes/min (log scale)
Trial

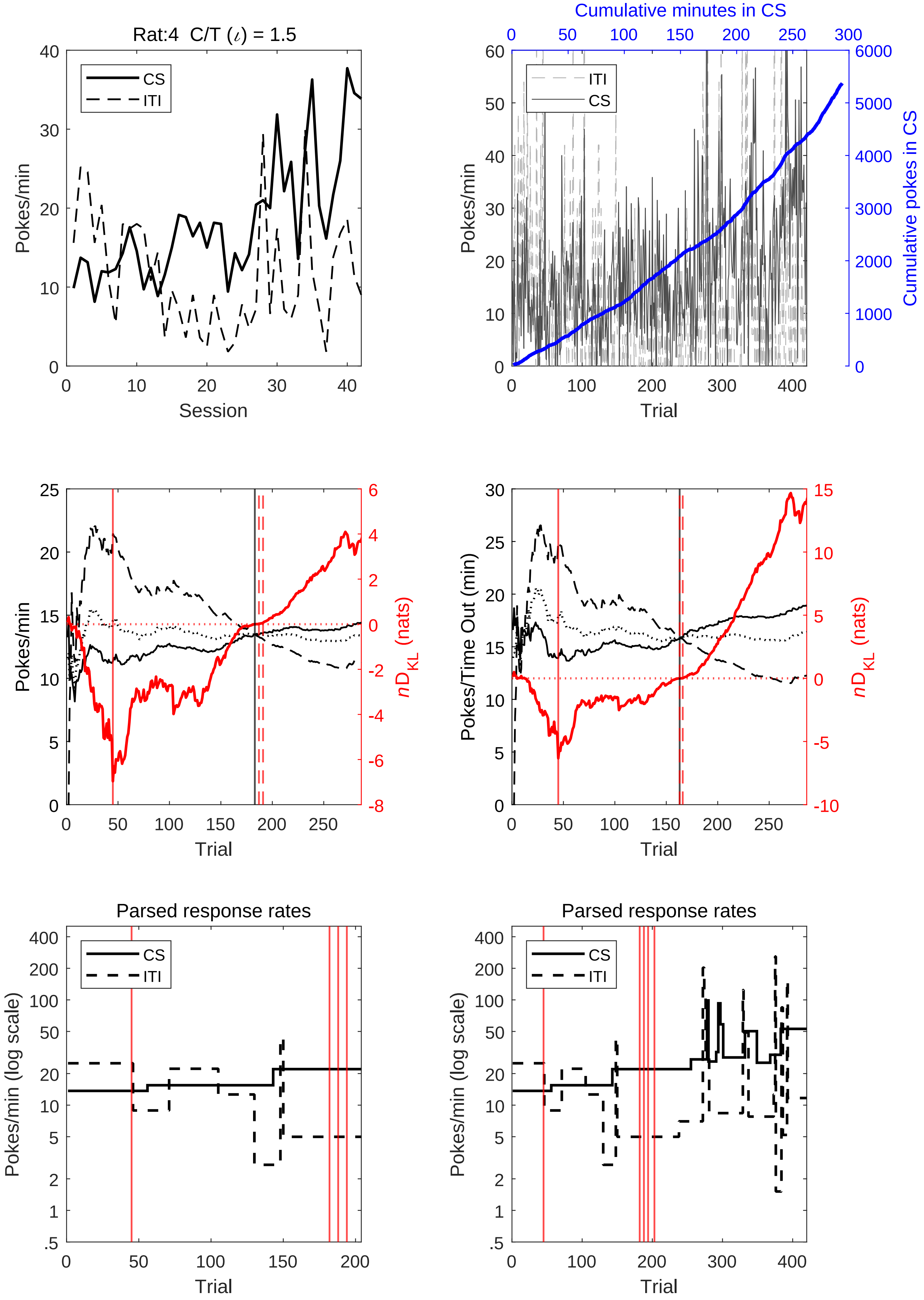
Rat:4 C/T (ι) = 1.5
CS
ITI
Pokes/min
Session
Cumulative minutes in CS
Cumulative pokes in CS
Trial
nD_KL (nats)
Pokes/Time Out (min)
Parsed response rates
Pokes/min (log scale)

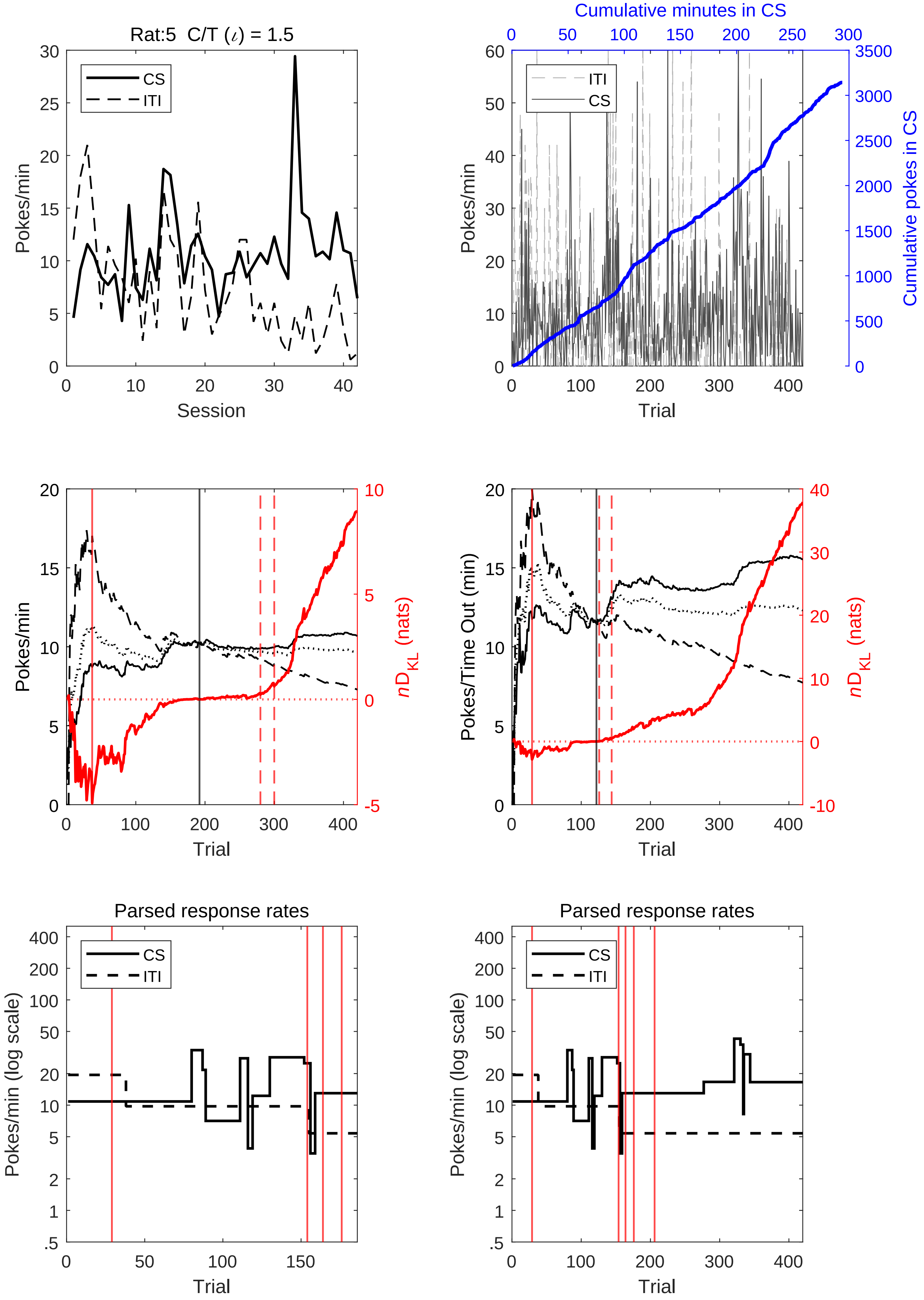
Rat:5 C/T (ι) = 1.5
CS
ITI
Pokes/min
Session
Cumulative minutes in CS
Cumulative pokes in CS
Trial
$nD_{KL}$ (nats)
Pokes/Time Out (min)
Parsed response rates
Pokes/min (log scale)

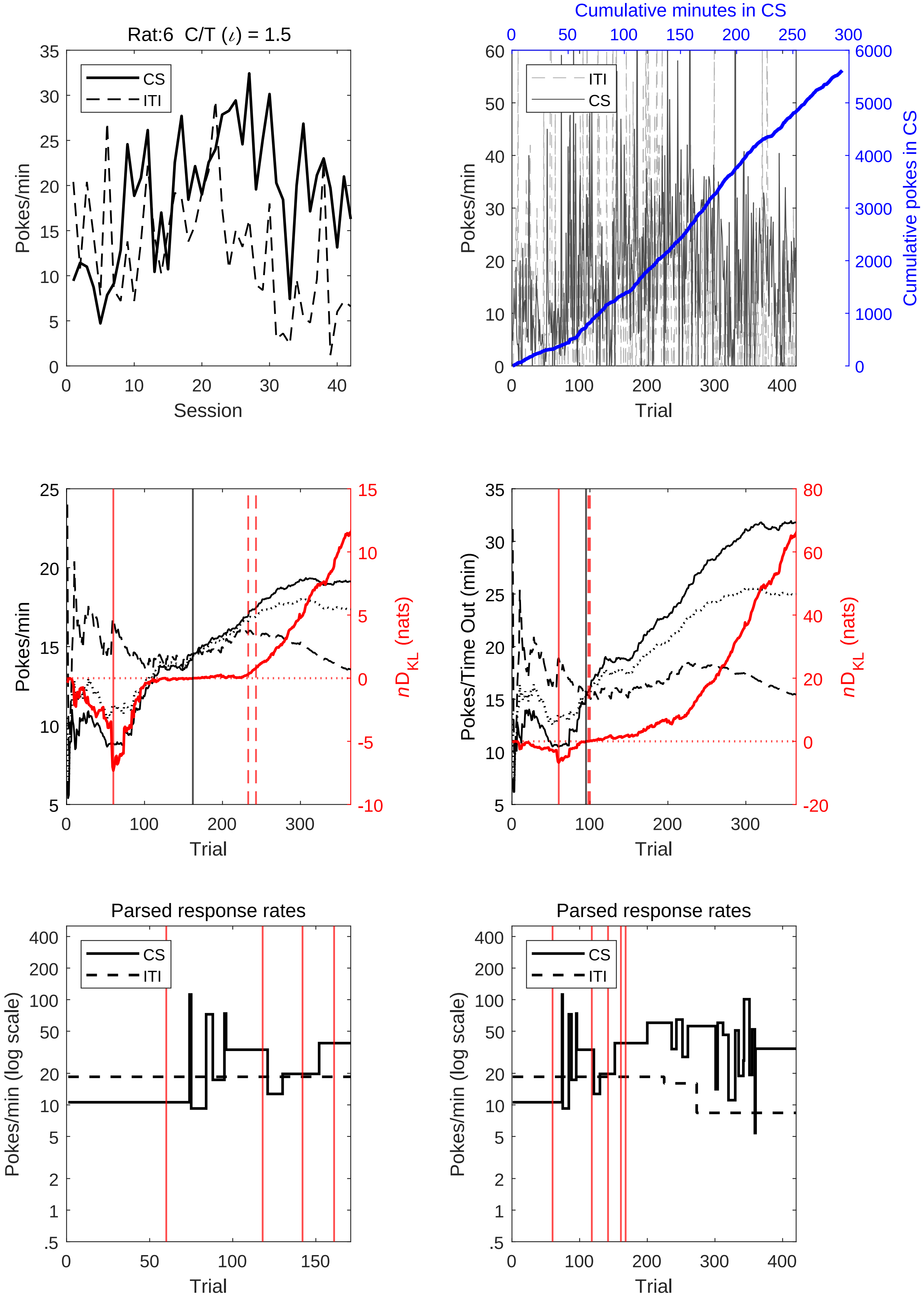
Rat:6 C/T (ι) = 1.5
CS
ITI
Pokes/min
Session
Cumulative minutes in CS
Cumulative pokes in CS
Trial
nD_KL (nats)
Pokes/Time Out (min)
Parsed response rates
Pokes/min (log scale)

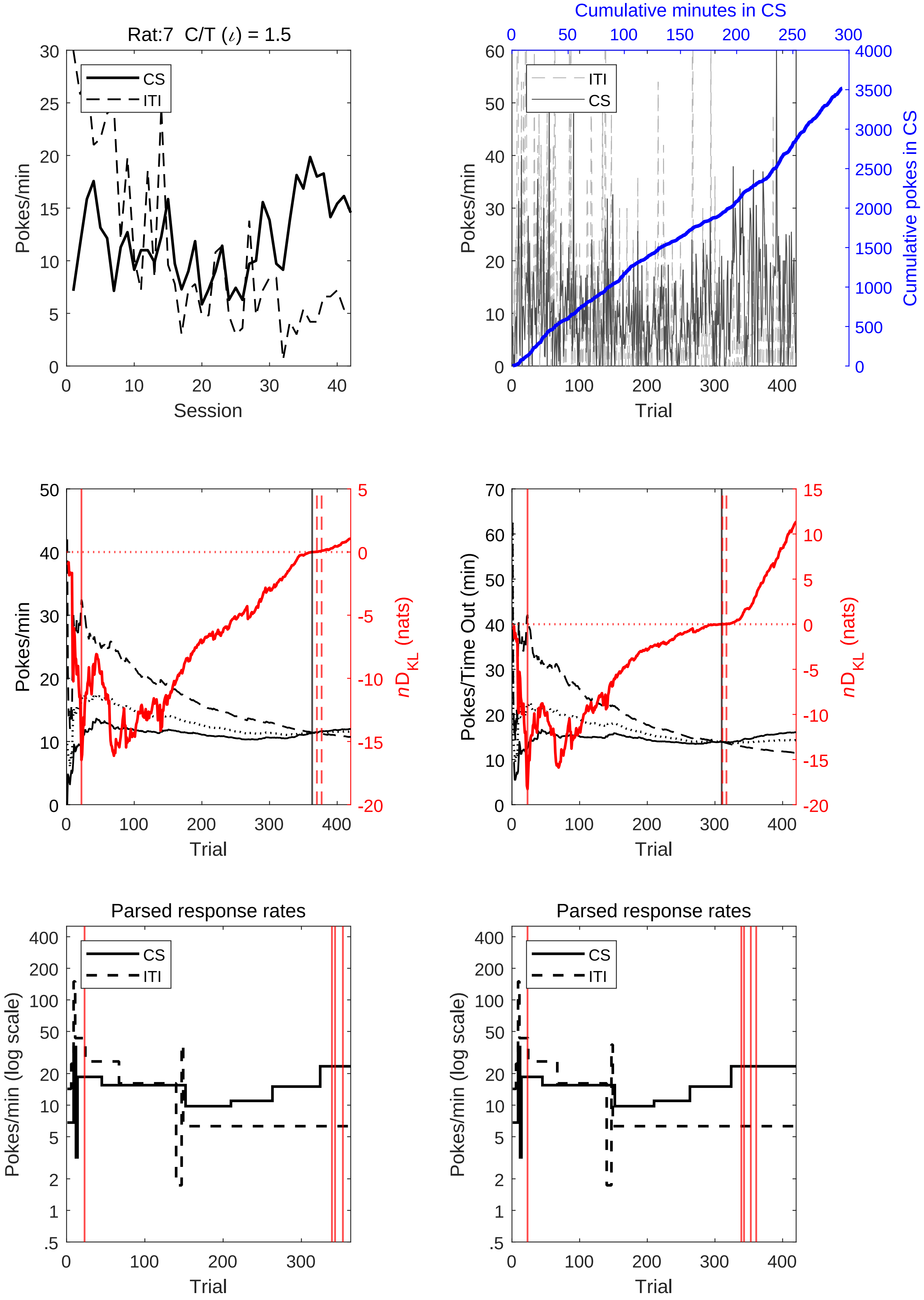

Rat:7 C/T (ι) = 1.5
CS
ITI
Pokes/min
Session
Cumulative minutes in CS
Cumulative pokes in CS
Trial
Pokes/Time Out (min)
nD_KL (nats)
Parsed response rates
Pokes/min (log scale)

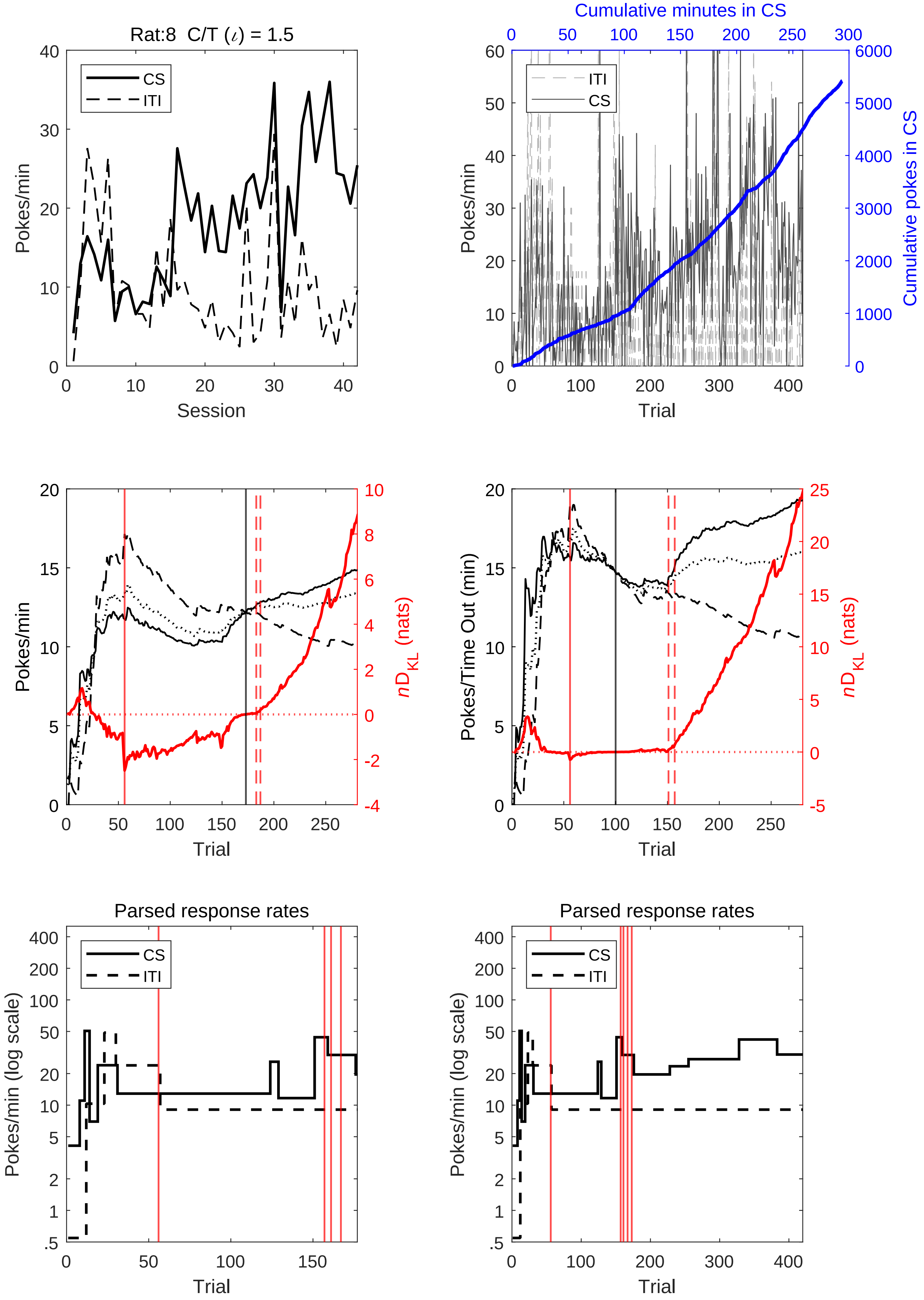

Rat:8 C/T ($\iota$) = 1.5
CS
ITI
Pokes/min
Session
Cumulative minutes in CS
Cumulative pokes in CS
Trial
Pokes/Time Out (min)
$nD_{KL}$ (nats)
Parsed response rates
Pokes/min (log scale)

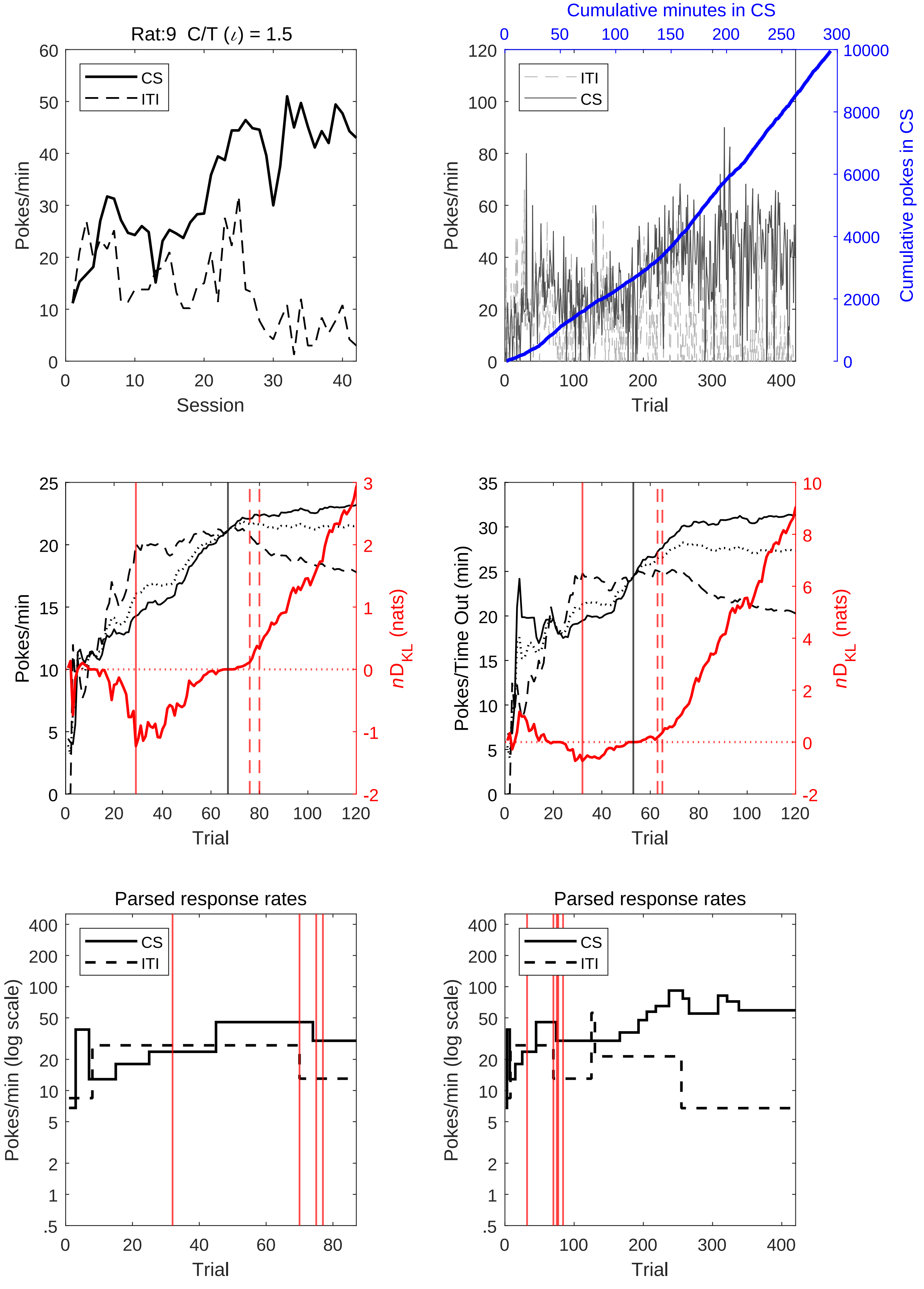

Rat:9 C/T (ι) = 1.5
Pokes/min
Session
CS
ITI
Cumulative minutes in CS
Cumulative pokes in CS
Trial
Pokes/Time Out (min)
nD_KL (nats)
Parsed response rates
Pokes/min (log scale)

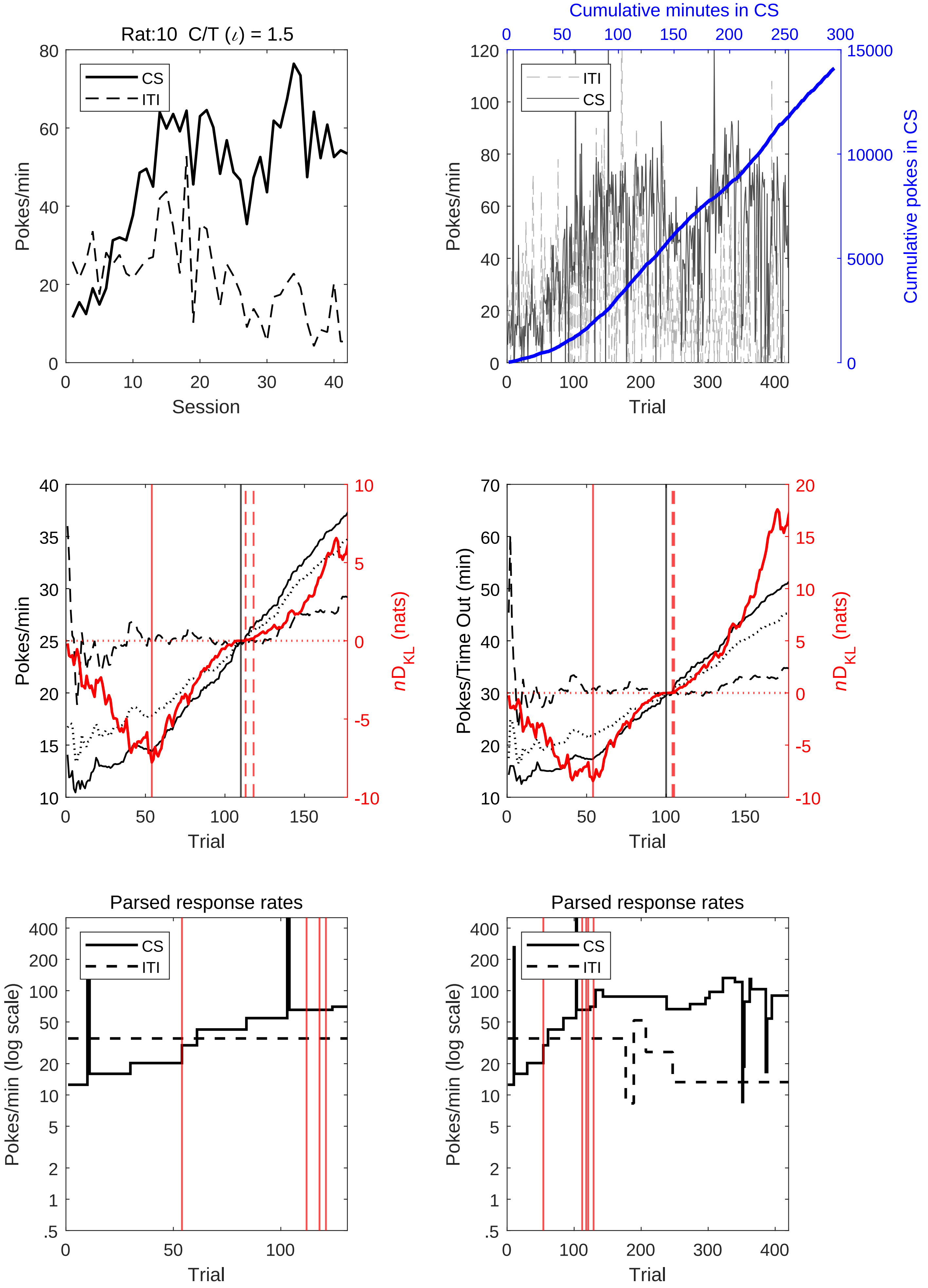

Rat:10 C/T ($\iota$) = 1.5
CS
ITI
Pokes/min
Session
Cumulative minutes in CS
ITI
CS
Pokes/min
Cumulative pokes in CS
Trial
Pokes/min
$nD_{KL}$ (nats)
Trial
Pokes/Time Out (min)
$nD_{KL}$ (nats)
Trial
Parsed response rates
CS
ITI
Pokes/min (log scale)
Trial
Parsed response rates
CS
ITI
Pokes/min (log scale)
Trial

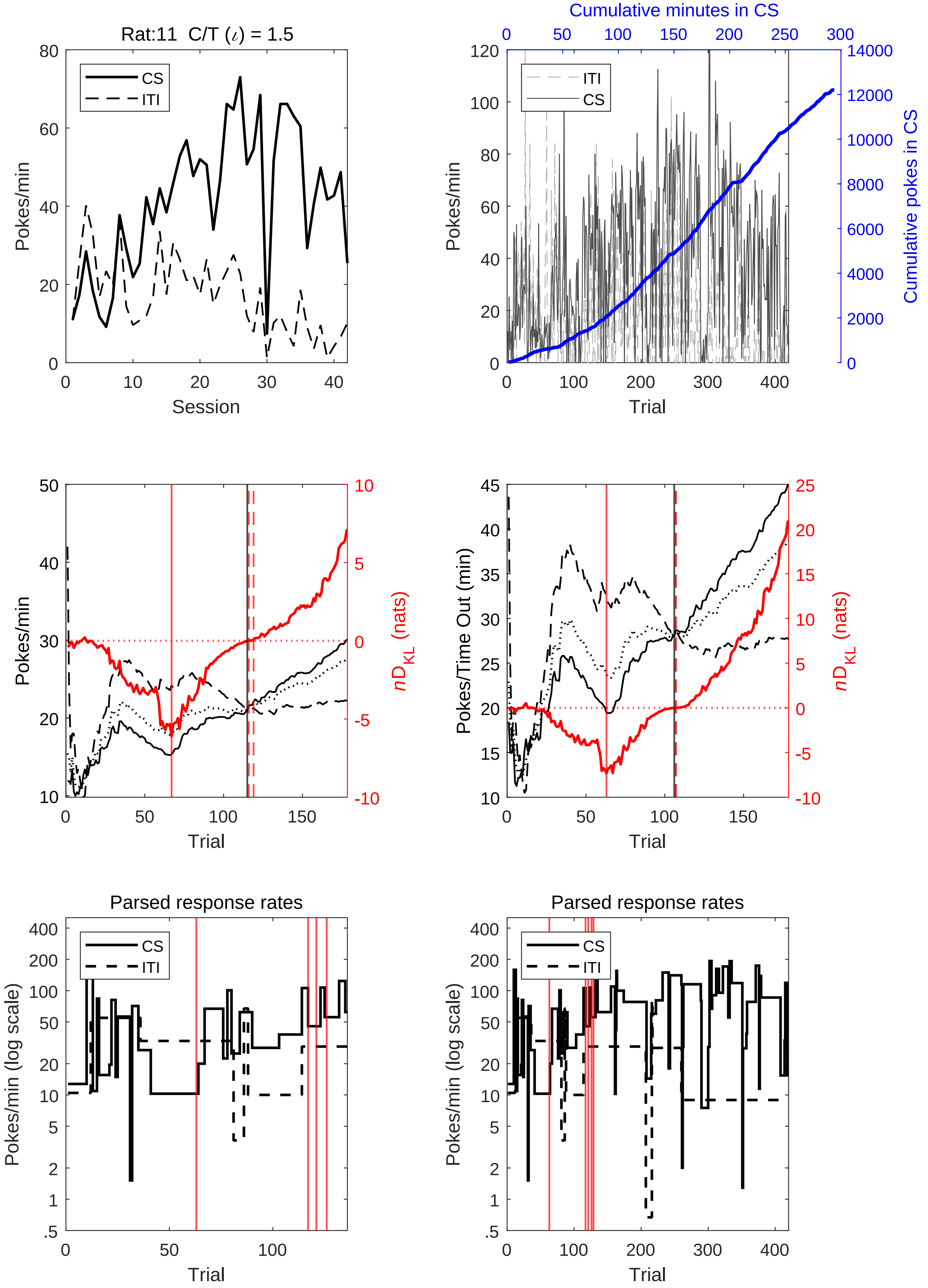

Rat:11 C/T (ι) = 1.5
CS
ITI
Pokes/min
Session
Cumulative minutes in CS
Cumulative pokes in CS
Trial
nD_KL (nats)
Pokes/Time Out (min)
Parsed response rates
Pokes/min (log scale)

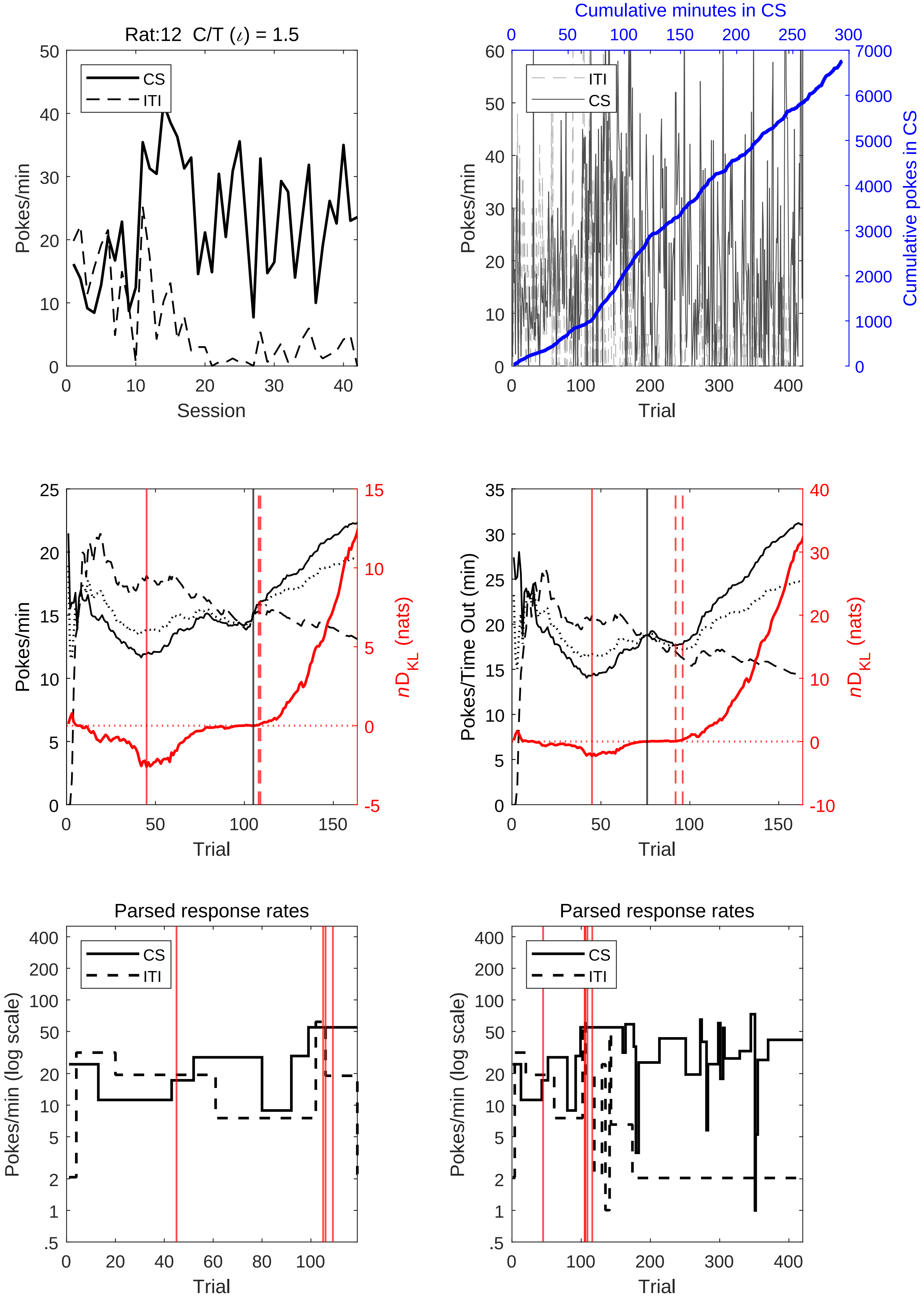
Rat:12 C/T ($\iota$) = 1.5
CS
ITI
Pokes/min
Session
Cumulative minutes in CS
Cumulative pokes in CS
Trial
$nD_{KL}$ (nats)
Pokes/Time Out (min)
Parsed response rates
Pokes/min (log scale)

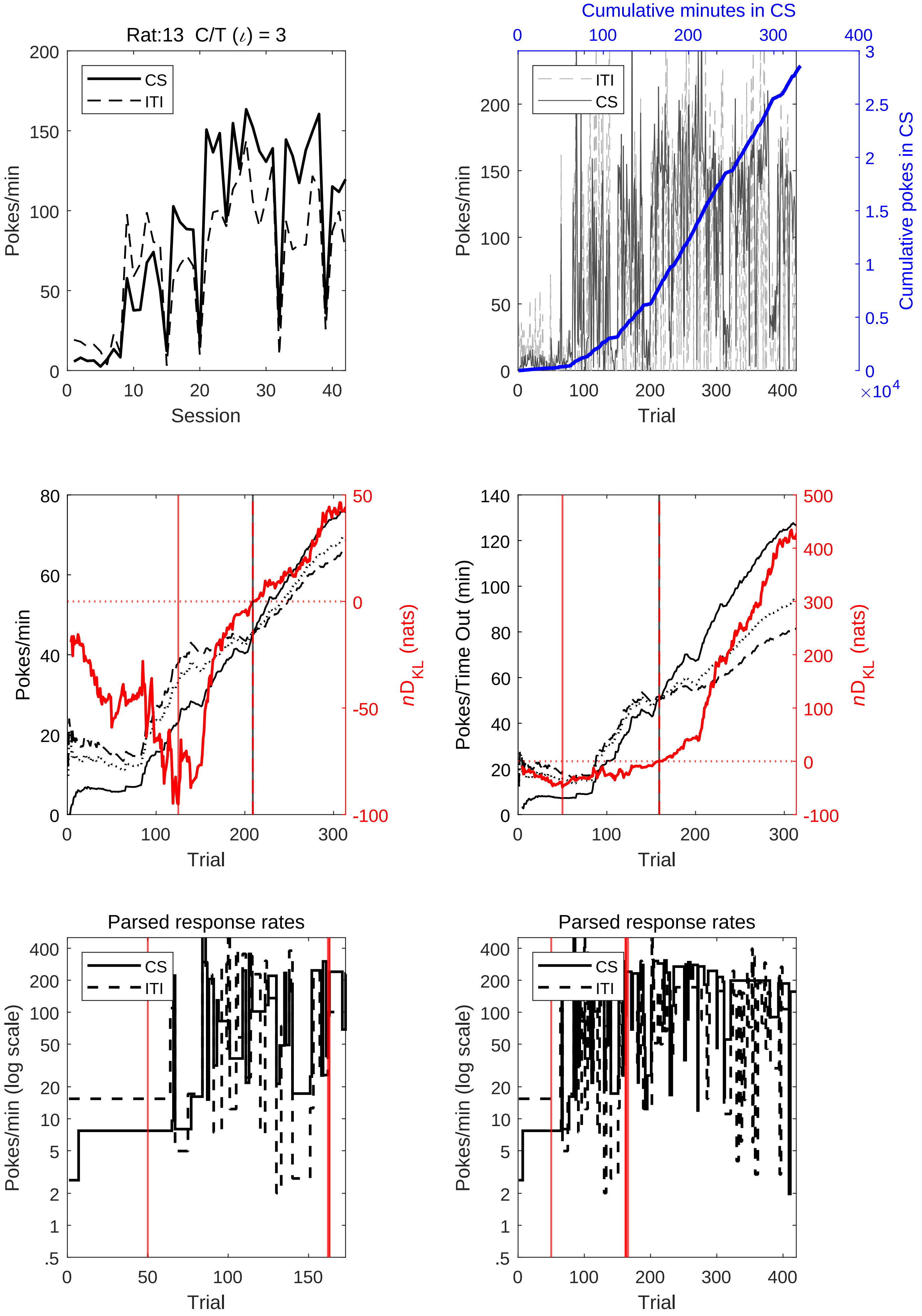
Rat:13 C/T (ι) = 3
Pokes/min
Session
CS
ITI
Cumulative minutes in CS
Cumulative pokes in CS
×10^4
Trial
Pokes/Time Out (min)
nD_KL (nats)
Parsed response rates
Pokes/min (log scale)

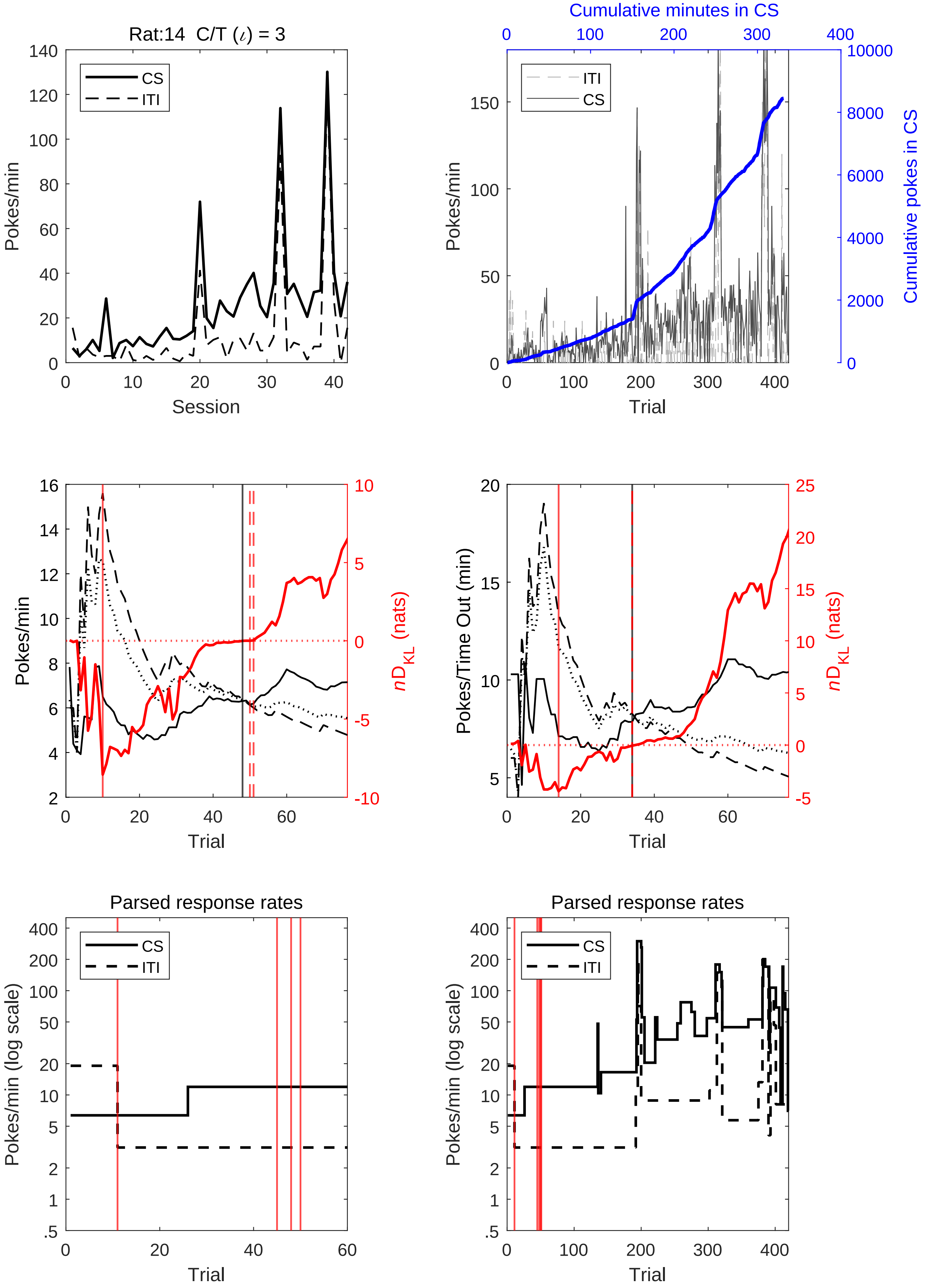
Rat:14 C/T (ι) = 3
CS
ITI
Pokes/min
Session
Cumulative minutes in CS
Cumulative pokes in CS
Trial
Pokes/Time Out (min)
nD_KL (nats)
Parsed response rates
Pokes/min (log scale)

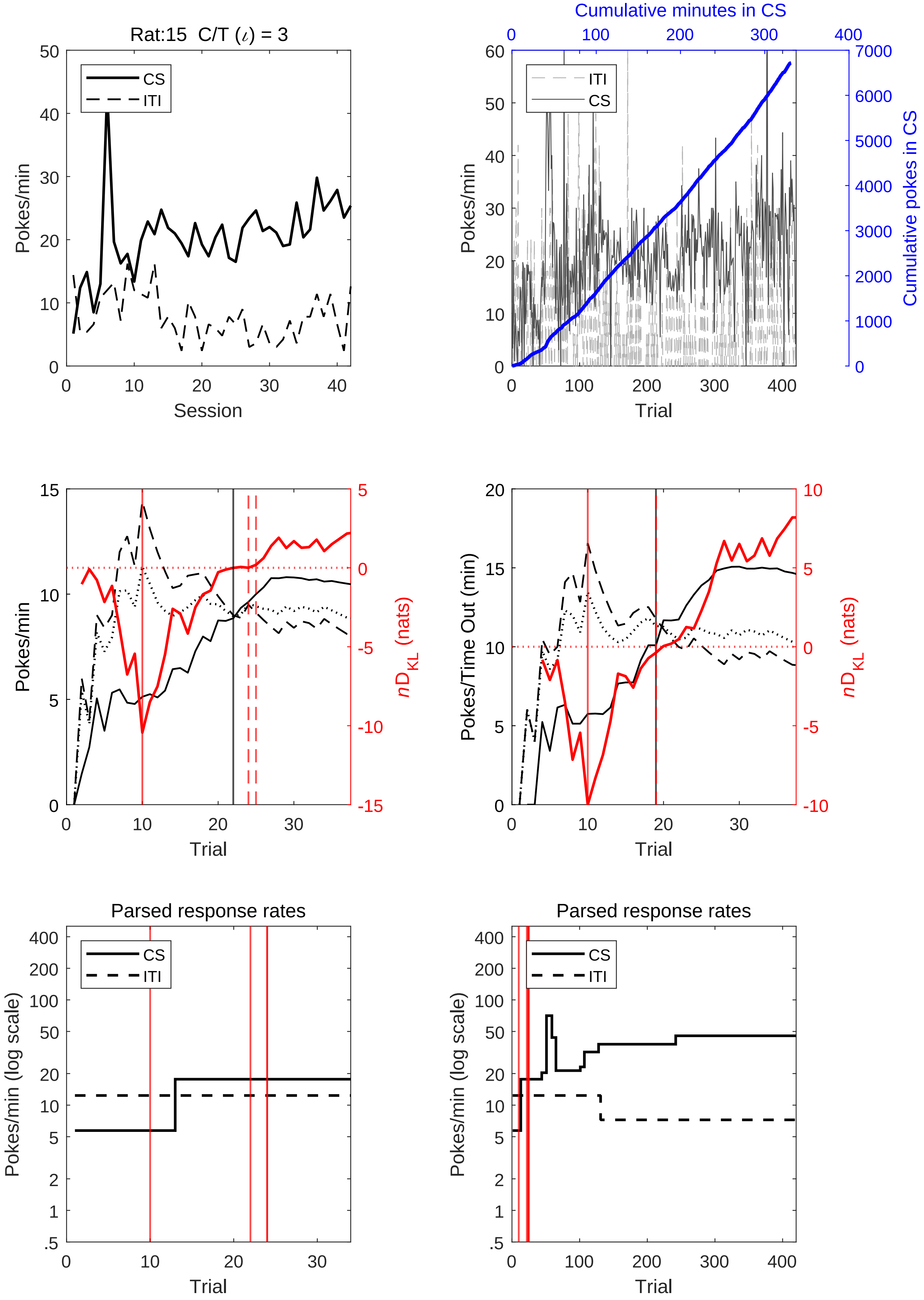

Rat:15 C/T (ι) = 3
CS
ITI
Pokes/min
Session
Cumulative minutes in CS
Cumulative pokes in CS
Trial
nD_KL (nats)
Pokes/Time Out (min)
Parsed response rates
Pokes/min (log scale)

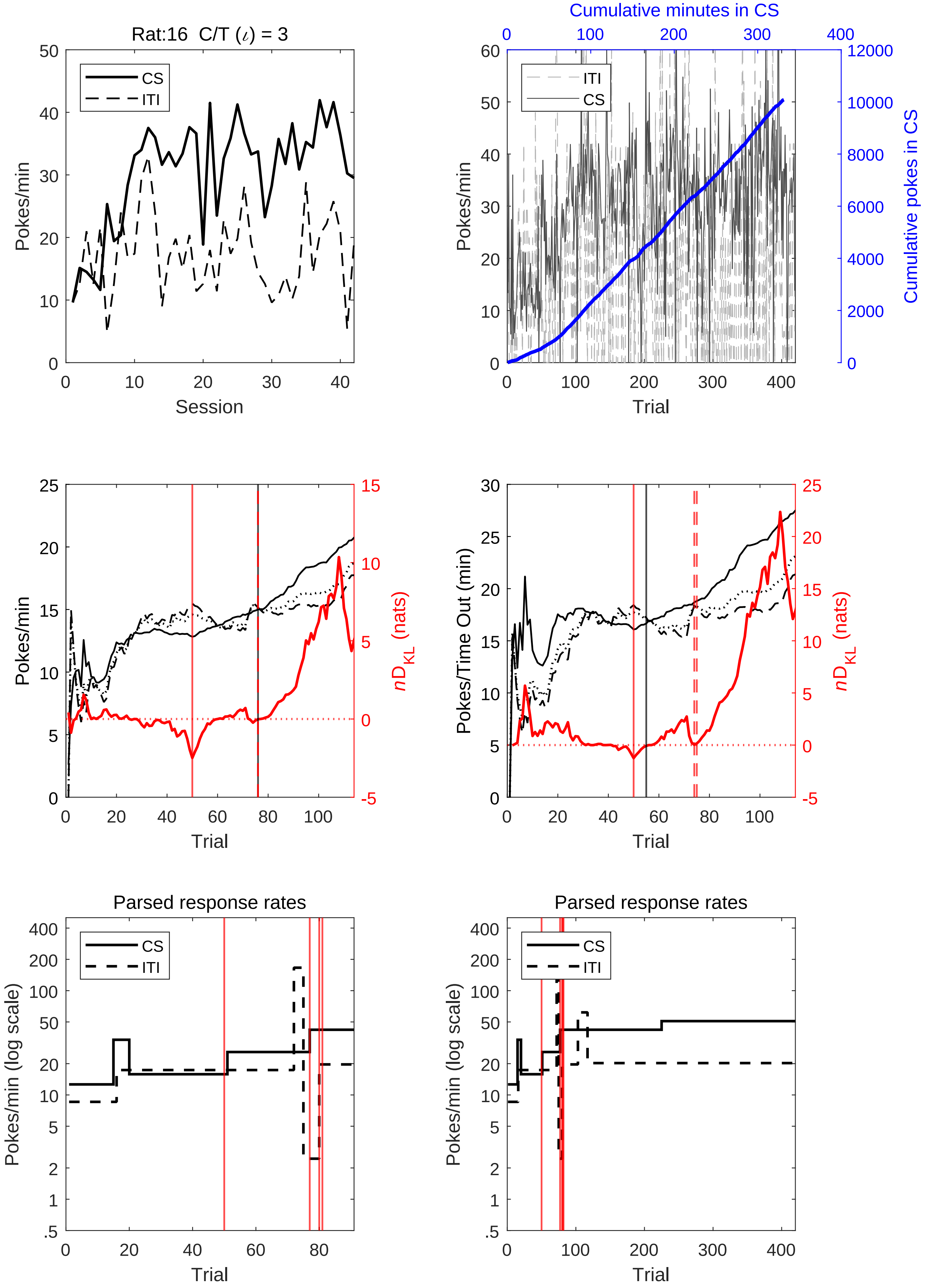
Rat:16 C/T (ι) = 3
CS
ITI
Pokes/min
Session
Cumulative minutes in CS
Cumulative pokes in CS
Trial
Pokes/Time Out (min)
nD_KL (nats)
Parsed response rates
Pokes/min (log scale)

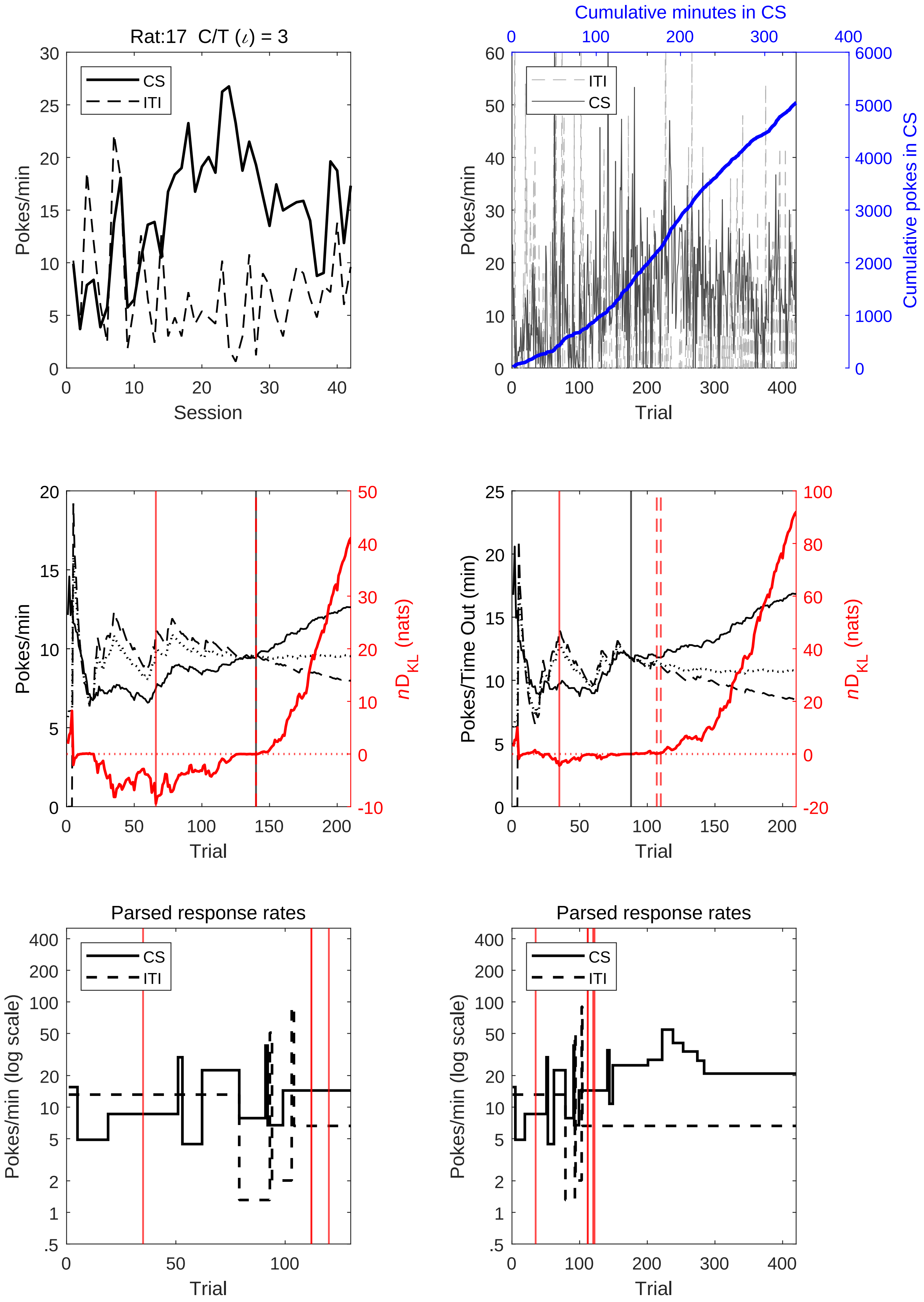
Rat:17 C/T (ι) = 3
CS
ITI
Pokes/min
Session
Cumulative minutes in CS
ITI
CS
Pokes/min
Cumulative pokes in CS
Trial
Pokes/min
nD_KL (nats)
Trial
Pokes/Time Out (min)
nD_KL (nats)
Trial
Parsed response rates
CS
ITI
Pokes/min (log scale)
Trial
Parsed response rates
CS
ITI
Pokes/min (log scale)
Trial

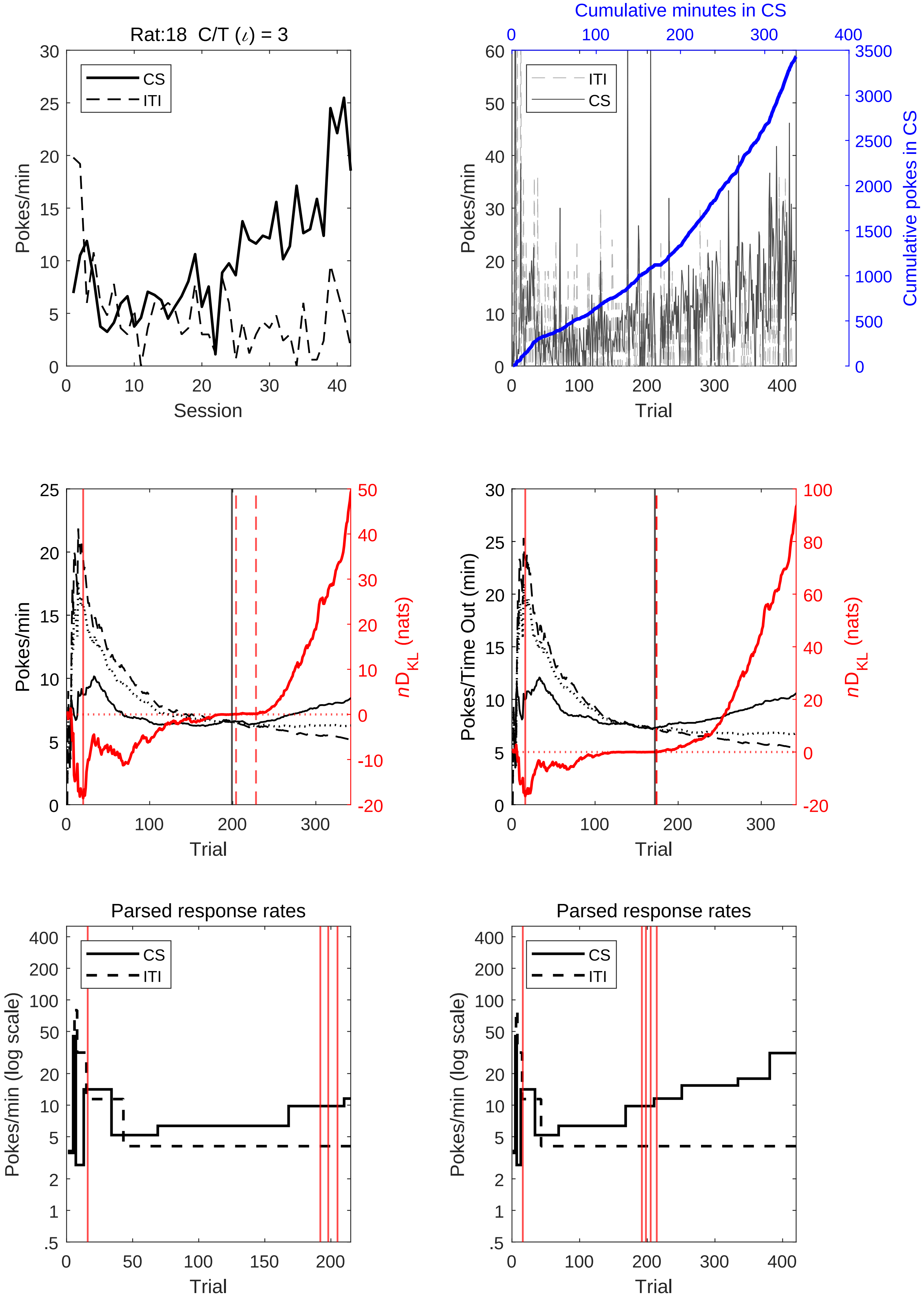
Rat:18 C/T (ι) = 3
CS
ITI
Pokes/min
Session
Cumulative minutes in CS
Cumulative pokes in CS
Trial
nD_KL (nats)
Pokes/Time Out (min)
Parsed response rates
Pokes/min (log scale)

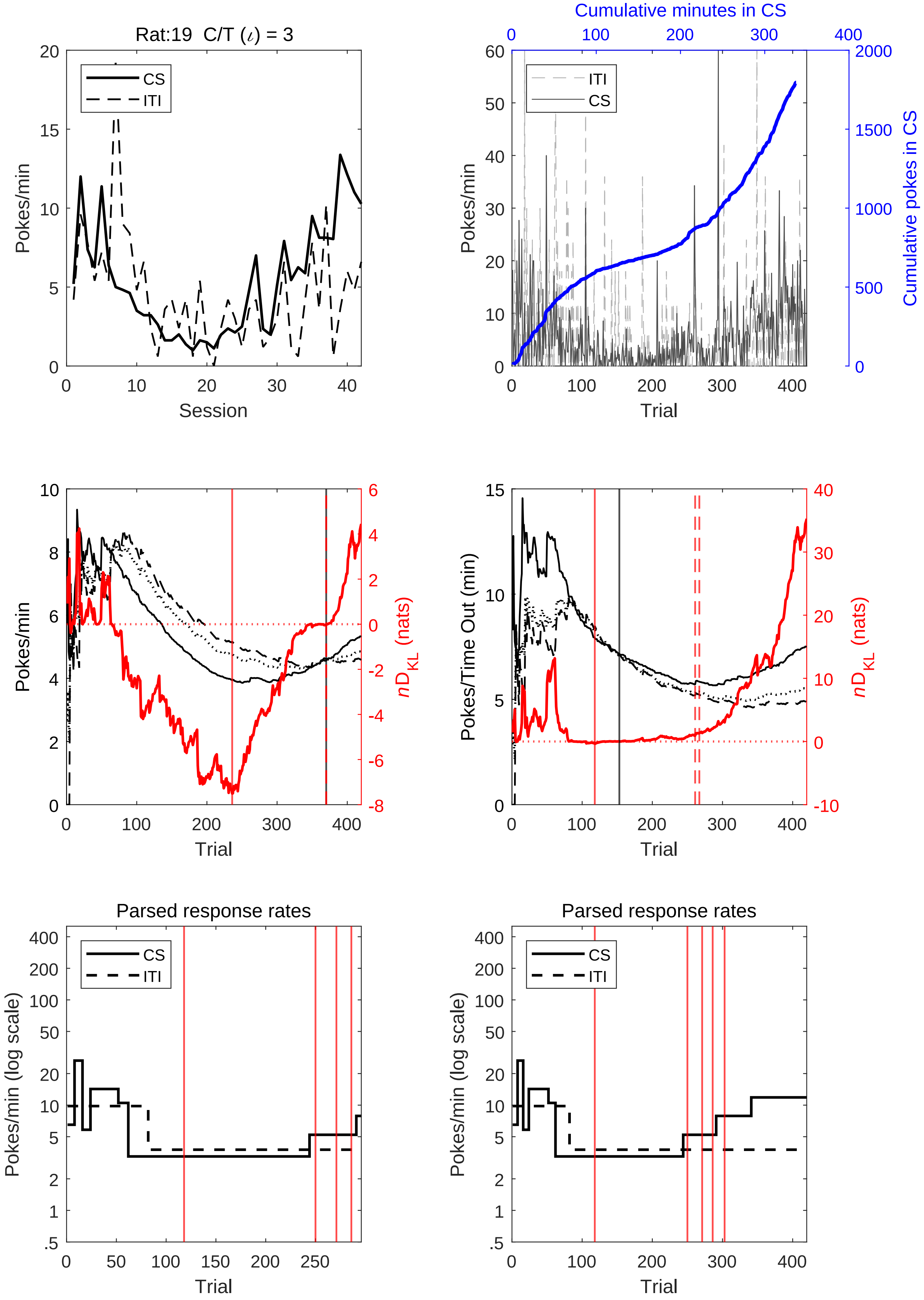

Rat:19 C/T ($\iota$) = 3
CS
ITI
Pokes/min
Session
Cumulative minutes in CS
Cumulative pokes in CS
Trial
nD_KL (nats)
Pokes/Time Out (min)
Parsed response rates
Pokes/min (log scale)

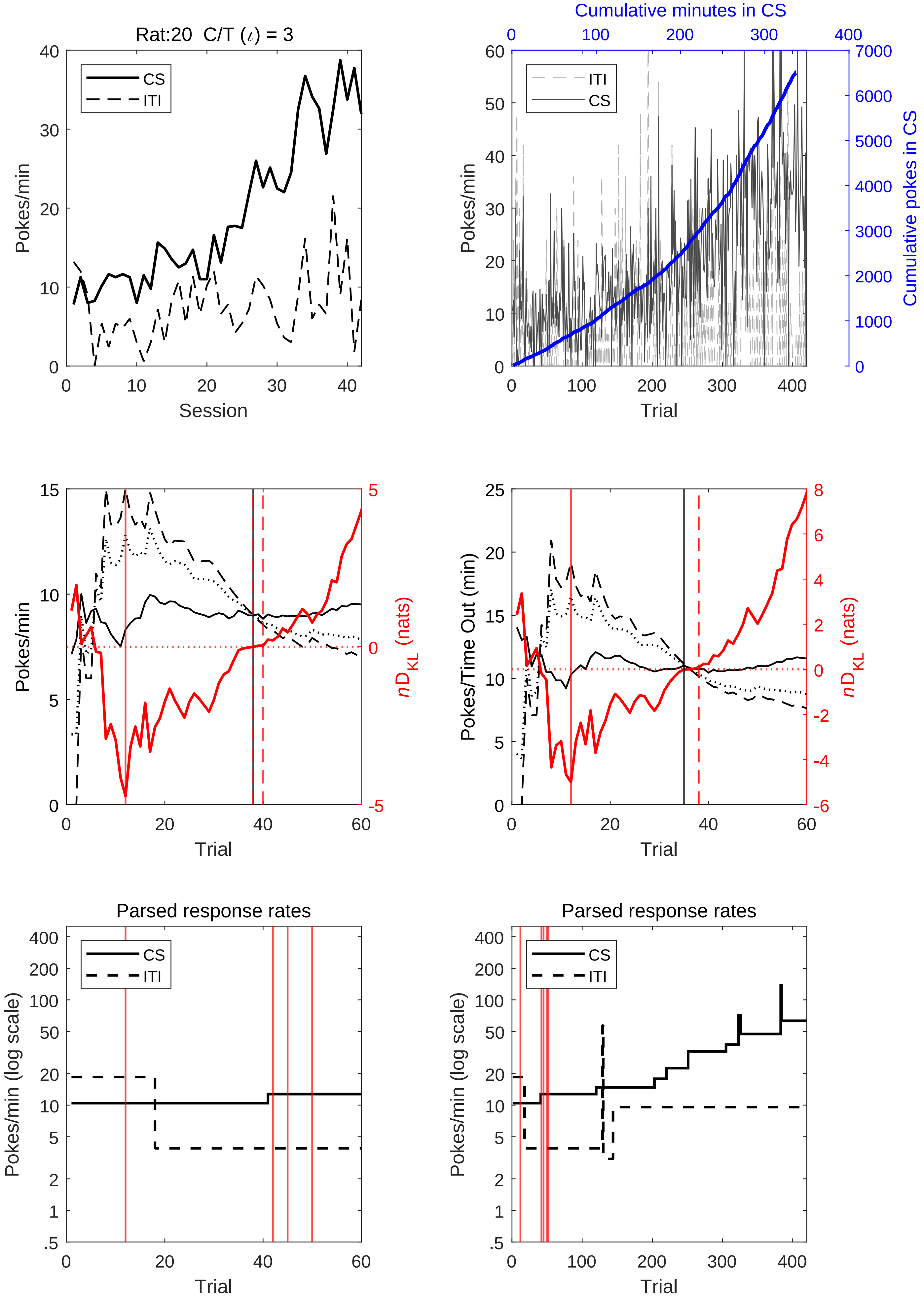

Rat:20 C/T (ι) = 3
CS
ITI
Pokes/min
Session
Cumulative minutes in CS
Cumulative pokes in CS
Trial
Pokes/Time Out (min)
nD_KL (nats)
Parsed response rates
Pokes/min (log scale)

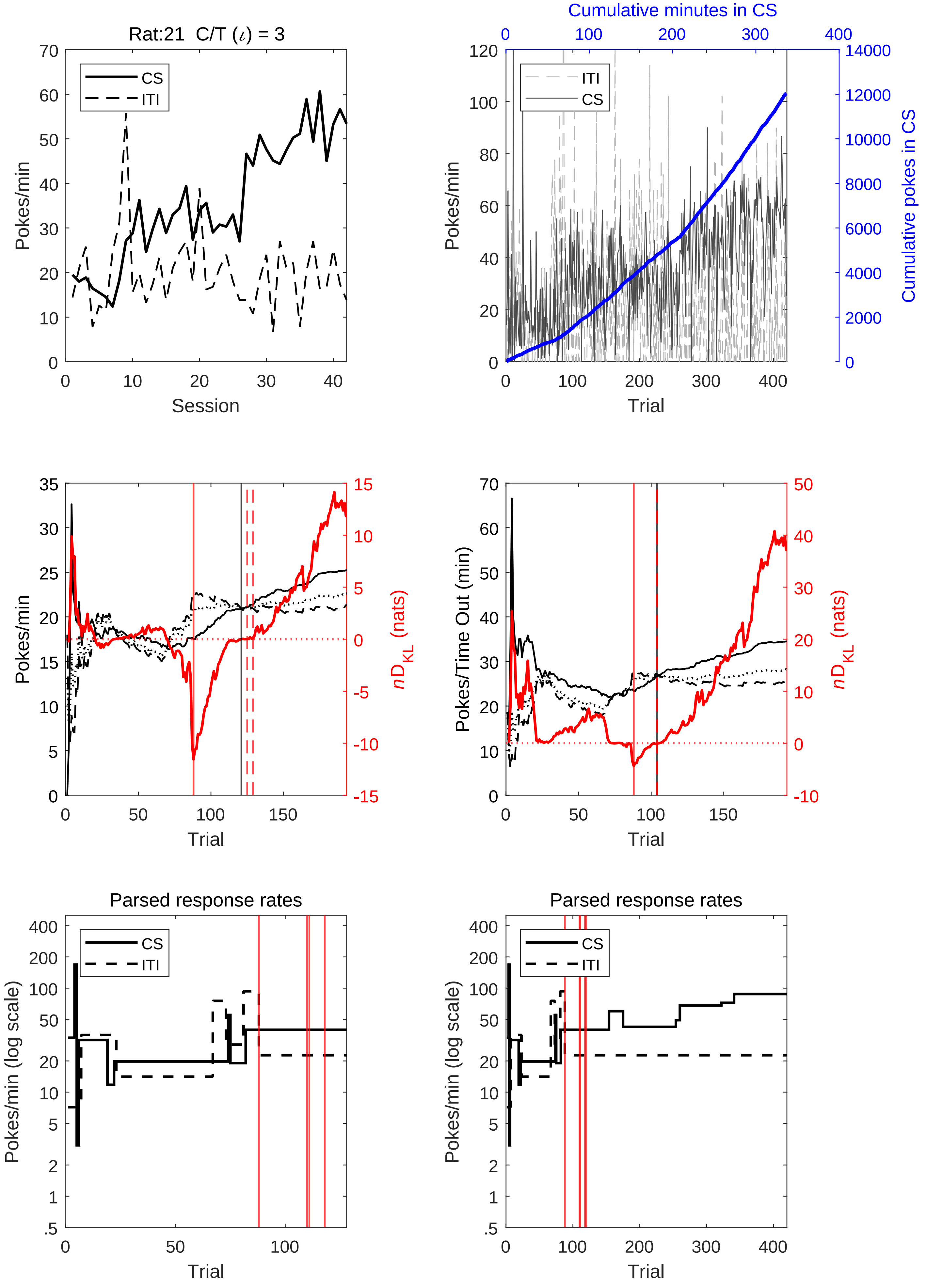
Rat:21 C/T (ι) = 3
CS
ITI
Pokes/min
Session
Cumulative minutes in CS
Cumulative pokes in CS
Trial
nD_KL (nats)
Pokes/Time Out (min)
Parsed response rates
Pokes/min (log scale)

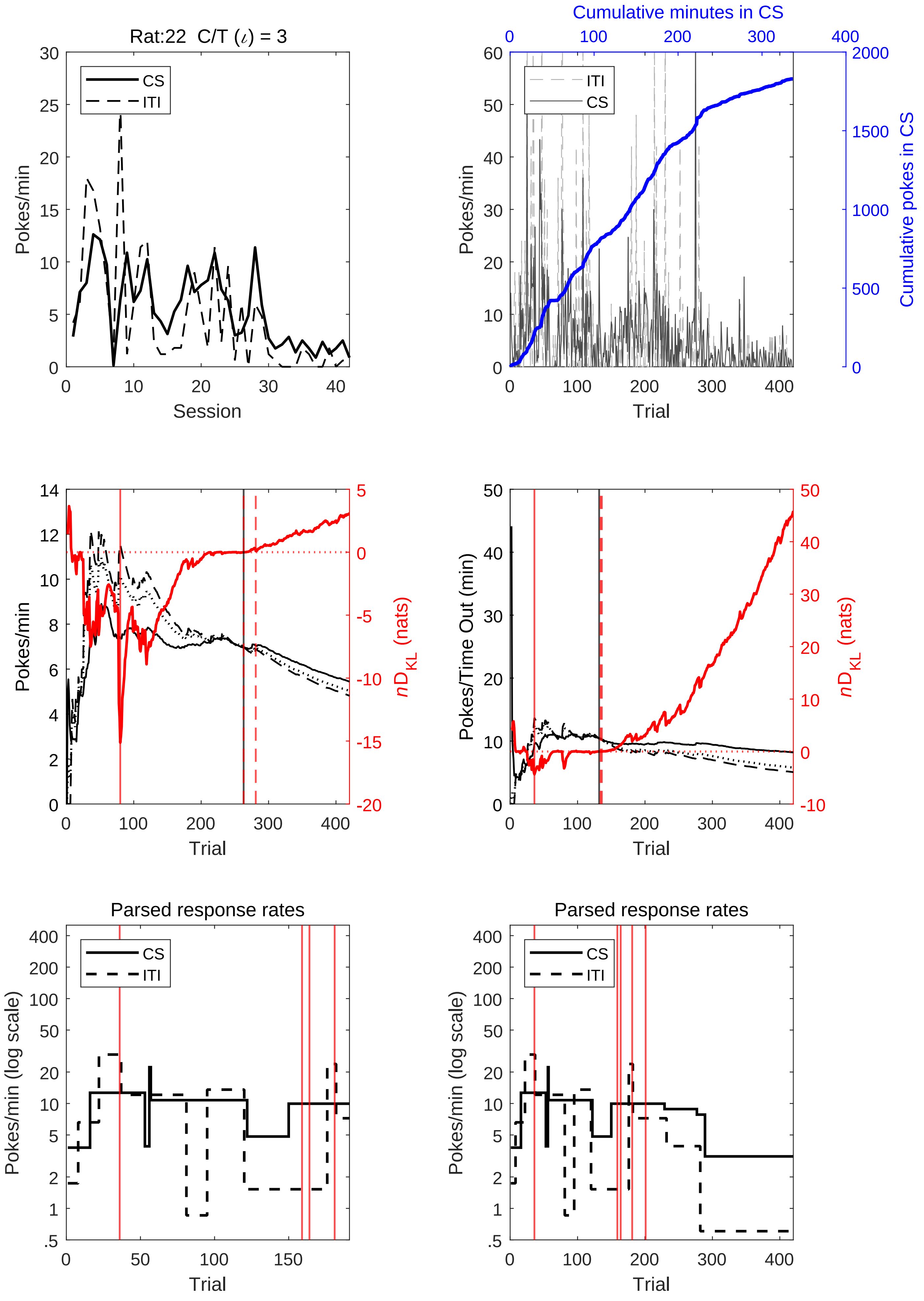

Rat:22 C/T (ι) = 3
CS
ITI
Pokes/min
Session
Cumulative minutes in CS
Cumulative pokes in CS
Trial
$nD_{KL}$ (nats)
Pokes/Time Out (min)
Parsed response rates
Pokes/min (log scale)

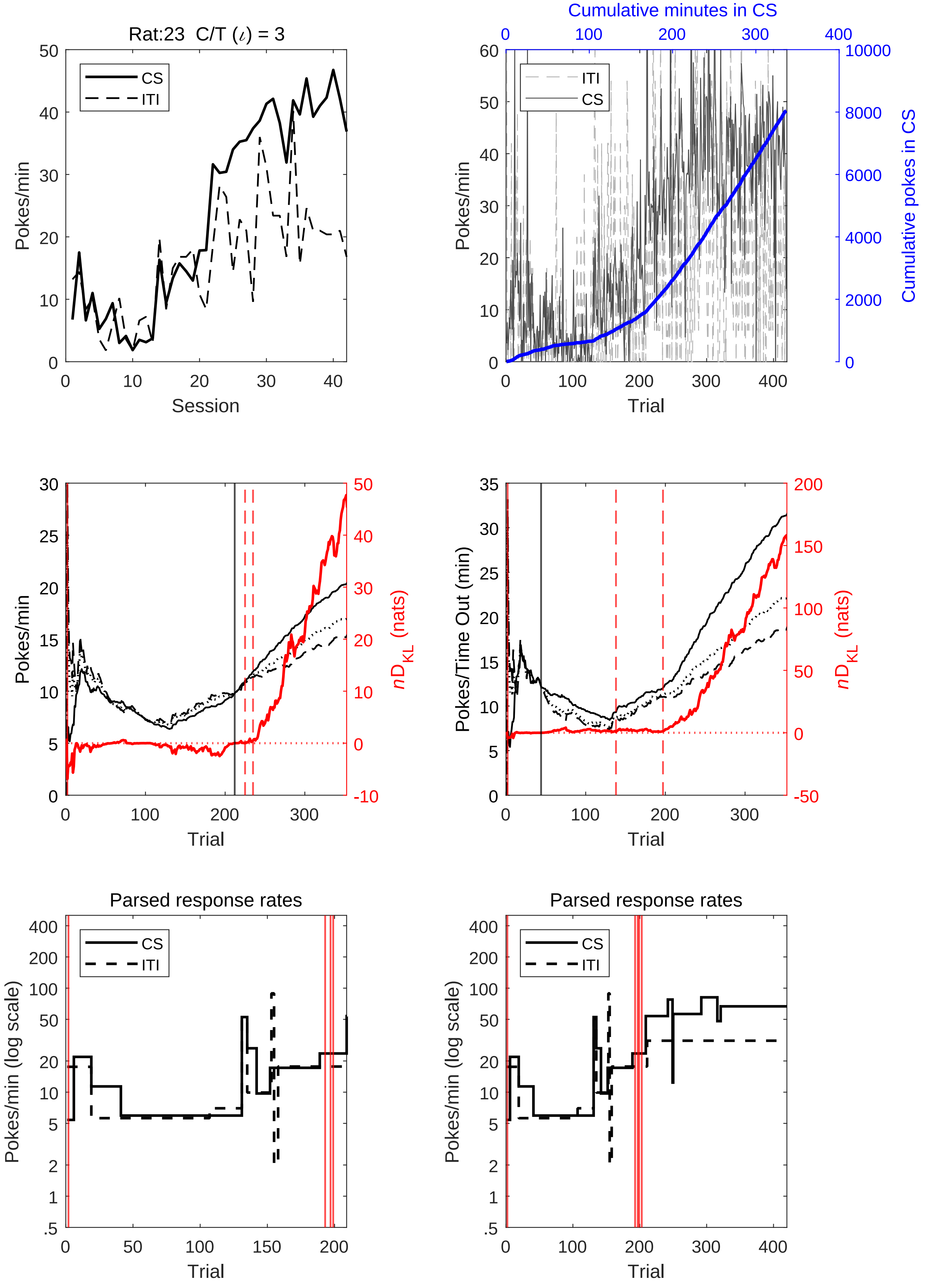
Rat:23 C/T (ι) = 3
CS
ITI
Pokes/min
Session
Cumulative minutes in CS
Cumulative pokes in CS
ITI
CS
Pokes/min
Trial
Pokes/min
nD_KL (nats)
Trial
Pokes/Time Out (min)
nD_KL (nats)
Trial
Parsed response rates
CS
ITI
Pokes/min (log scale)
Trial
Parsed response rates
CS
ITI
Pokes/min (log scale)
Trial

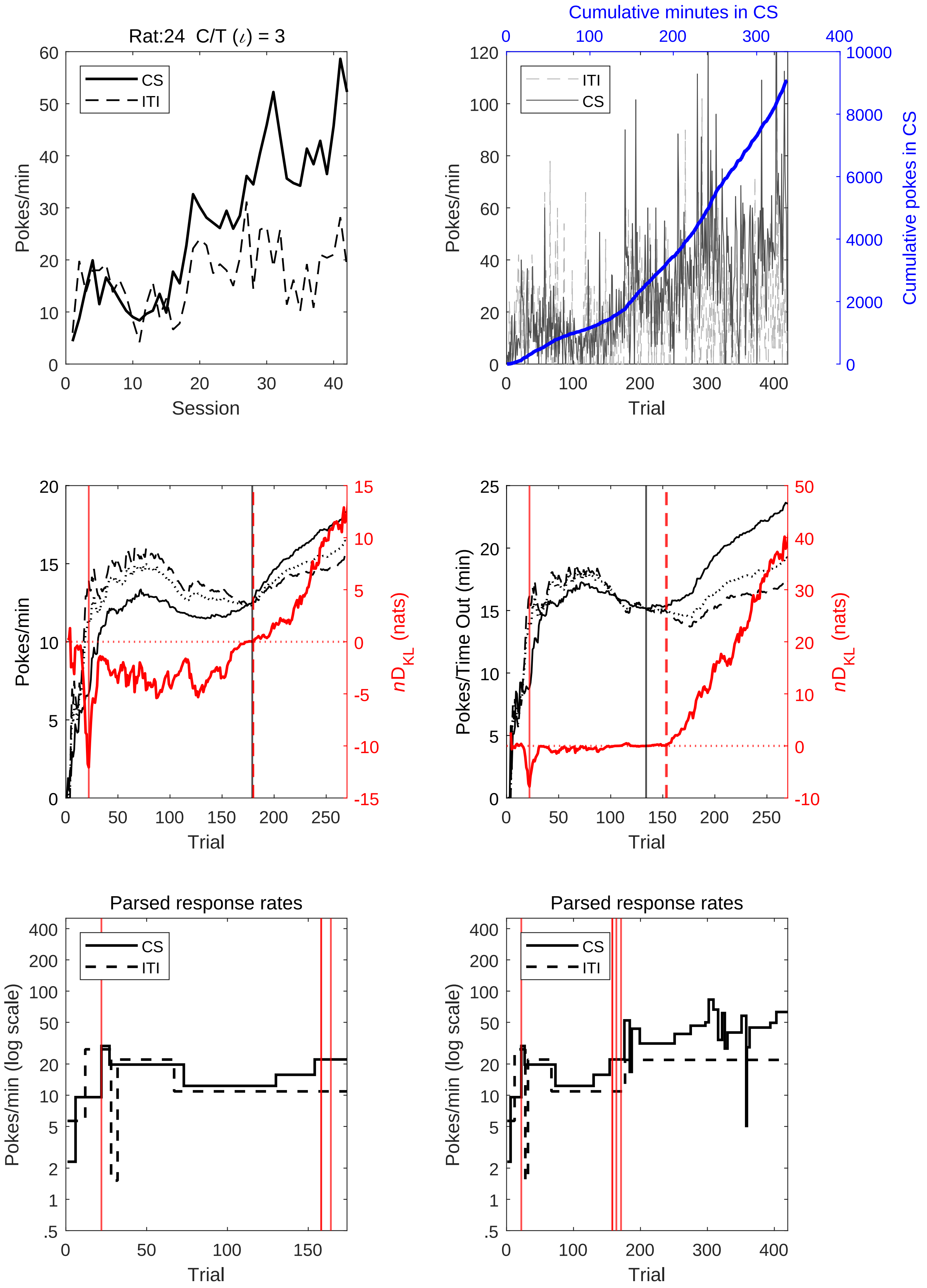
Rat:24 C/T (ι) = 3
CS
ITI
Pokes/min
Session
Cumulative minutes in CS
Cumulative pokes in CS
Trial
nD_KL (nats)
Pokes/Time Out (min)
Parsed response rates
Pokes/min (log scale)

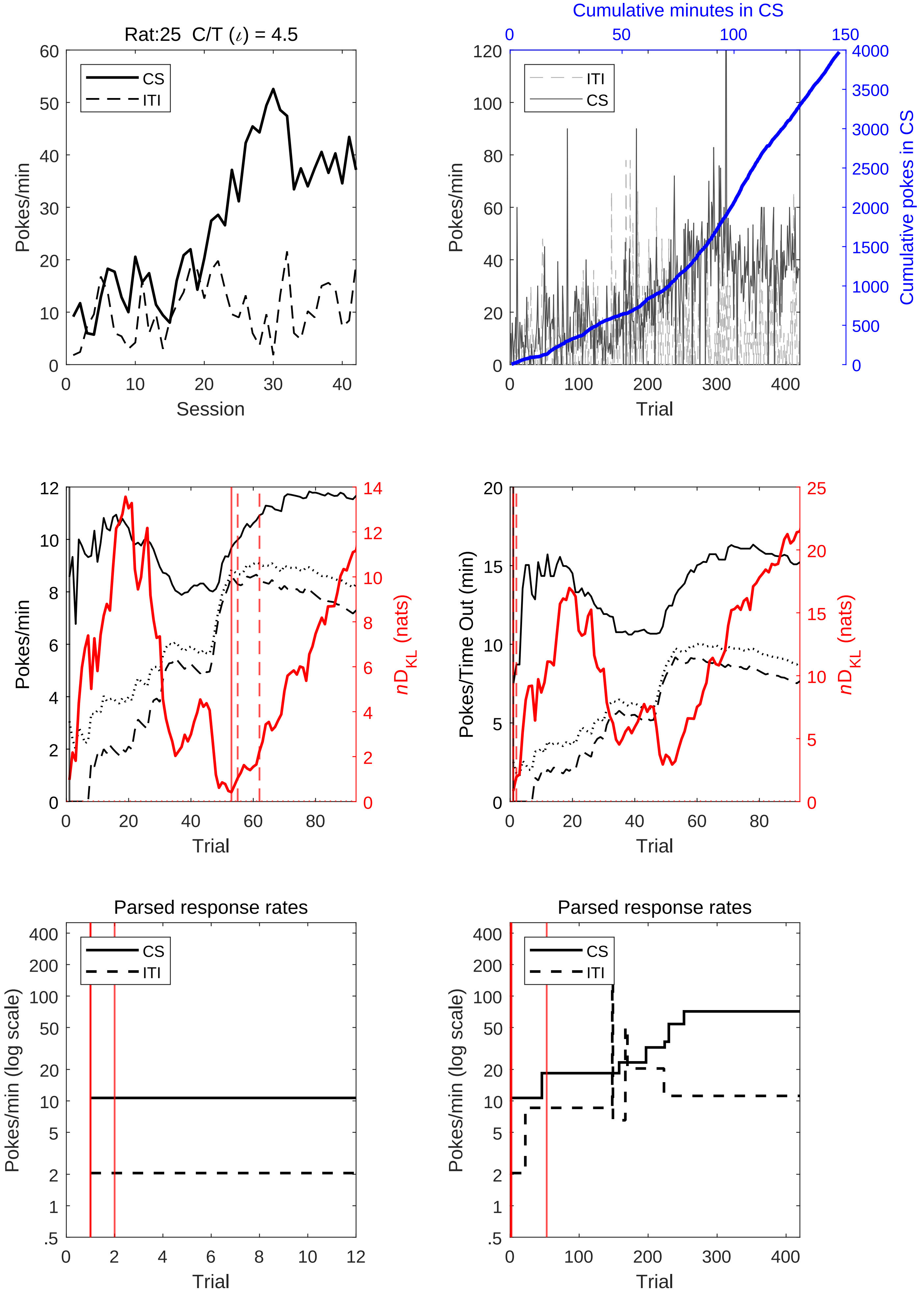

Rat:25 C/T ($\iota$) = 4.5
CS
ITI
Pokes/min
Session
Cumulative minutes in CS
Cumulative pokes in CS
Trial
Pokes/Time Out (min)
$n$D$_{KL}$ (nats)
Parsed response rates
Pokes/min (log scale)

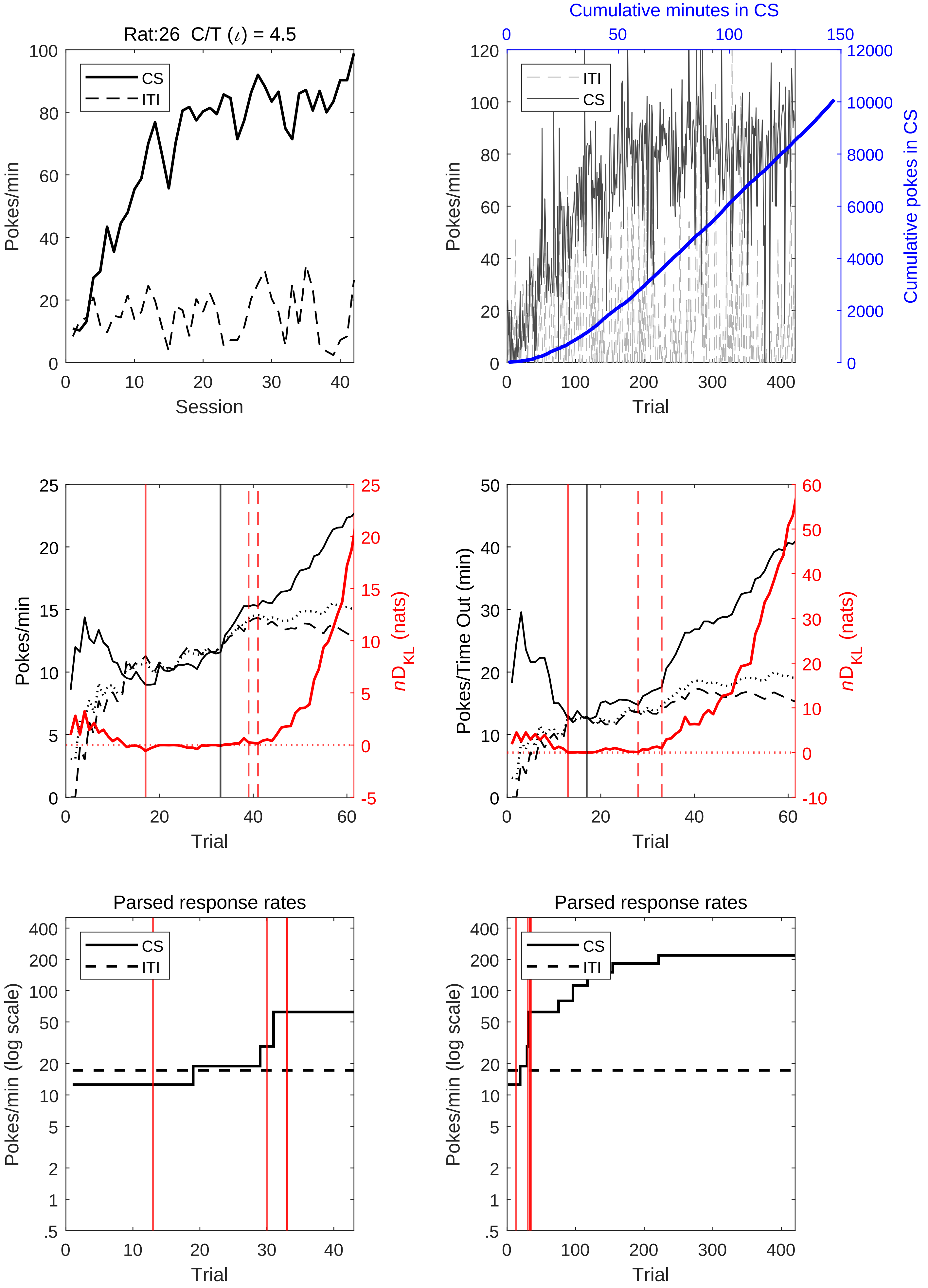
Rat:26 C/T (ι) = 4.5
CS
ITI
Pokes/min
Session
Cumulative minutes in CS
Cumulative pokes in CS
Trial
nD_KL (nats)
Pokes/Time Out (min)
Parsed response rates
Pokes/min (log scale)

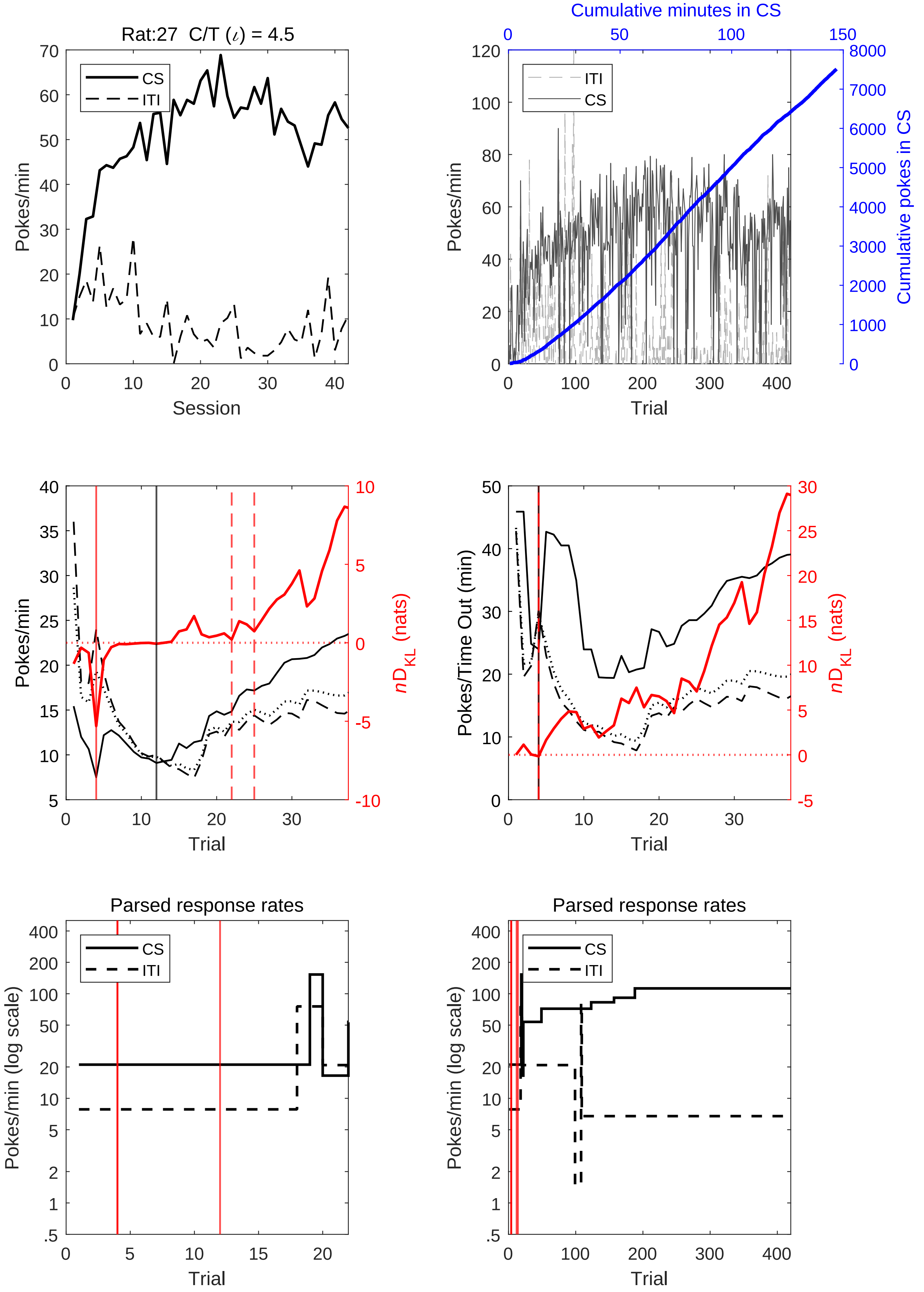

Rat:27 C/T (ι) = 4.5
CS
ITI
Pokes/min
Session
Cumulative minutes in CS
Cumulative pokes in CS
Trial
nD_KL (nats)
Pokes/Time Out (min)
Parsed response rates
Pokes/min (log scale)

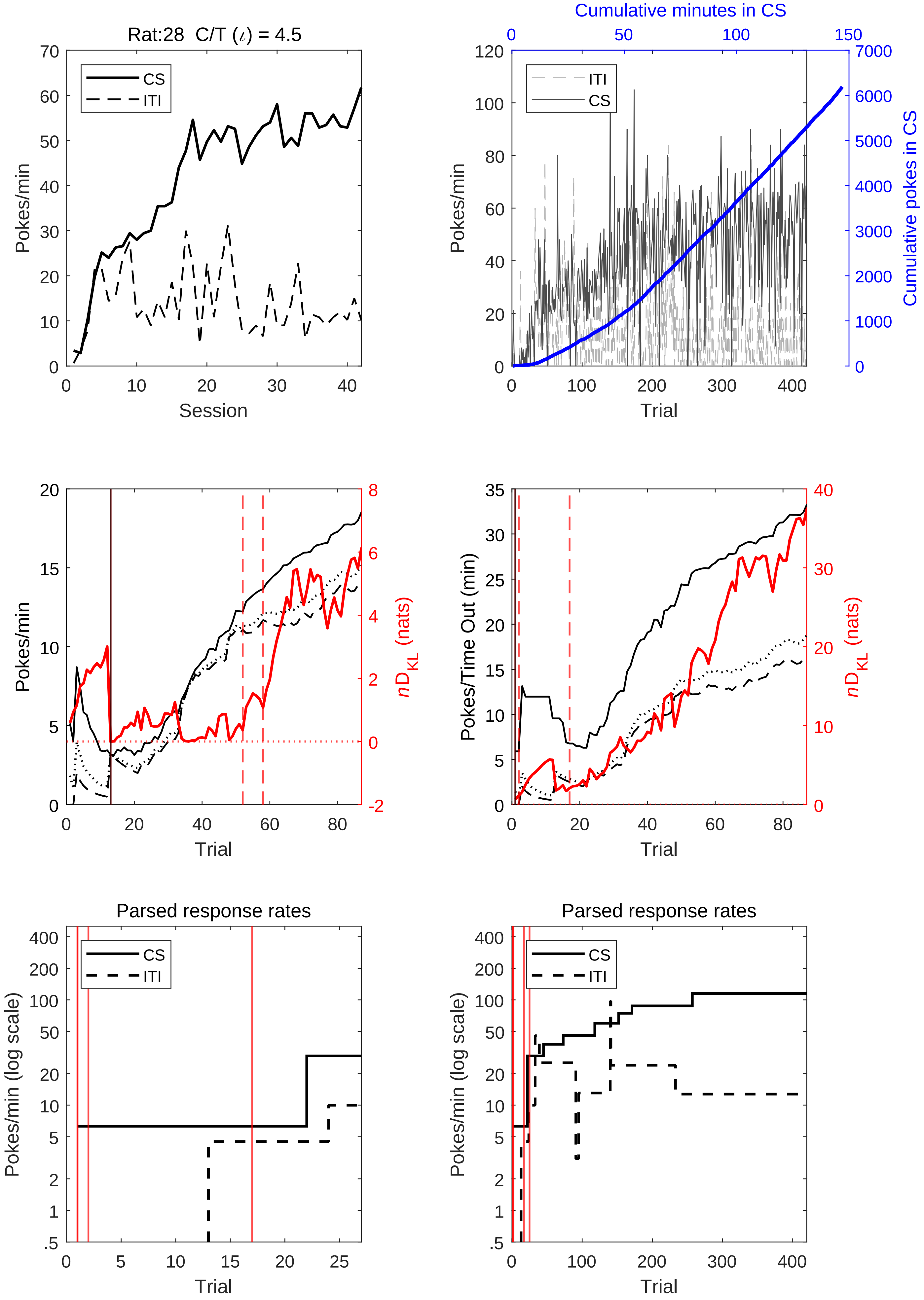
Rat:28 C/T (ι) = 4.5
CS
ITI
Pokes/min
Session
Cumulative minutes in CS
Cumulative pokes in CS
Trial
Pokes/min
nD_KL (nats)
Pokes/Time Out (min)
Parsed response rates
Pokes/min (log scale)

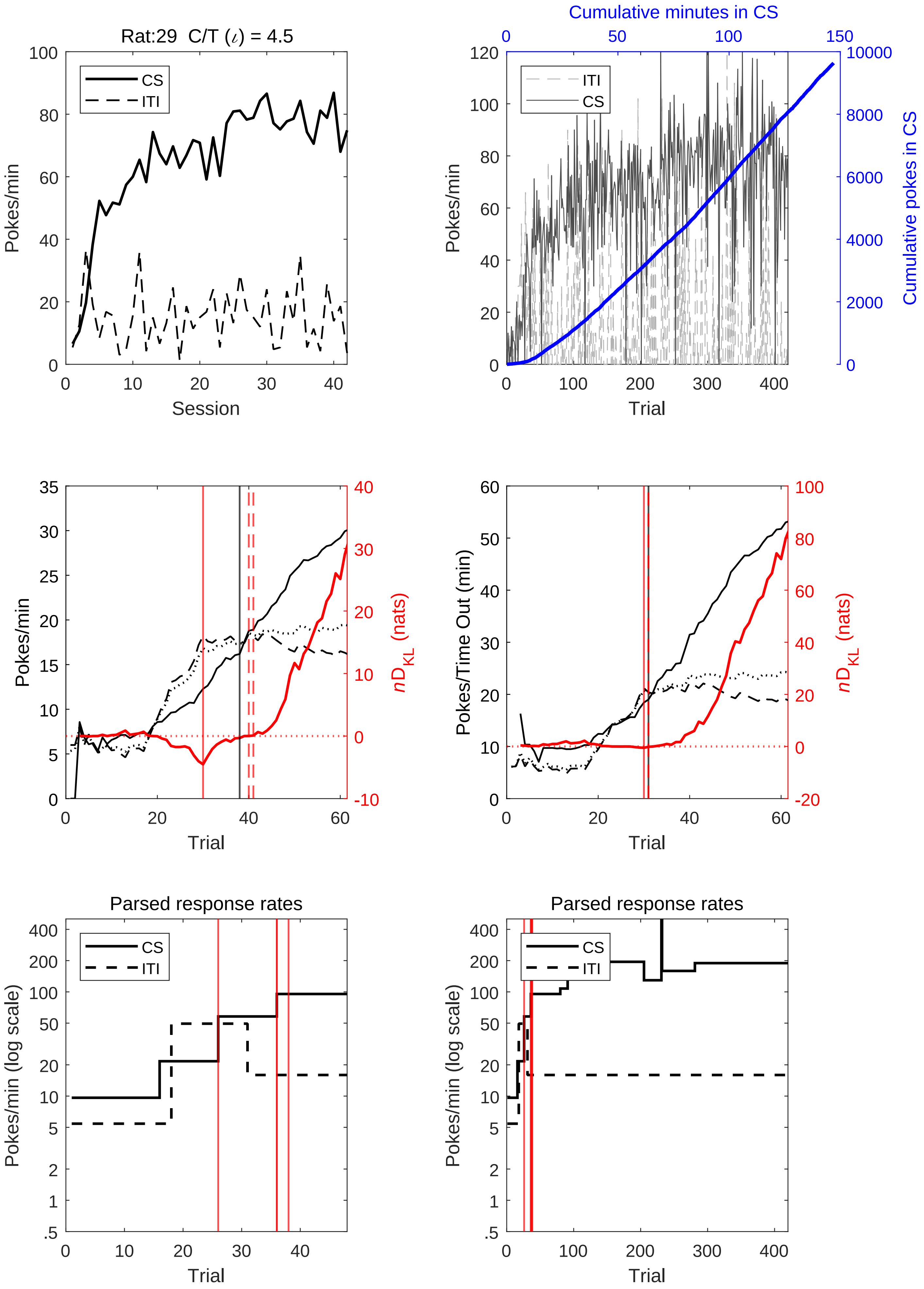
Rat:29 C/T (ι) = 4.5
CS
ITI
Pokes/min
Session
Cumulative minutes in CS
ITI
CS
Pokes/min
Cumulative pokes in CS
Trial
Pokes/min
$nD_{KL}$ (nats)
Trial
Pokes/Time Out (min)
$nD_{KL}$ (nats)
Trial
Parsed response rates
CS
ITI
Pokes/min (log scale)
Trial
Parsed response rates
CS
ITI
Pokes/min (log scale)
Trial

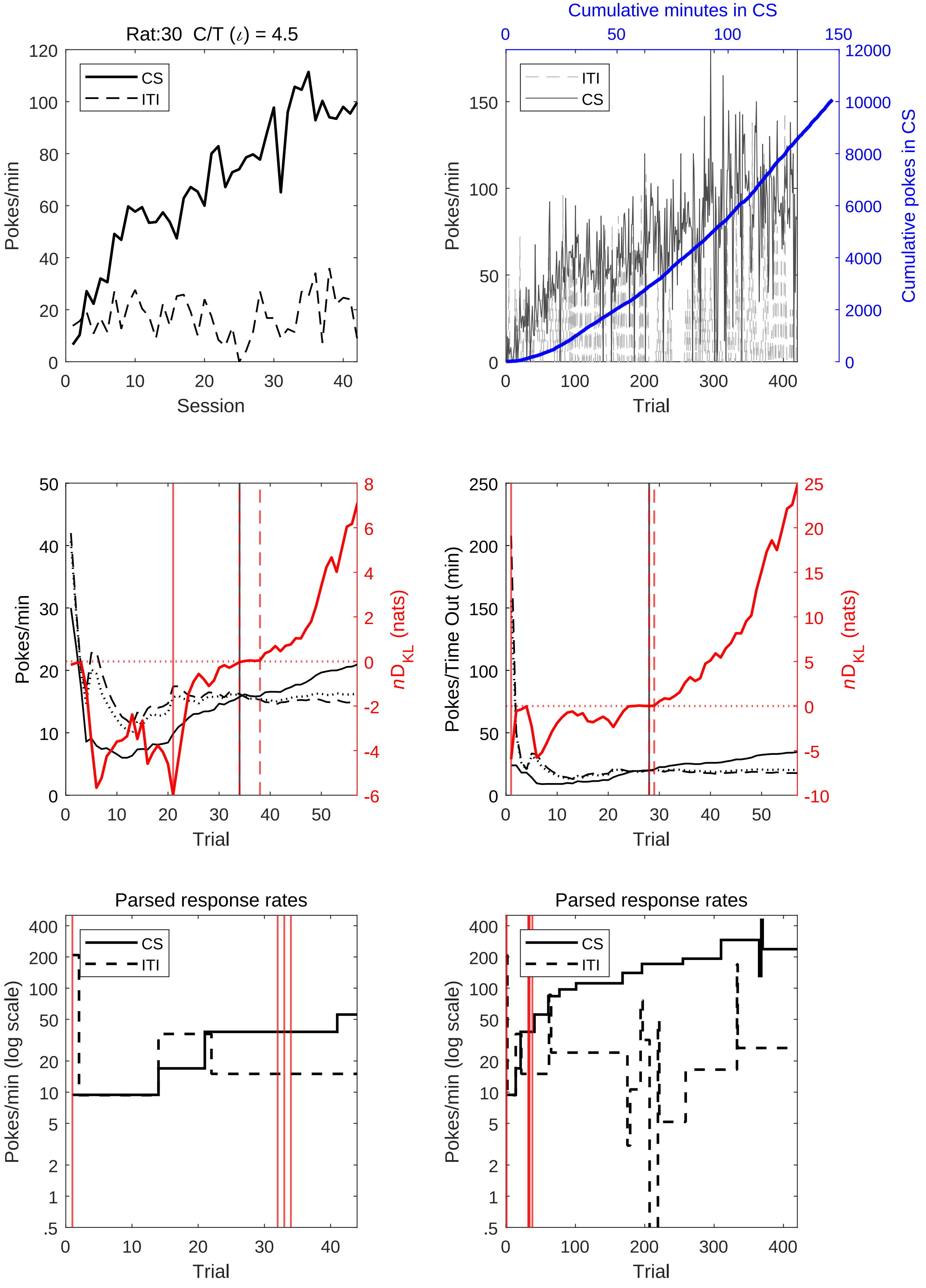
Rat:30 C/T (ι) = 4.5
CS
ITI
Pokes/min
Session
Cumulative minutes in CS
Cumulative pokes in CS
Trial
nD_KL (nats)
Pokes/Time Out (min)
Parsed response rates
Pokes/min (log scale)

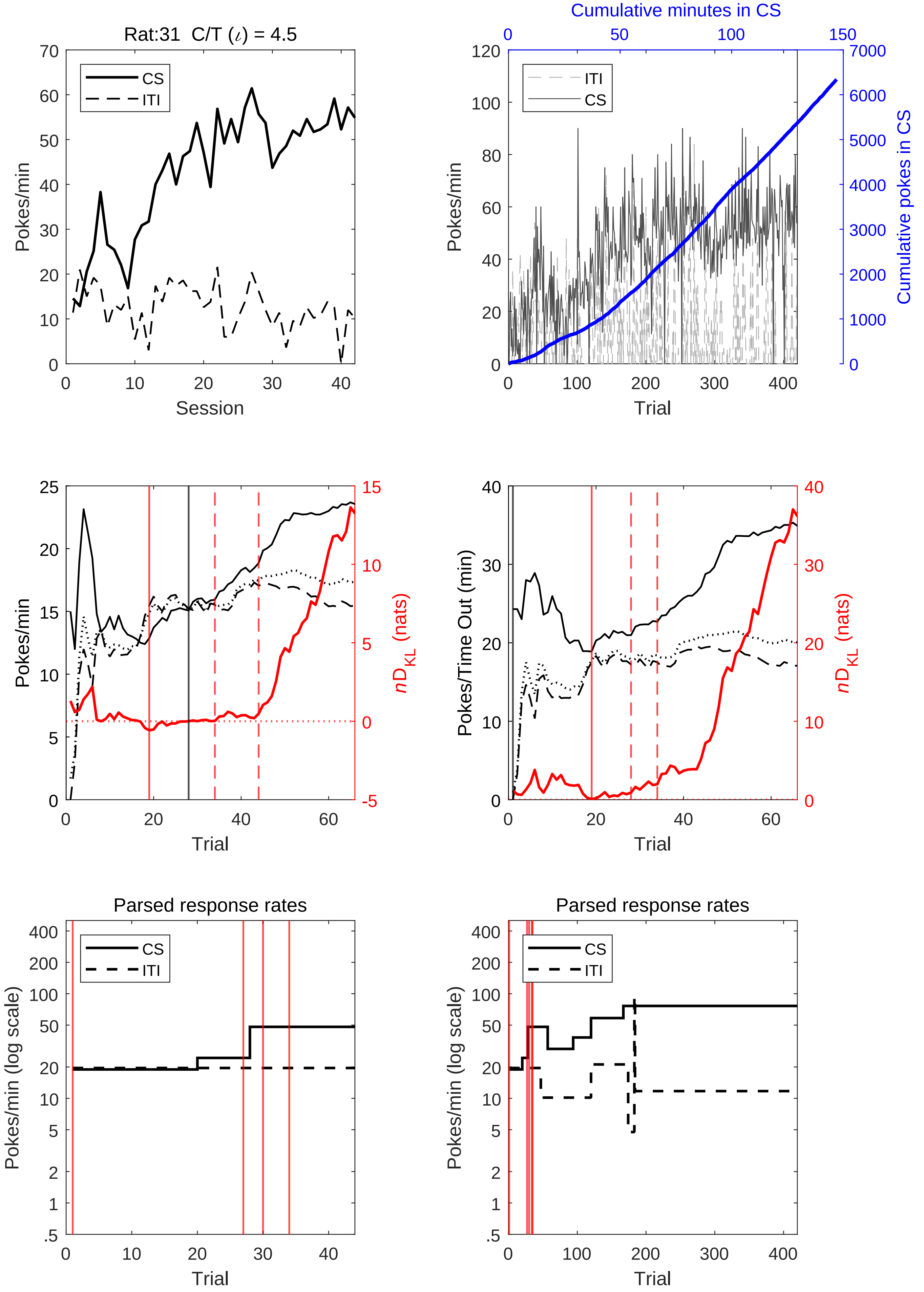

Rat:31 C/T (ι) = 4.5
CS
ITI
Pokes/min
Session
Cumulative minutes in CS
Cumulative pokes in CS
Trial
Pokes/Time Out (min)
nD_KL (nats)
Parsed response rates
Pokes/min (log scale)

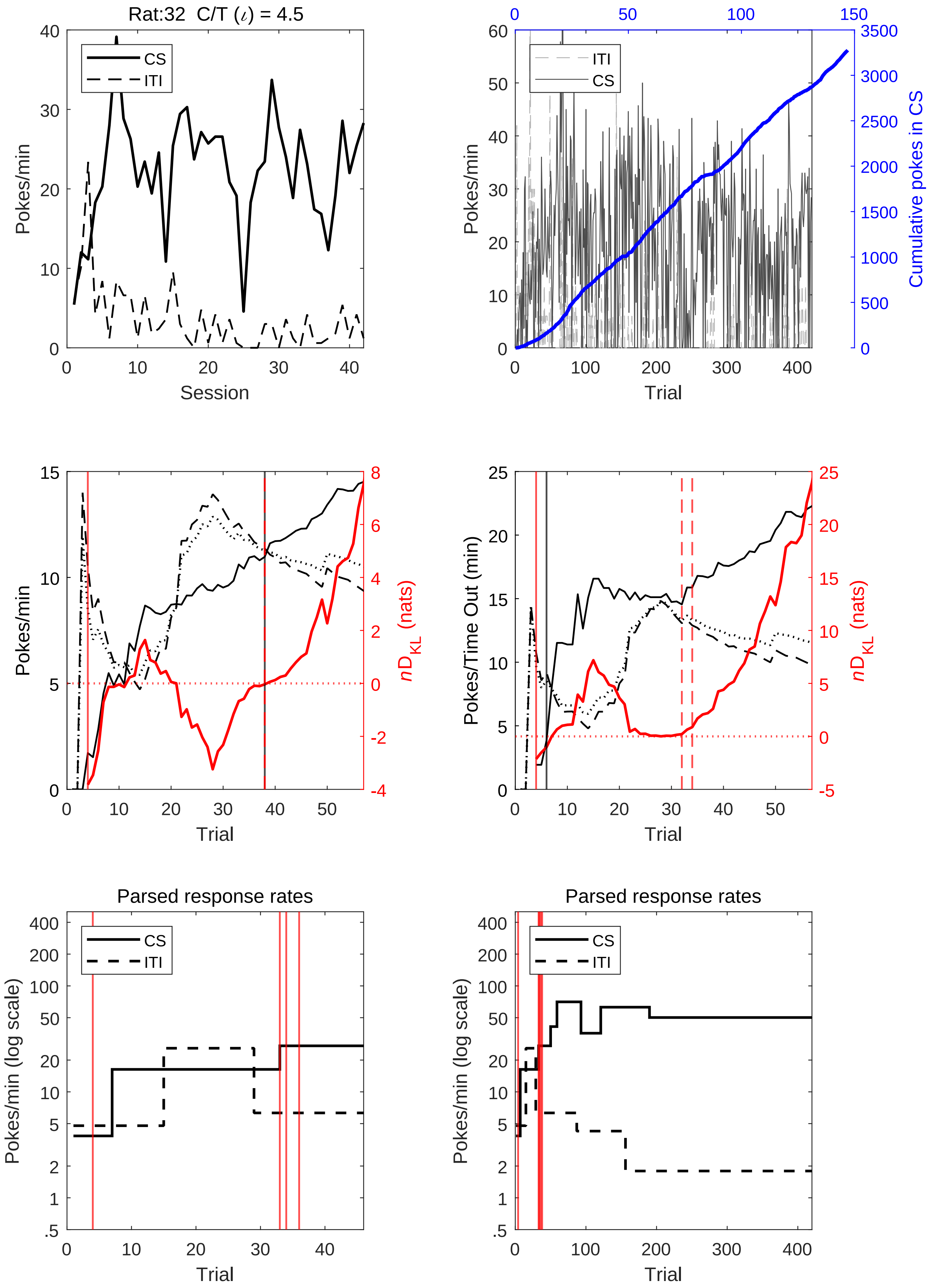

Rat:32 C/T (ι) = 4.5
CS
ITI
Pokes/min
Session
Cumulative minutes in CS
Cumulative pokes in CS
Trial
nD_KL (nats)
Pokes/Time Out (min)
Parsed response rates
Pokes/min (log scale)

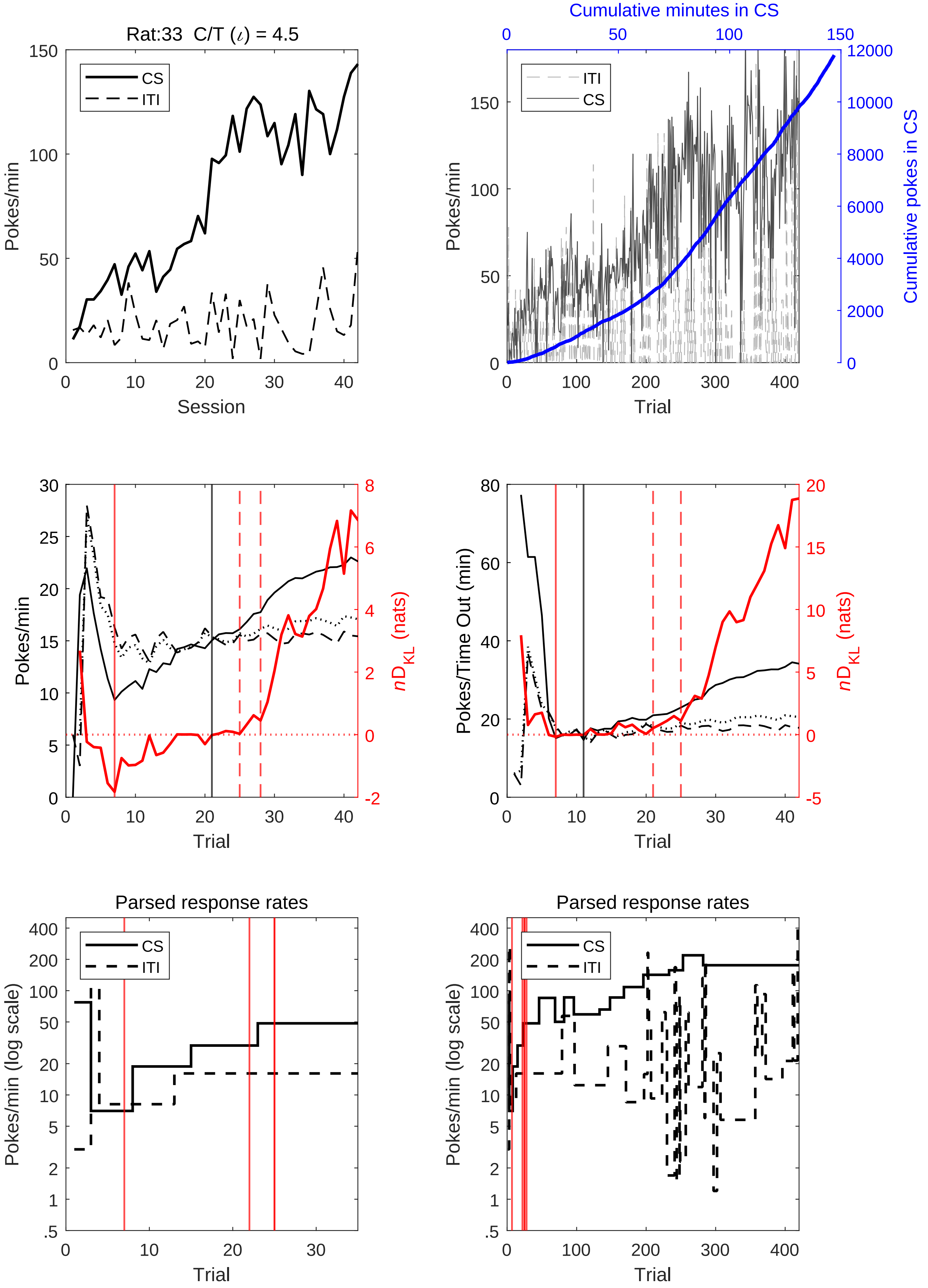
Rat:33  C/T (ι) = 4.5
CS
ITI
Pokes/min
Session
Cumulative minutes in CS
Cumulative pokes in CS
Trial
nD_KL (nats)
Pokes/Time Out (min)
Parsed response rates
Pokes/min (log scale)

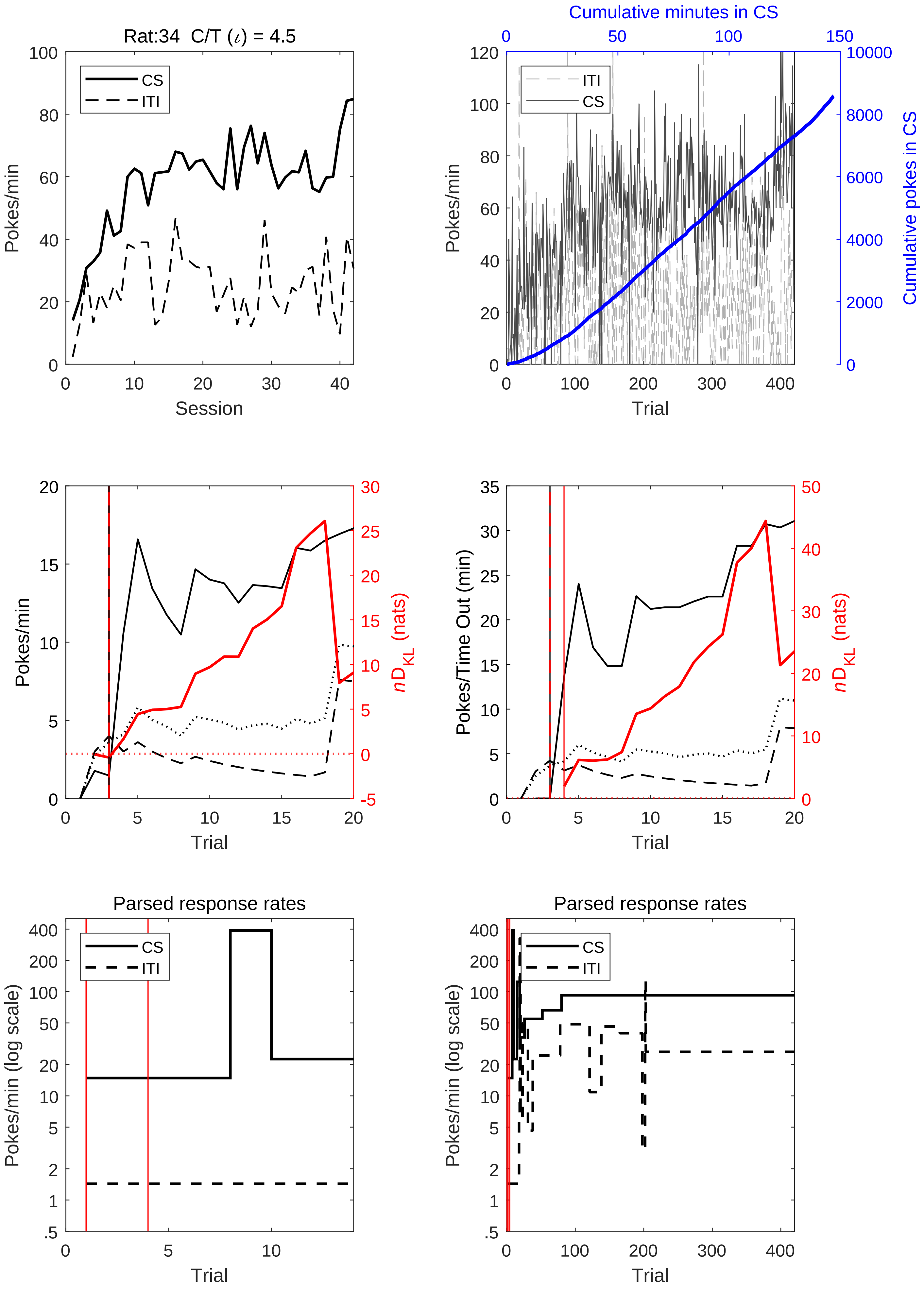
Rat:34 C/T (ι) = 4.5
CS
ITI
Pokes/min
Session
Cumulative minutes in CS
ITI
CS
Pokes/min
Trial
Cumulative pokes in CS
Pokes/min
Trial
$nD_{KL}$ (nats)
Pokes/Time Out (min)
Trial
$nD_{KL}$ (nats)
Parsed response rates
CS
ITI
Pokes/min (log scale)
Trial
Parsed response rates
CS
ITI
Pokes/min (log scale)
Trial

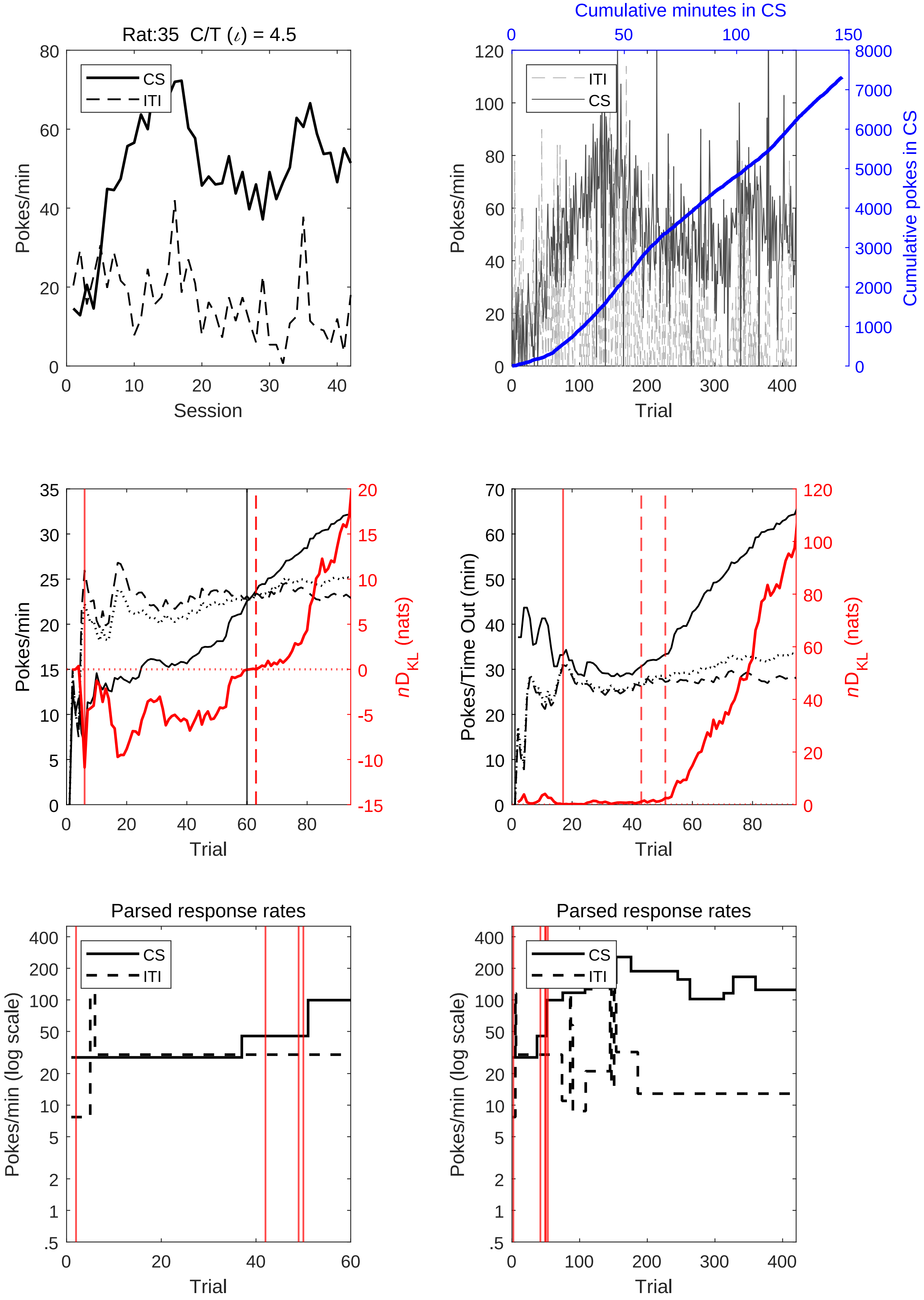

Rat:35 C/T (ι) = 4.5
CS
ITI
Pokes/min
Session
Cumulative minutes in CS
Cumulative pokes in CS
Trial
Pokes/min
nD_KL (nats)
Pokes/Time Out (min)
Parsed response rates
Pokes/min (log scale)

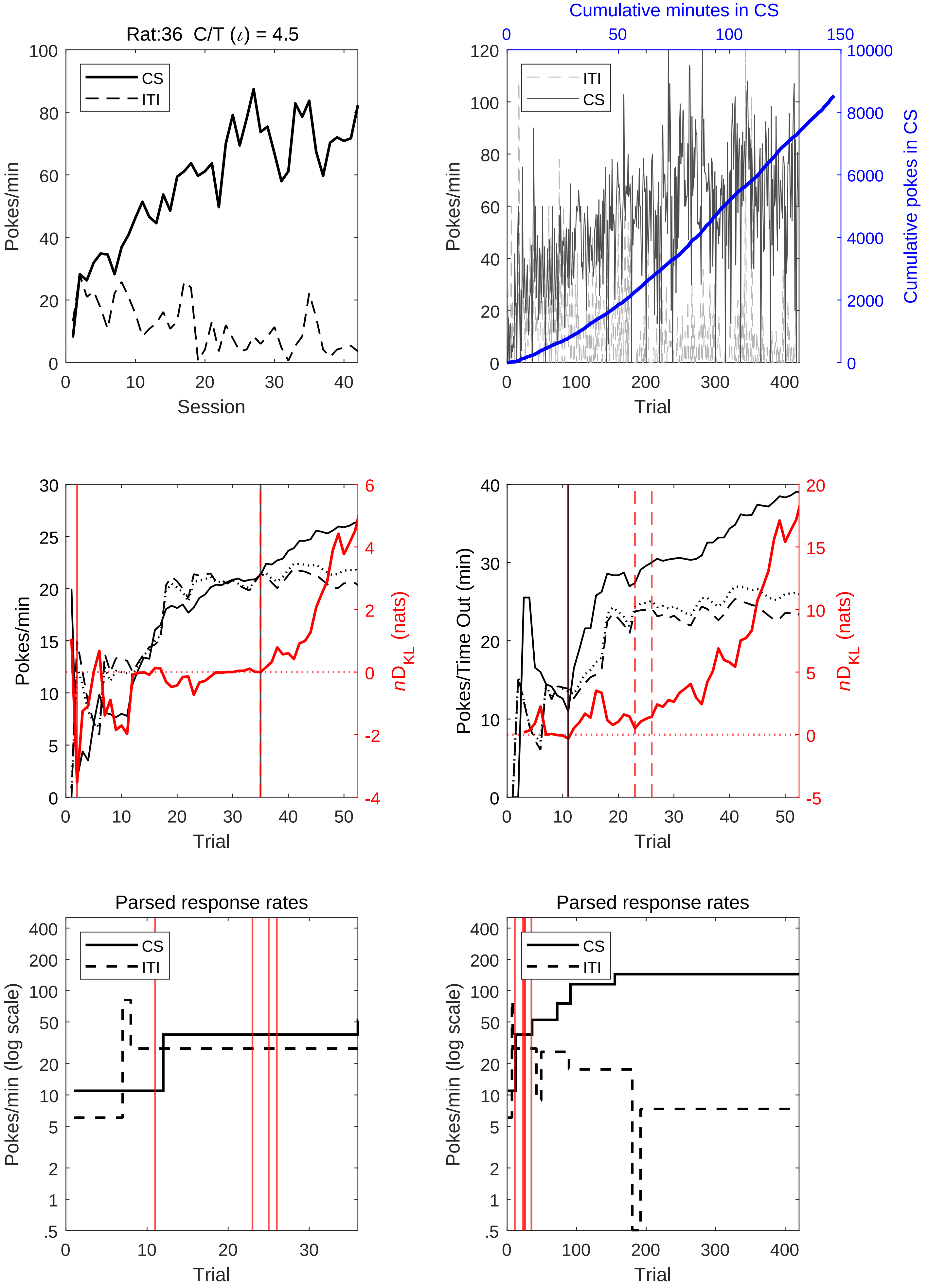

Rat:36 C/T (ι) = 4.5
CS
ITI
Pokes/min
Session
Cumulative minutes in CS
Cumulative pokes in CS
Trial
Pokes/Time Out (min)
nD_KL (nats)
Parsed response rates
Pokes/min (log scale)

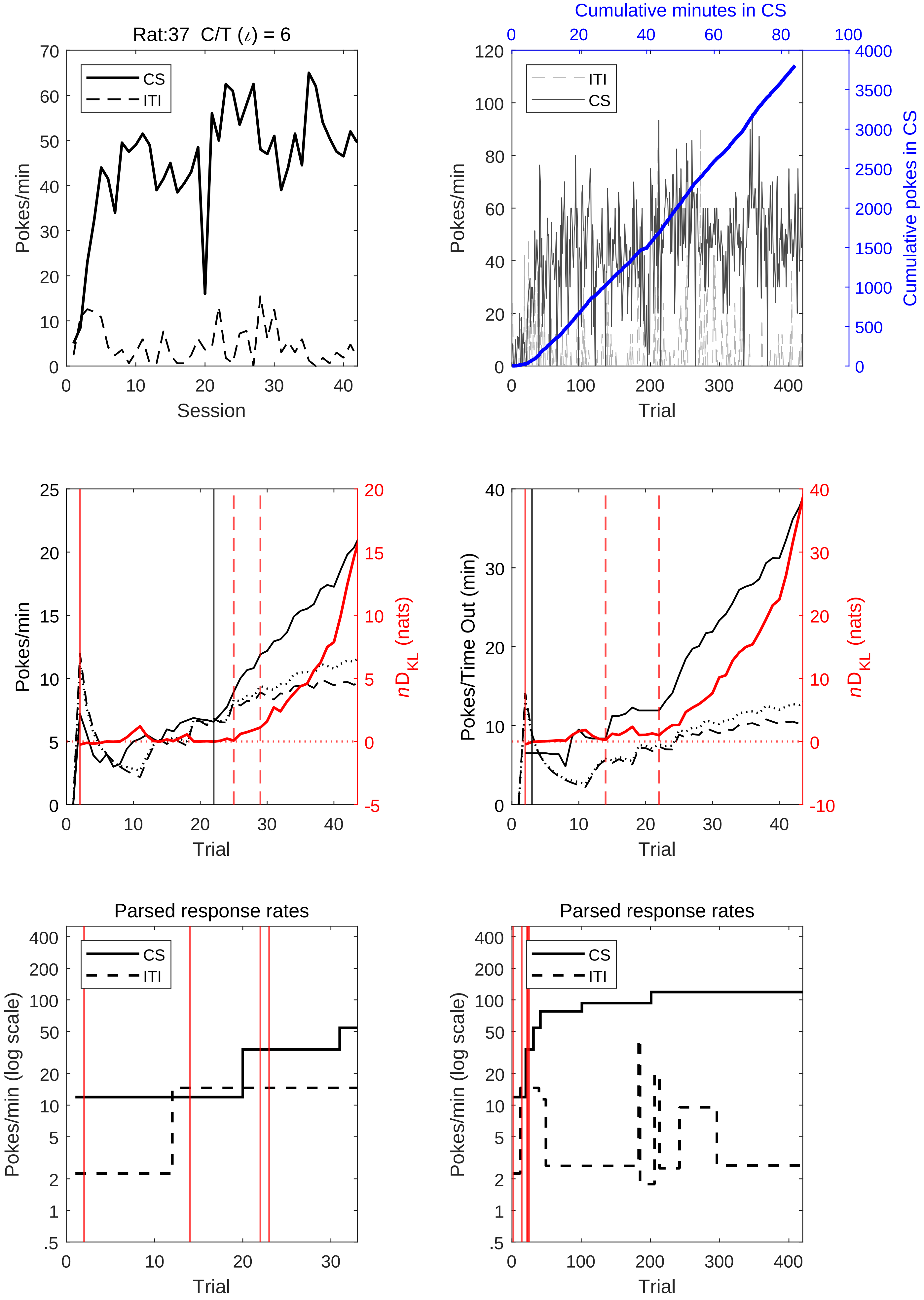
Rat:37 C/T ($\iota$) = 6
CS
ITI
Pokes/min
Session
Cumulative minutes in CS
ITI
CS
Pokes/min
Cumulative pokes in CS
Trial
Pokes/min
$nD_{KL}$ (nats)
Trial
Pokes/Time Out (min)
$nD_{KL}$ (nats)
Trial
Parsed response rates
CS
ITI
Pokes/min (log scale)
Trial
Parsed response rates
CS
ITI
Pokes/min (log scale)
Trial

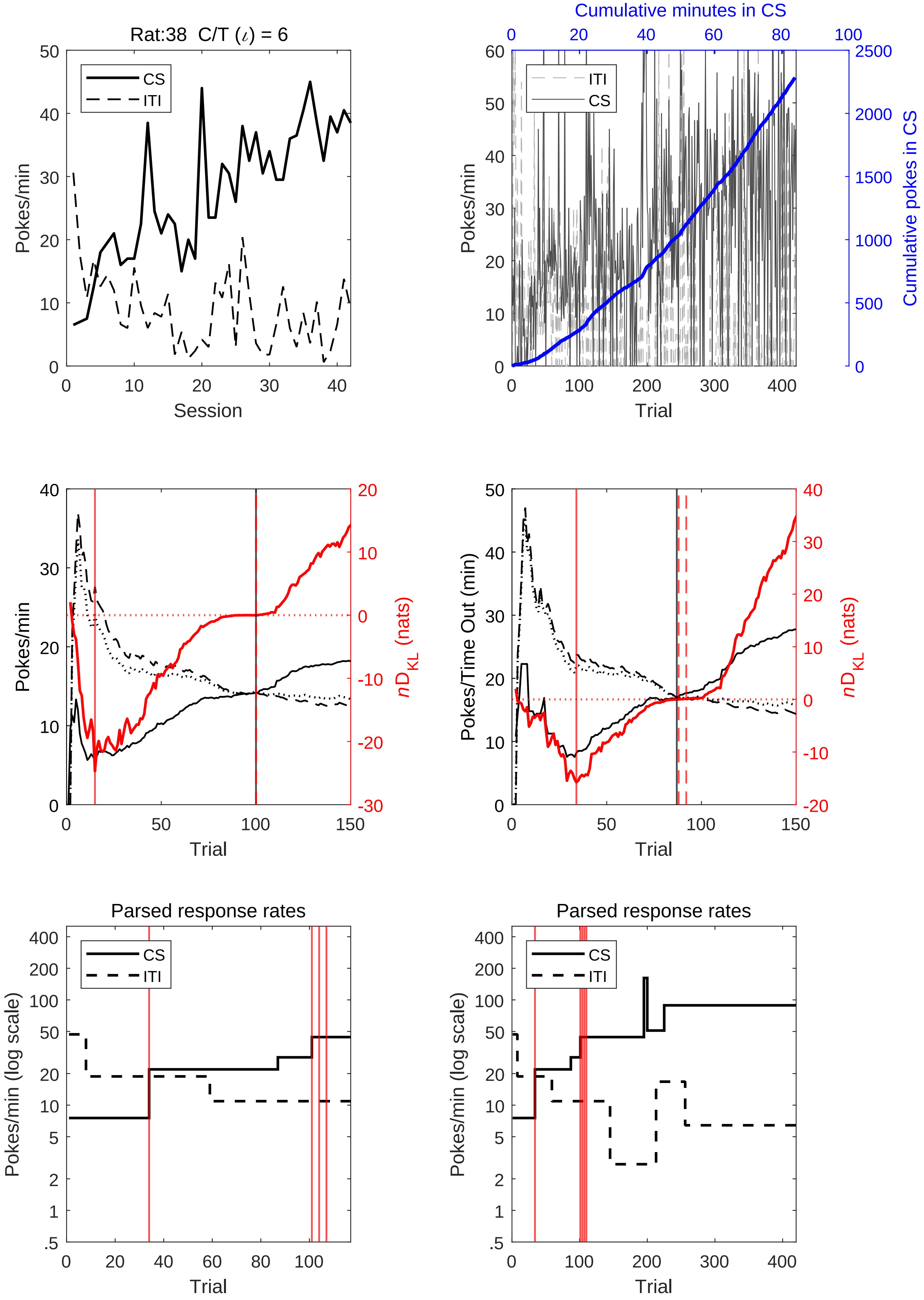

Rat:38 C/T (ι) = 6
CS
ITI
Pokes/min
Session
Cumulative minutes in CS
ITI
CS
Pokes/min
Cumulative pokes in CS
Trial
Pokes/min
nD_KL (nats)
Trial
Pokes/Time Out (min)
nD_KL (nats)
Trial
Parsed response rates
CS
ITI
Pokes/min (log scale)
Trial
Parsed response rates
CS
ITI
Pokes/min (log scale)
Trial

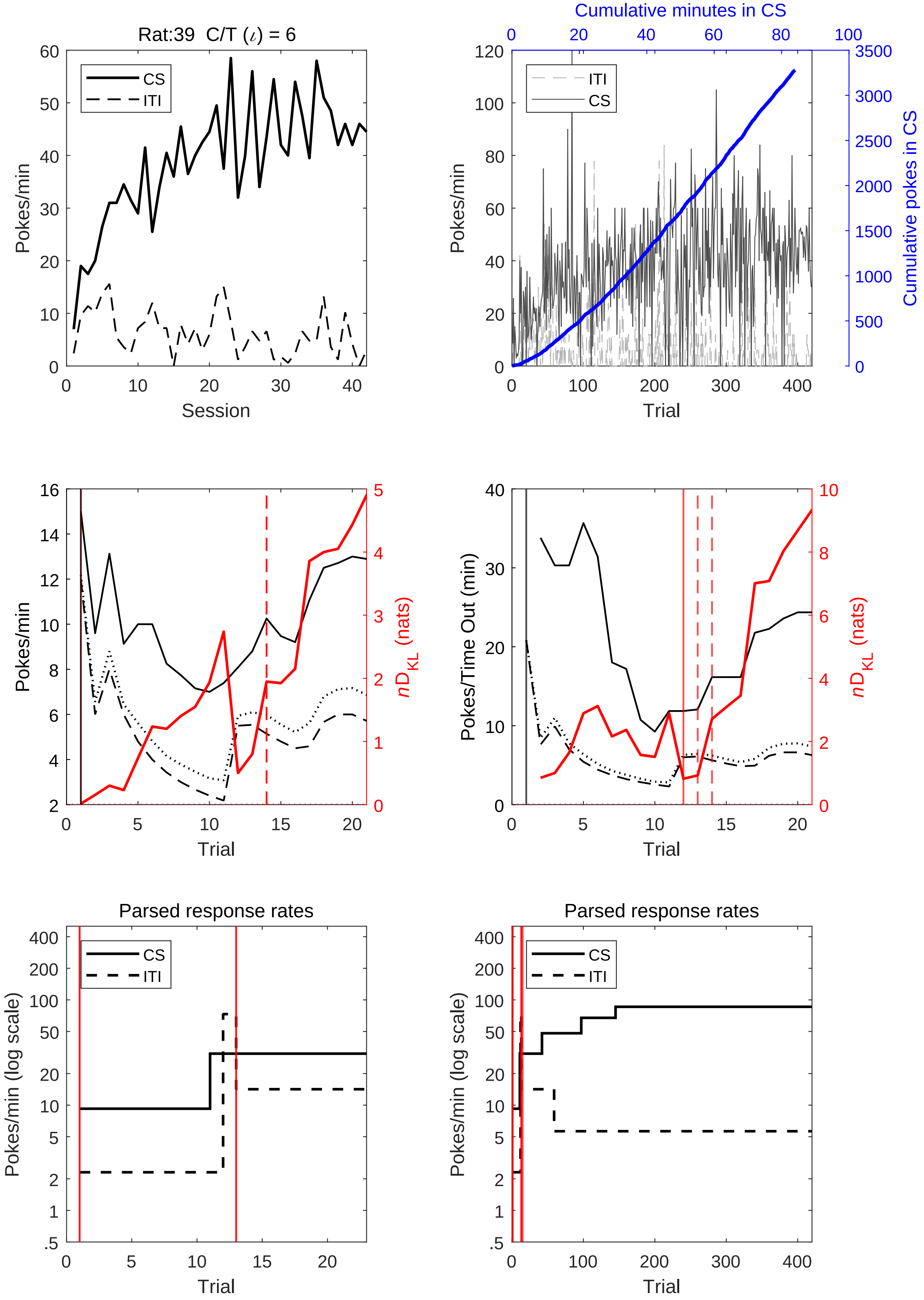

Rat:39 C/T (ι) = 6
CS
ITI
Pokes/min
Session
Cumulative minutes in CS
Cumulative pokes in CS
Trial
nD_KL (nats)
Pokes/Time Out (min)
Parsed response rates
Pokes/min (log scale)

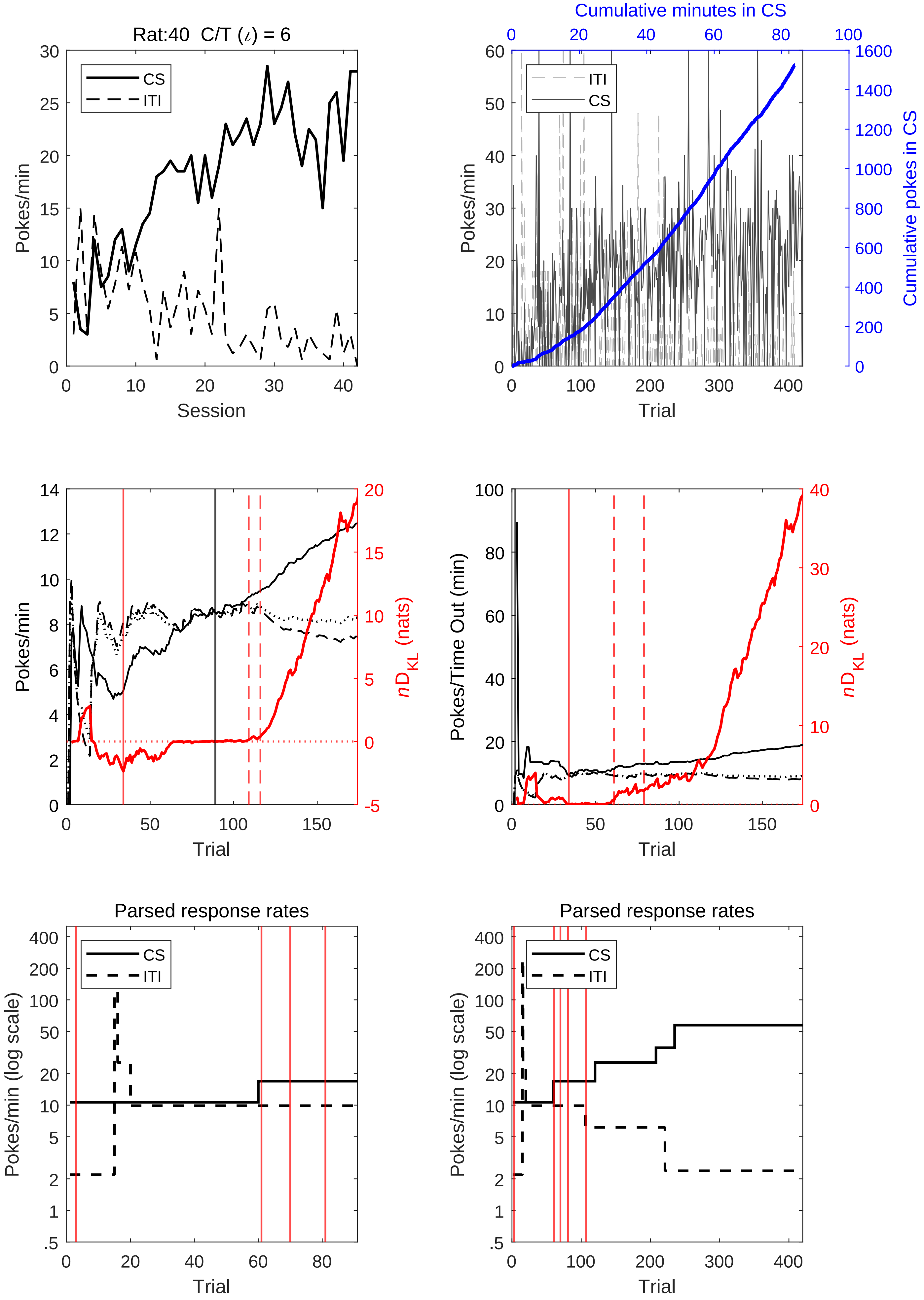

Rat:40 C/T (ι) = 6
CS
ITI
Pokes/min
Session
Cumulative minutes in CS
Cumulative pokes in CS
Trial
nD_KL (nats)
Pokes/Time Out (min)
Parsed response rates
Pokes/min (log scale)

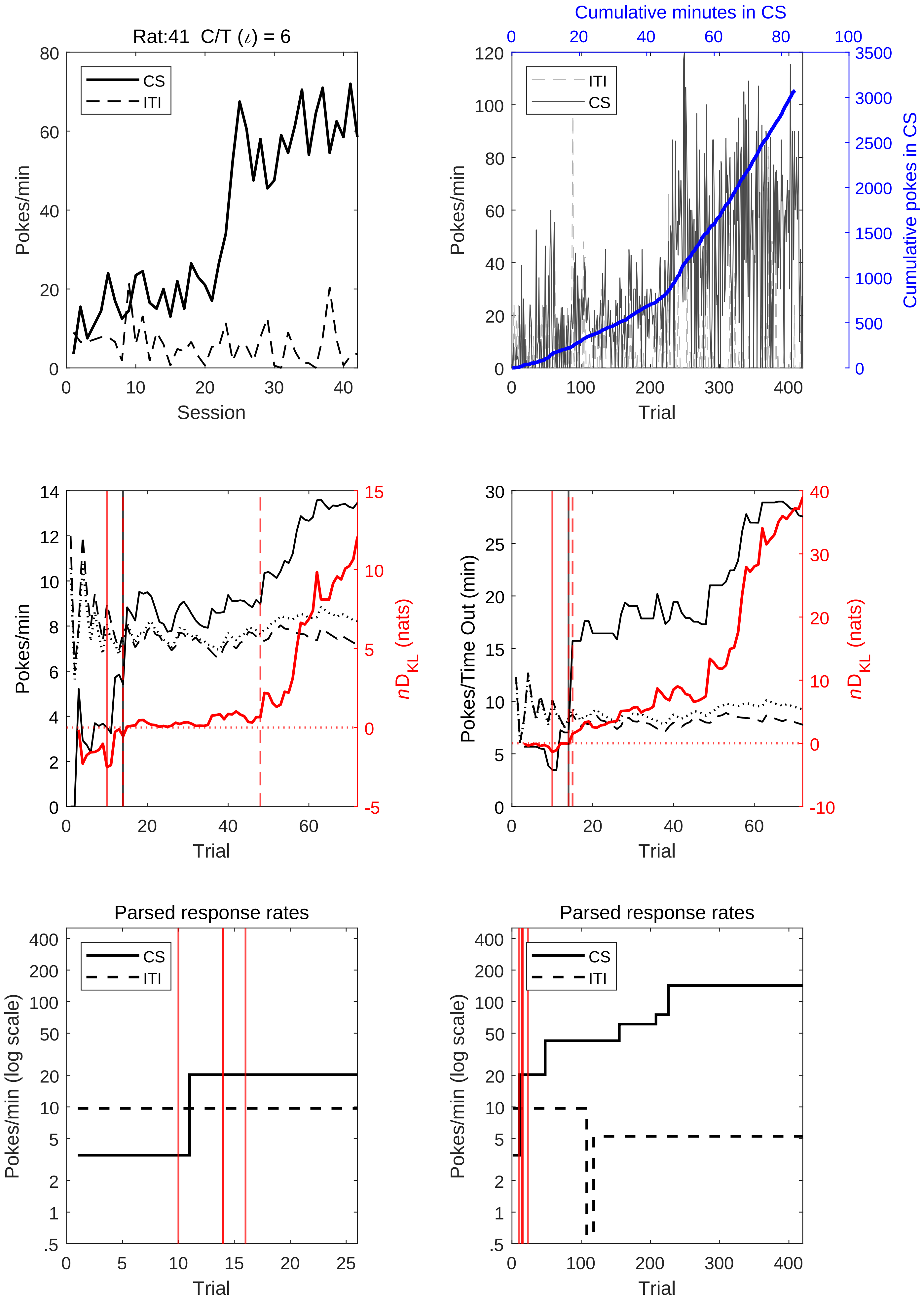
Rat:41 C/T (ι) = 6
CS
ITI
Pokes/min
Session
Cumulative minutes in CS
Cumulative pokes in CS
Trial
Pokes/Time Out (min)
nD_KL (nats)
Parsed response rates
Pokes/min (log scale)

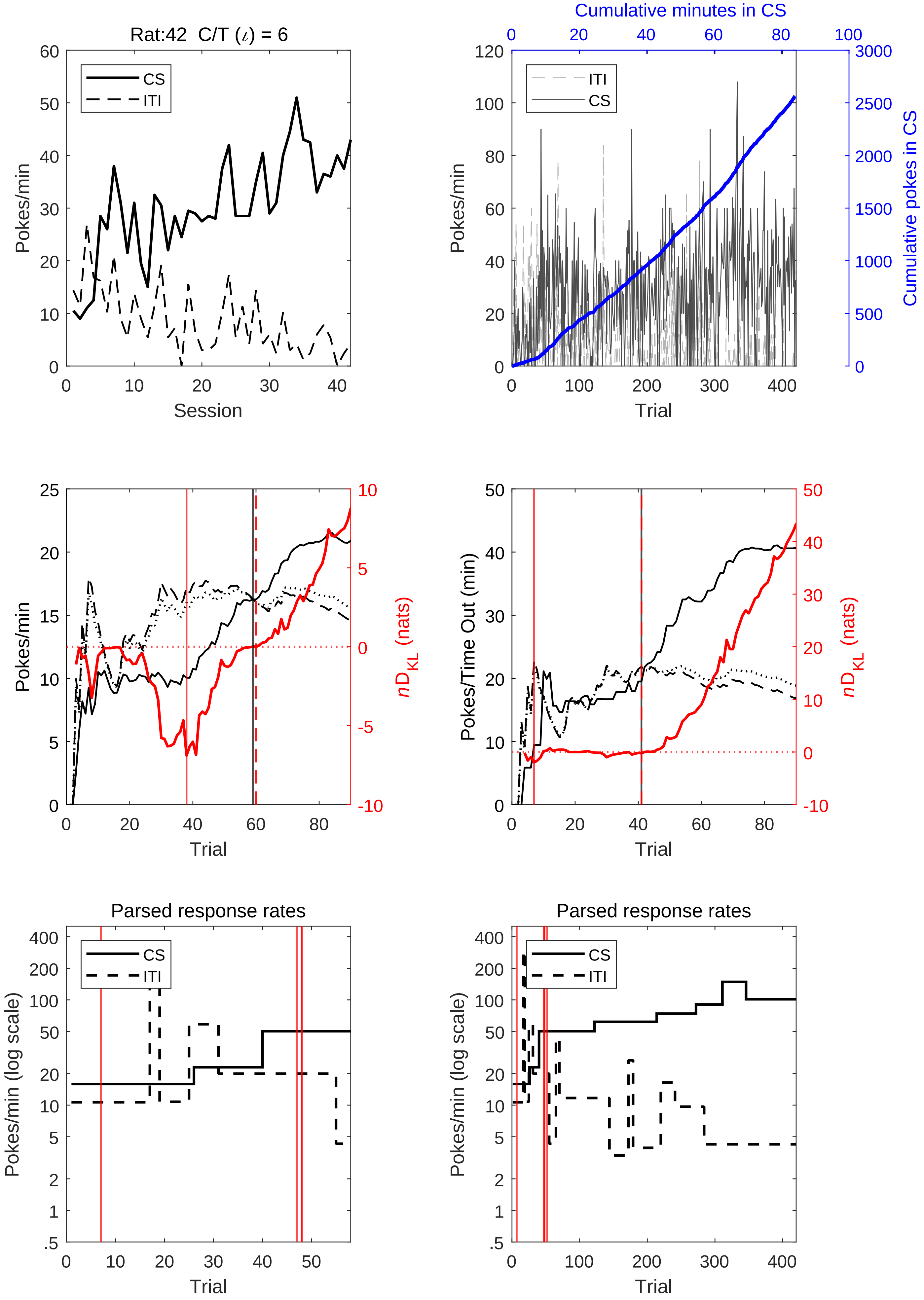
Rat:42 C/T (ι) = 6
CS
ITI
Pokes/min
Session
Cumulative minutes in CS
Cumulative pokes in CS
Trial
nD_KL (nats)
Pokes/Time Out (min)
Parsed response rates
Pokes/min (log scale)

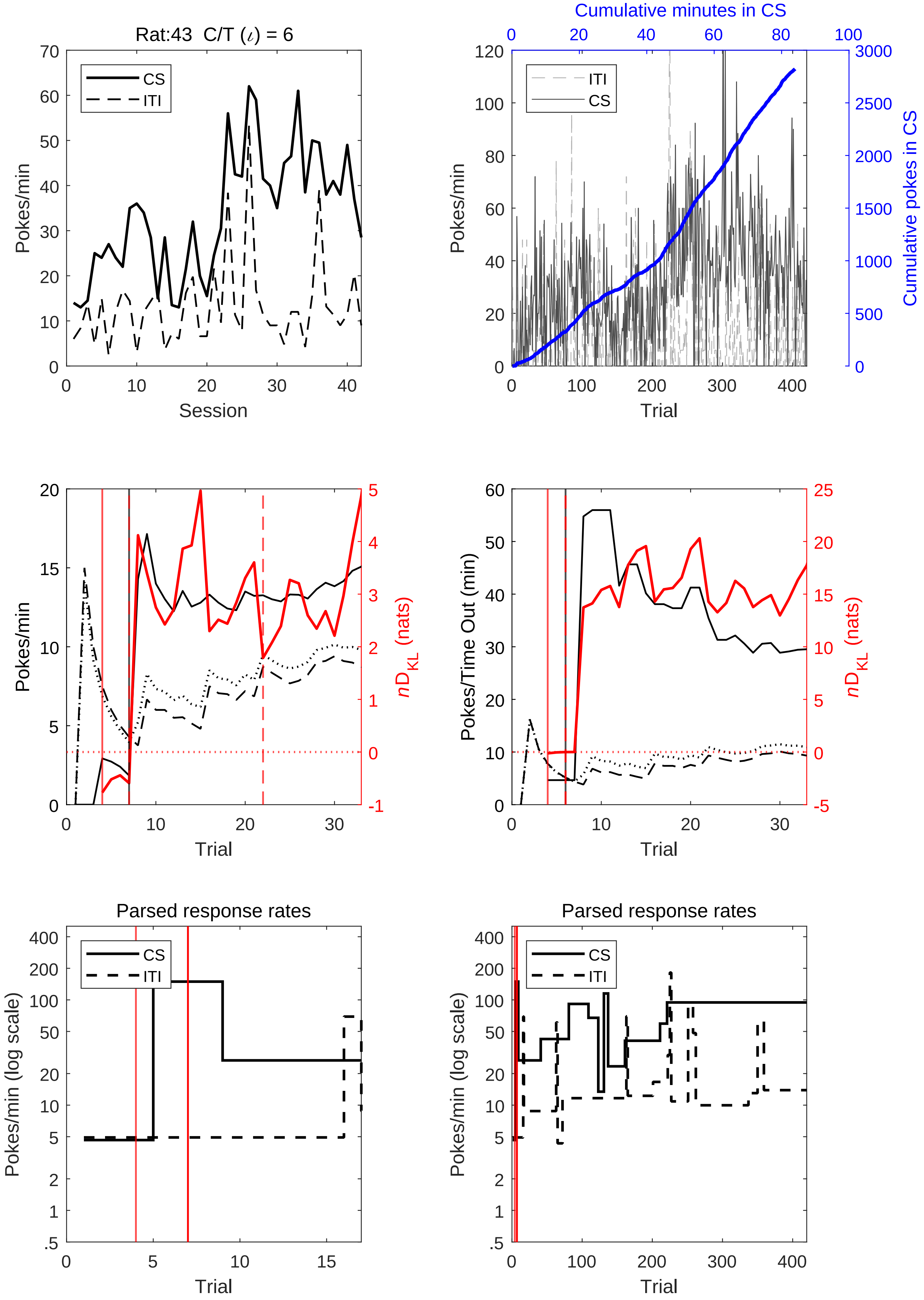
Rat:43 C/T (ι) = 6
CS
ITI
Pokes/min
Session
Cumulative minutes in CS
Cumulative pokes in CS
Trial
nD_KL (nats)
Pokes/Time Out (min)
Parsed response rates
Pokes/min (log scale)

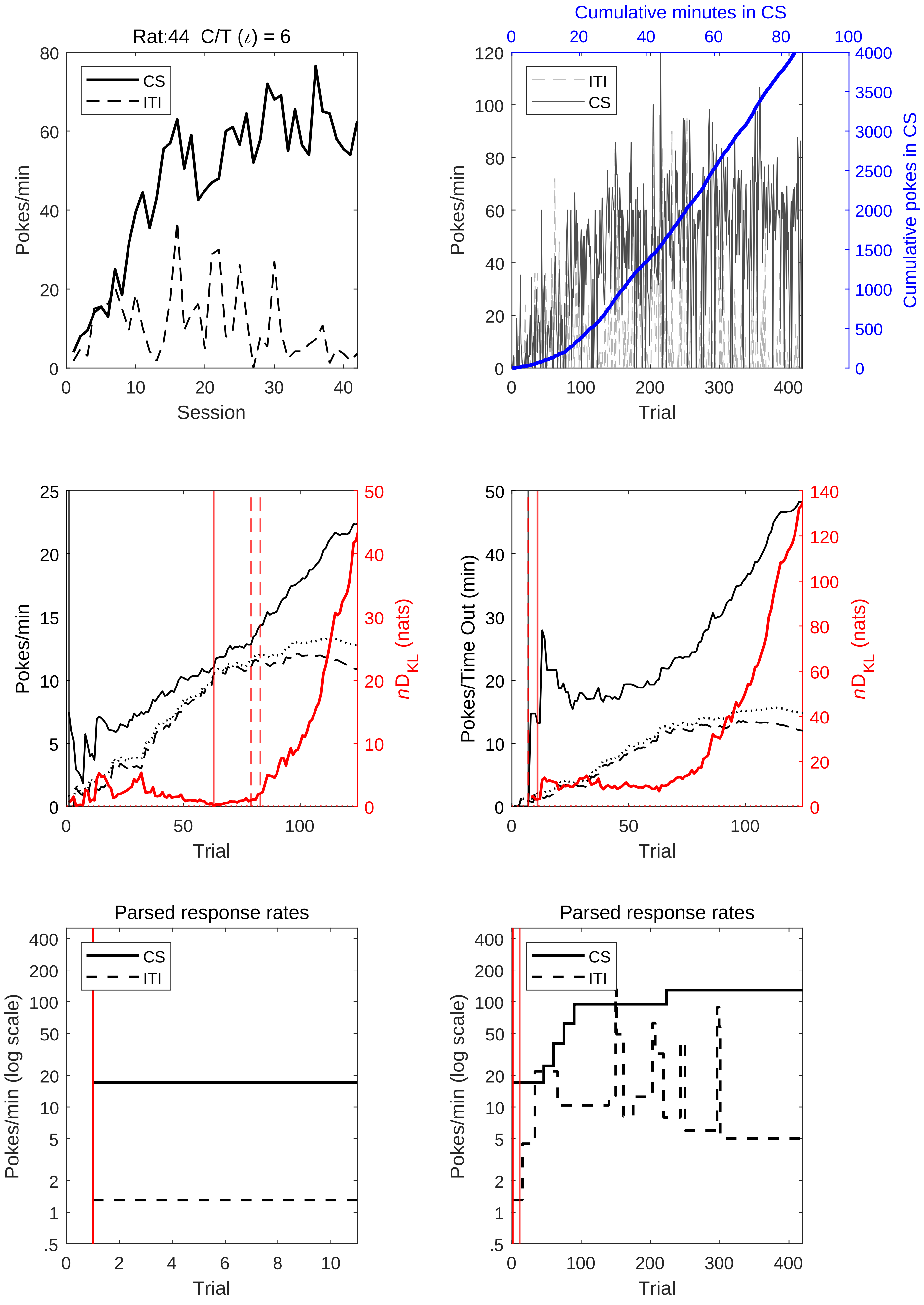
Rat:44 C/T (ι) = 6
CS
ITI
Pokes/min
Session
Cumulative minutes in CS
Cumulative pokes in CS
Trial
Pokes/Time Out (min)
nD_KL (nats)
Parsed response rates
Pokes/min (log scale)

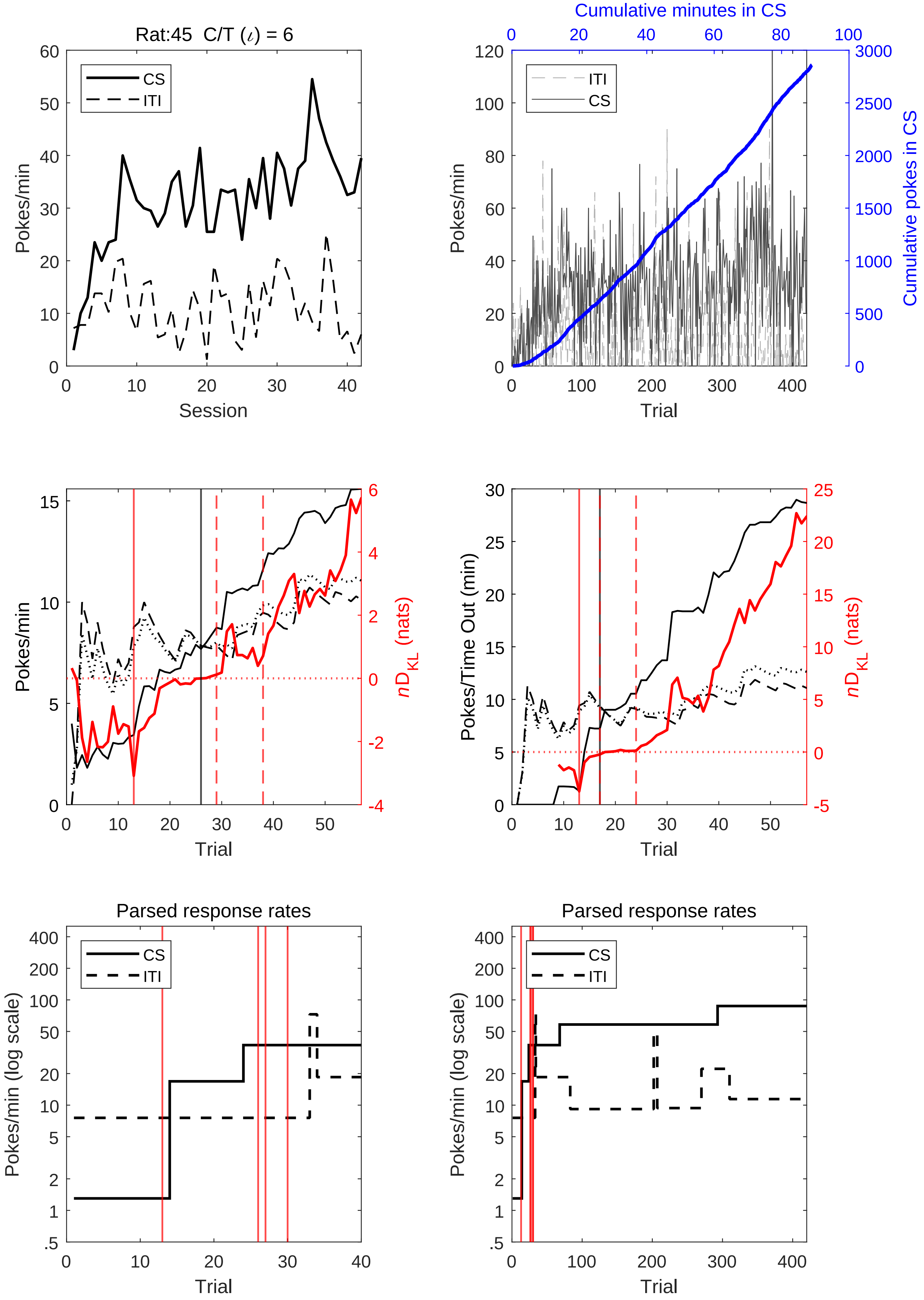
Rat:45 C/T (ι) = 6
CS
ITI
Pokes/min
Session
Cumulative minutes in CS
Cumulative pokes in CS
Trial
nD_KL (nats)
Pokes/Time Out (min)
Parsed response rates
Pokes/min (log scale)

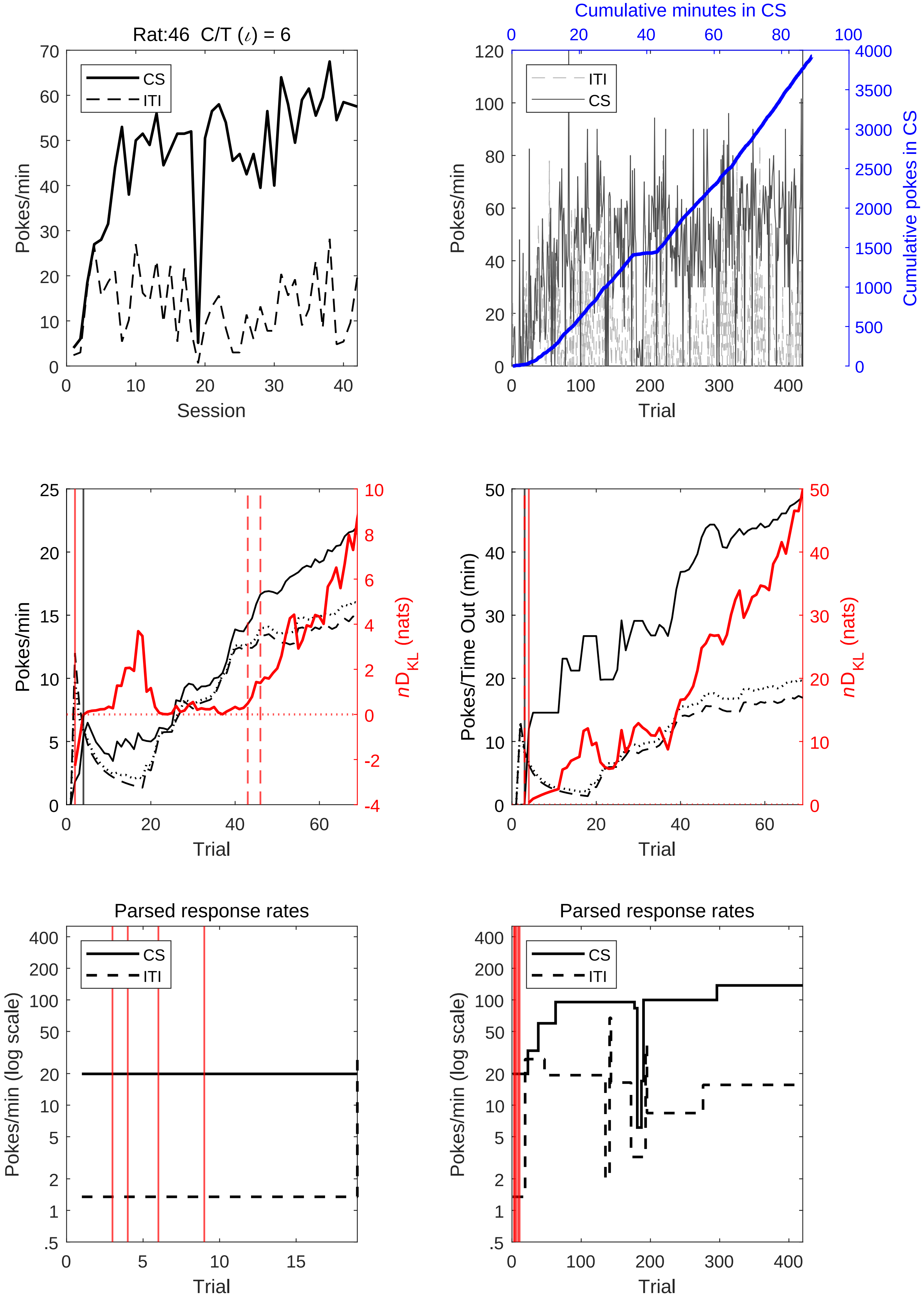
Rat:46 C/T (ι) = 6
CS
ITI
Pokes/min
Session
Cumulative minutes in CS
Cumulative pokes in CS
Trial
Pokes/Time Out (min)
nD_KL (nats)
Parsed response rates
Pokes/min (log scale)

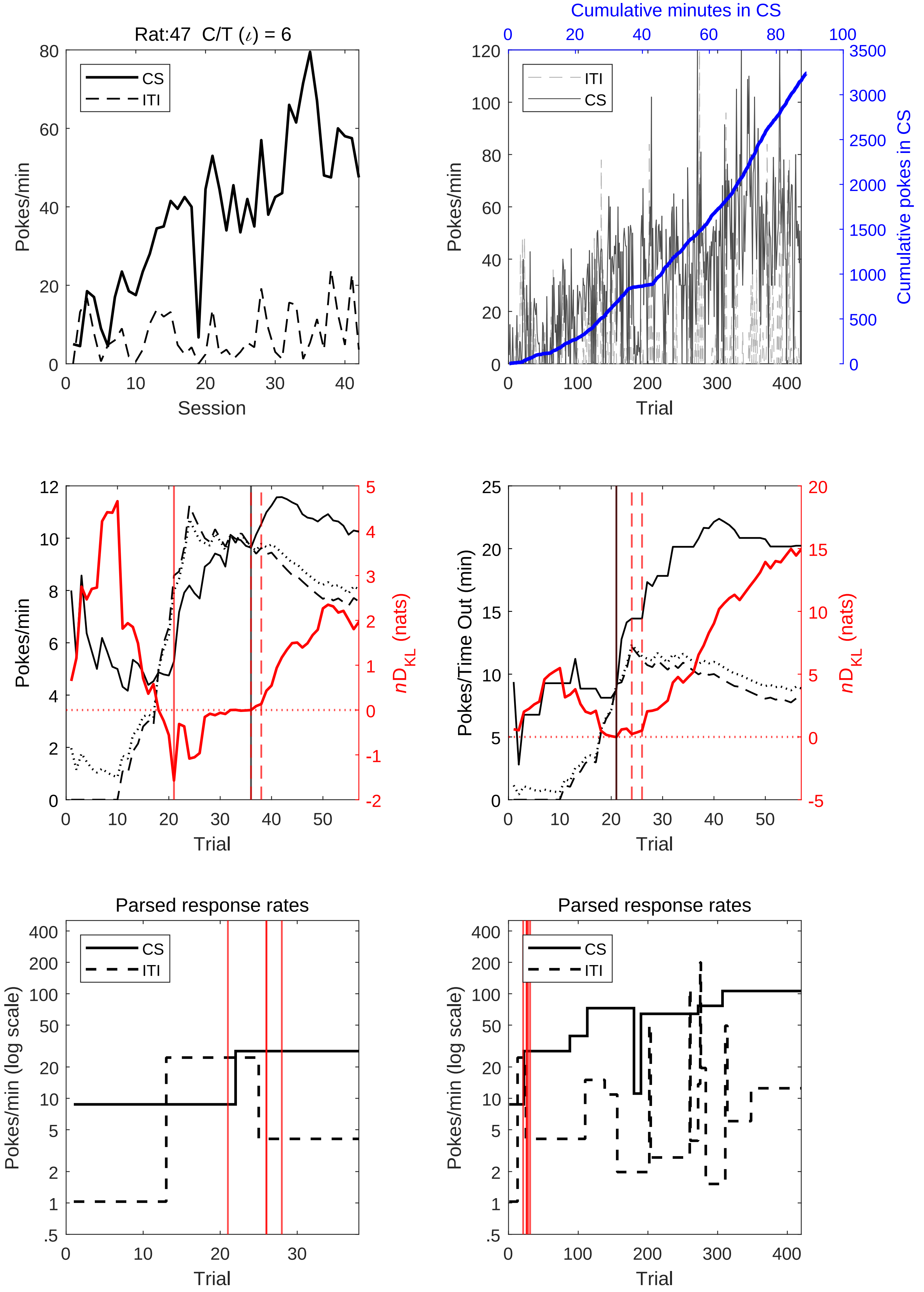

Rat:47 C/T (ι) = 6
CS
ITI
Pokes/min
Session
Cumulative minutes in CS
Cumulative pokes in CS
Trial
nD_KL (nats)
Pokes/Time Out (min)
Parsed response rates
Pokes/min (log scale)

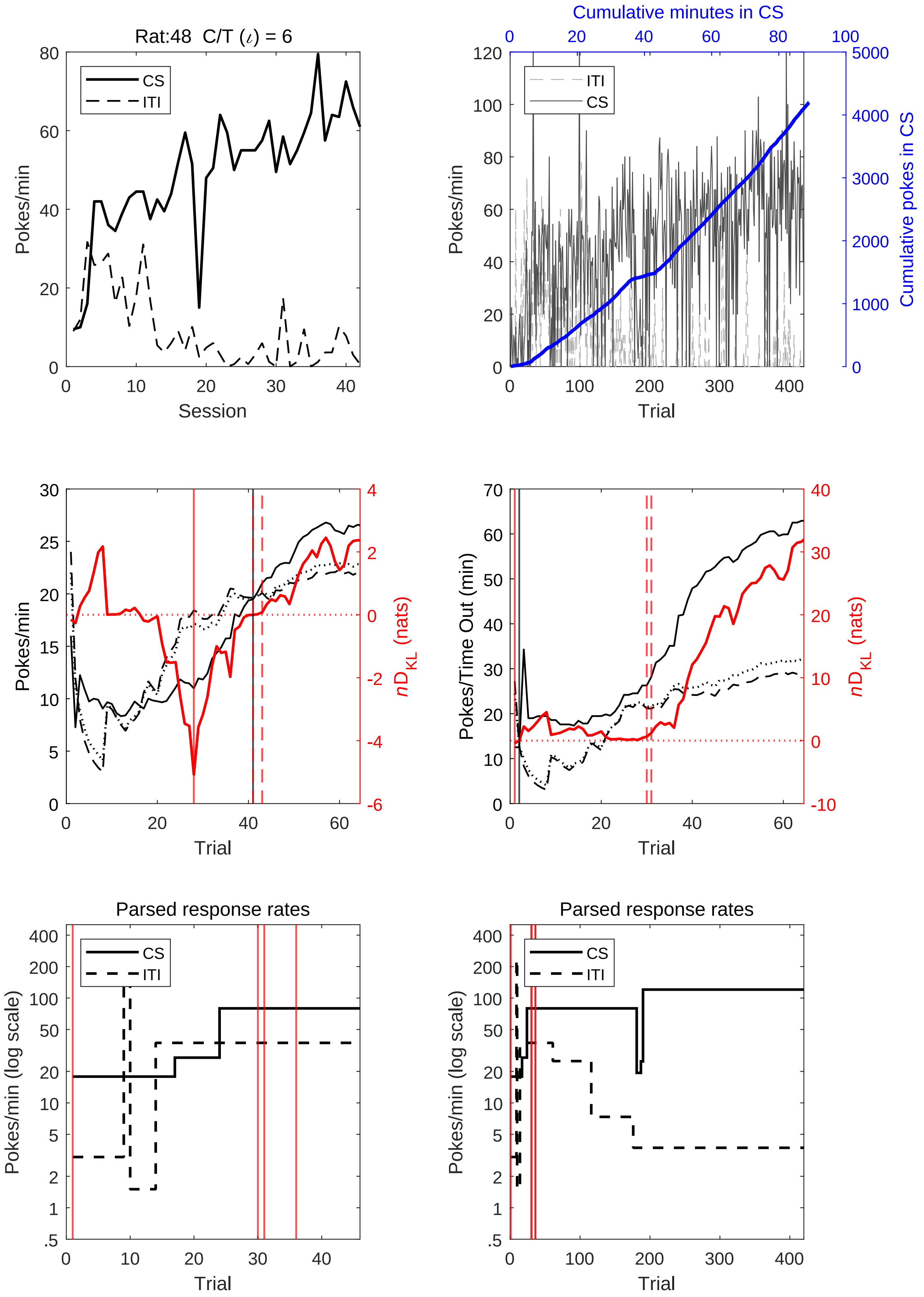

Rat:48 C/T (ι) = 6
CS
ITI
Pokes/min
Session
Cumulative minutes in CS
Cumulative pokes in CS
ITI
CS
Pokes/min
Trial
Pokes/min
$nD_{KL}$ (nats)
Trial
Pokes/Time Out (min)
$nD_{KL}$ (nats)
Trial
Parsed response rates
CS
ITI
Pokes/min (log scale)
Trial
Parsed response rates
CS
ITI
Pokes/min (log scale)
Trial

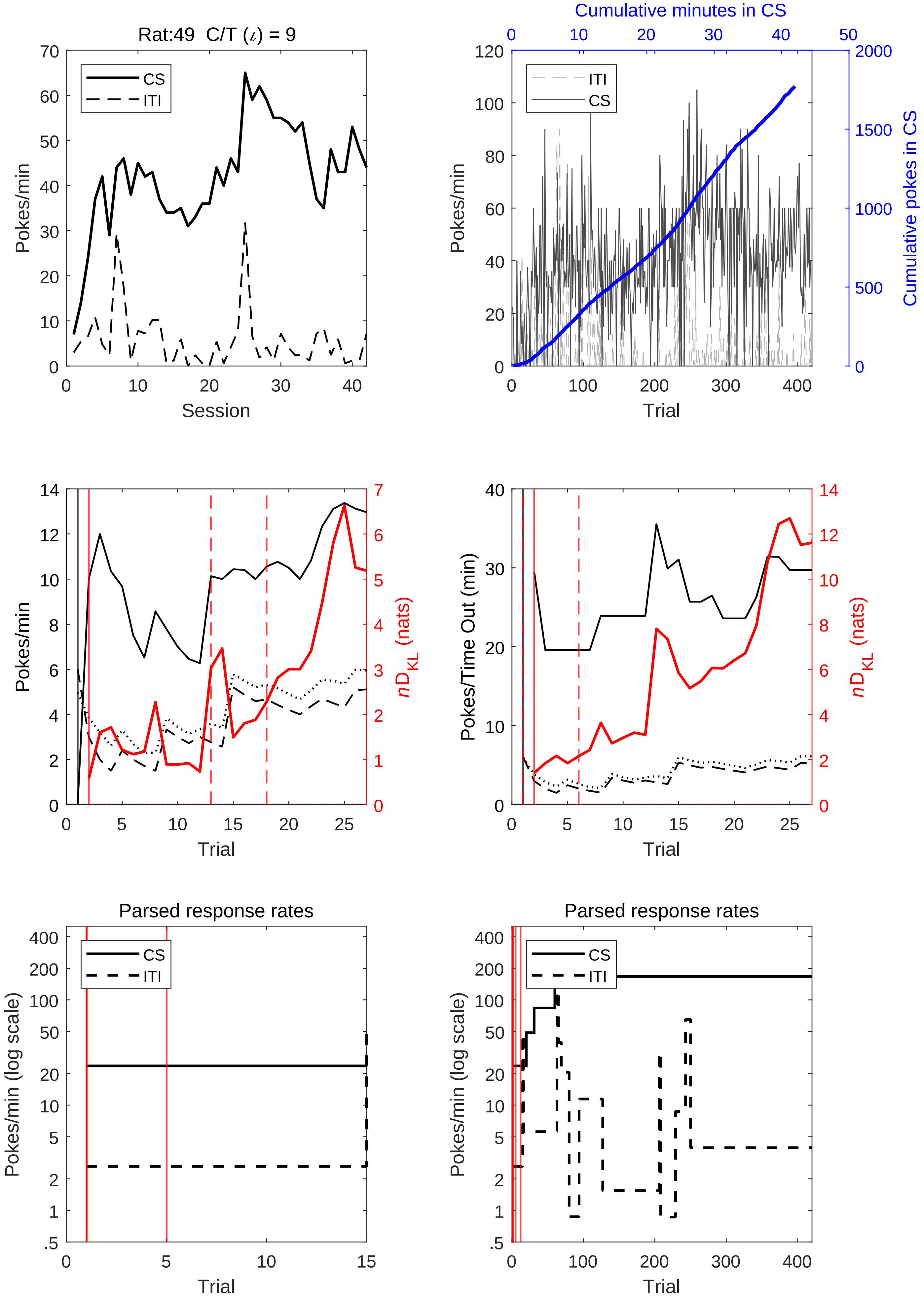
Rat:49 C/T (ι) = 9
CS
ITI
Pokes/min
Session
Cumulative minutes in CS
Cumulative pokes in CS
Trial
Pokes/min
nD_KL (nats)
Pokes/Time Out (min)
Parsed response rates
Pokes/min (log scale)

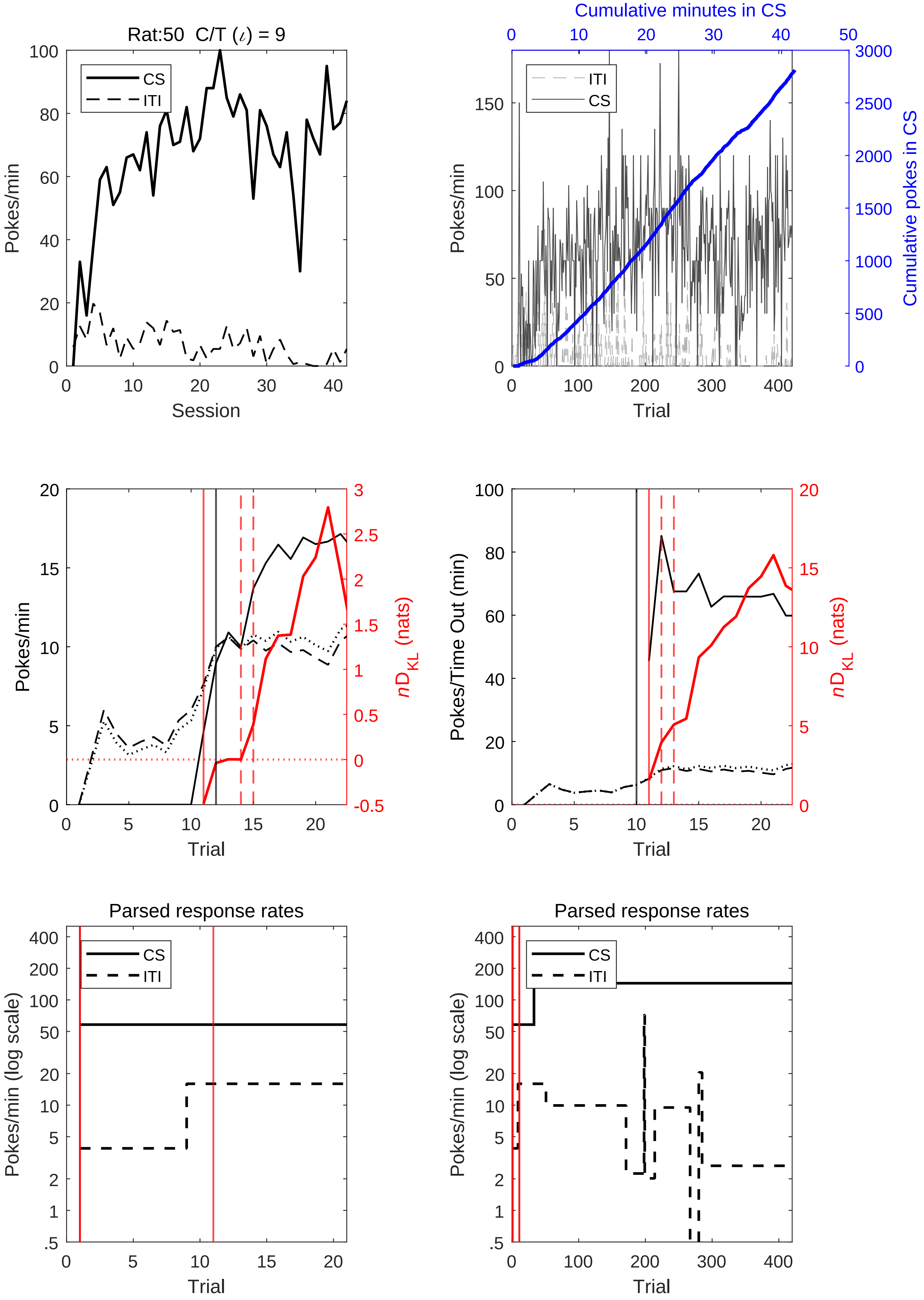
Rat:50 C/T (ι) = 9
CS
ITI
Pokes/min
Session
Cumulative minutes in CS
Cumulative pokes in CS
Trial
nD_KL (nats)
Pokes/Time Out (min)
Parsed response rates
Pokes/min (log scale)

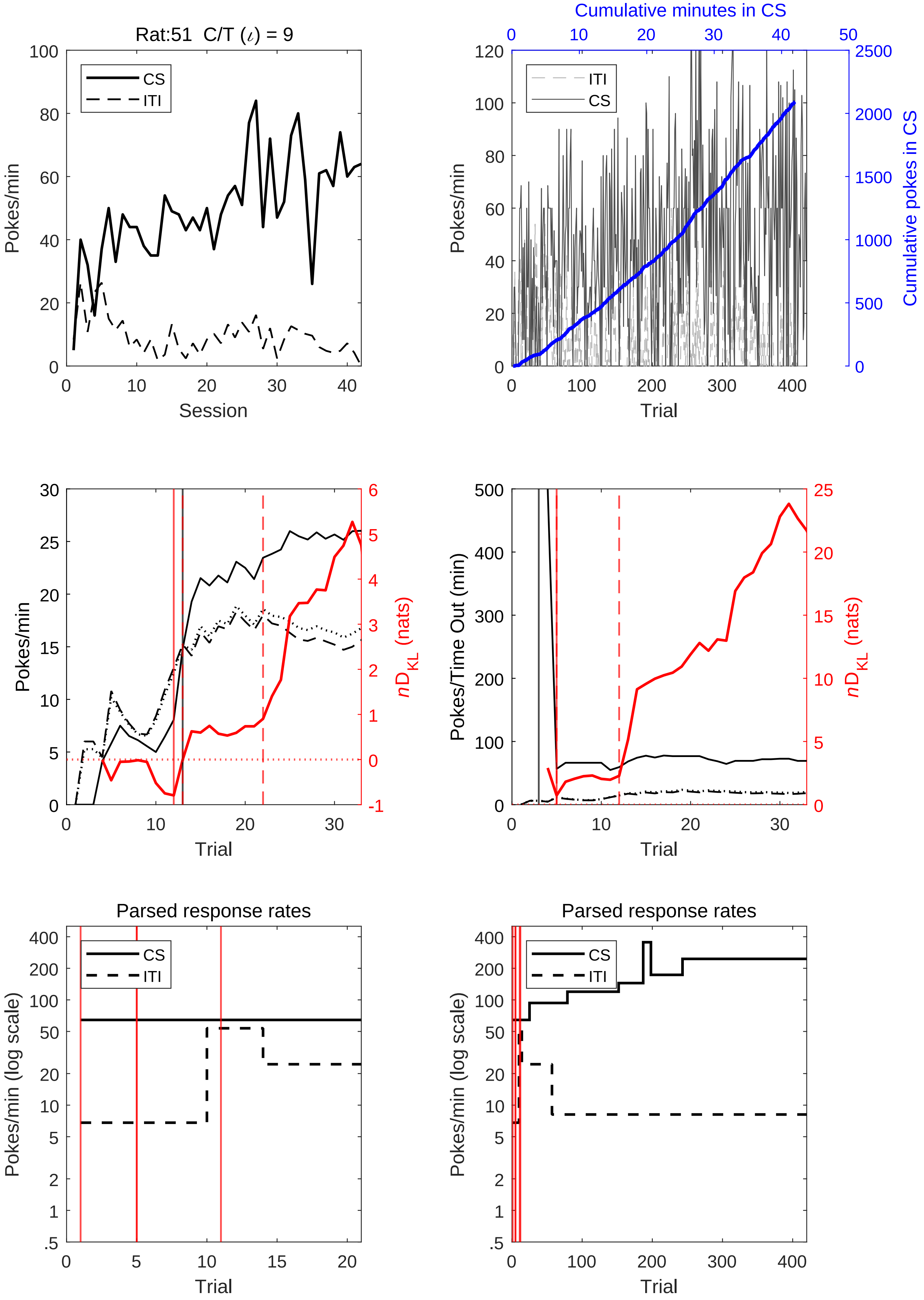

Rat:51 C/T (ι) = 9
CS
ITI
Pokes/min
Session
Cumulative minutes in CS
ITI
CS
Pokes/min
Cumulative pokes in CS
Trial
Pokes/min
nD_KL (nats)
Trial
Pokes/Time Out (min)
nD_KL (nats)
Trial
Parsed response rates
CS
ITI
Pokes/min (log scale)
Trial
Parsed response rates
CS
ITI
Pokes/min (log scale)
Trial

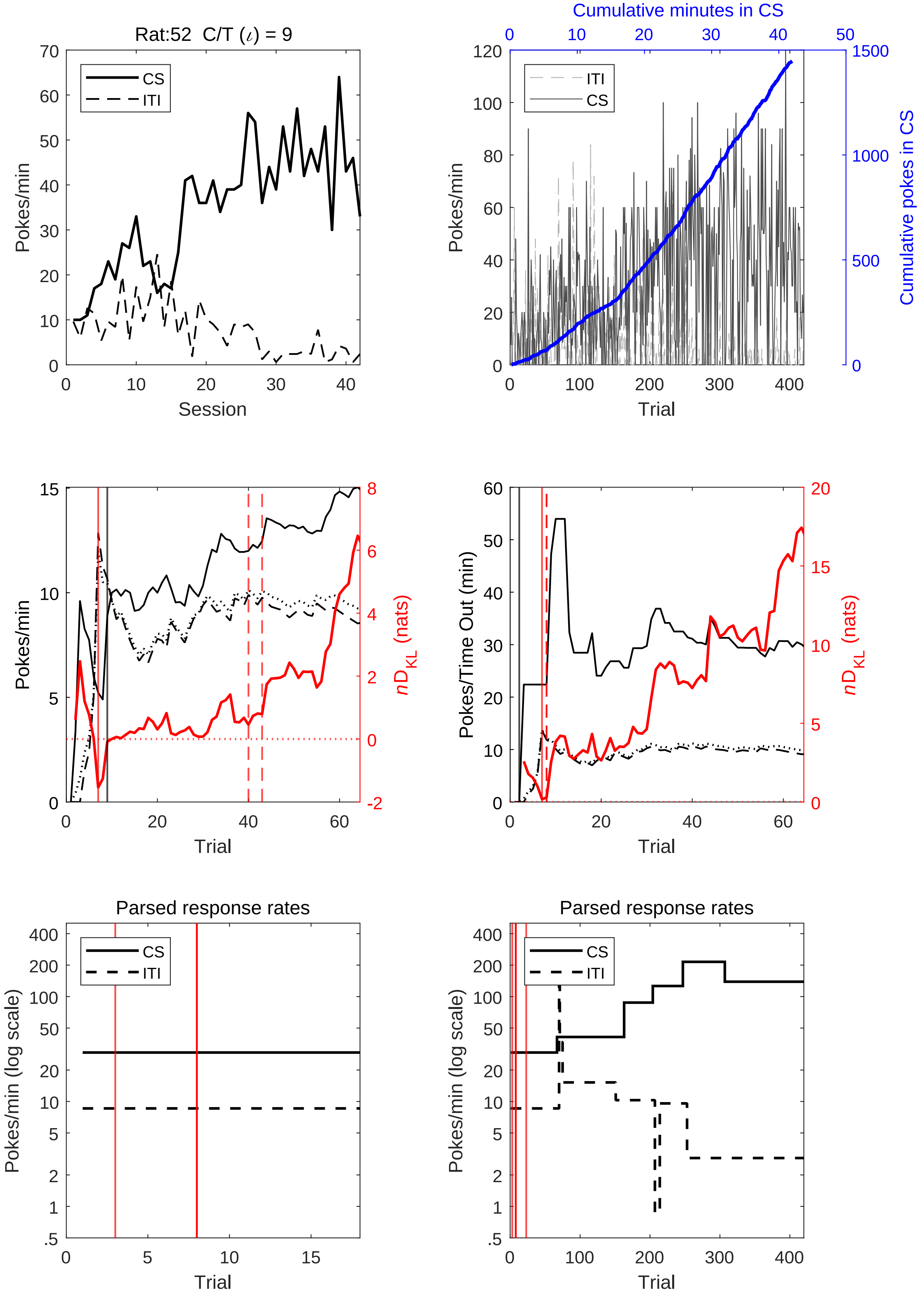
Rat:52 C/T (ι) = 9
CS
ITI
Pokes/min
Session
Cumulative minutes in CS
Cumulative pokes in CS
Trial
Pokes/Time Out (min)
nD_KL (nats)
Parsed response rates
Pokes/min (log scale)

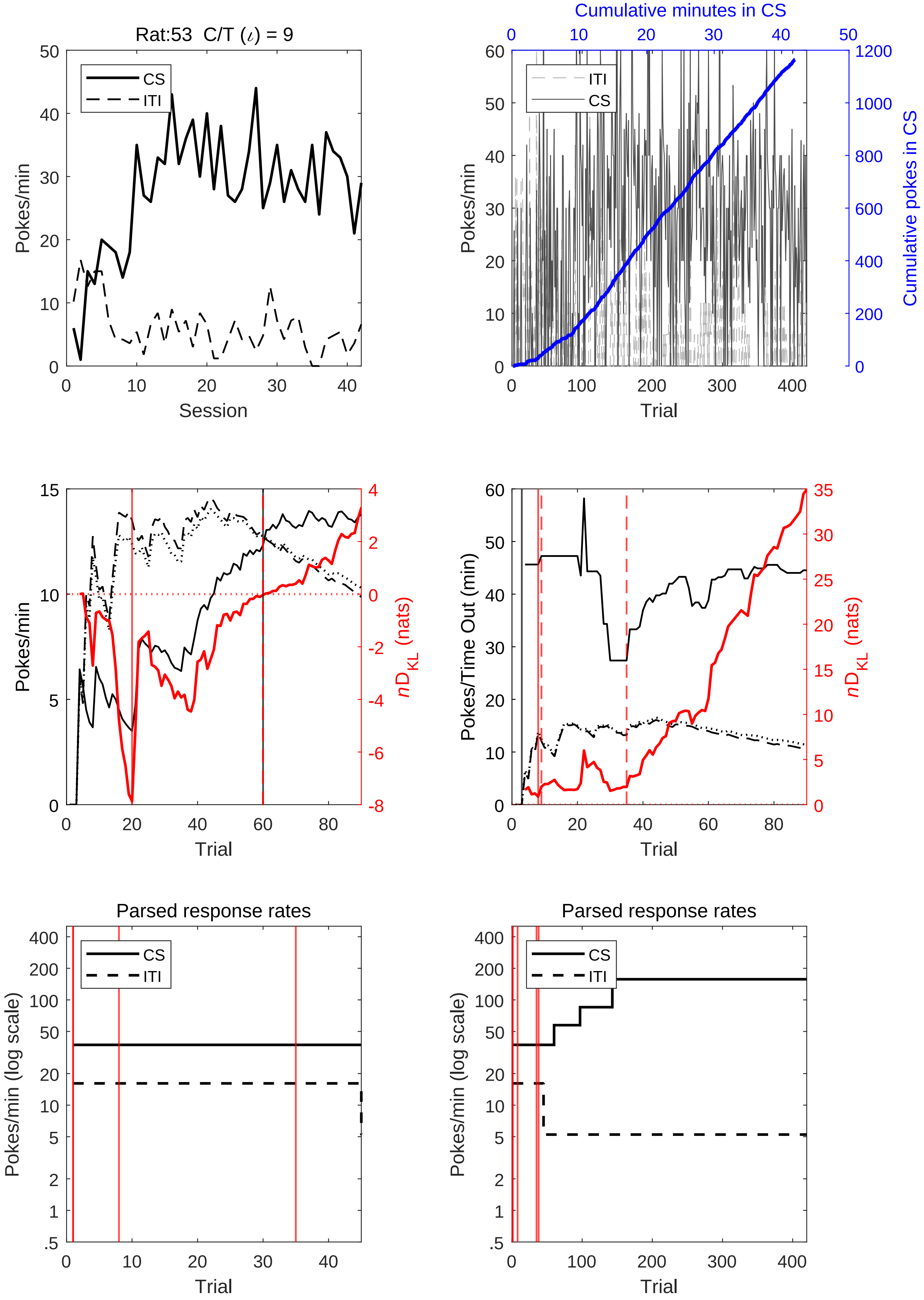
Rat:53 C/T (ι) = 9
CS
ITI
Pokes/min
Session
Cumulative minutes in CS
Cumulative pokes in CS
Trial
nD_KL (nats)
Pokes/Time Out (min)
Parsed response rates
Pokes/min (log scale)

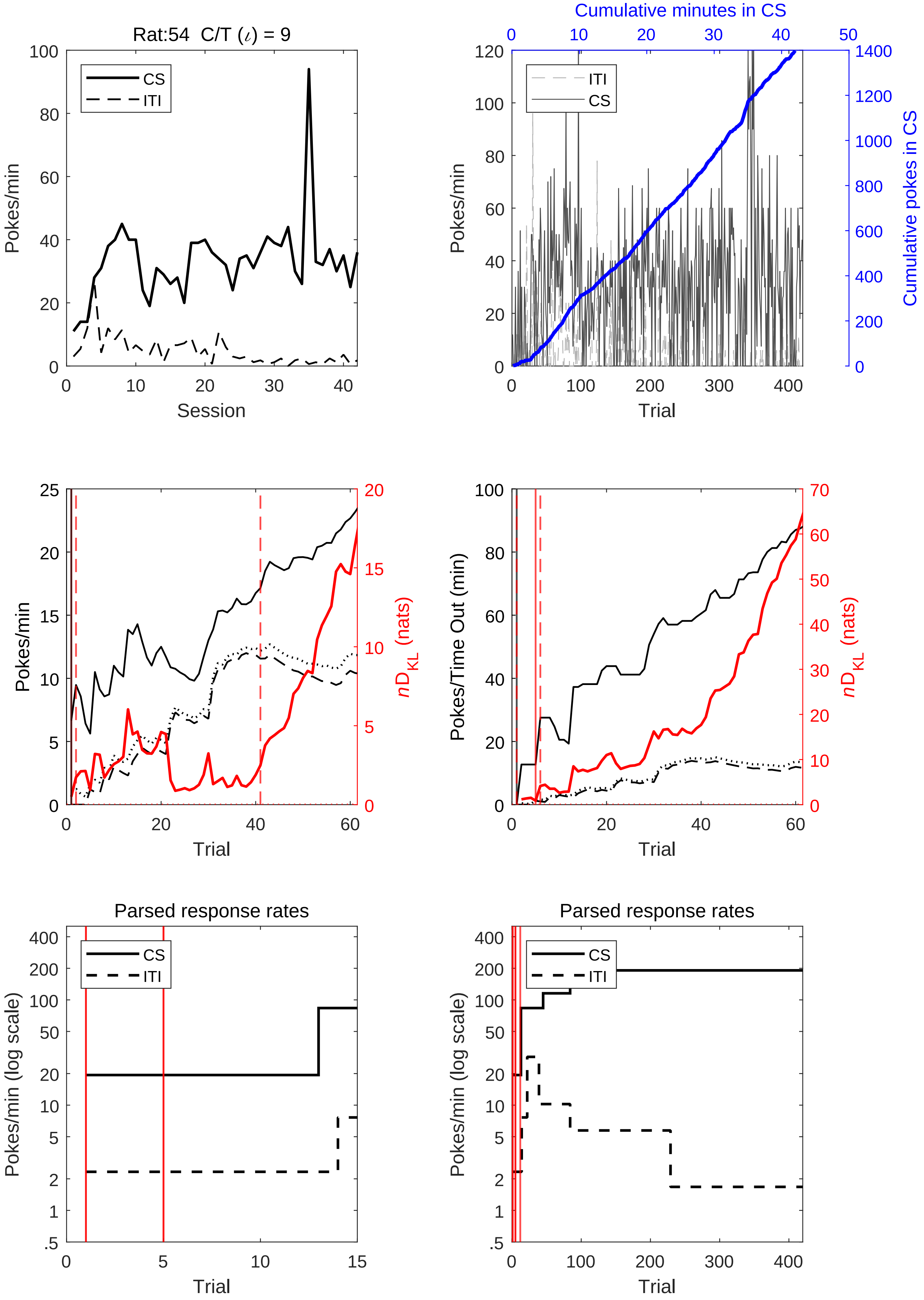
Rat:54 C/T (ι) = 9
CS
ITI
Pokes/min
Session
Cumulative minutes in CS
Cumulative pokes in CS
Trial
Pokes/Time Out (min)
$nD_{KL}$ (nats)
Parsed response rates
Pokes/min (log scale)

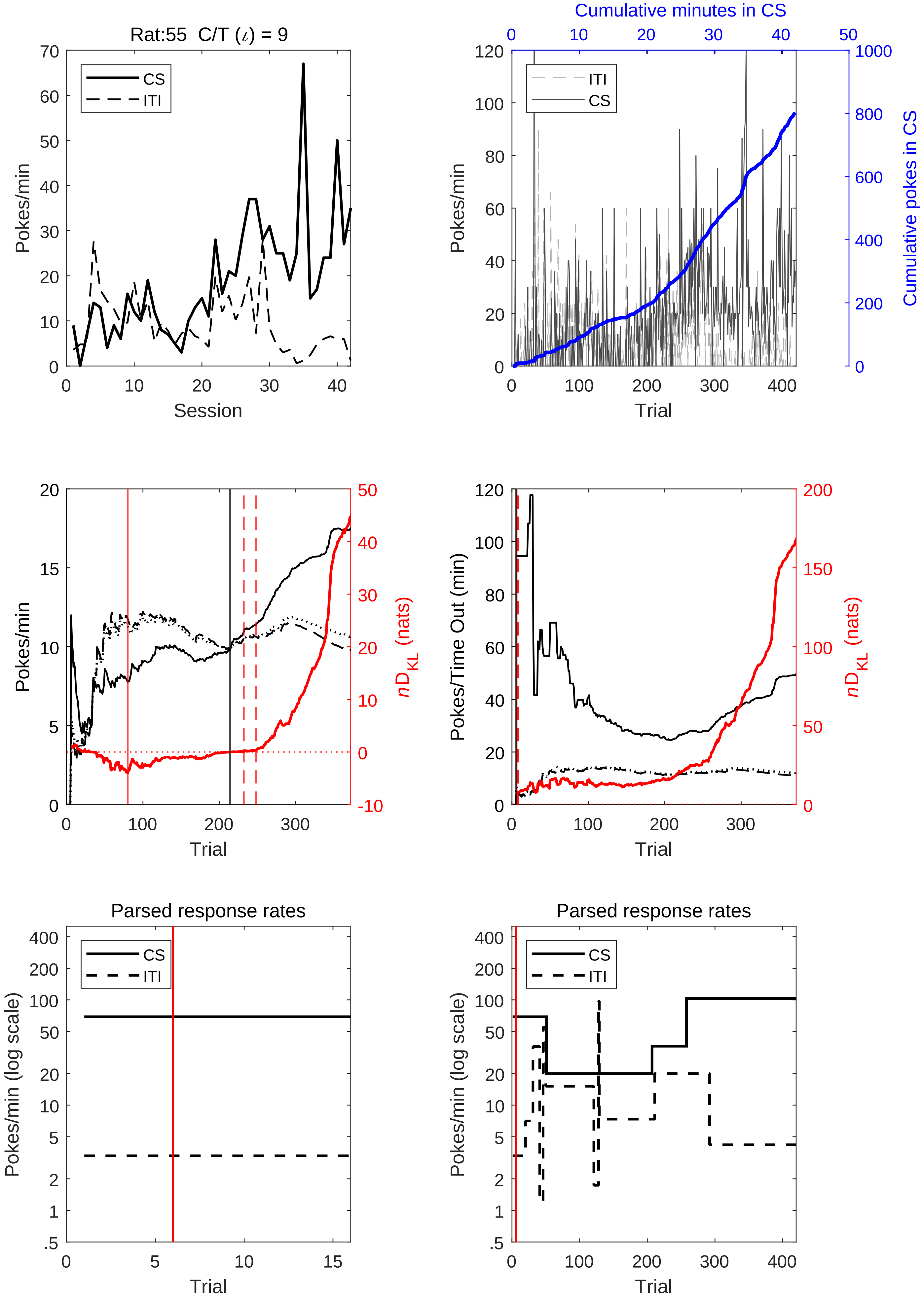
Rat:55 C/T (ι) = 9
Pokes/min
Session
CS
ITI
Cumulative minutes in CS
Pokes/min
Cumulative pokes in CS
Trial
ITI
CS
Pokes/min
nD_KL (nats)
Trial
Pokes/Time Out (min)
nD_KL (nats)
Trial
Parsed response rates
Pokes/min (log scale)
Trial
CS
ITI
Parsed response rates
Pokes/min (log scale)
Trial
CS
ITI

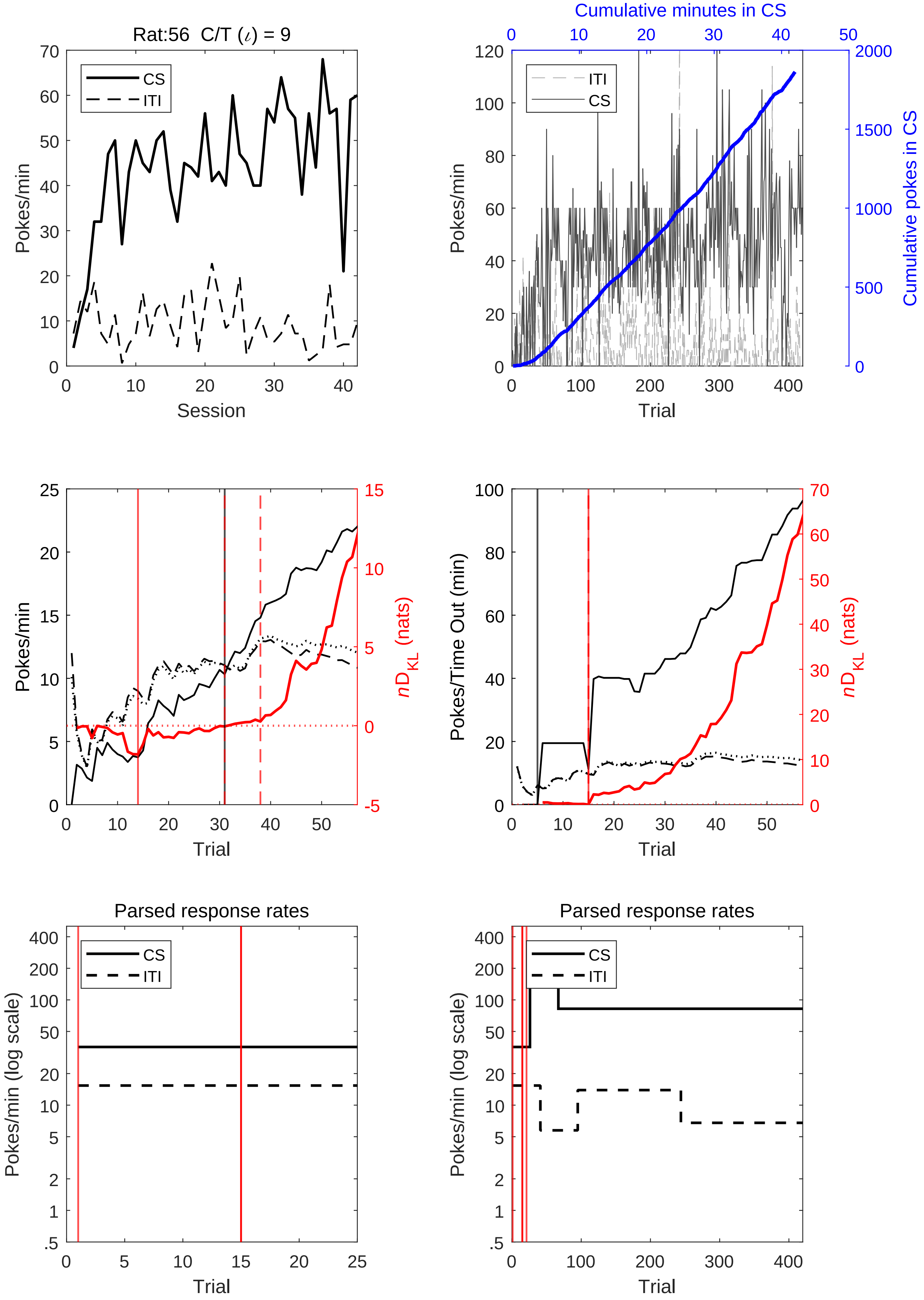
Rat:56 C/T (ι) = 9
CS
ITI
Pokes/min
Session
Cumulative minutes in CS
Cumulative pokes in CS
Trial
nD_KL (nats)
Pokes/Time Out (min)
Parsed response rates
Pokes/min (log scale)

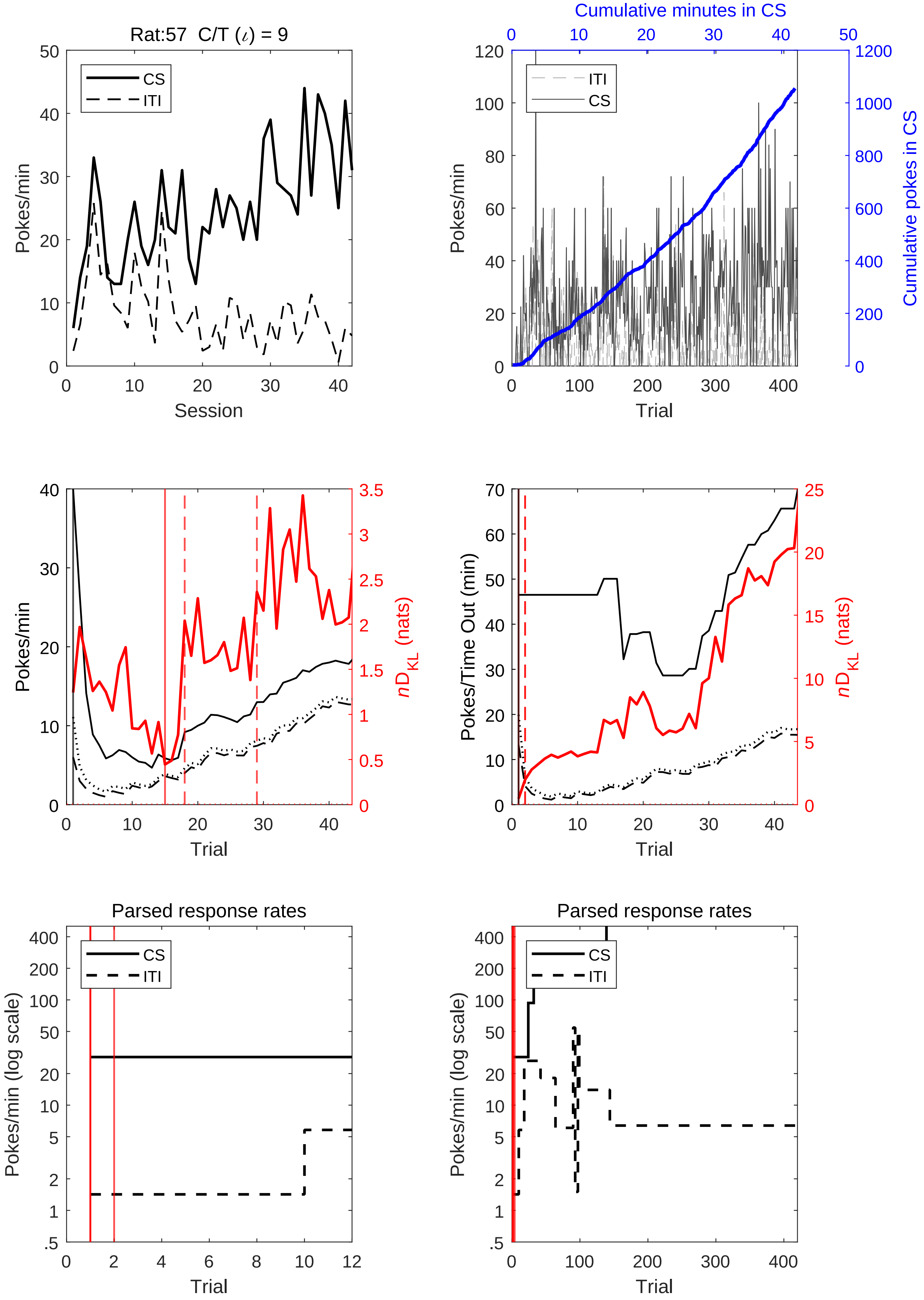
Rat:57 C/T (ι) = 9
CS
ITI
Pokes/min
Session
Cumulative minutes in CS
Cumulative pokes in CS
Pokes/min
Trial
Pokes/min
nD_KL (nats)
Trial
Pokes/Time Out (min)
nD_KL (nats)
Trial
Parsed response rates
Pokes/min (log scale)
Trial
Parsed response rates
Pokes/min (log scale)
Trial

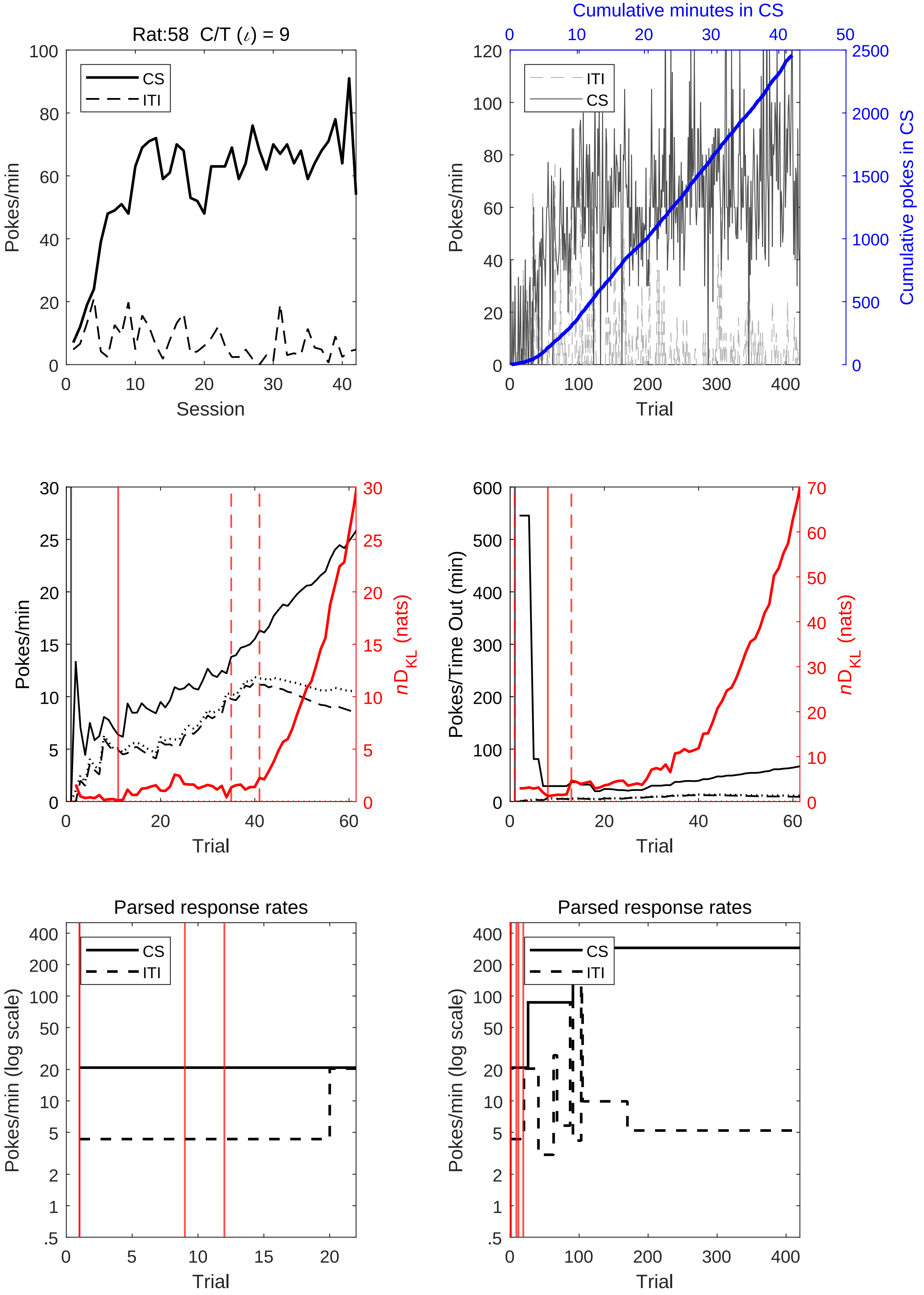
Rat:58 C/T (ι) = 9
Pokes/min
Session
CS
ITI
Cumulative minutes in CS
Cumulative pokes in CS
Trial
nD_KL (nats)
Pokes/Time Out (min)
Parsed response rates
Pokes/min (log scale)

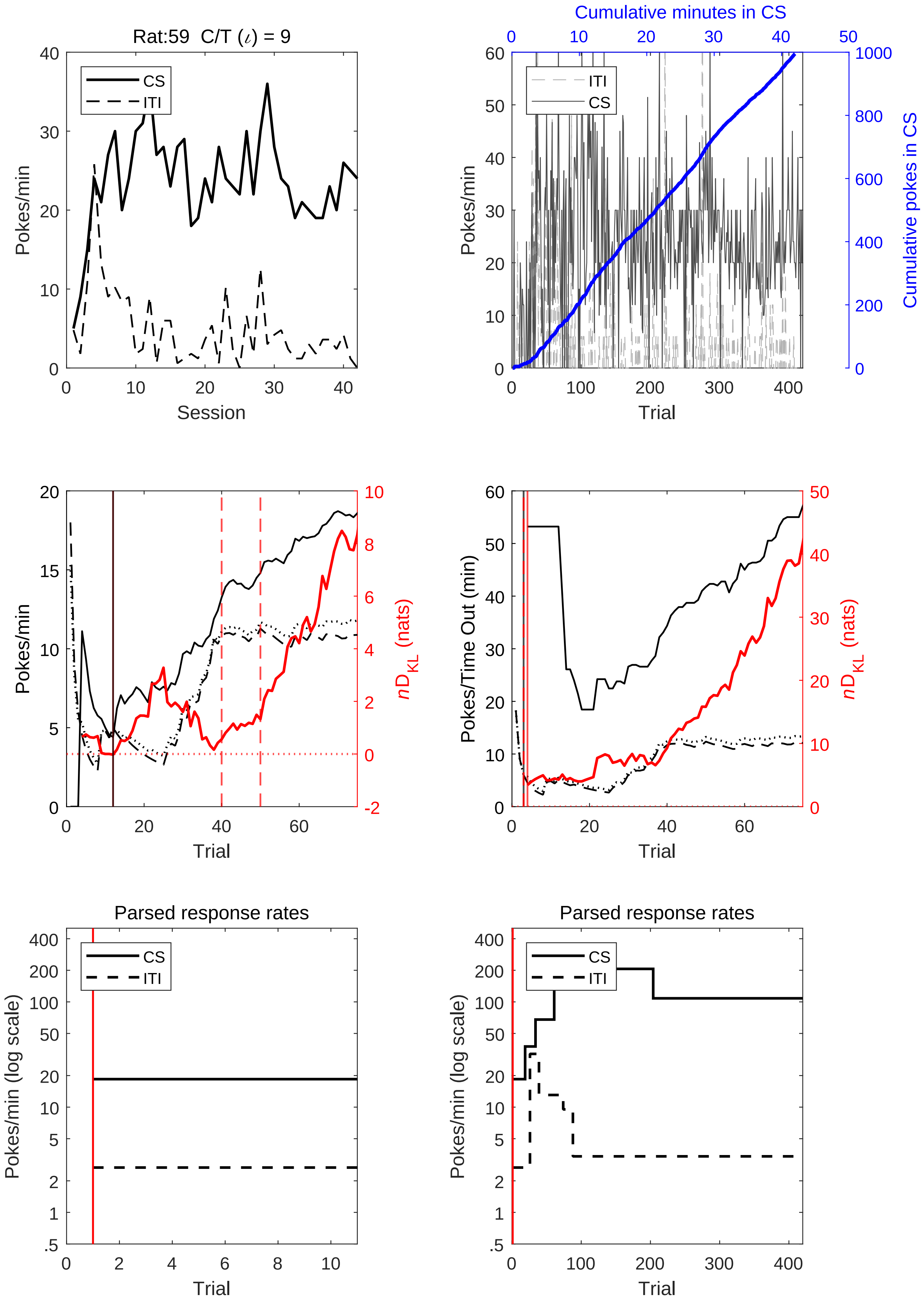
Rat:59 C/T (ι) = 9
CS
ITI
Pokes/min
Session
Cumulative minutes in CS
Pokes/min
Cumulative pokes in CS
Trial
Pokes/min
nD_KL (nats)
Trial
Pokes/Time Out (min)
nD_KL (nats)
Trial
Parsed response rates
Pokes/min (log scale)
Trial
Parsed response rates
Pokes/min (log scale)
Trial

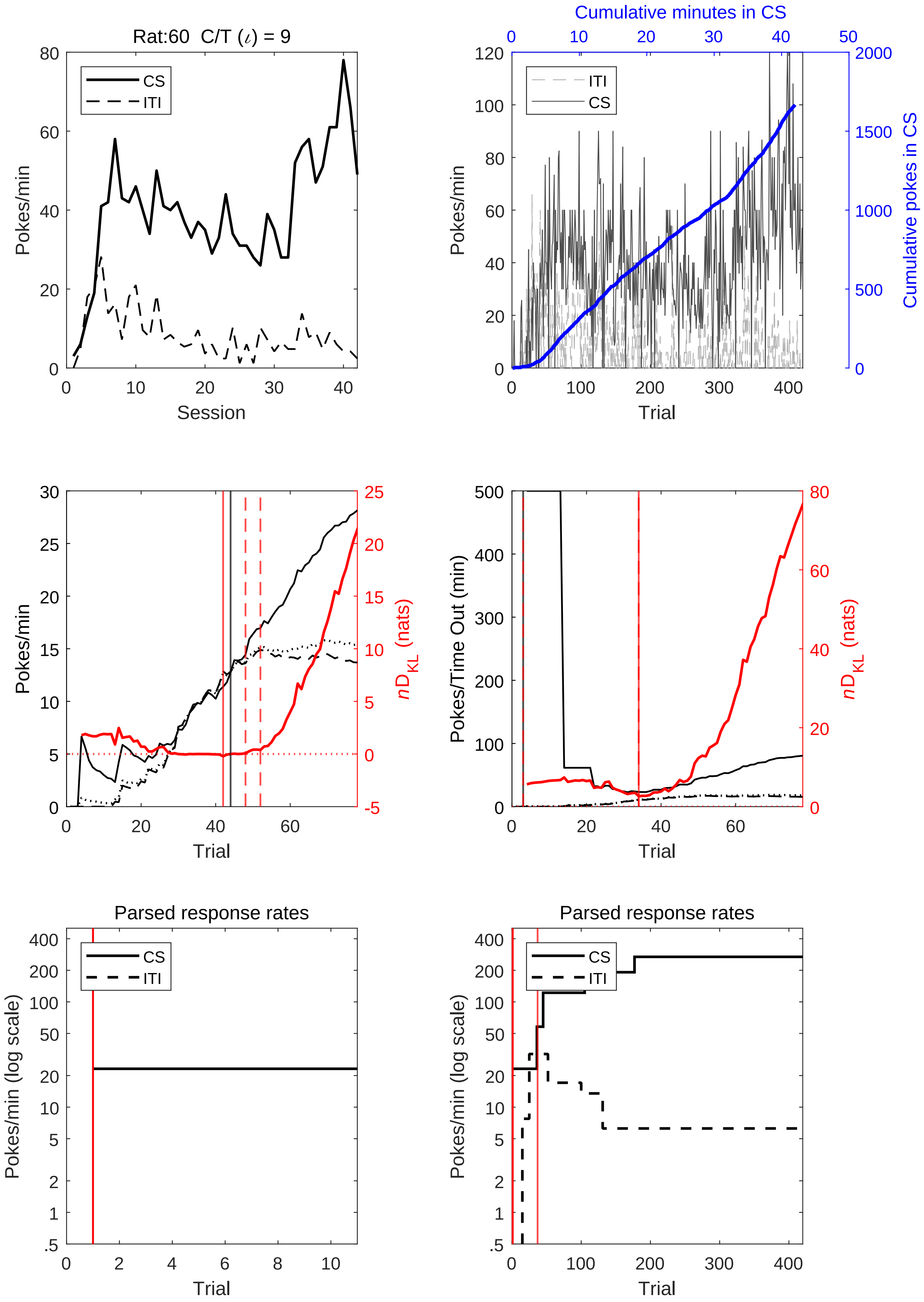
Rat:60 C/T (ι) = 9
CS
ITI
Pokes/min
Session
Cumulative minutes in CS
ITI
CS
Pokes/min
Cumulative pokes in CS
Trial
Pokes/min
nD_KL (nats)
Trial
Pokes/Time Out (min)
nD_KL (nats)
Trial
Parsed response rates
CS
ITI
Pokes/min (log scale)
Trial
Parsed response rates
CS
ITI
Pokes/min (log scale)
Trial

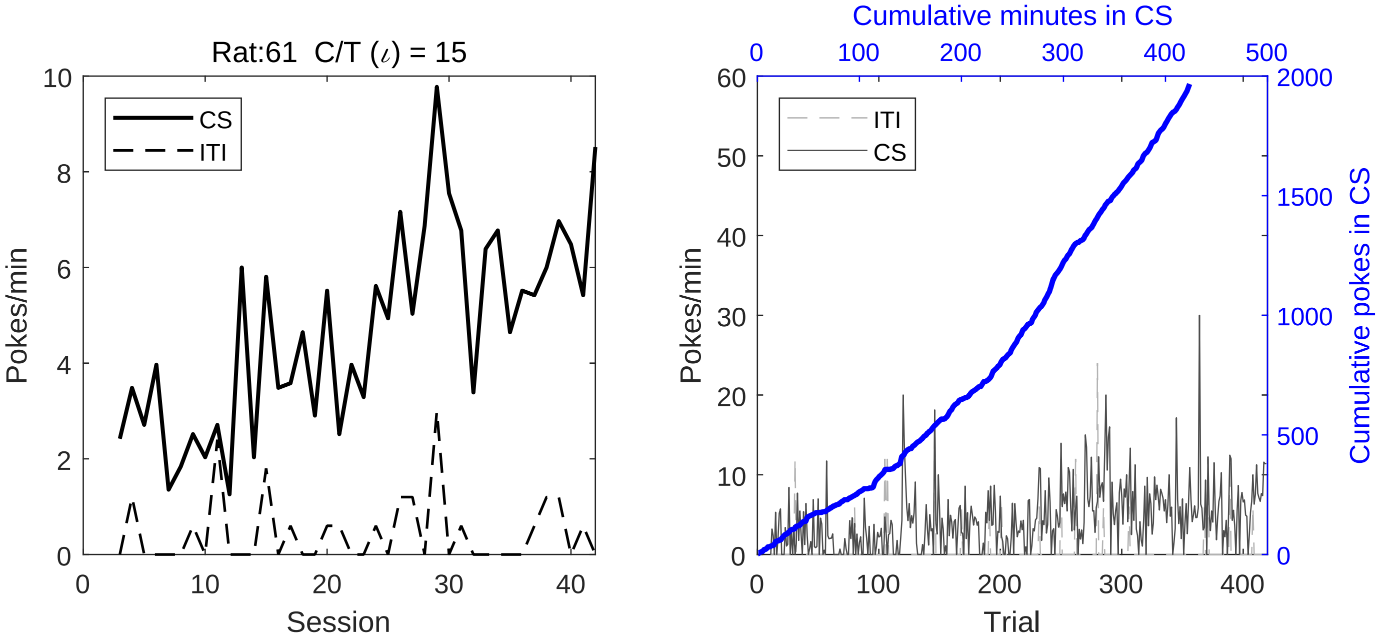

Rat:61 C/T (ι) = 15
CS
ITI
Pokes/min
Session
Cumulative minutes in CS
ITI
CS
Pokes/min
Cumulative pokes in CS
Trial

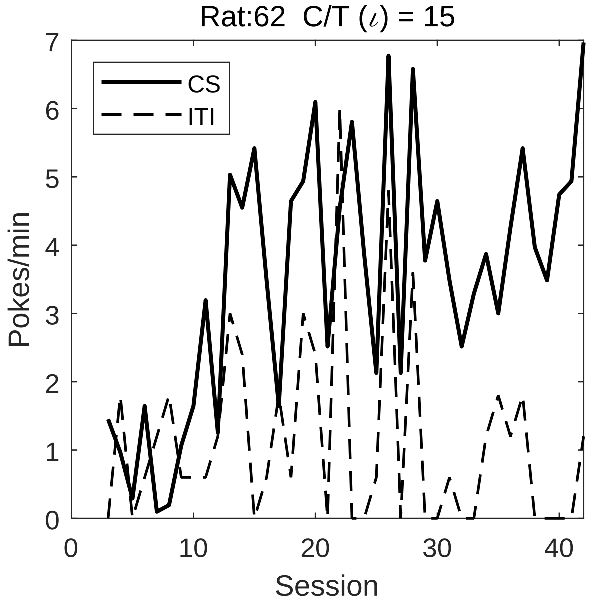
Rat:62 C/T (ι) = 15
CS
ITI
Pokes/min
Session
0
1
2
3
4
5
6
7
10
20
30
40

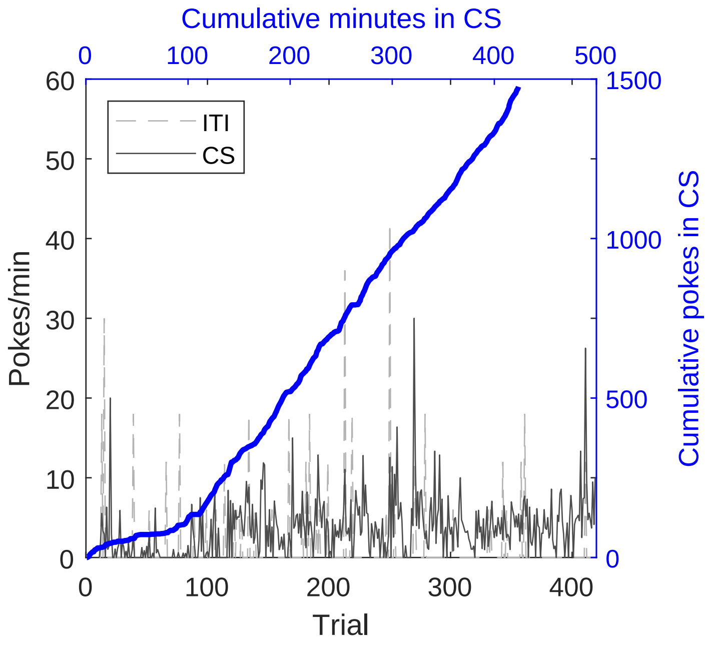
Cumulative minutes in CS
ITI
CS
Pokes/min
Trial
Cumulative pokes in CS
0
100
200
300
400
500
10
20
30
40
50
60
1000
1500

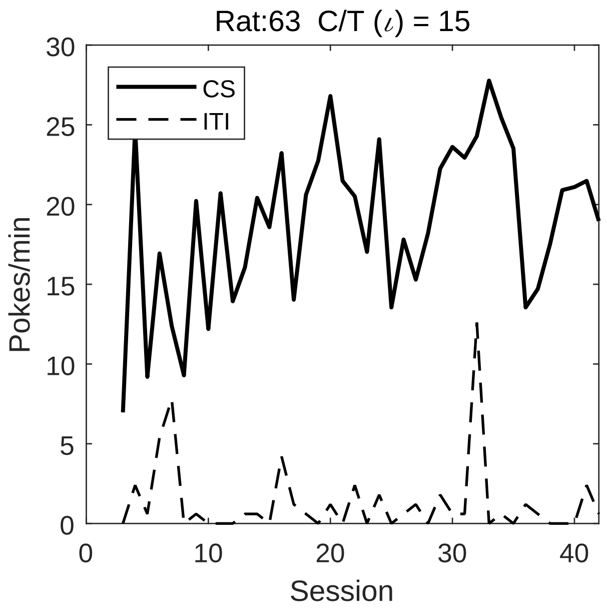

Rat:63 C/T ($\iota$) = 15
CS
ITI
Pokes/min
Session
0
5
10
15
20
25
30
0
10
20
30
40


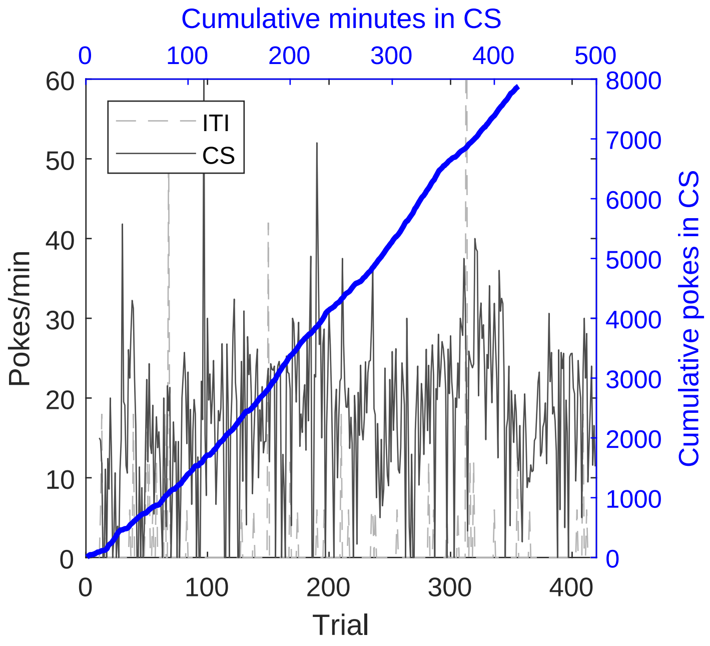

Cumulative minutes in CS
ITI
CS
Pokes/min
Trial
Cumulative pokes in CS
0
100
200
300
400
500
0
10
20
30
40
50
60
1000
2000
3000
4000
5000
6000
7000
8000

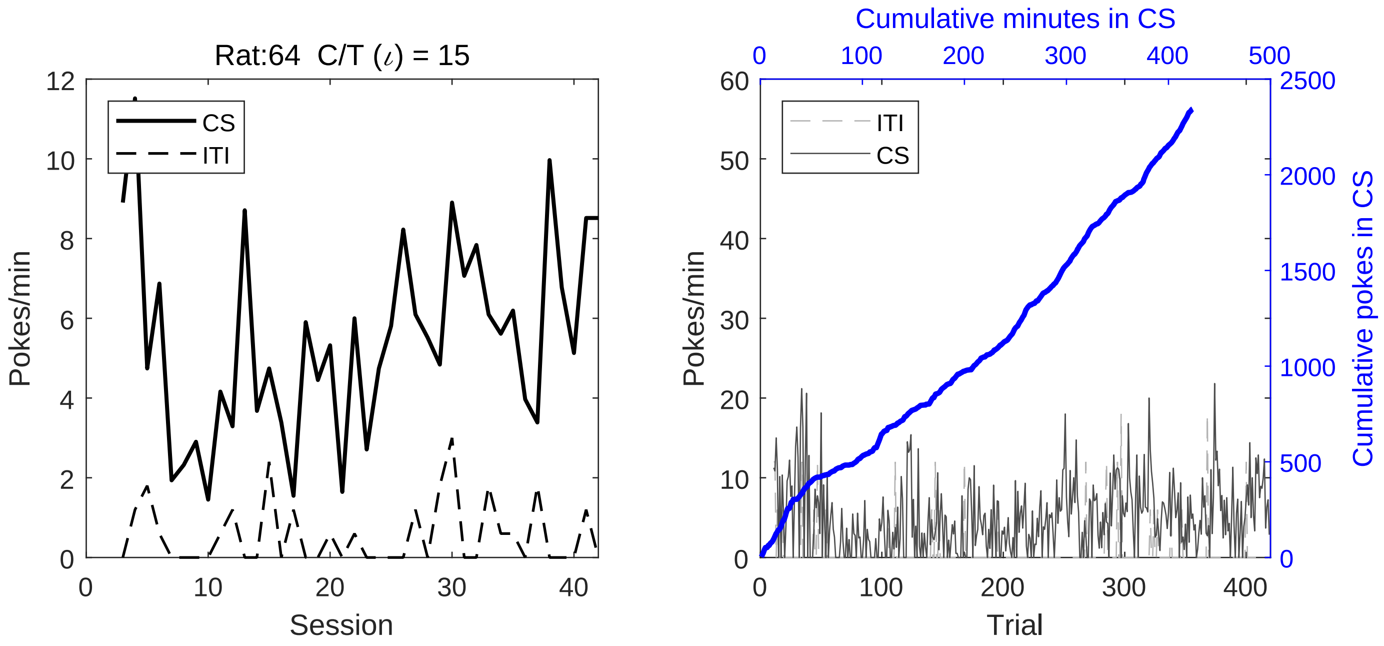
Rat:64 C/T (ι) = 15
CS
ITI
Pokes/min
Session
Cumulative minutes in CS
ITI
CS
Pokes/min
Trial
Cumulative pokes in CS

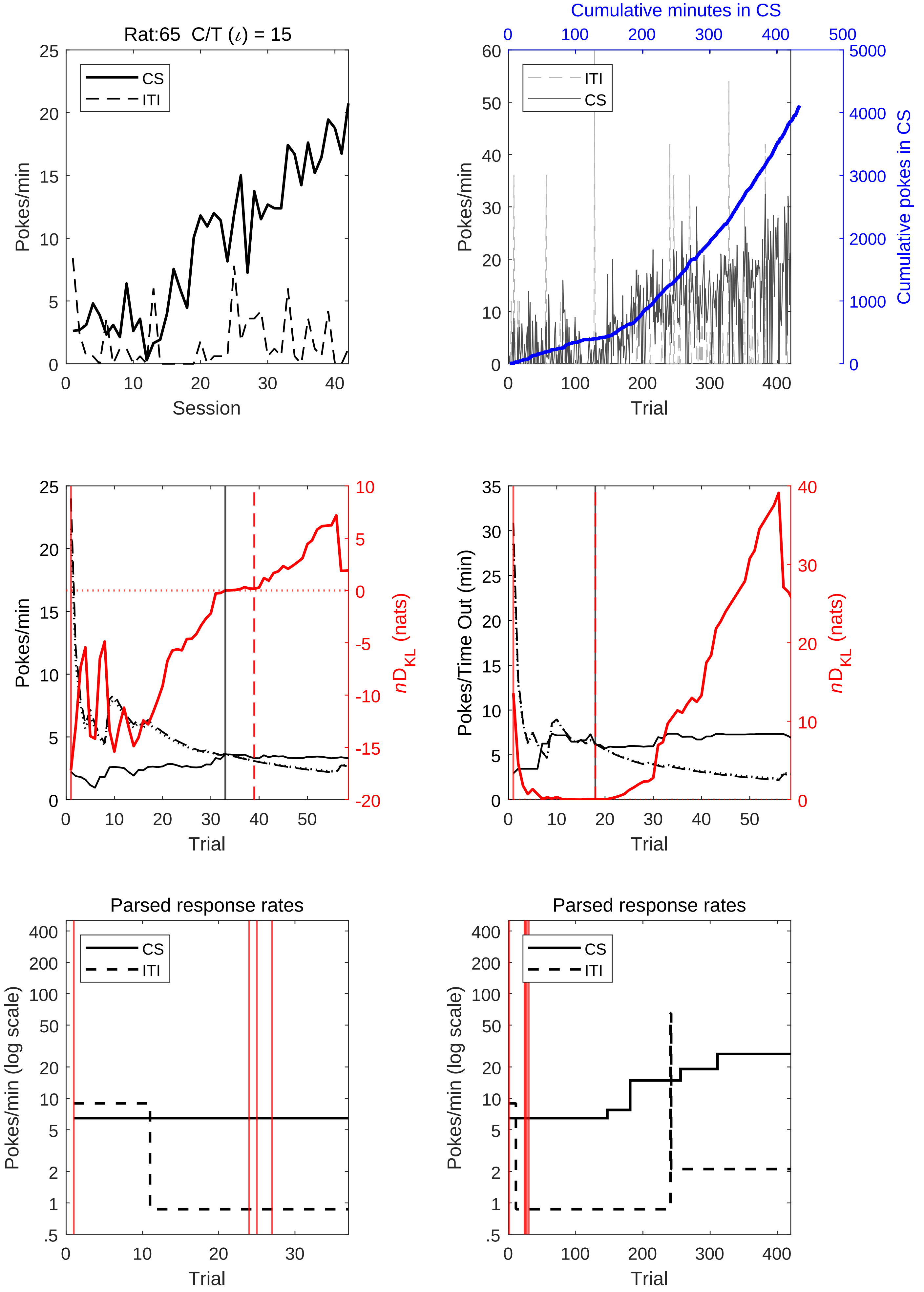

Rat:65 C/T (ι) = 15
Pokes/min
Session
CS
ITI
Cumulative minutes in CS
Cumulative pokes in CS
Trial
nD_KL (nats)
Pokes/Time Out (min)
Parsed response rates
Pokes/min (log scale)

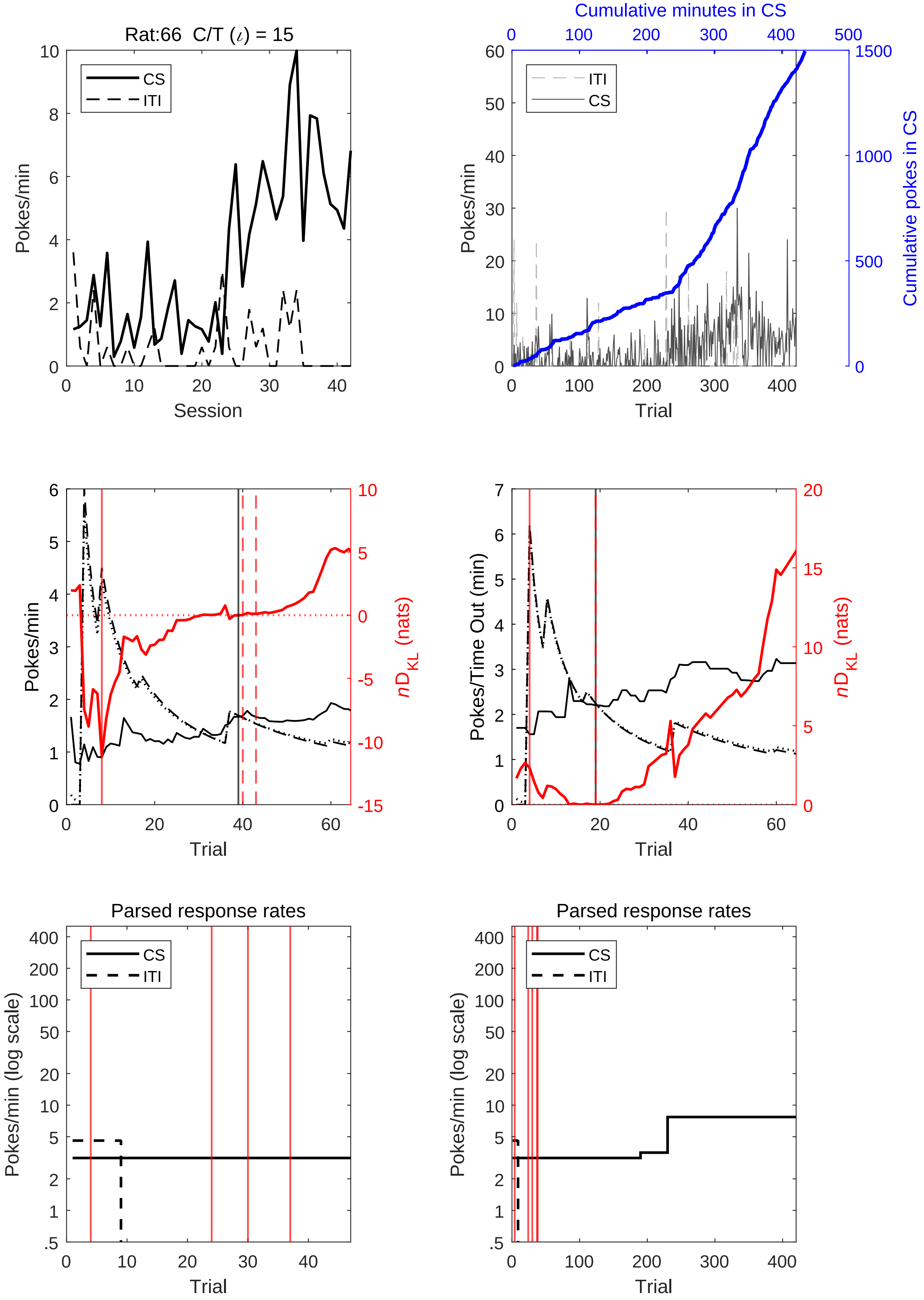
Rat:66 C/T (ι) = 15
CS
ITI
Pokes/min
Session
Cumulative minutes in CS
Cumulative pokes in CS
Pokes/min
Trial
Pokes/min
nD_KL (nats)
Trial
Pokes/Time Out (min)
nD_KL (nats)
Trial
Parsed response rates
Pokes/min (log scale)
Trial
Parsed response rates
Pokes/min (log scale)
Trial

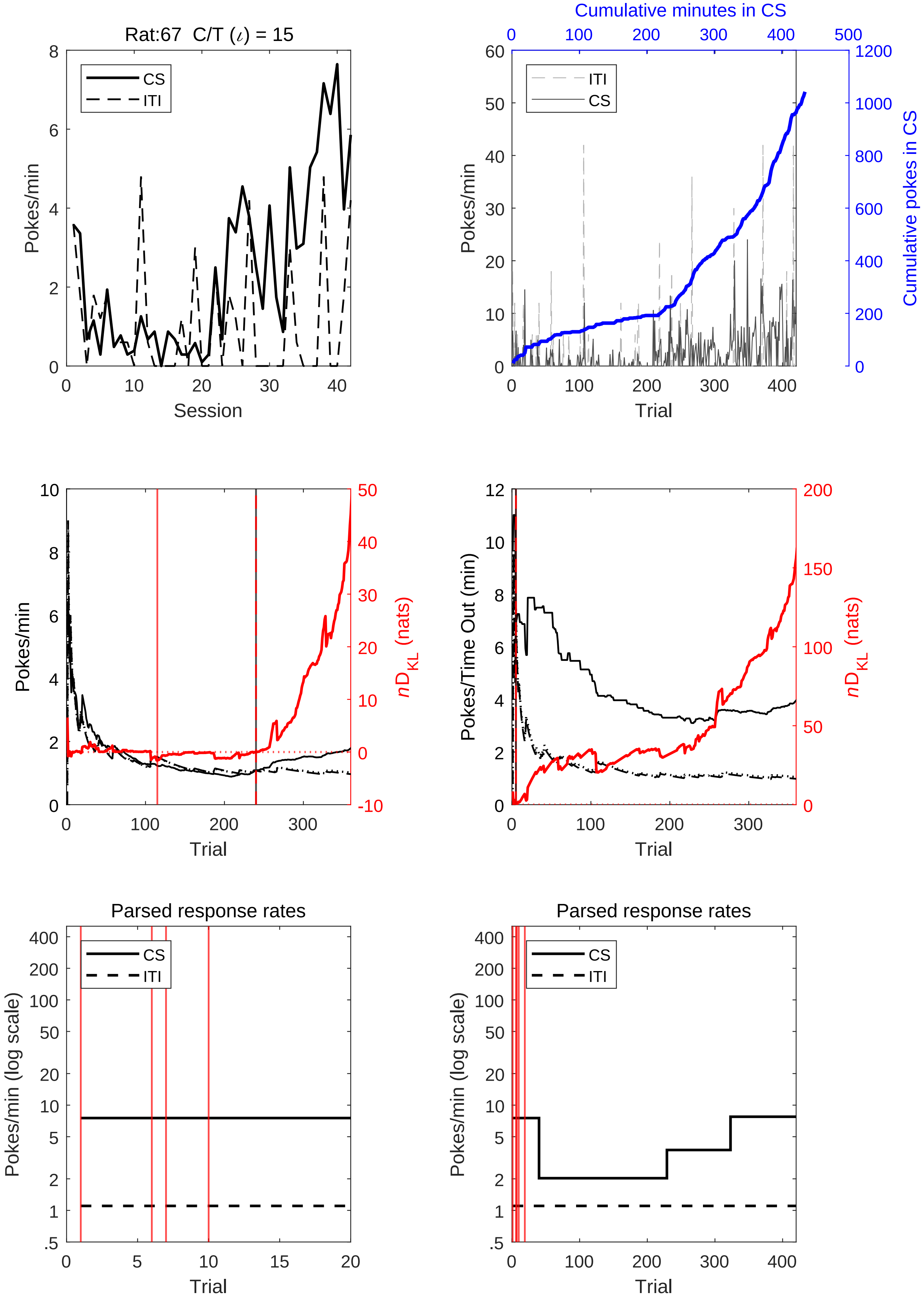
Rat:67 C/T (ι) = 15
CS
ITI
Pokes/min
Session
Cumulative minutes in CS
Cumulative pokes in CS
ITI
CS
Pokes/min
Trial
Pokes/min
$nD_{KL}$ (nats)
Trial
Pokes/Time Out (min)
$nD_{KL}$ (nats)
Trial
Parsed response rates
CS
ITI
Pokes/min (log scale)
Trial
Parsed response rates
CS
ITI
Pokes/min (log scale)
Trial

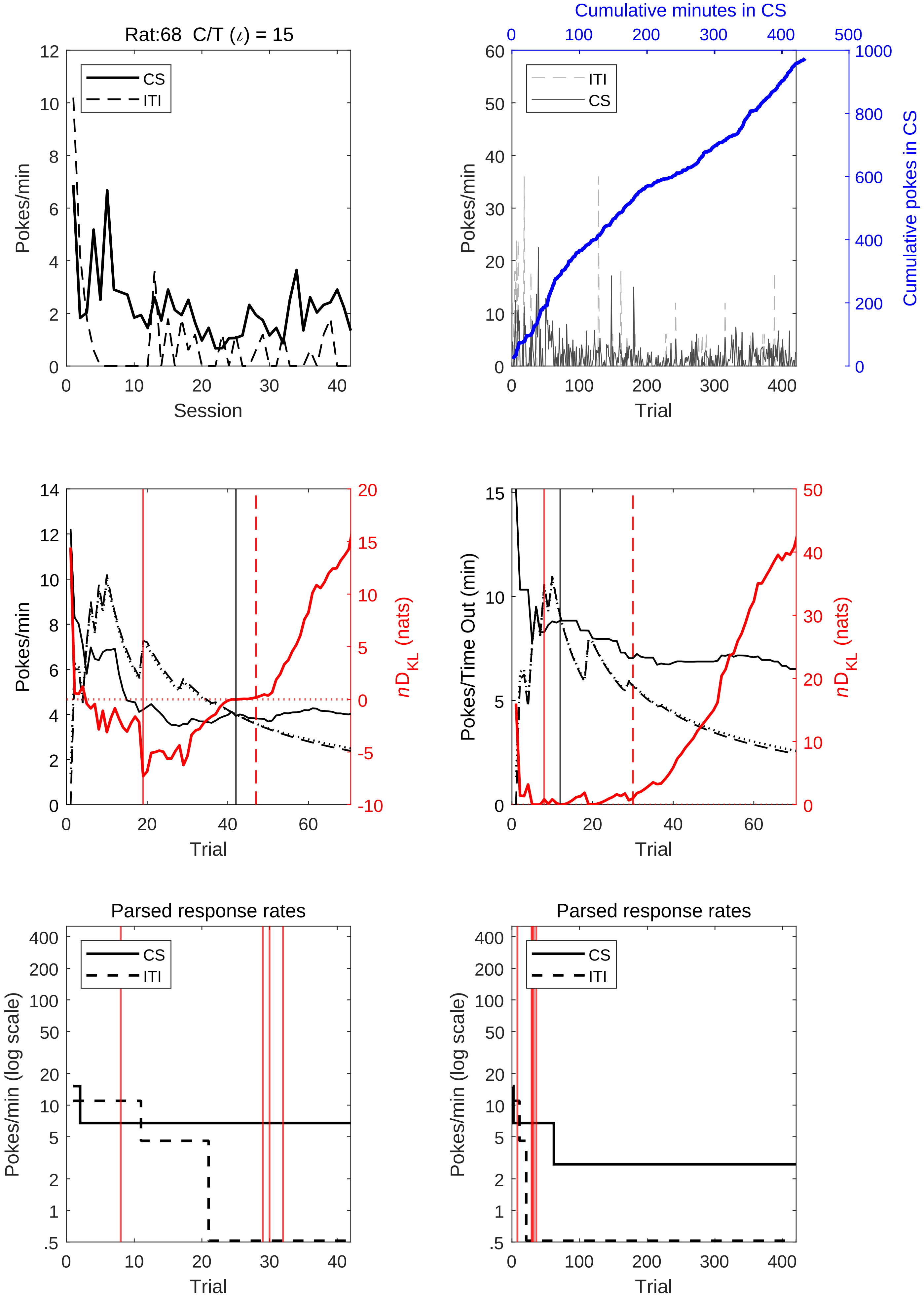

Rat:68 C/T (ι) = 15
CS
ITI
Pokes/min
Session
Cumulative minutes in CS
Cumulative pokes in CS
Trial
Pokes/Time Out (min)
$nD_{KL}$ (nats)
Parsed response rates
Pokes/min (log scale)

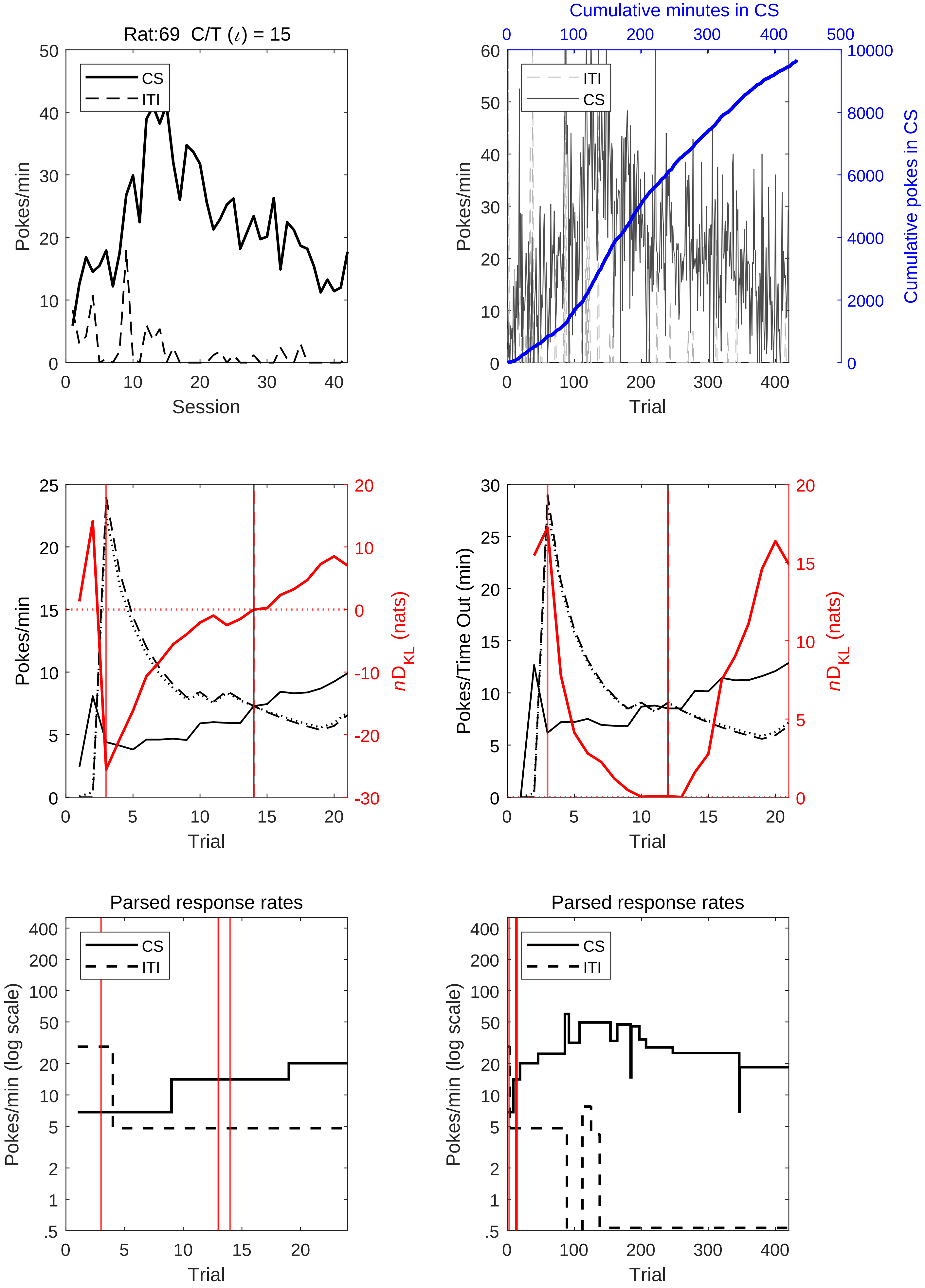

Rat:69 C/T (ι) = 15
CS
ITI
Pokes/min
Session
Cumulative minutes in CS
Cumulative pokes in CS
Trial
nD_KL (nats)
Pokes/Time Out (min)
Parsed response rates
Pokes/min (log scale)

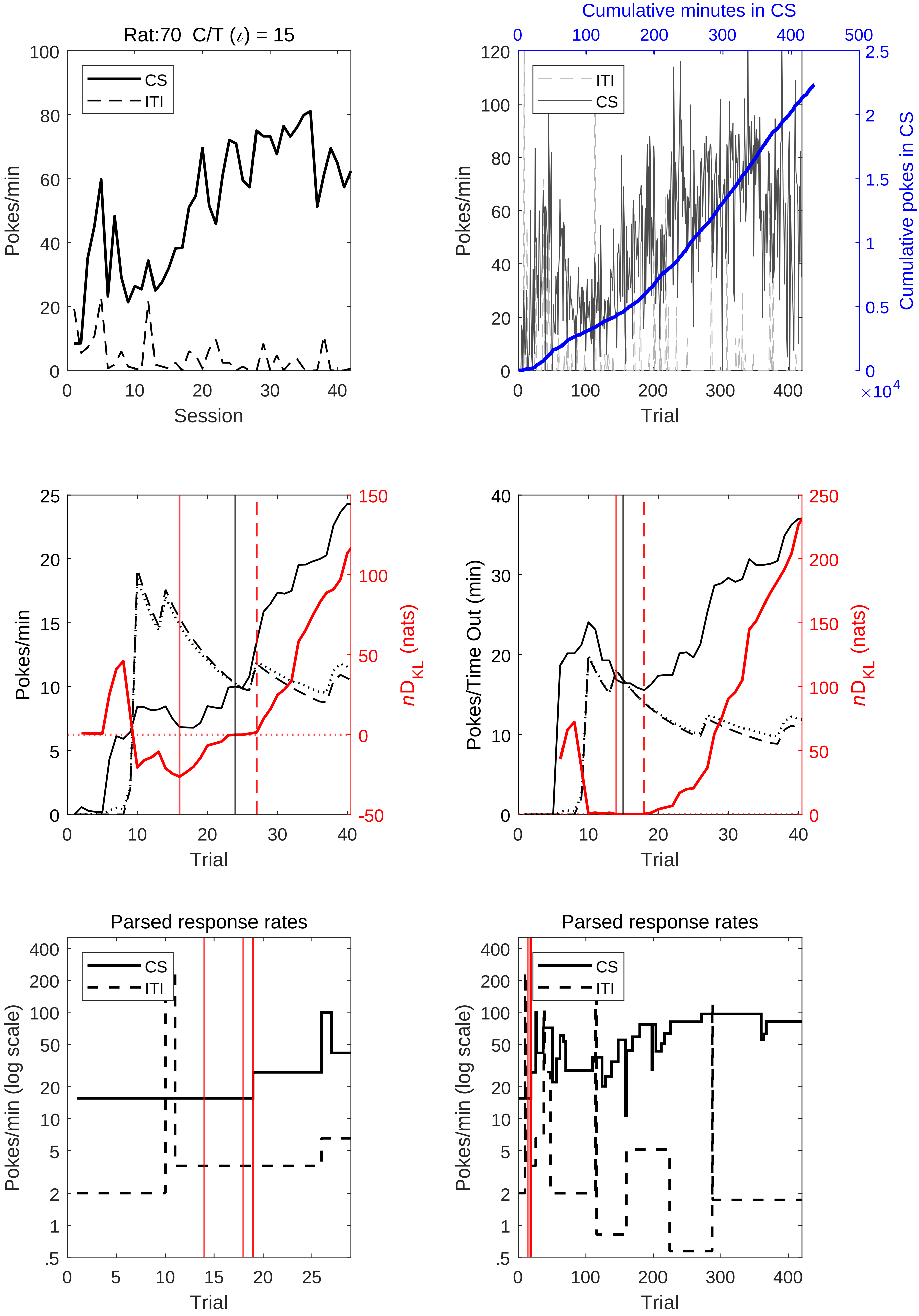
Rat:70 C/T (ι) = 15
CS
ITI
Pokes/min
Session
Cumulative minutes in CS
Cumulative pokes in CS
×10^4
Trial
nD_KL (nats)
Pokes/Time Out (min)
Parsed response rates
Pokes/min (log scale)

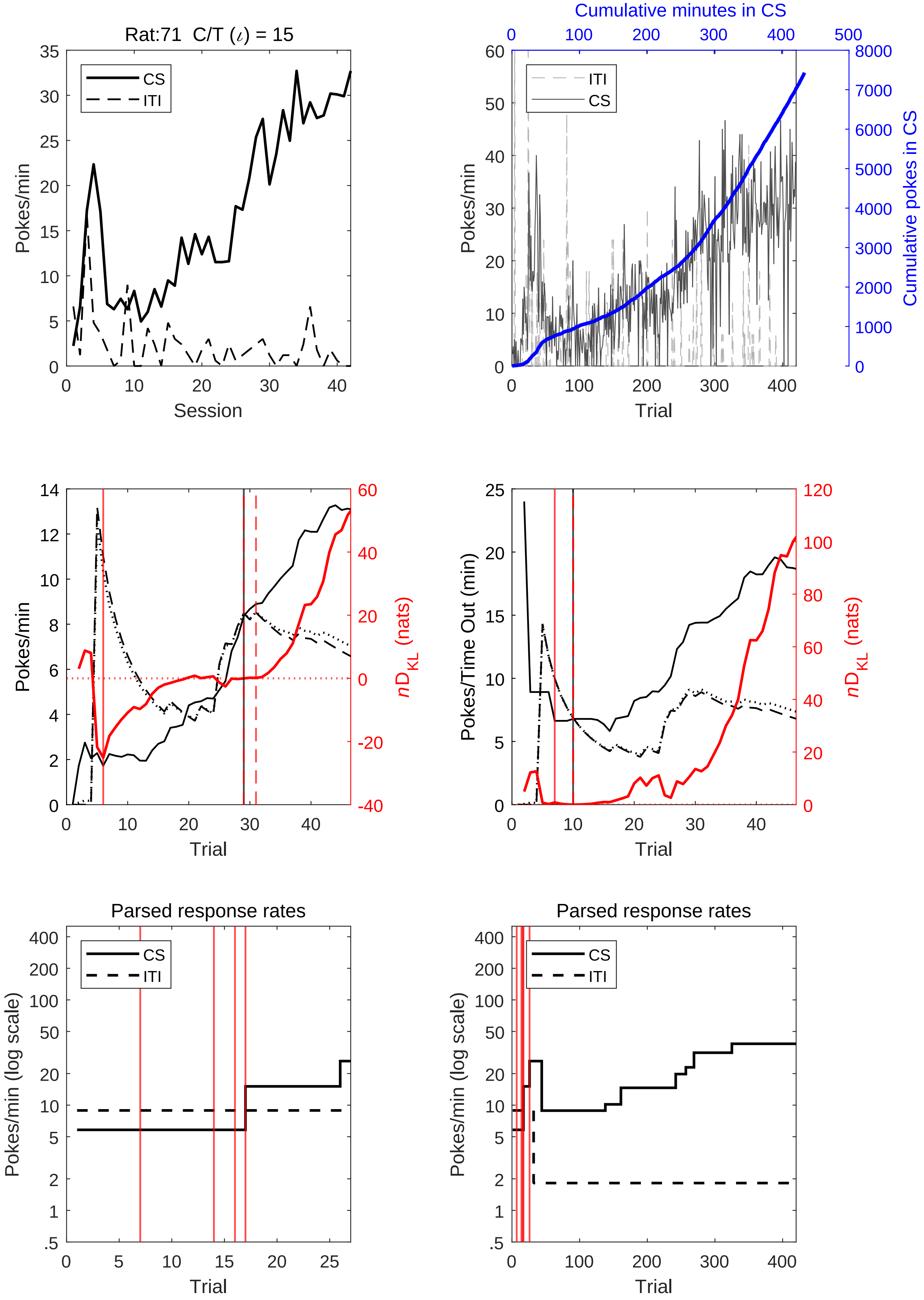
Rat:71 C/T (ι) = 15
Pokes/min
Session
CS
ITI
Cumulative minutes in CS
Cumulative pokes in CS
Trial
nD_KL (nats)
Pokes/Time Out (min)
Parsed response rates
Pokes/min (log scale)

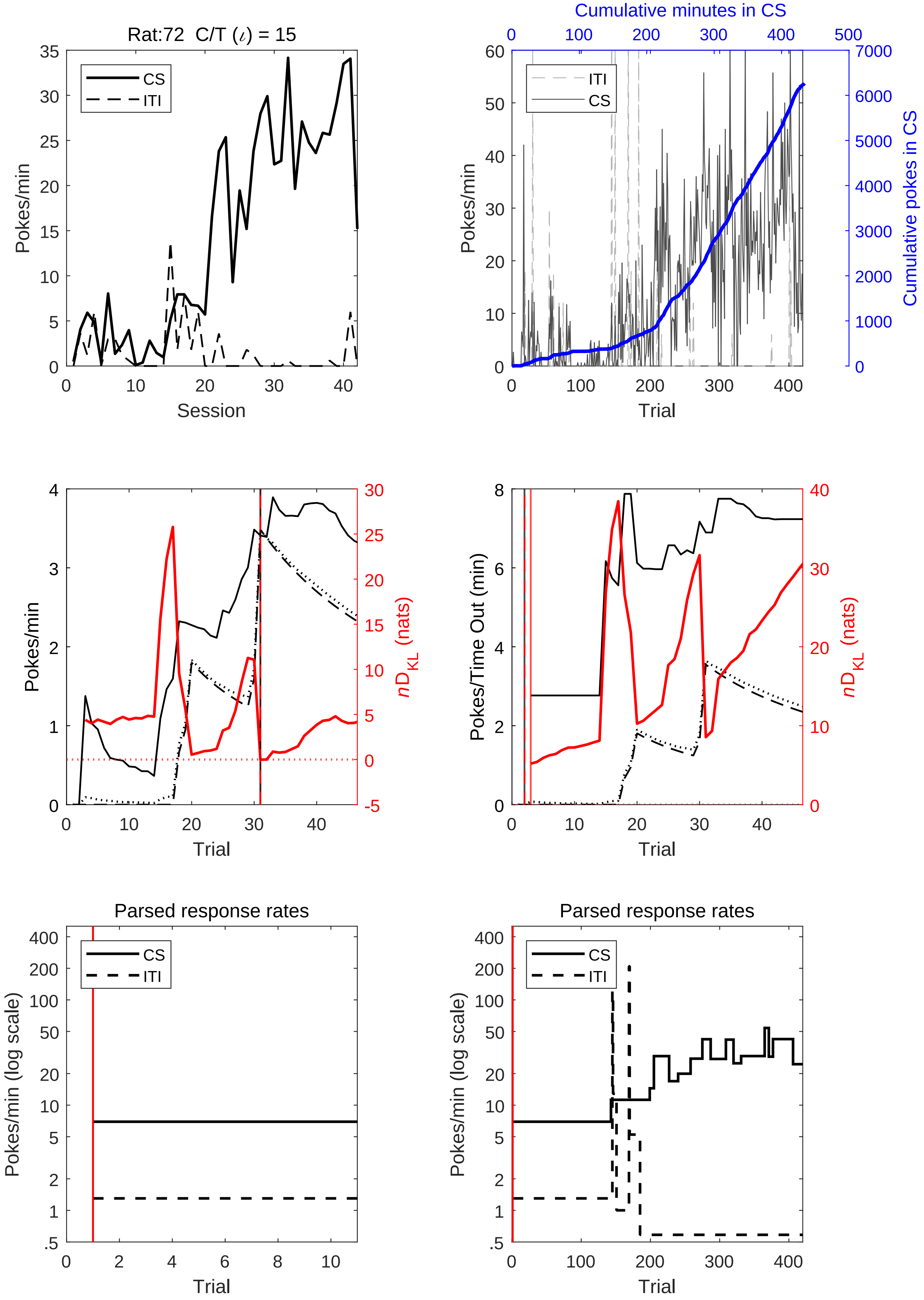
Rat:72 C/T (ι) = 15
CS
ITI
Pokes/min
Session
Cumulative minutes in CS
Cumulative pokes in CS
Trial
nD_KL (nats)
Pokes/Time Out (min)
Parsed response rates
Pokes/min (log scale)

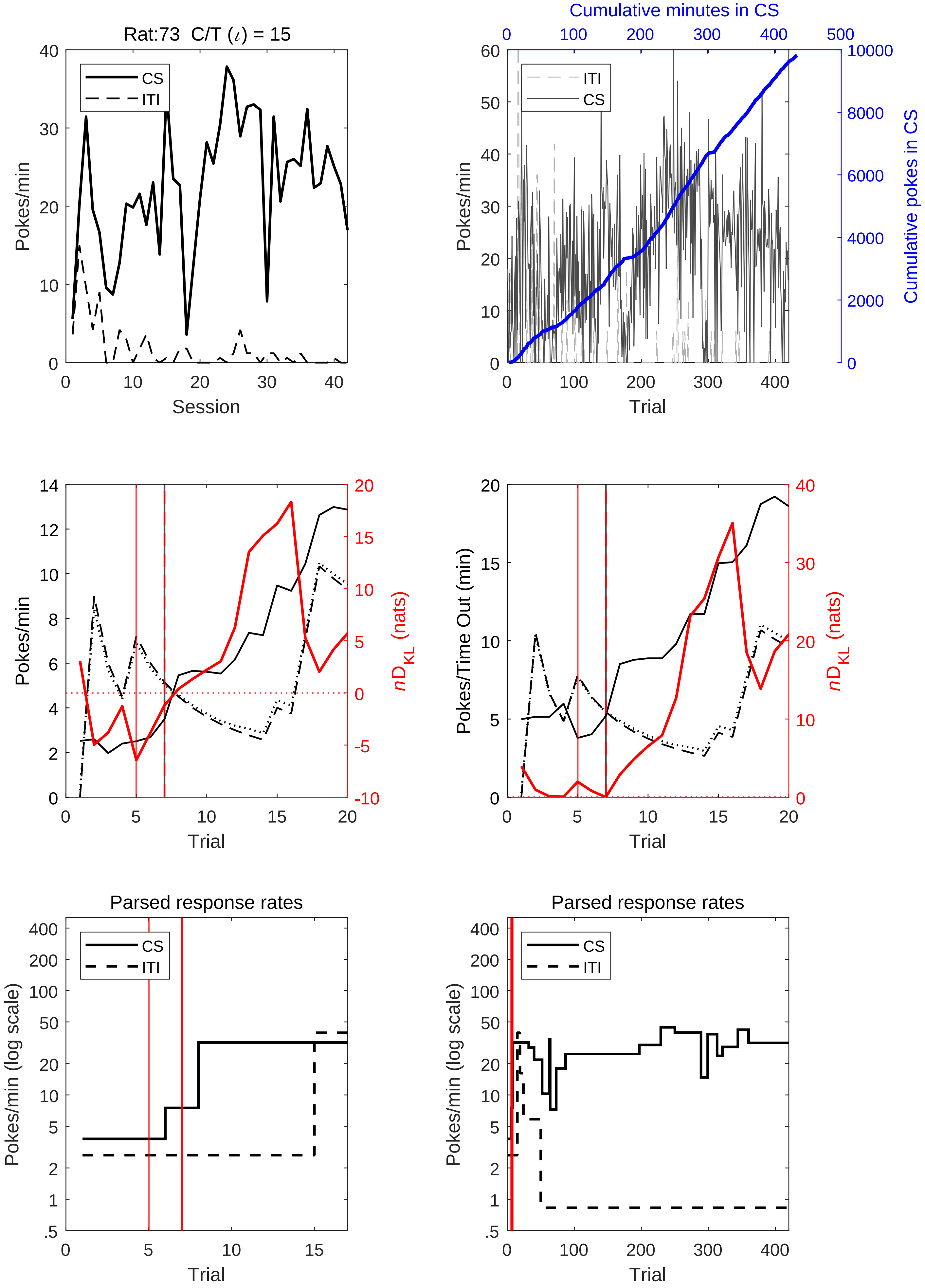
Rat:73 C/T (ι) = 15
CS
ITI
Pokes/min
Session
Cumulative minutes in CS
Cumulative pokes in CS
Trial
Pokes/Time Out (min)
nD_KL (nats)
Parsed response rates
Pokes/min (log scale)

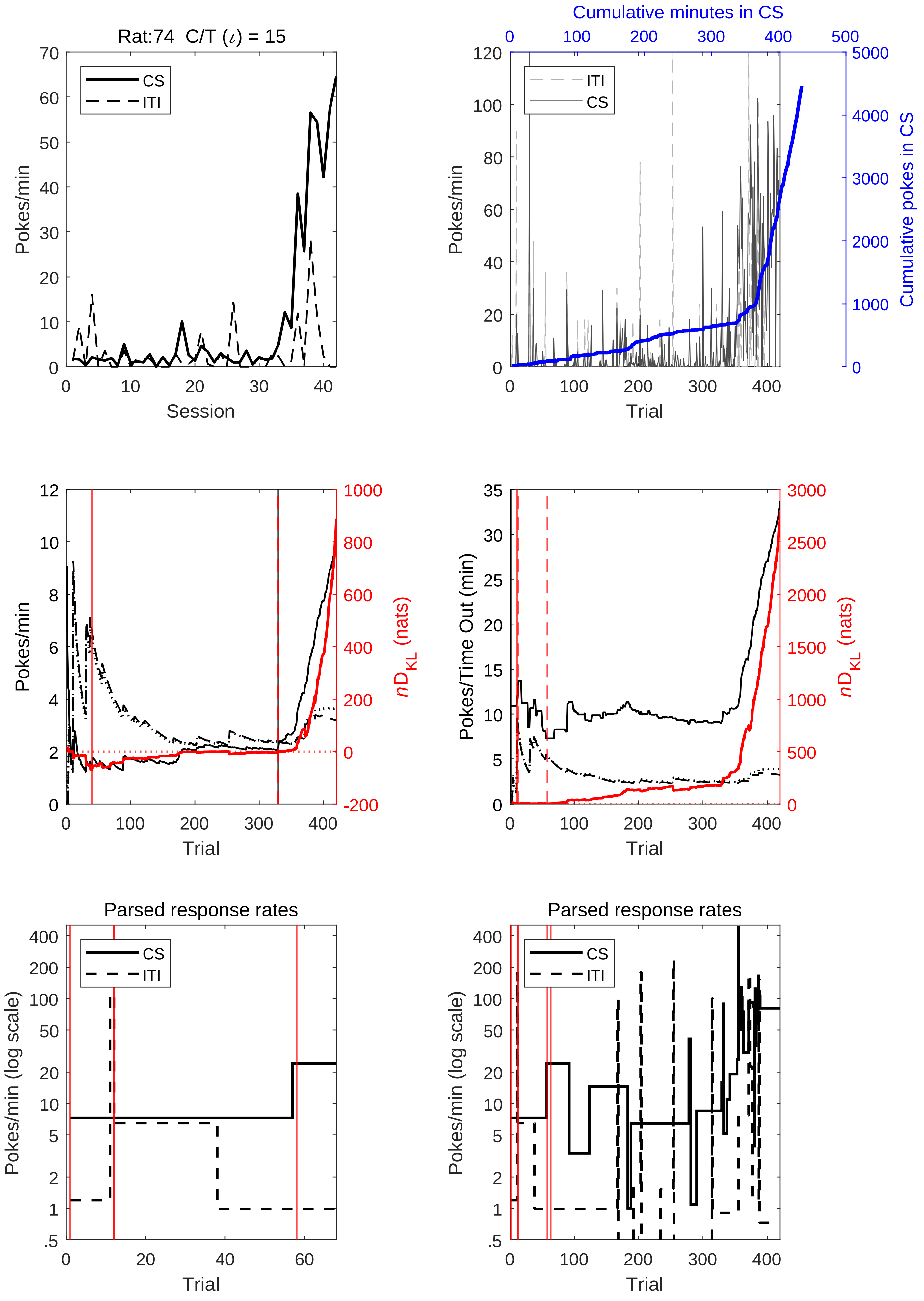
Rat:74 C/T (ι) = 15
CS
ITI
Pokes/min
Session
Cumulative minutes in CS
Cumulative pokes in CS
Trial
nD_KL (nats)
Pokes/Time Out (min)
Parsed response rates
Pokes/min (log scale)

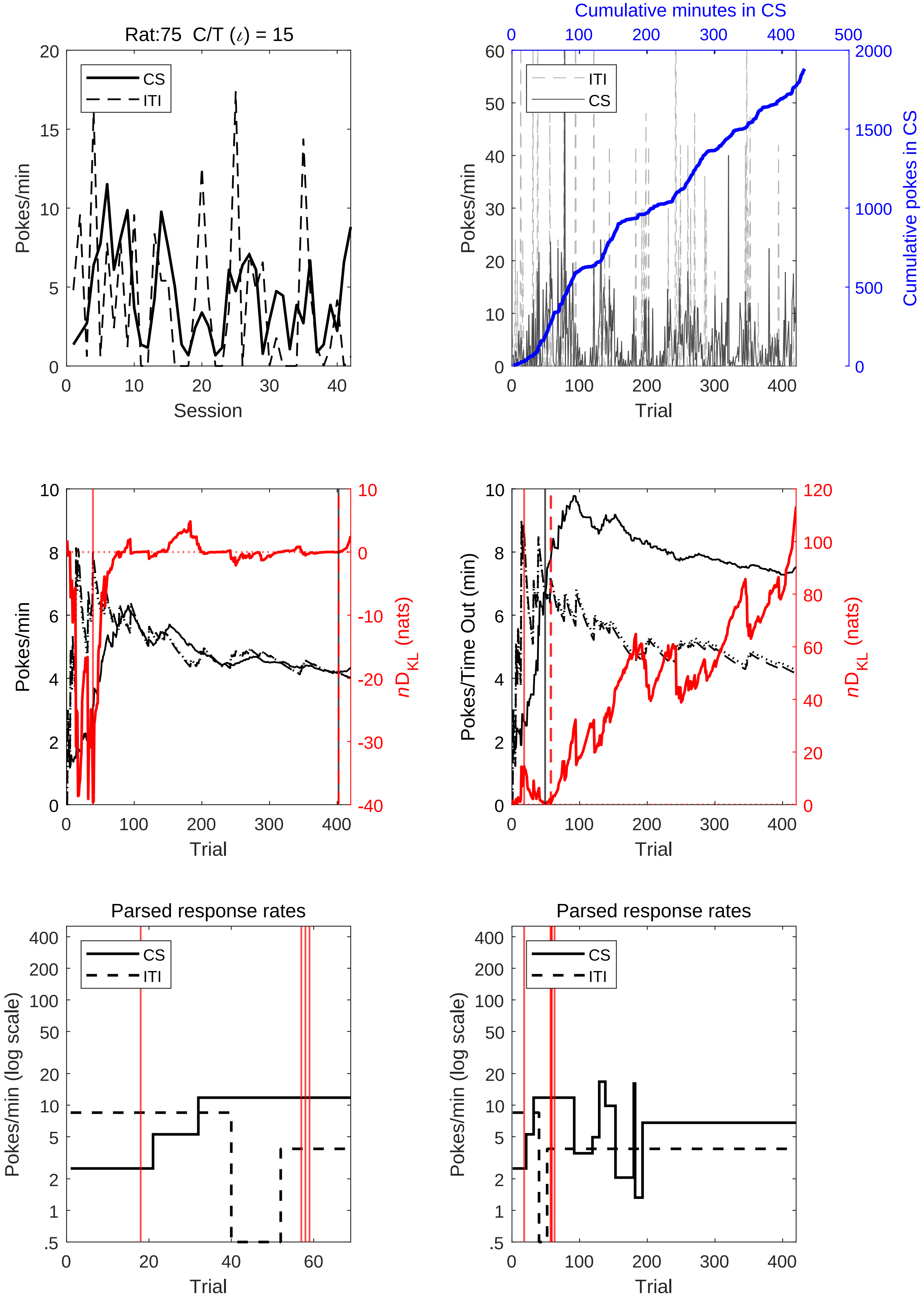
Rat:75 C/T (ι) = 15
CS
ITI
Pokes/min
Session
Cumulative minutes in CS
Cumulative pokes in CS
Trial
nD_KL (nats)
Pokes/Time Out (min)
Parsed response rates
Pokes/min (log scale)

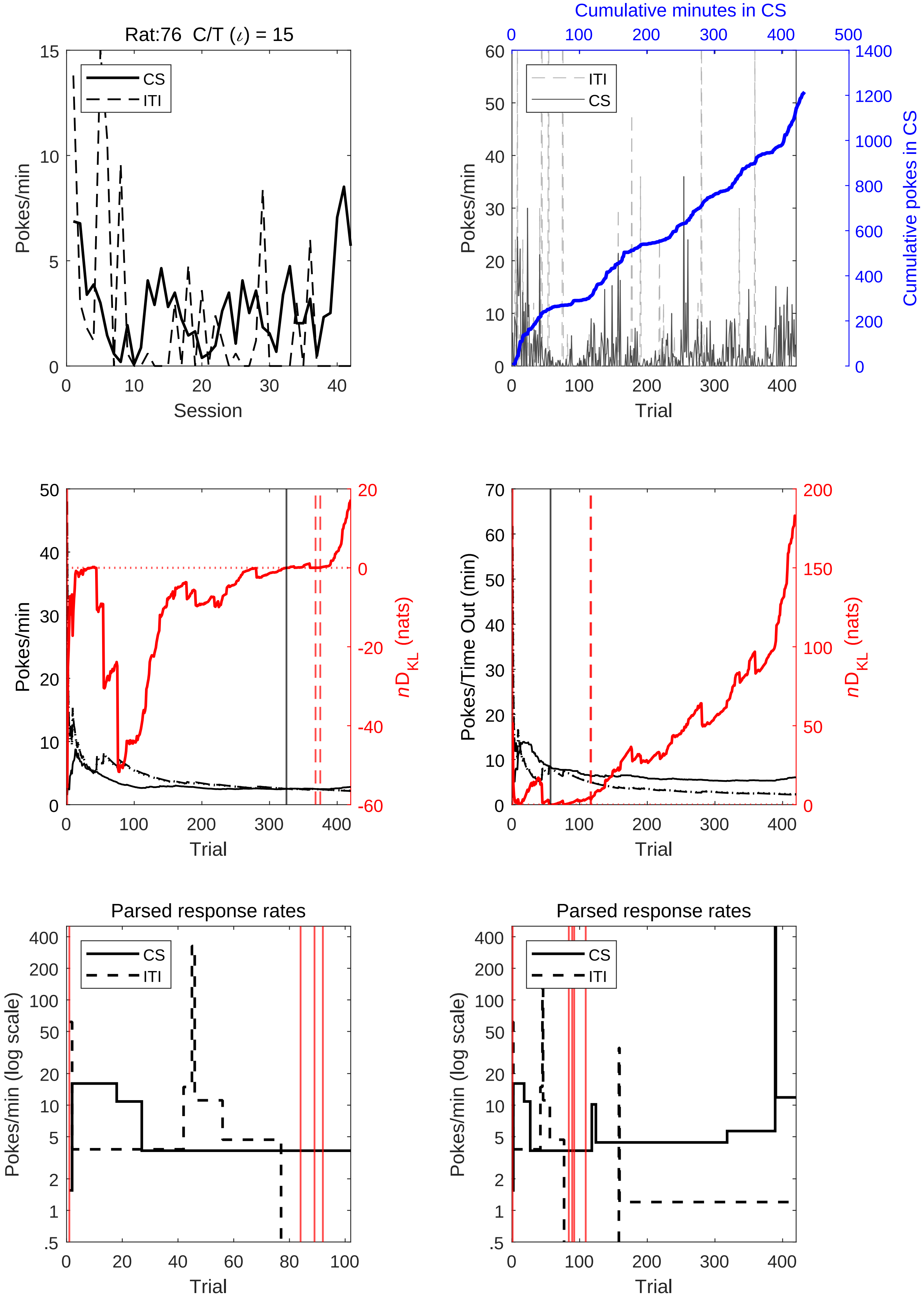

Rat:76 C/T ($\iota$) = 15
CS
ITI
Pokes/min
Session
0
5
10
15
20
30
40
Cumulative minutes in CS
0
100
200
300
400
500
ITI
CS
Pokes/min
Trial
Cumulative pokes in CS
1400
1200
1000
800
600
400
200
60
50
Pokes/min
$nD_{KL}$ (nats)
-20
-40
-60
Pokes/Time Out (min)
70
150
Parsed response rates
Pokes/min (log scale)
.5
1
2
5
10
20
50
100
200
400
80

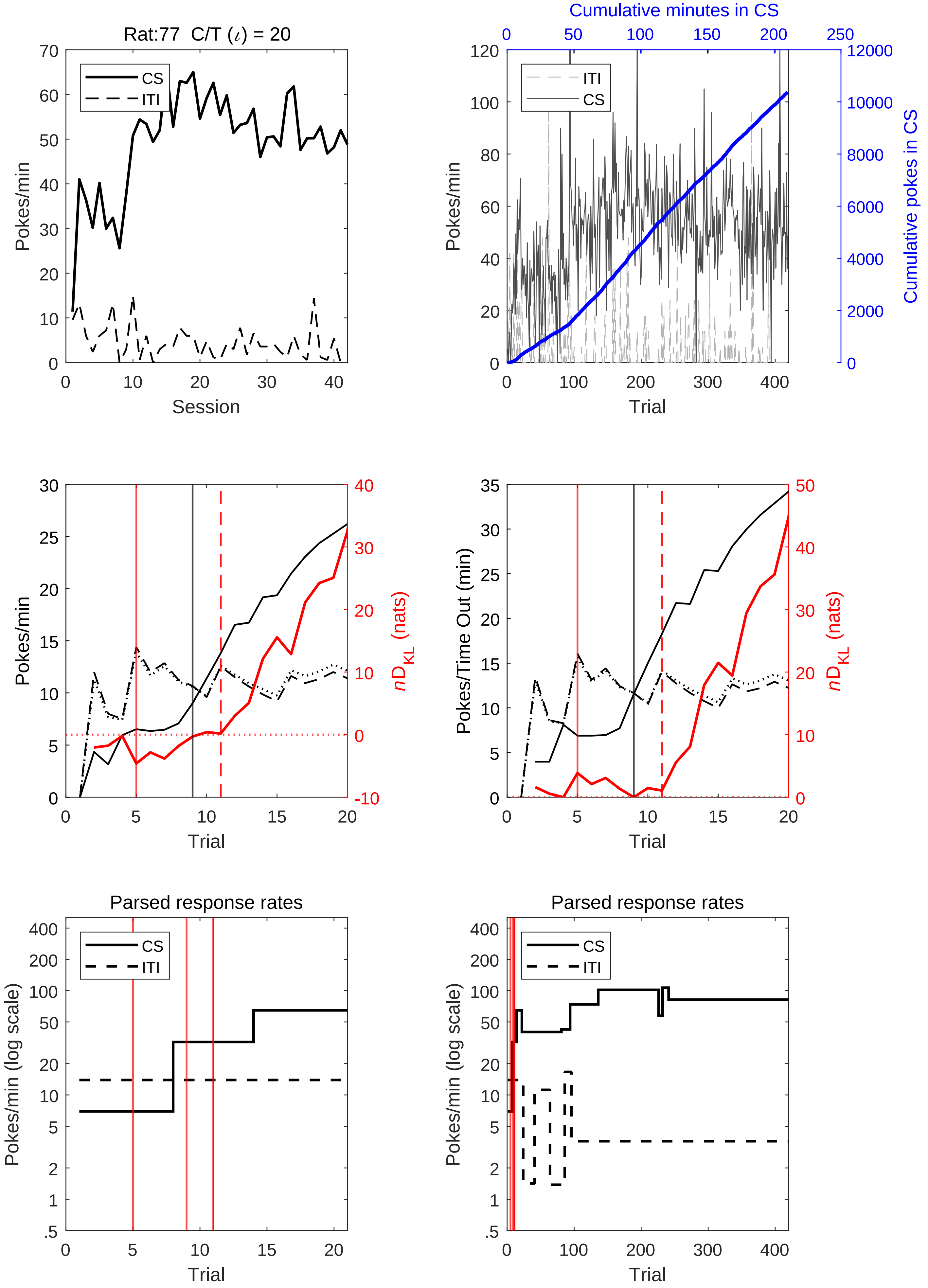
Rat:77 C/T (ι) = 20
CS
ITI
Pokes/min
Session
Cumulative minutes in CS
Cumulative pokes in CS
Trial
nD_KL (nats)
Pokes/Time Out (min)
Parsed response rates
Pokes/min (log scale)

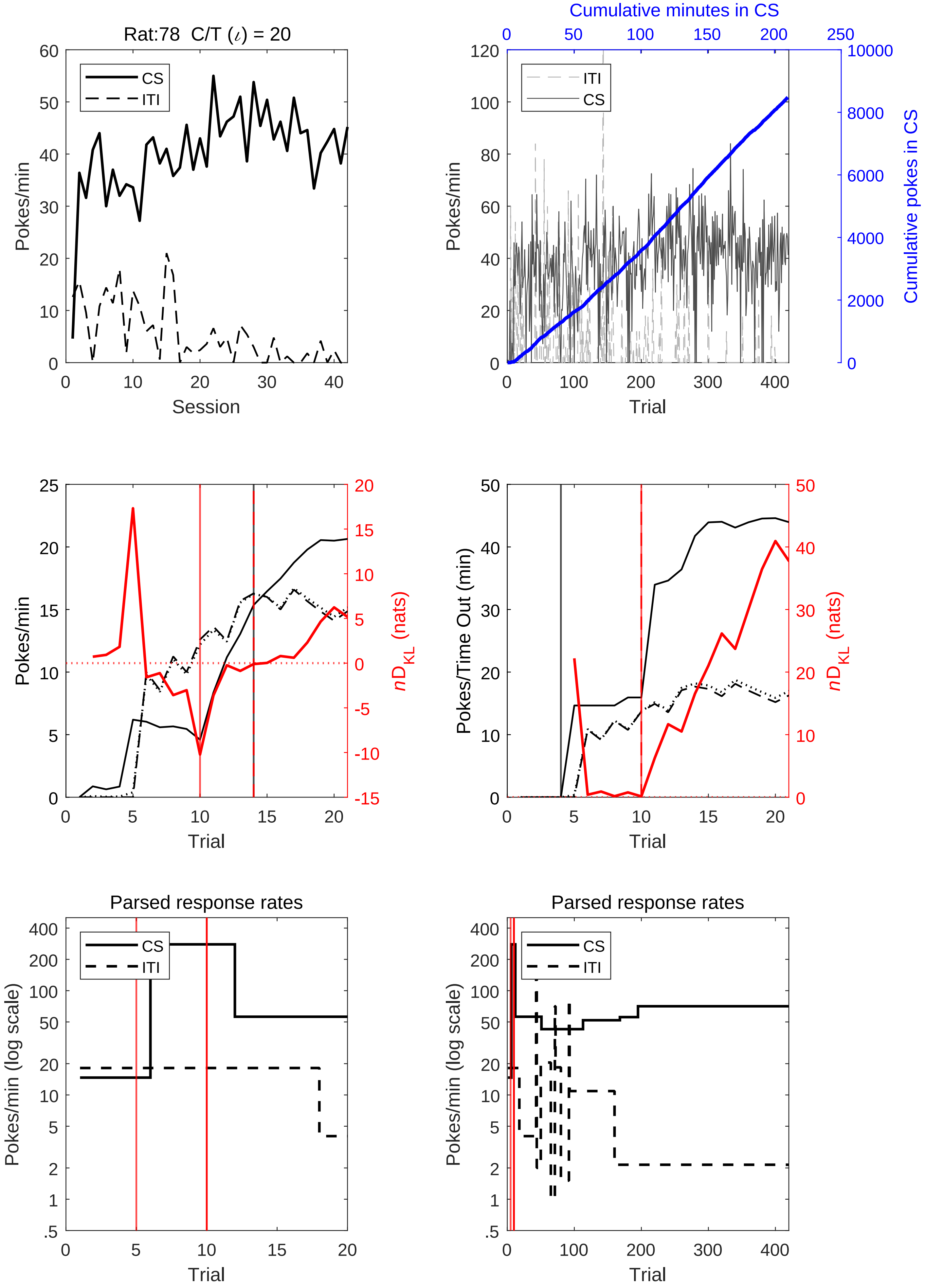
Rat:78 C/T ($\iota$) = 20
CS
ITI
Pokes/min
Session
Cumulative minutes in CS
ITI
CS
Pokes/min
Cumulative pokes in CS
Trial
Pokes/min
$nD_{KL}$ (nats)
Trial
Pokes/Time Out (min)
$nD_{KL}$ (nats)
Trial
Parsed response rates
CS
ITI
Pokes/min (log scale)
Trial
Parsed response rates
CS
ITI
Pokes/min (log scale)
Trial

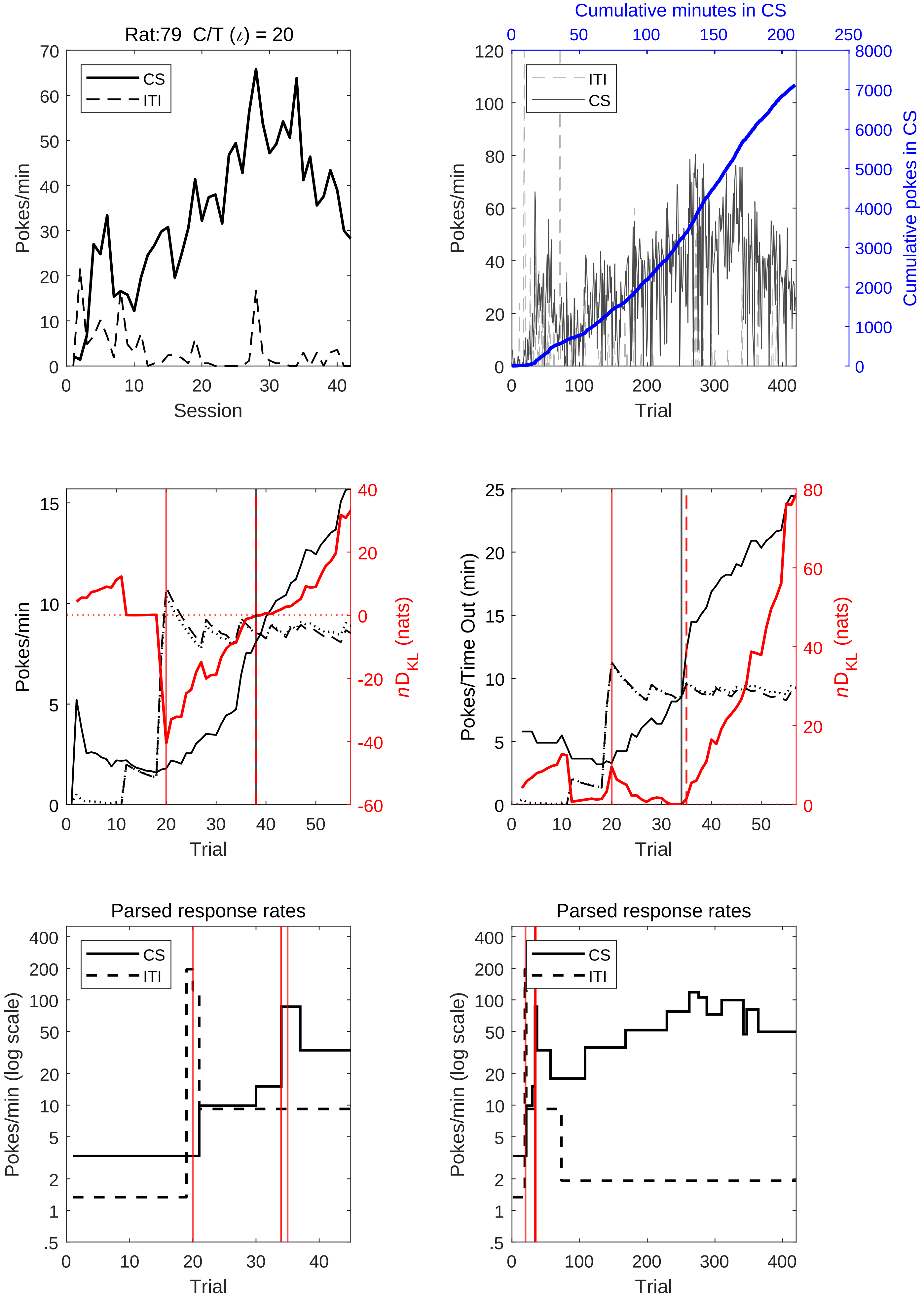
Rat:79 C/T (ι) = 20
Pokes/min
Session
CS
ITI
Cumulative minutes in CS
Cumulative pokes in CS
Trial
nD_KL (nats)
Pokes/Time Out (min)
Parsed response rates
Pokes/min (log scale)

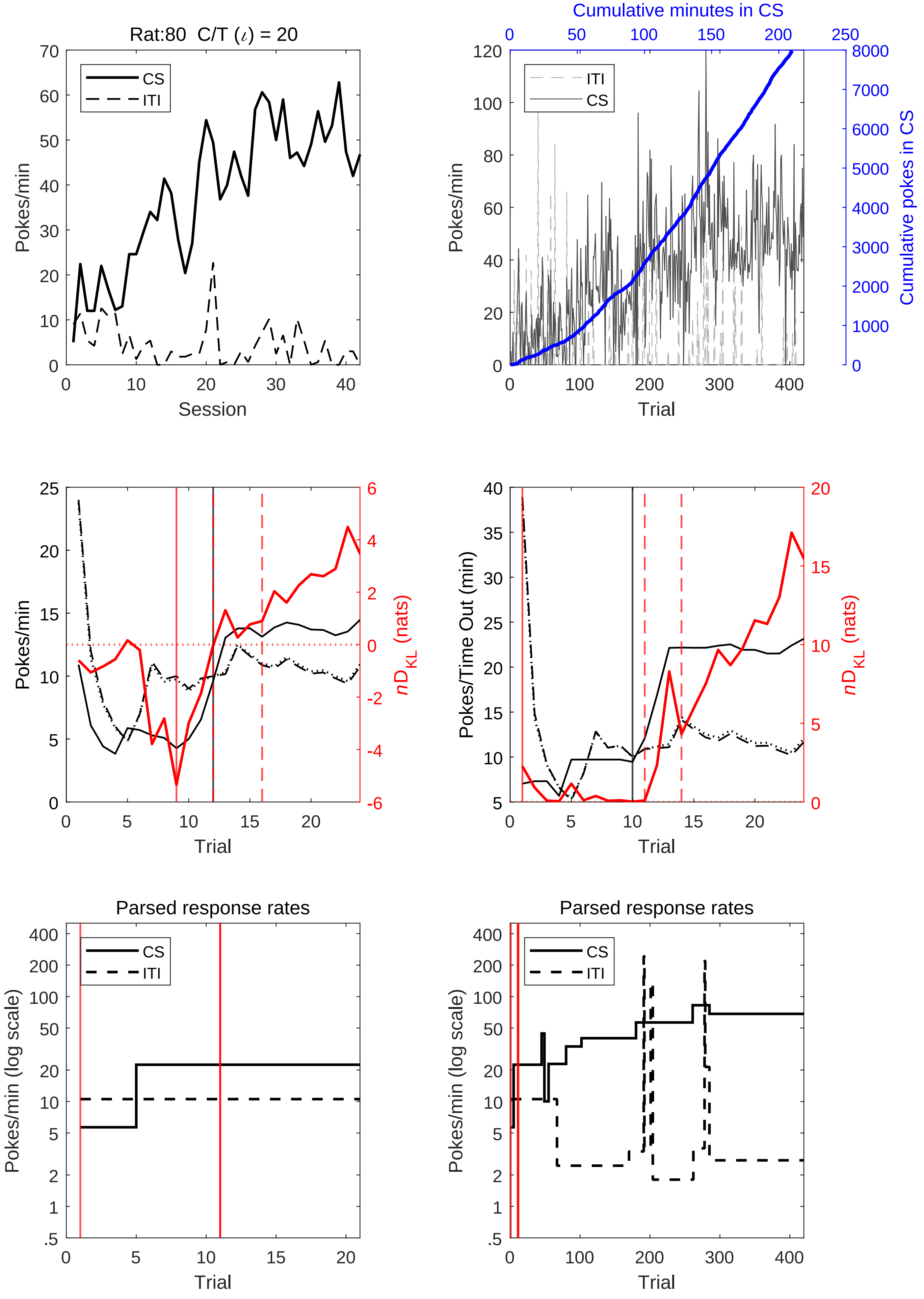
Rat:80 C/T (ι) = 20
CS
ITI
Pokes/min
Session
Cumulative minutes in CS
Cumulative pokes in CS
Trial
nD_KL (nats)
Pokes/Time Out (min)
Parsed response rates
Pokes/min (log scale)

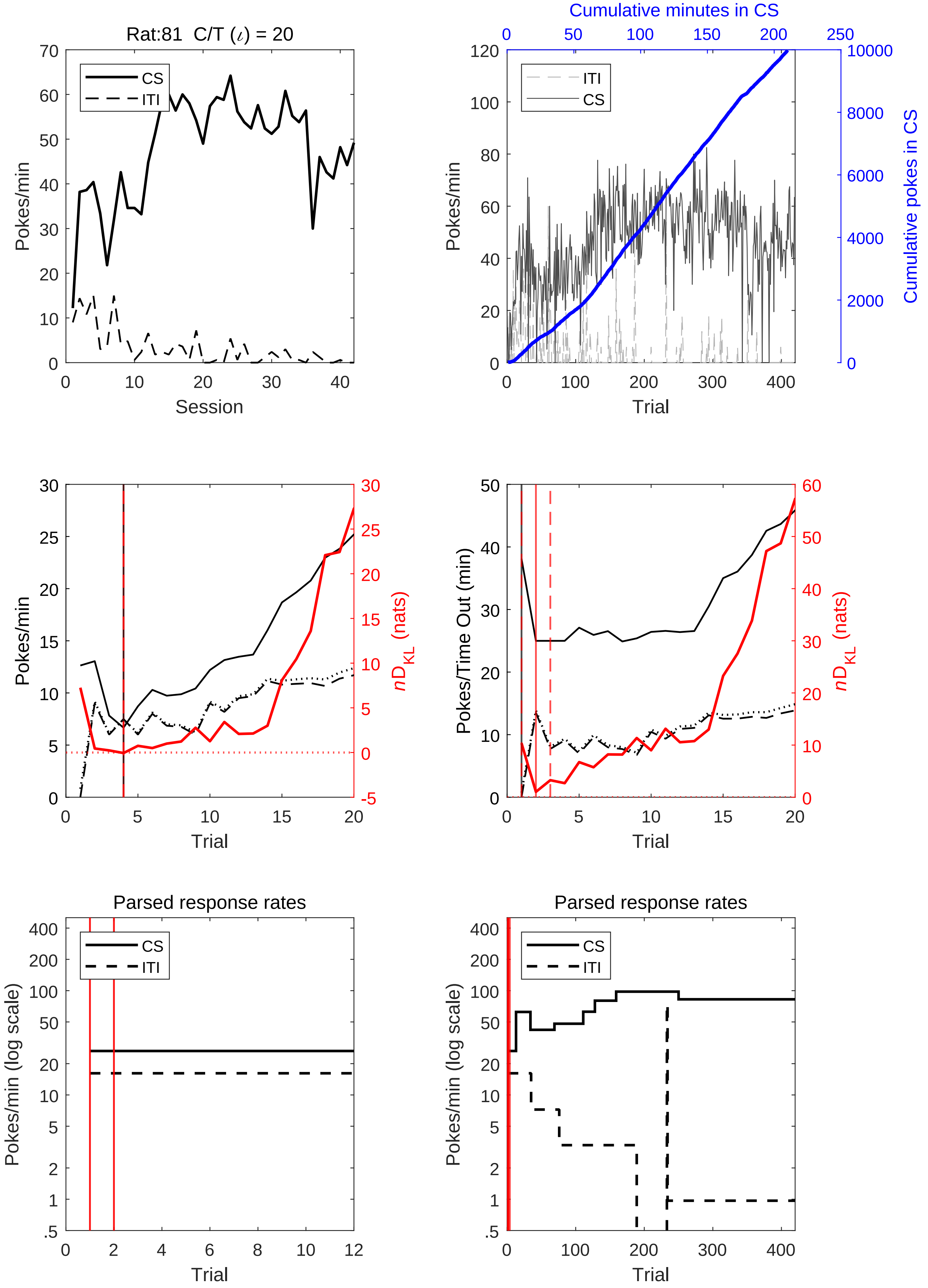

Rat:81 C/T (ι) = 20
CS
ITI
Pokes/min
Session
Cumulative minutes in CS
Cumulative pokes in CS
Trial
nD_KL (nats)
Pokes/Time Out (min)
Parsed response rates
Pokes/min (log scale)

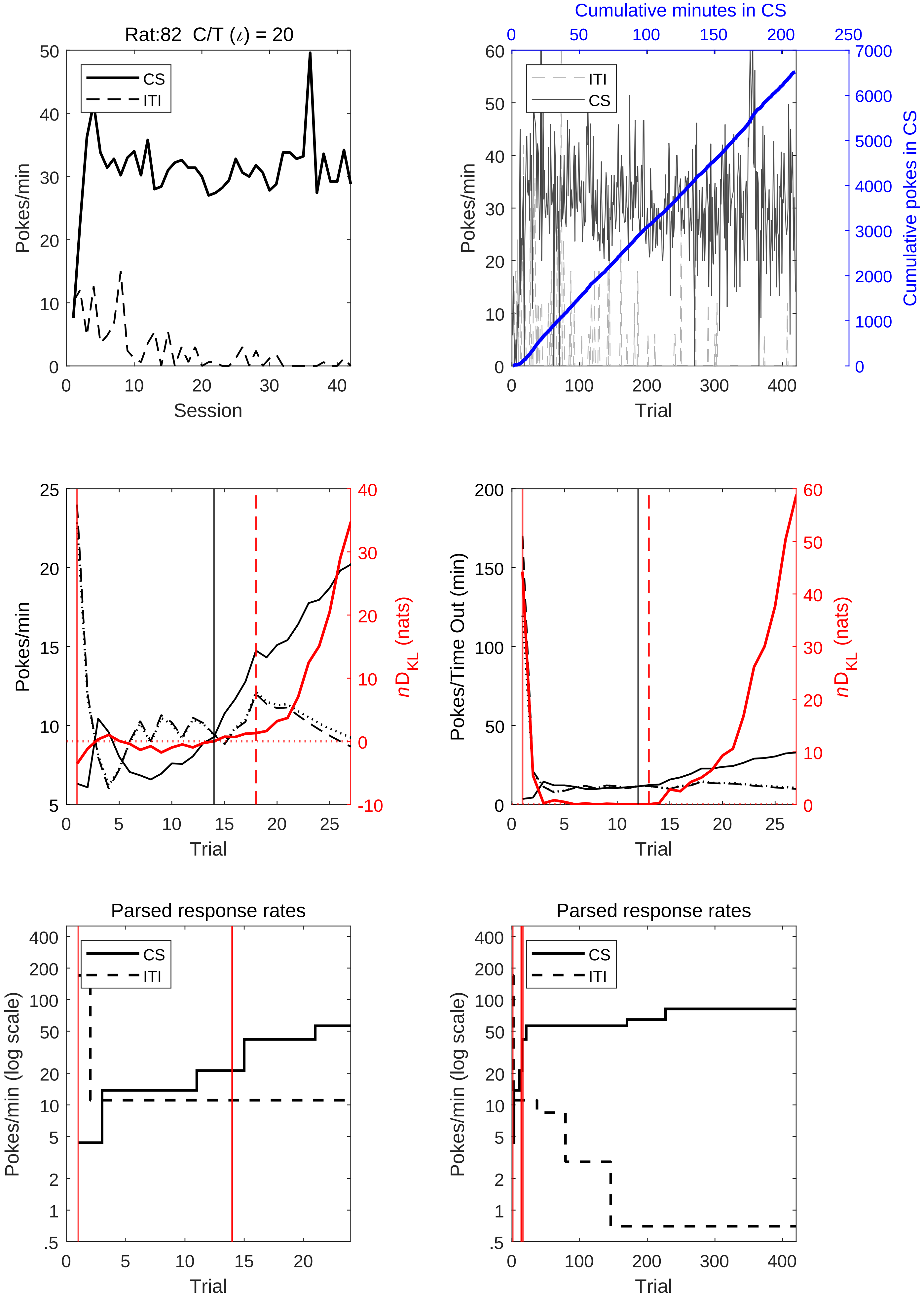

Rat:82 C/T (ι) = 20
CS
ITI
Pokes/min
Session
Cumulative minutes in CS
Cumulative pokes in CS
Trial
Pokes/Time Out (min)
$nD_{KL}$ (nats)
Parsed response rates
Pokes/min (log scale)

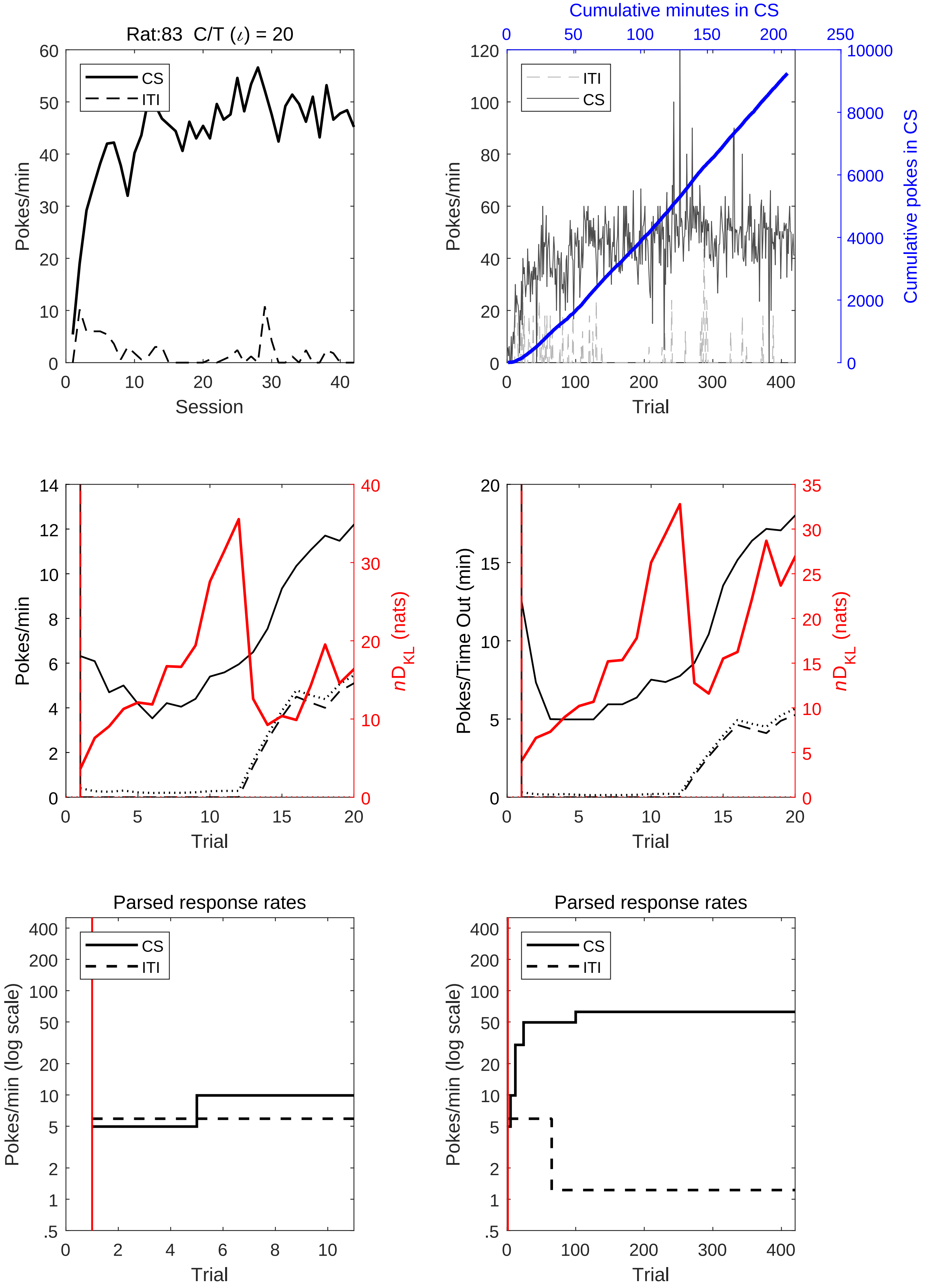

Rat:83 C/T (ι) = 20
CS
ITI
Pokes/min
Session
Cumulative minutes in CS
Cumulative pokes in CS
Trial
$nD_{KL}$ (nats)
Pokes/Time Out (min)
Parsed response rates
Pokes/min (log scale)

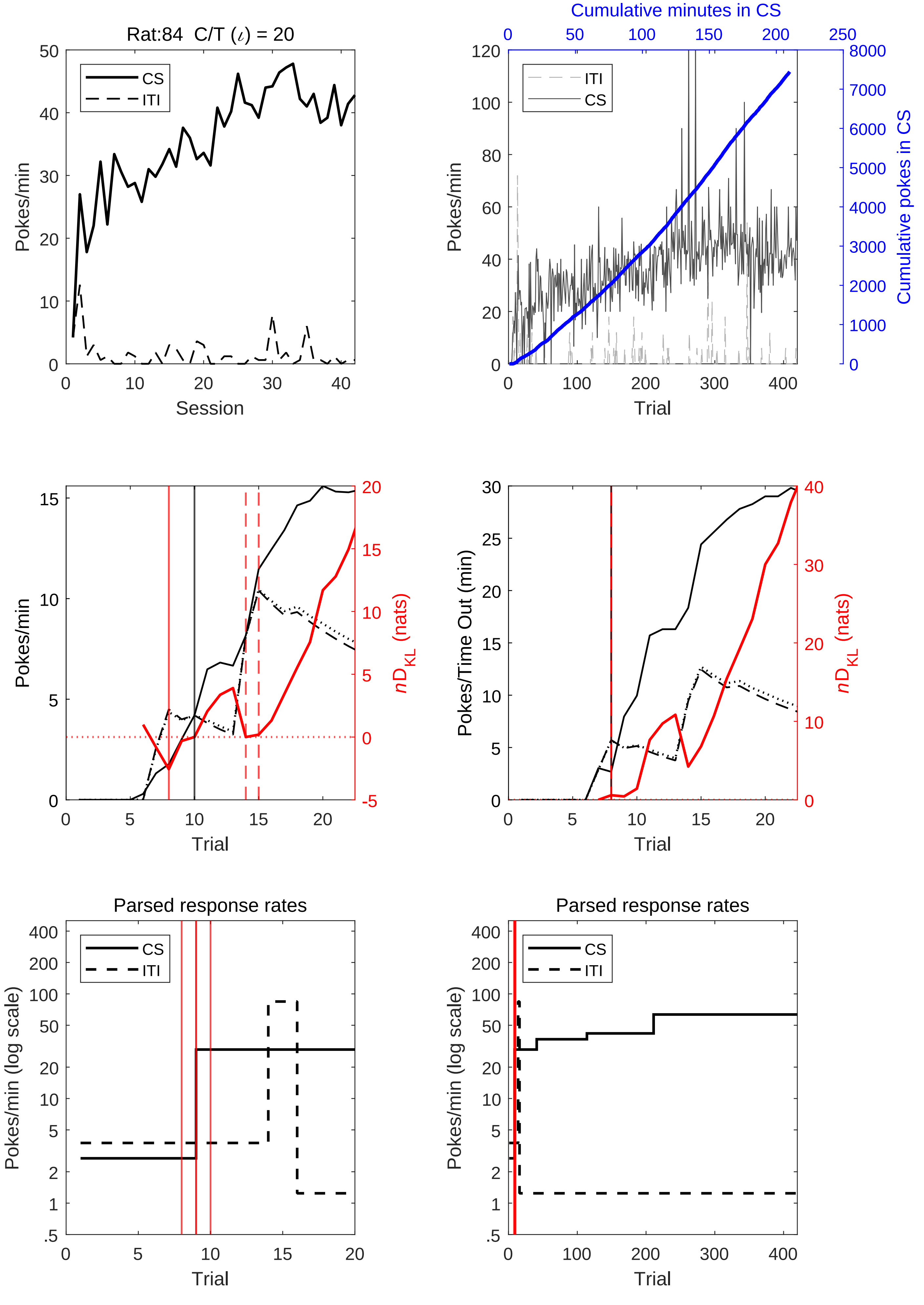
Rat:84 C/T (ι) = 20
CS
ITI
Pokes/min
Session
Cumulative minutes in CS
ITI
CS
Pokes/min
Cumulative pokes in CS
Trial
Pokes/min
nD_KL (nats)
Trial
Pokes/Time Out (min)
nD_KL (nats)
Trial
Parsed response rates
CS
ITI
Pokes/min (log scale)
Trial
Parsed response rates
CS
ITI
Pokes/min (log scale)
Trial

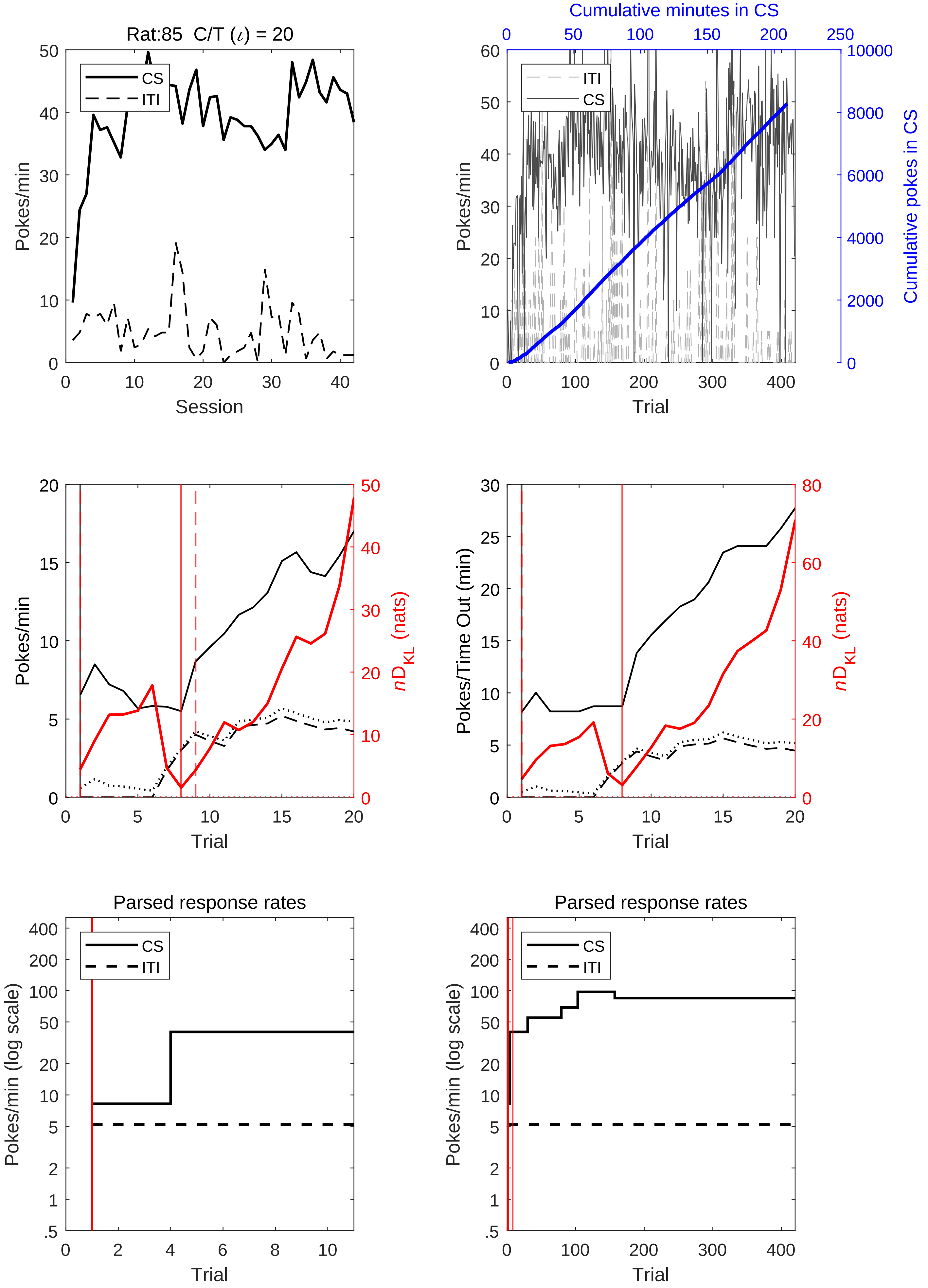
Rat:85 C/T (ι) = 20
CS
ITI
Pokes/min
Session
Cumulative minutes in CS
Cumulative pokes in CS
Trial
nD_KL (nats)
Pokes/Time Out (min)
Parsed response rates
Pokes/min (log scale)

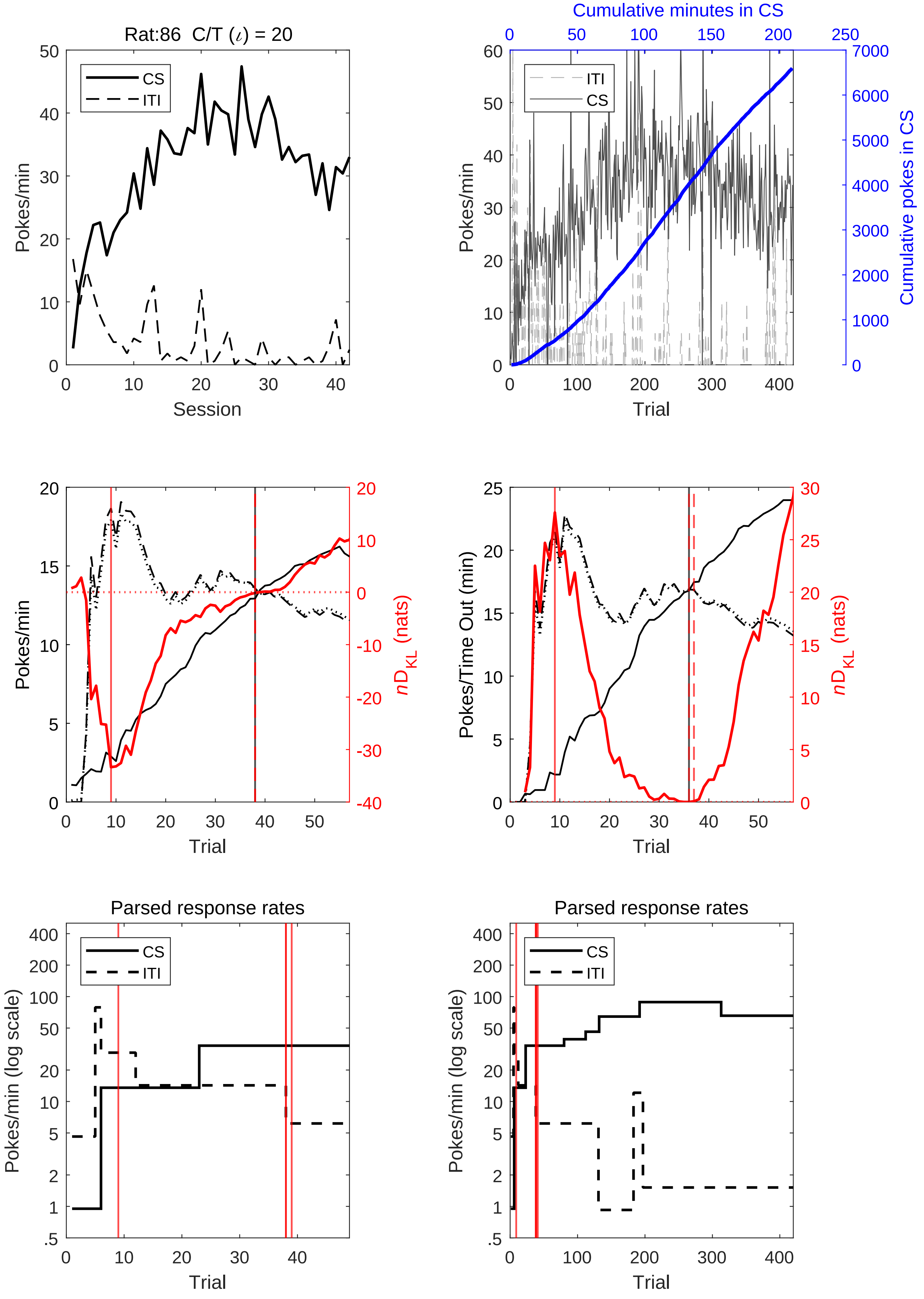
Rat:86  C/T (ι) = 20
CS
ITI
Pokes/min
Session
0
10
20
30
40
50
Cumulative minutes in CS
0
50
100
150
200
250
ITI
CS
Pokes/min
Cumulative pokes in CS
Trial
0
1000
2000
3000
4000
5000
6000
7000
60
100
200
300
400
Pokes/min
nD_KL (nats)
Trial
-40
-30
-20
-10
0
10
20
15
5
Pokes/Time Out (min)
nD_KL (nats)
Trial
25
Parsed response rates
CS
ITI
Pokes/min (log scale)
Trial
.5
1
2
5
10
20
50
100
200
400
Parsed response rates
CS
ITI
Pokes/min (log scale)
Trial

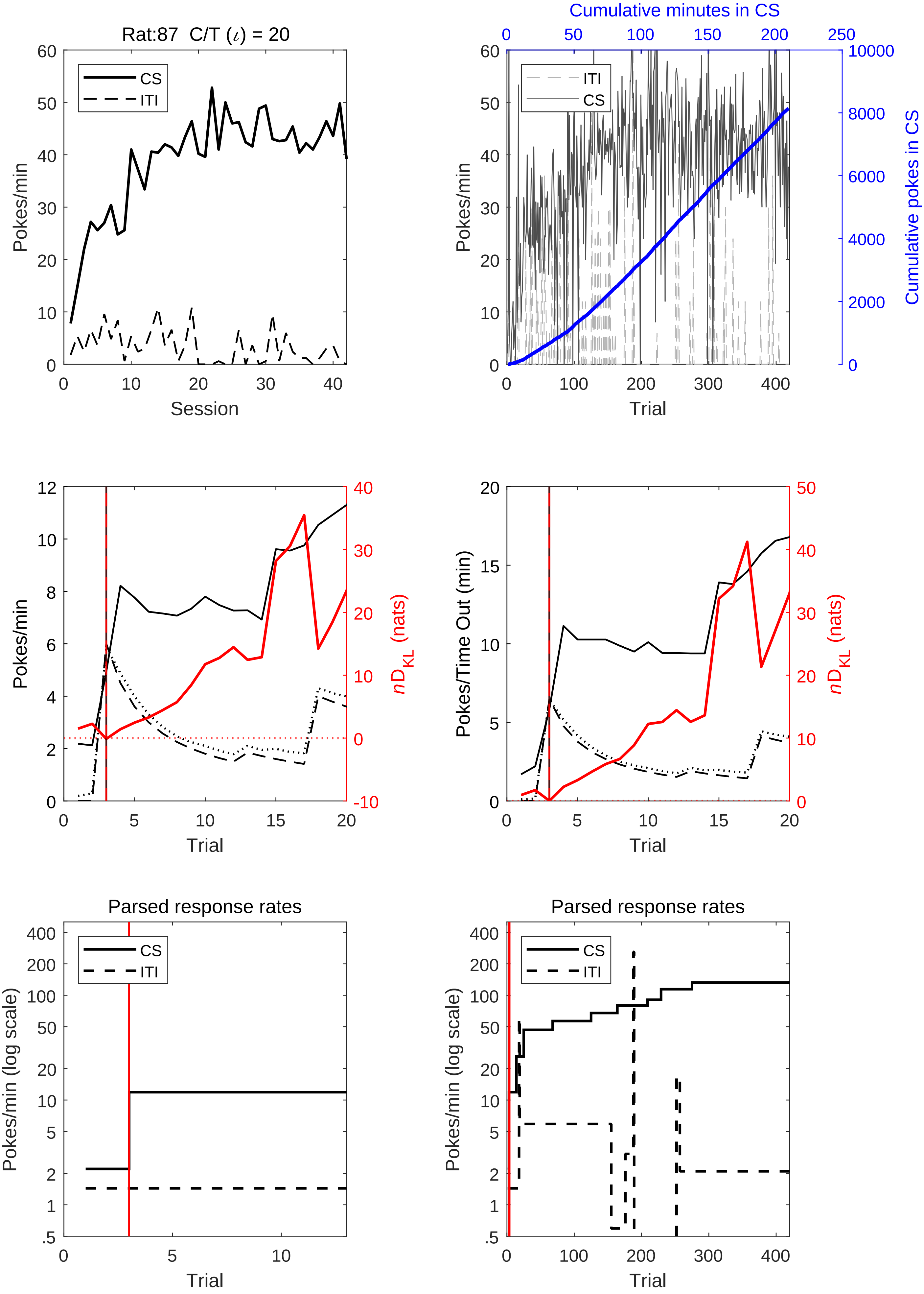
Rat:87 C/T (ι) = 20
Pokes/min
Session
CS
ITI
Cumulative minutes in CS
Cumulative pokes in CS
Trial
Pokes/min
nD_KL (nats)
Pokes/Time Out (min)
Parsed response rates
Pokes/min (log scale)

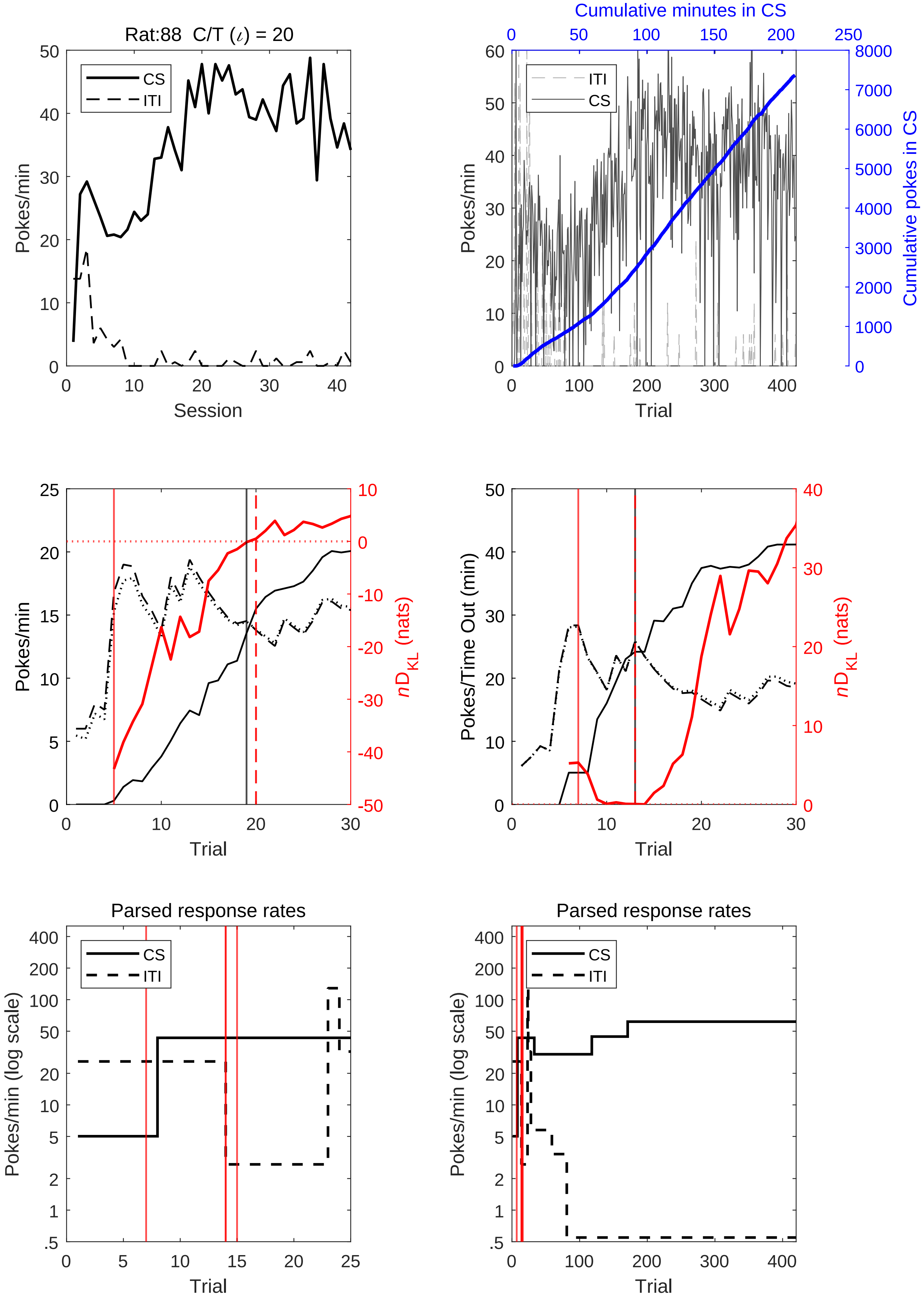
Rat:88 C/T (ι) = 20
CS
ITI
Pokes/min
Session
Cumulative minutes in CS
Cumulative pokes in CS
Trial
$nD_{KL}$ (nats)
Pokes/Time Out (min)
Parsed response rates
Pokes/min (log scale)

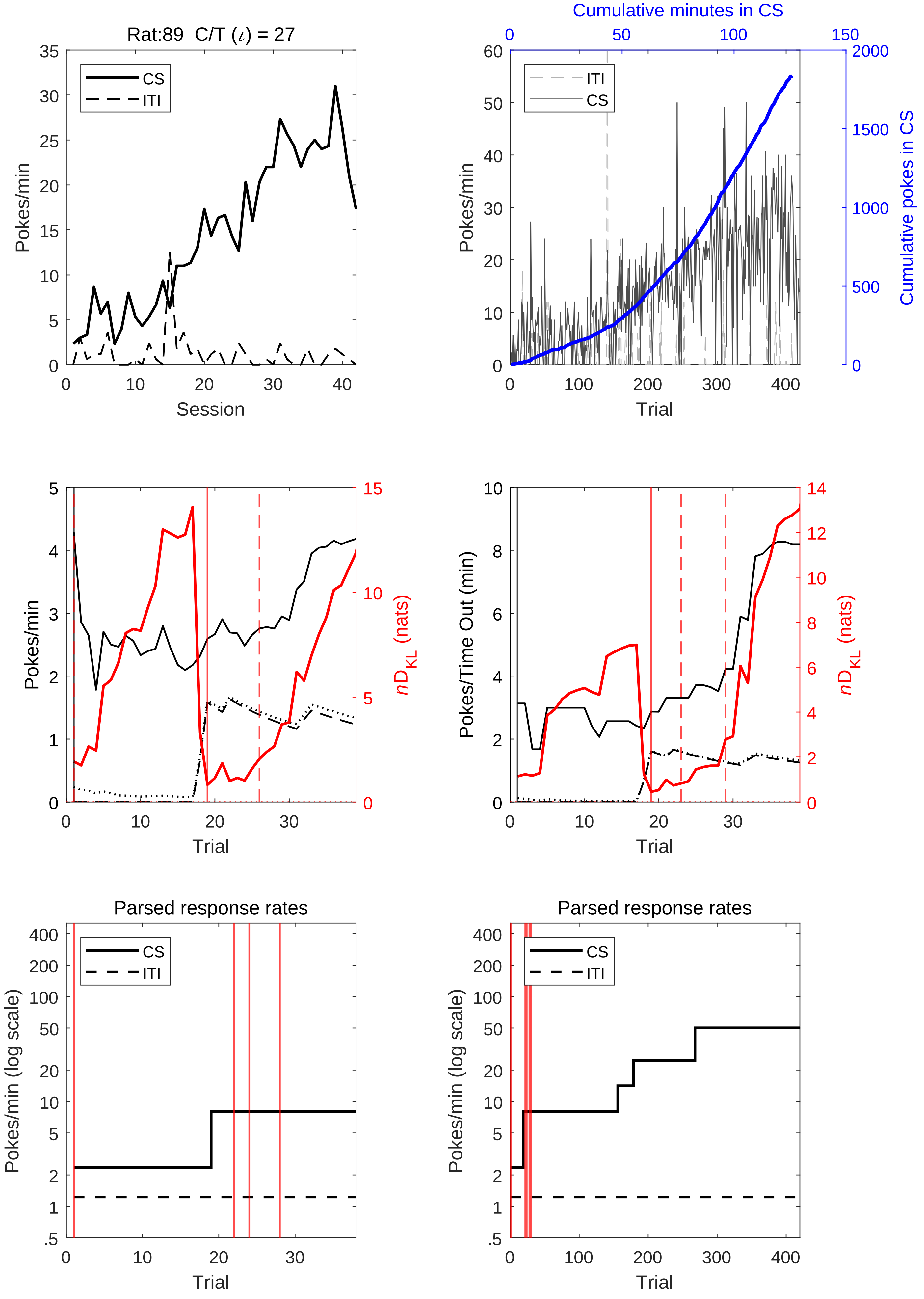
Rat:89 C/T ($\iota$) = 27
CS
ITI
Pokes/min
Session
Cumulative minutes in CS
ITI
CS
Pokes/min
Cumulative pokes in CS
Trial
Pokes/min
$nD_{KL}$ (nats)
Trial
Pokes/Time Out (min)
$nD_{KL}$ (nats)
Trial
Parsed response rates
CS
ITI
Pokes/min (log scale)
Trial
Parsed response rates
CS
ITI
Pokes/min (log scale)
Trial

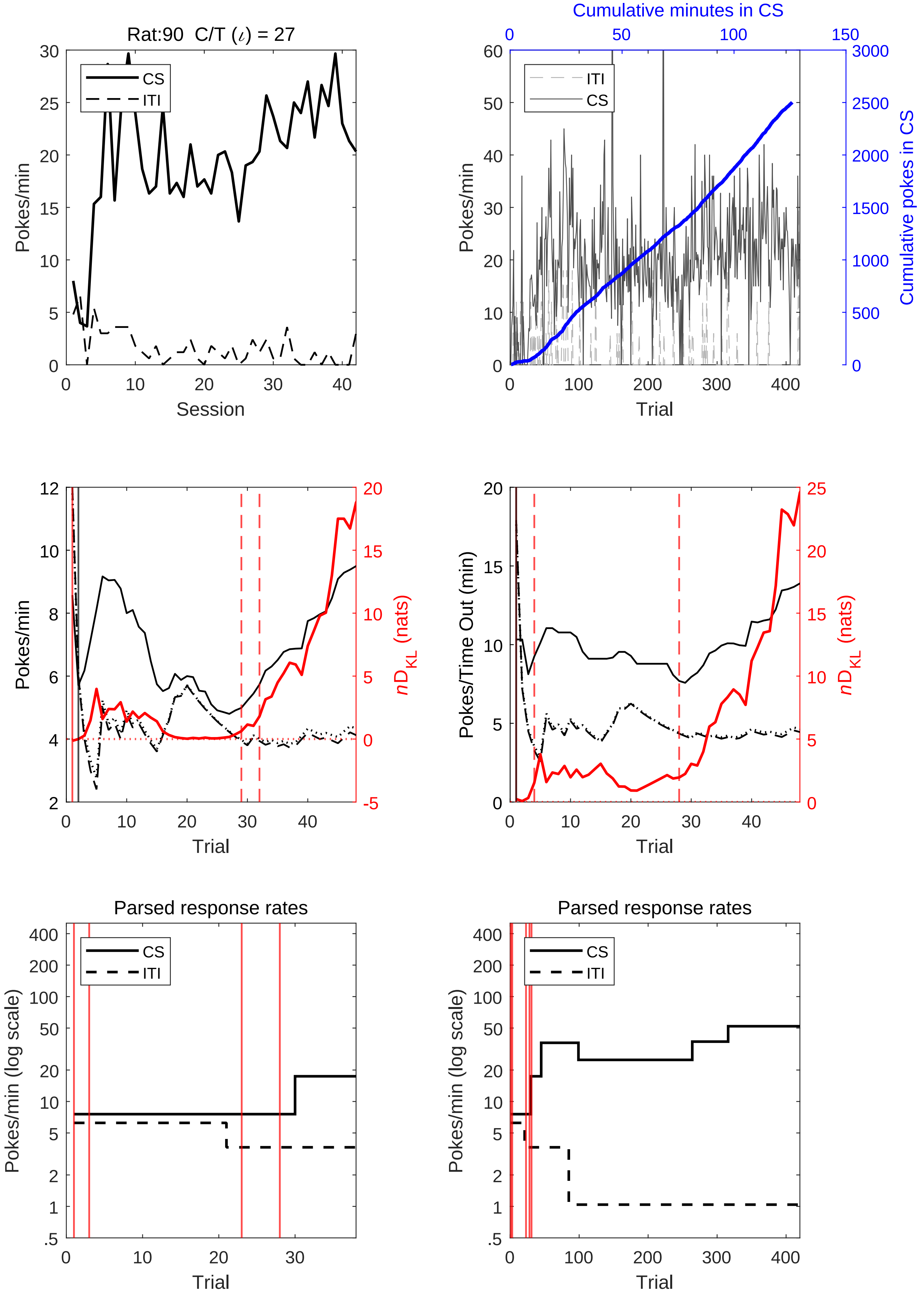
Rat:90 C/T ($\iota$) = 27
Pokes/min
Session
CS
ITI
Cumulative minutes in CS
Cumulative pokes in CS
Trial
Pokes/min
ITI
CS
Pokes/min
Trial
$nD_{KL}$ (nats)
Pokes/Time Out (min)
Trial
$nD_{KL}$ (nats)
Parsed response rates
Pokes/min (log scale)
Trial
CS
ITI
Parsed response rates
Pokes/min (log scale)
Trial
CS
ITI

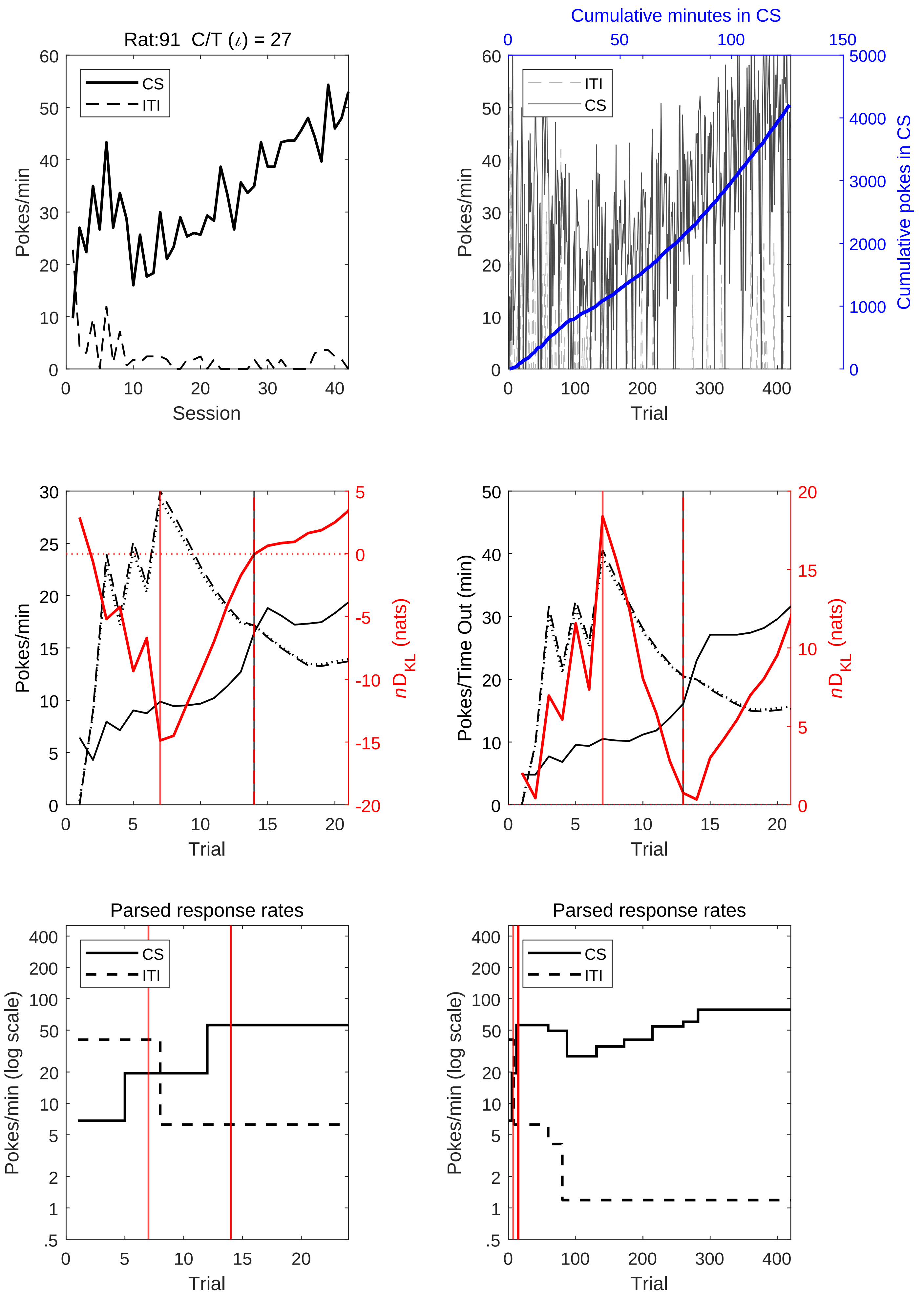

Rat:91 C/T (ι) = 27
CS
ITI
Pokes/min
Session
Cumulative minutes in CS
Cumulative pokes in CS
Trial
Pokes/Time Out (min)
nD_KL (nats)
Parsed response rates
Pokes/min (log scale)

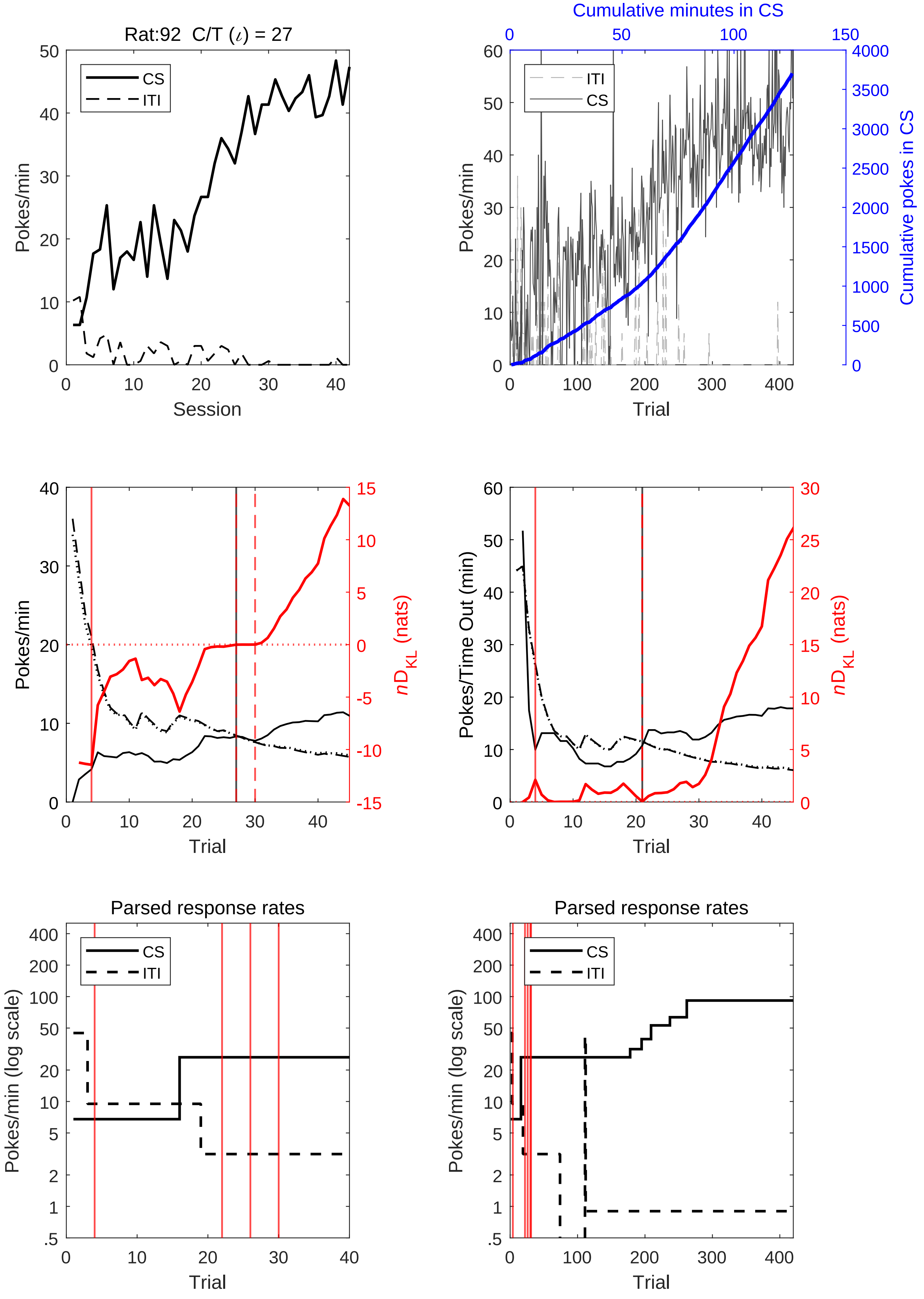

Rat:92 C/T (ι) = 27
CS
ITI
Pokes/min
Session
Cumulative minutes in CS
Cumulative pokes in CS
ITI
CS
Pokes/min
Trial
Pokes/min
$nD_{KL}$ (nats)
Trial
Pokes/Time Out (min)
$nD_{KL}$ (nats)
Trial
Parsed response rates
CS
ITI
Pokes/min (log scale)
Trial
Parsed response rates
CS
ITI
Pokes/min (log scale)
Trial

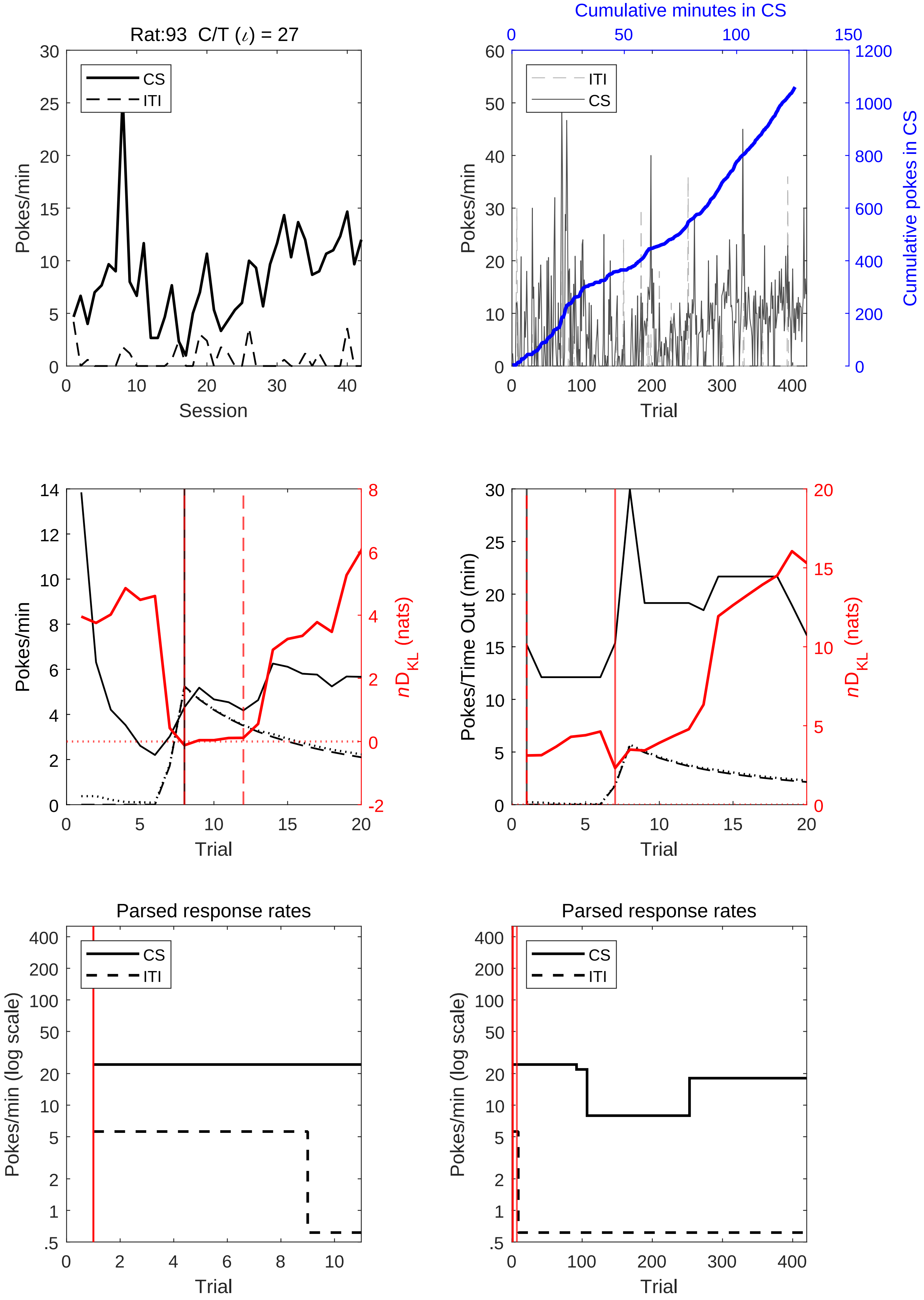

Rat:93 C/T (ι) = 27
CS
ITI
Pokes/min
Session
Cumulative minutes in CS
Cumulative pokes in CS
Pokes/min
Trial
Pokes/min
nD_KL (nats)
Trial
Pokes/Time Out (min)
nD_KL (nats)
Trial
Parsed response rates
Pokes/min (log scale)
Trial
Parsed response rates
Pokes/min (log scale)
Trial

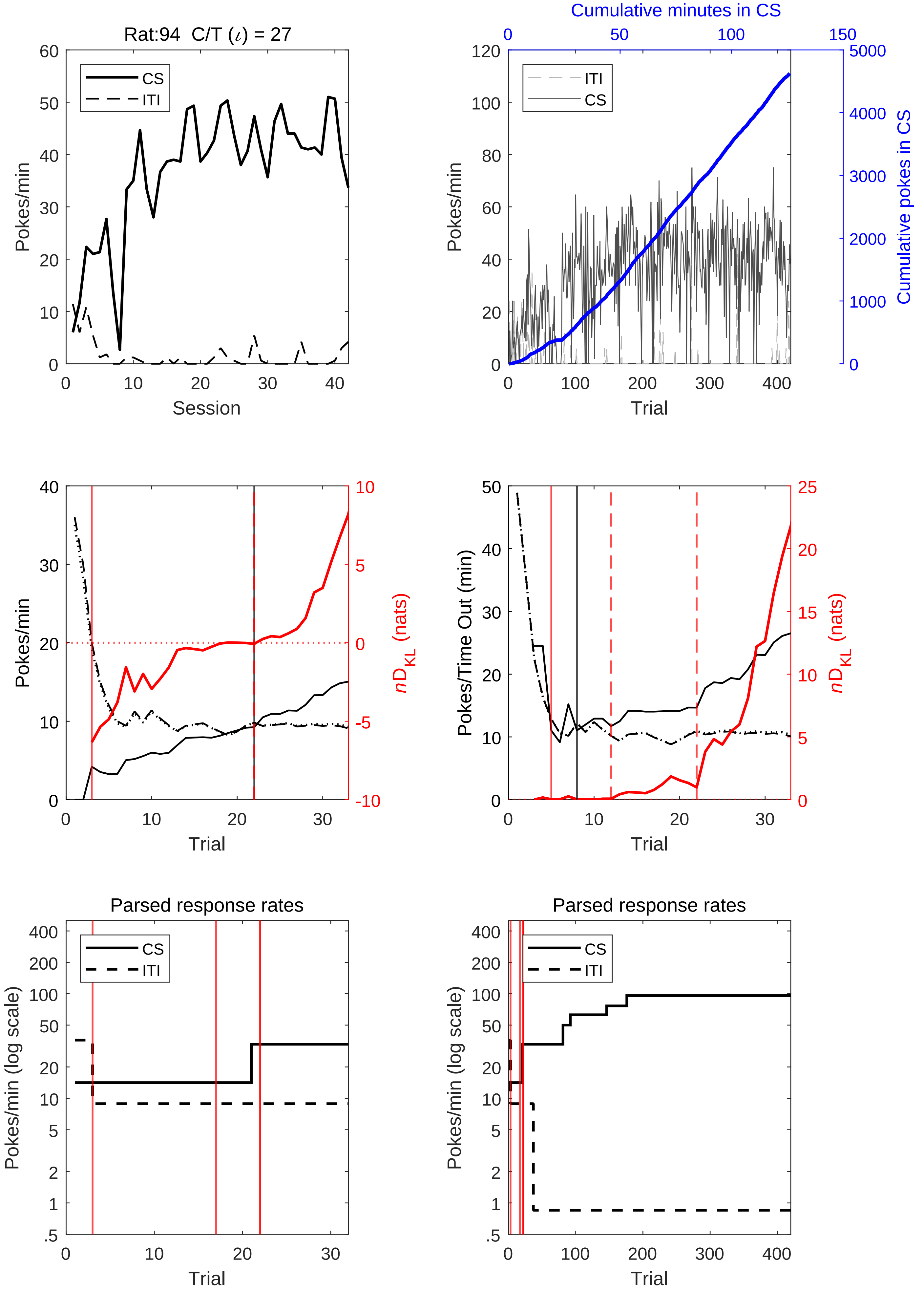

Rat:94 C/T ($\iota$) = 27
CS
ITI
Pokes/min
Session
Cumulative minutes in CS
Cumulative pokes in CS
Trial
nD_KL (nats)
Pokes/Time Out (min)
Parsed response rates
Pokes/min (log scale)

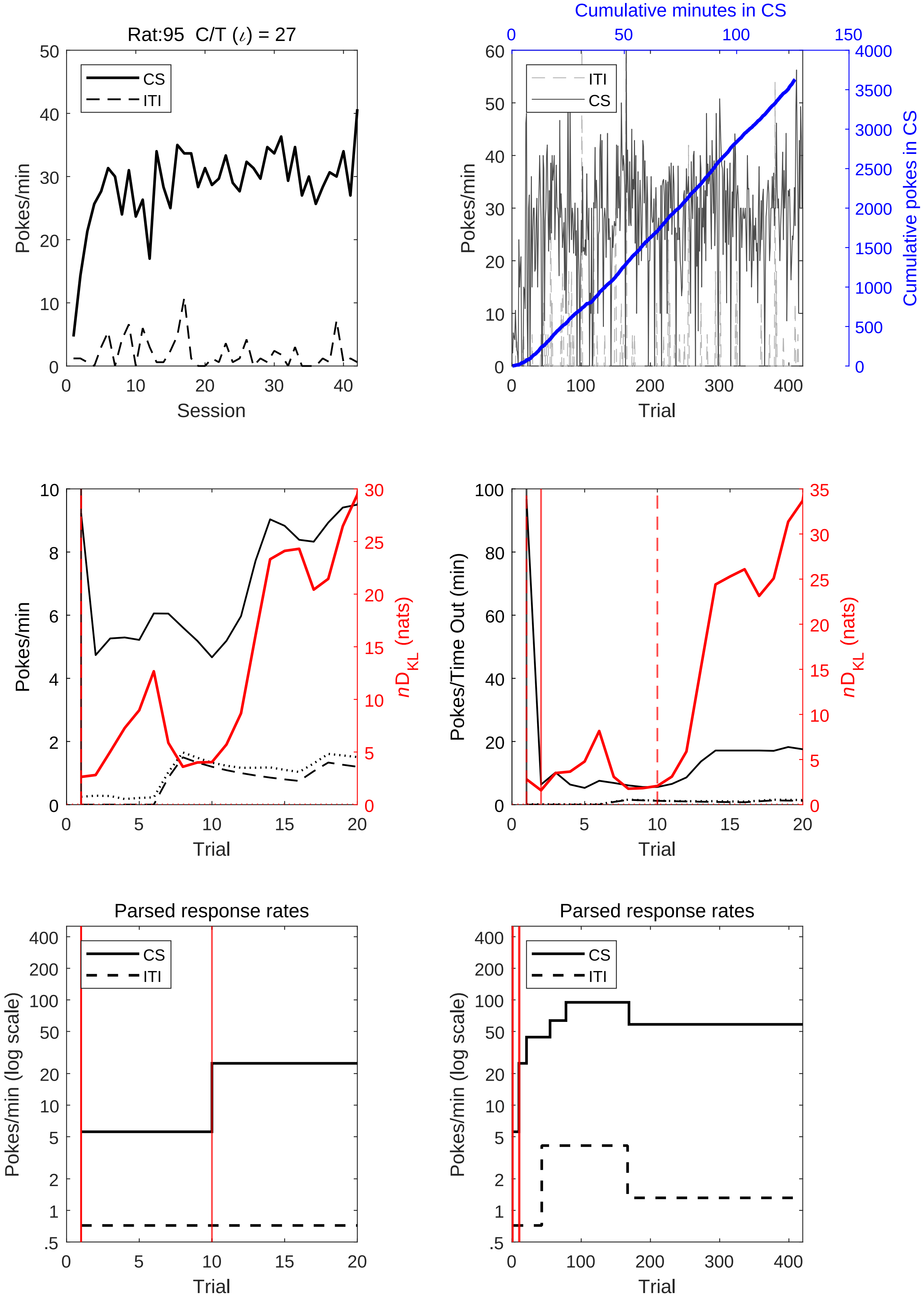
Rat:95 C/T ($\iota$) = 27
CS
ITI
Pokes/min
Session
Cumulative minutes in CS
ITI
CS
Pokes/min
Cumulative pokes in CS
Trial
Pokes/min
$n\mathrm{D}_{\mathrm{KL}}$ (nats)
Trial
Pokes/Time Out (min)
$n\mathrm{D}_{\mathrm{KL}}$ (nats)
Trial
Parsed response rates
CS
ITI
Pokes/min (log scale)
Trial
Parsed response rates
CS
ITI
Pokes/min (log scale)
Trial

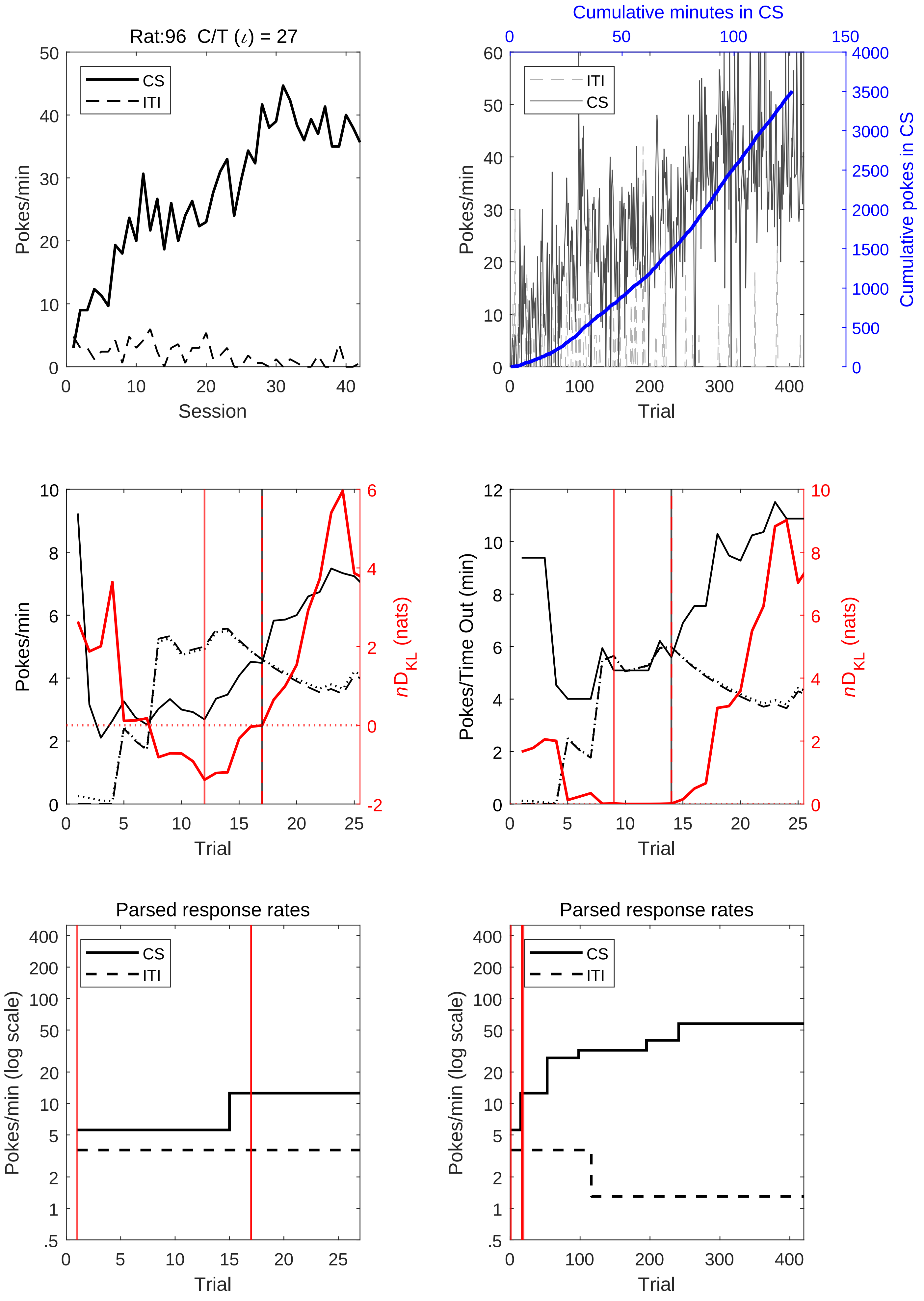

Rat:96 C/T (ι) = 27
CS
ITI
Pokes/min
Session
Cumulative minutes in CS
Cumulative pokes in CS
Trial
nD_KL (nats)
Pokes/Time Out (min)
Parsed response rates
Pokes/min (log scale)

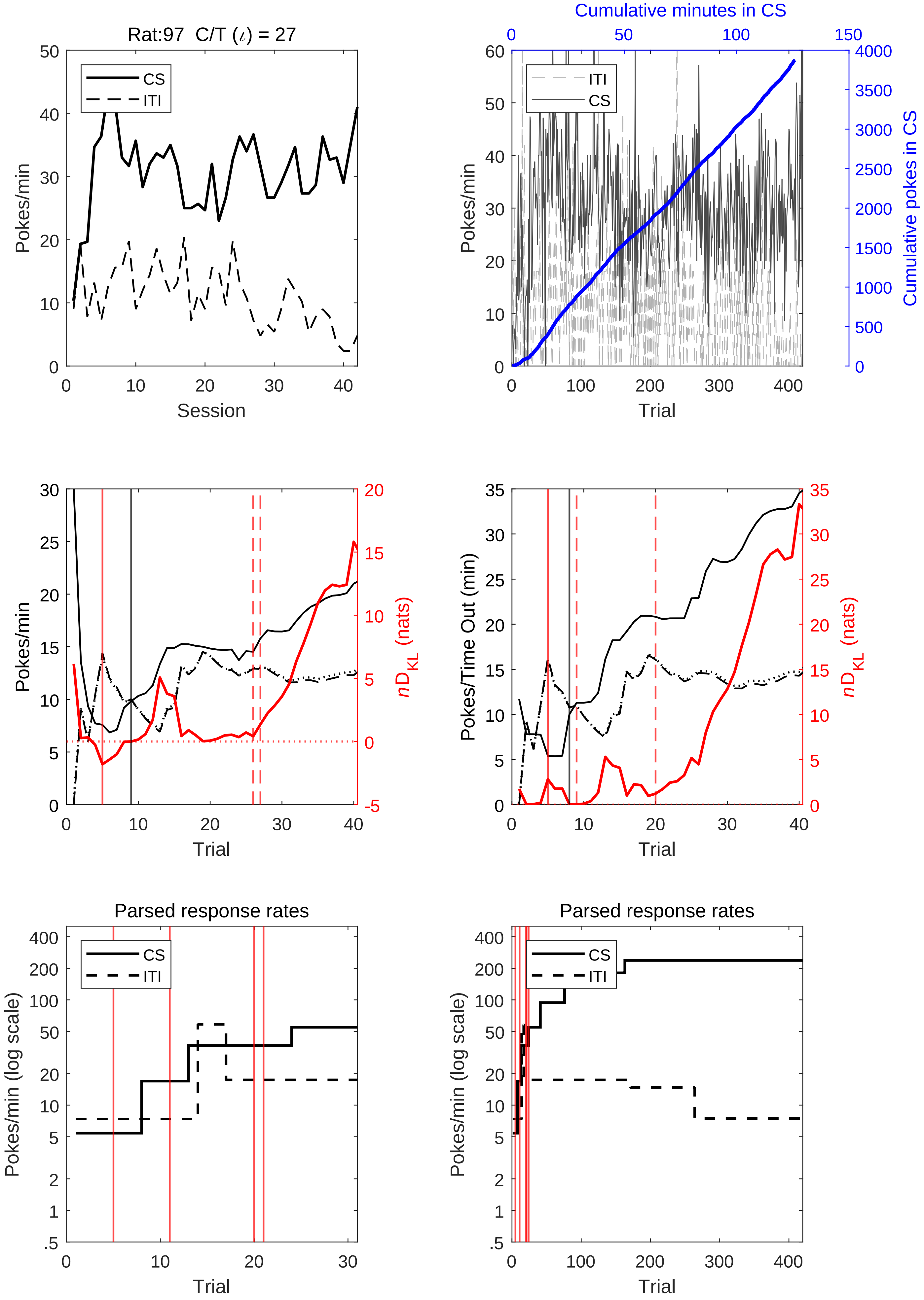
Rat:97 C/T (ι) = 27
CS
ITI
Pokes/min
Session
Cumulative minutes in CS
Cumulative pokes in CS
Trial
Pokes/Time Out (min)
nD_KL (nats)
Parsed response rates
Pokes/min (log scale)

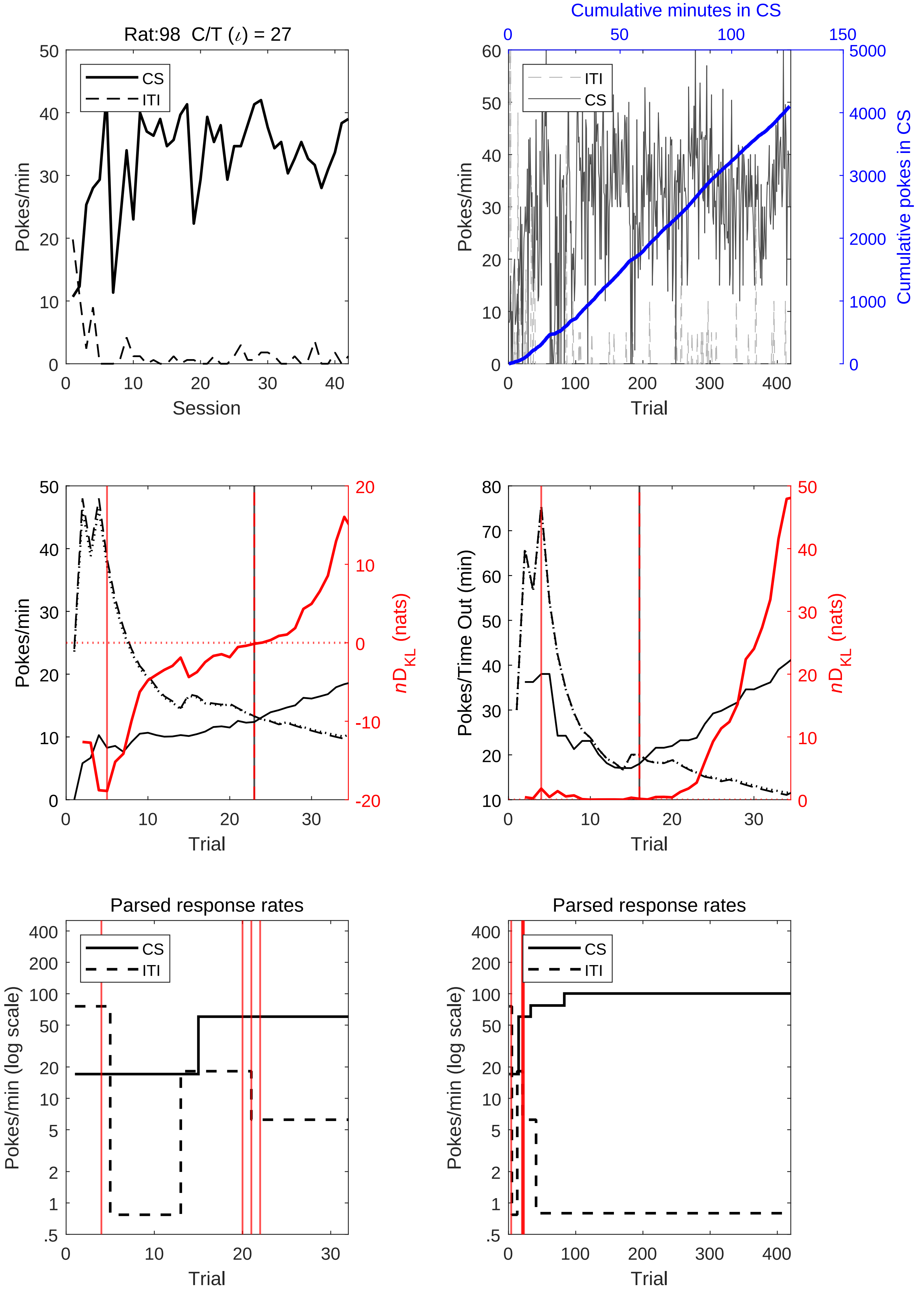
Rat:98 C/T (ι) = 27
CS
ITI
Pokes/min
Session
Cumulative minutes in CS
Cumulative pokes in CS
Trial
nD_KL (nats)
Pokes/Time Out (min)
Parsed response rates
Pokes/min (log scale)

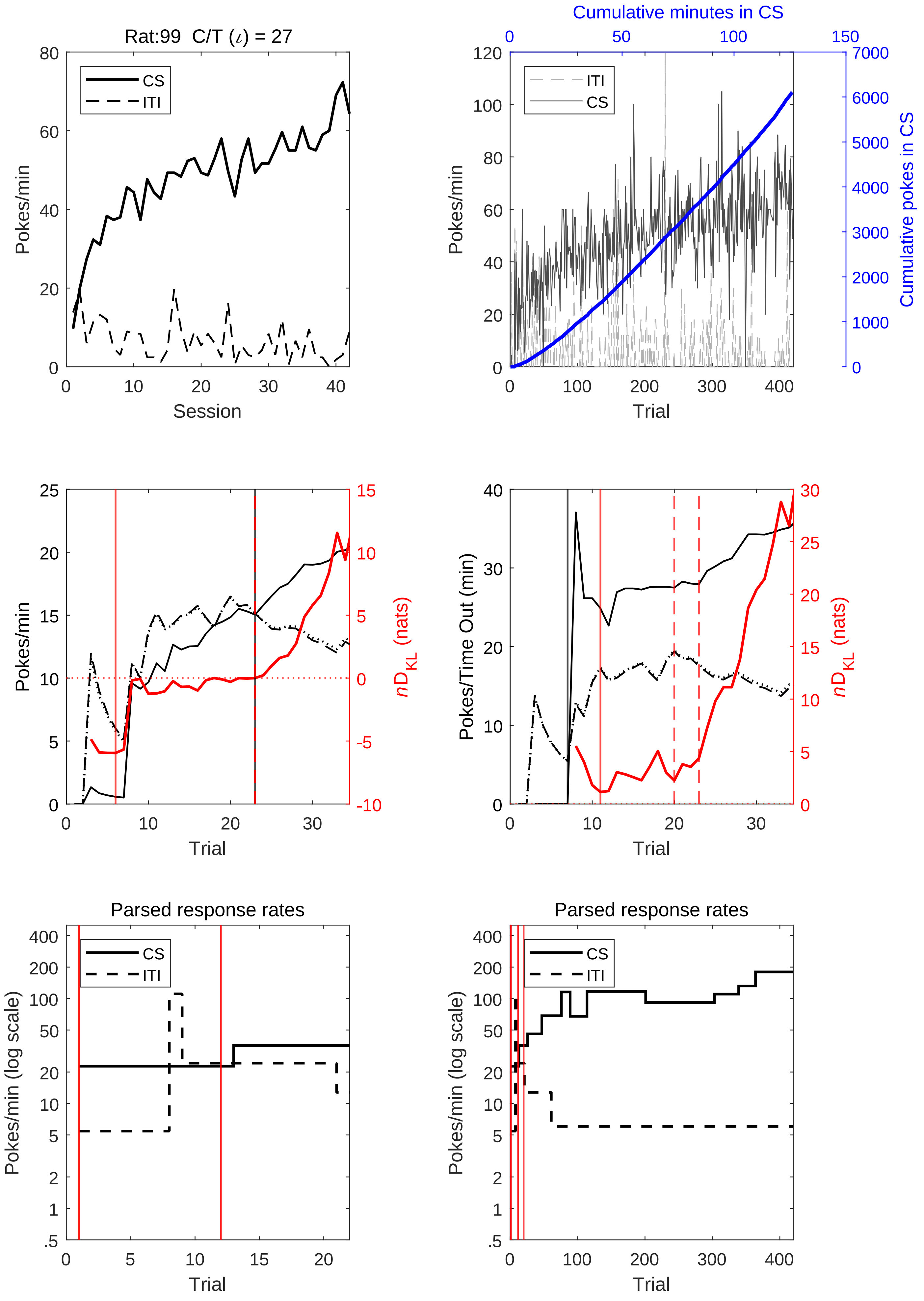
Rat:99 C/T (ι) = 27
CS
ITI
Pokes/min
Session
Cumulative minutes in CS
ITI
CS
Pokes/min
Cumulative pokes in CS
Trial
Pokes/min
nD_KL (nats)
Trial
Pokes/Time Out (min)
nD_KL (nats)
Trial
Parsed response rates
CS
ITI
Pokes/min (log scale)
Trial
Parsed response rates
CS
ITI
Pokes/min (log scale)
Trial

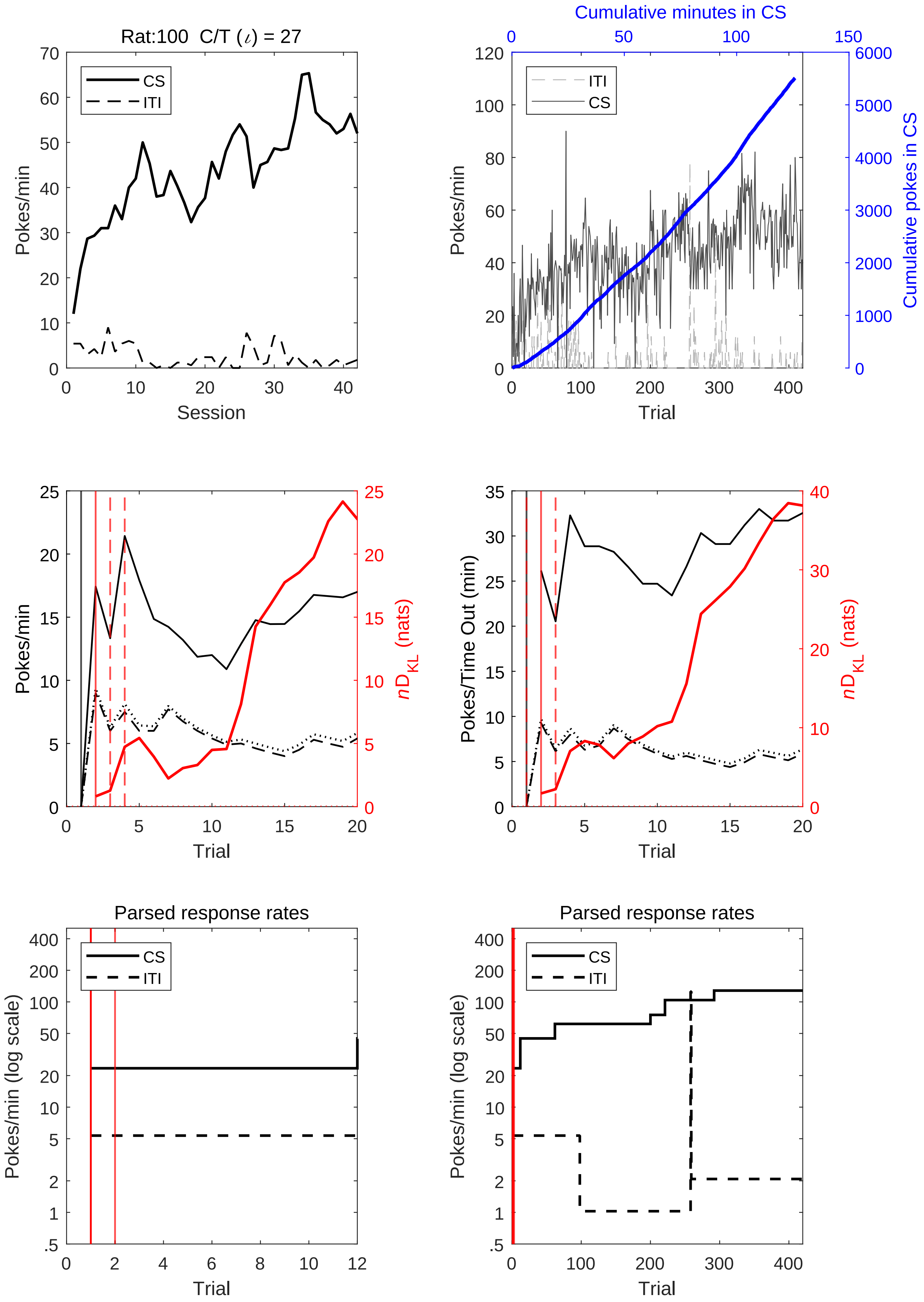
Rat:100 C/T (ι) = 27
CS
ITI
Pokes/min
Session
Cumulative minutes in CS
Cumulative pokes in CS
Trial
Pokes/Time Out (min)
nD_KL (nats)
Parsed response rates
Pokes/min (log scale)

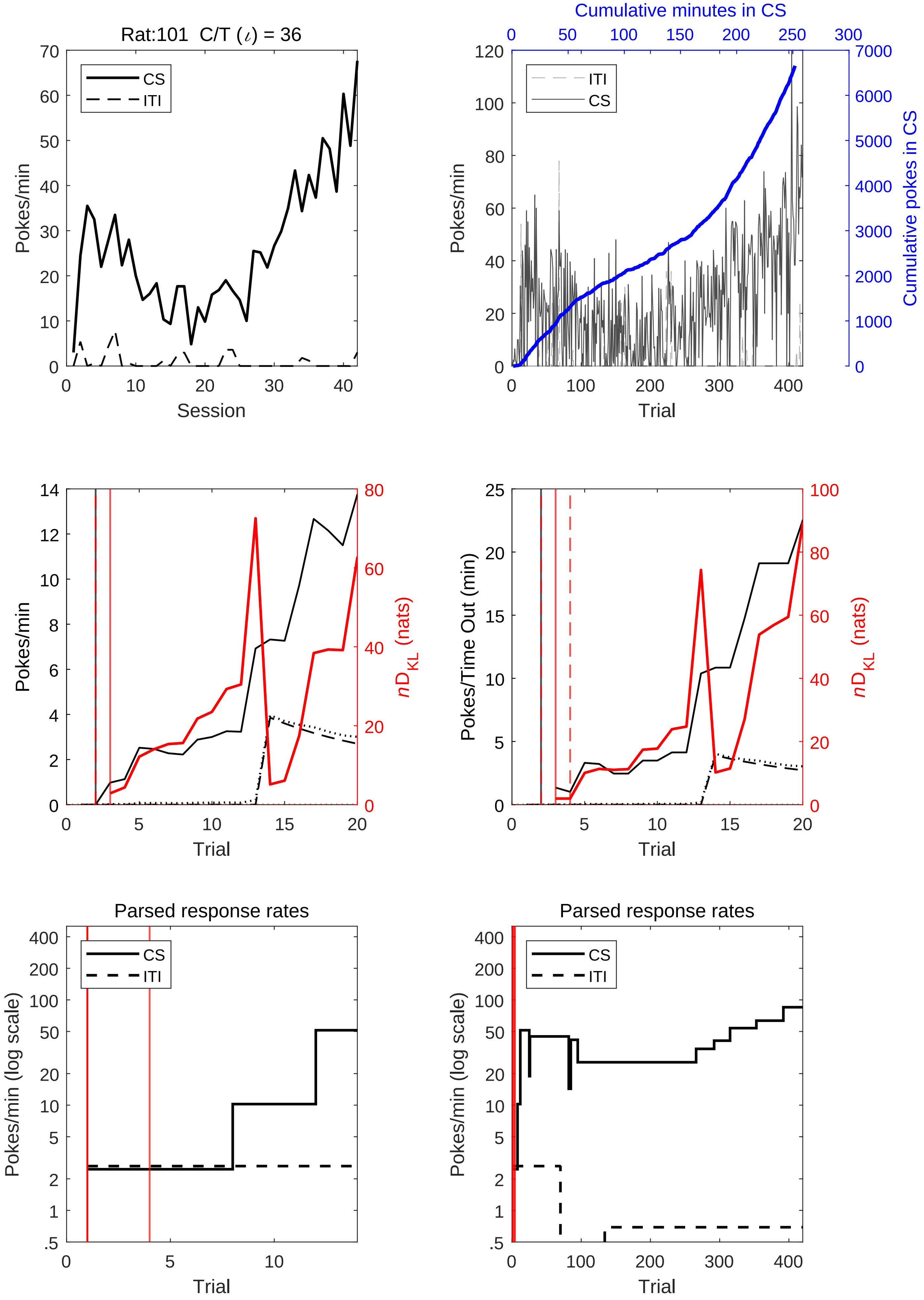
Rat:101 C/T (ι) = 36
CS
ITI
Pokes/min
Session
Cumulative minutes in CS
Cumulative pokes in CS
Trial
$nD_{KL}$ (nats)
Pokes/Time Out (min)
Parsed response rates
Pokes/min (log scale)

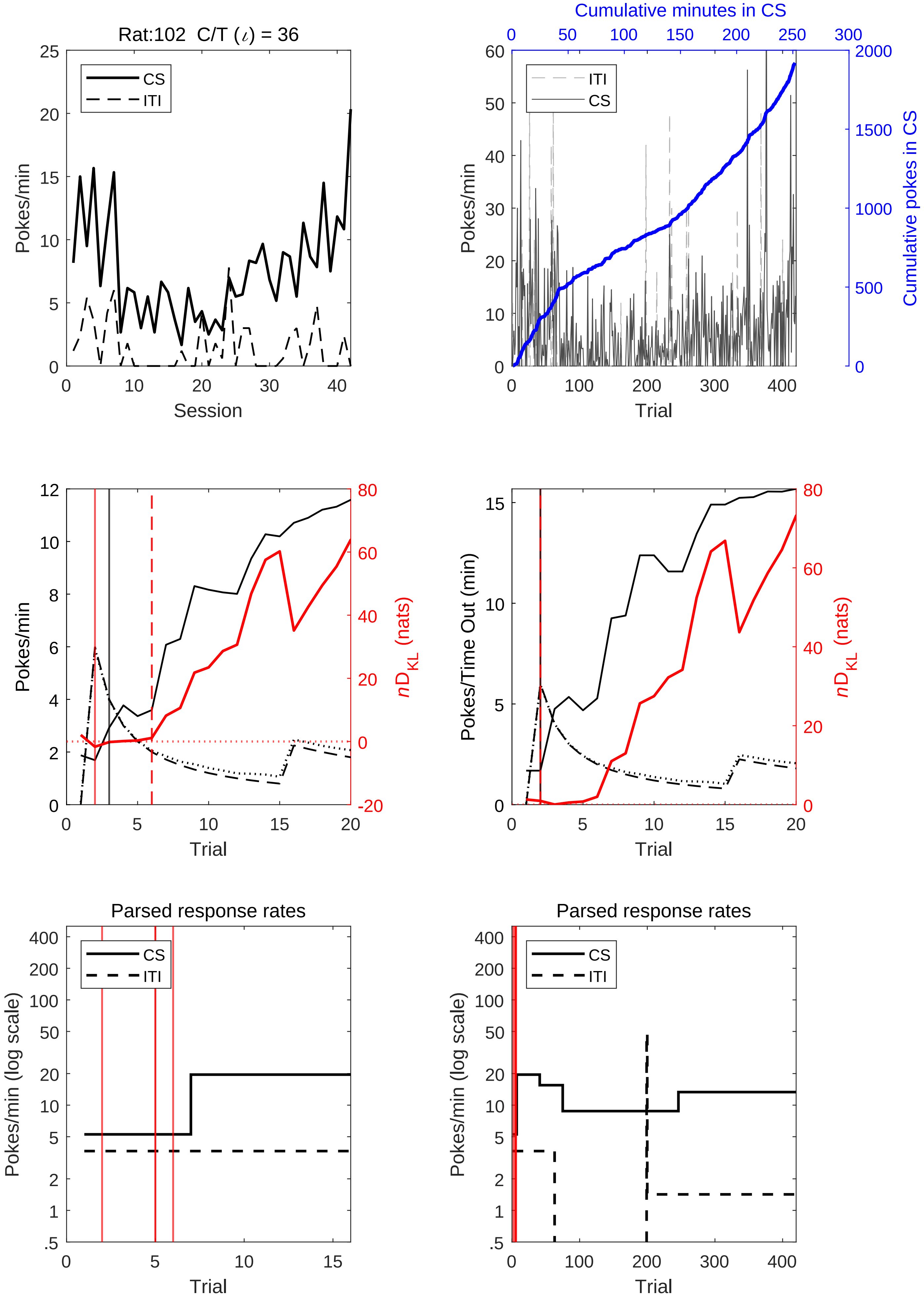
Rat:102 C/T (ι) = 36
CS
ITI
Pokes/min
Session
Cumulative minutes in CS
ITI
CS
Pokes/min
Cumulative pokes in CS
Trial
Pokes/min
nD_KL (nats)
Trial
Pokes/Time Out (min)
nD_KL (nats)
Trial
Parsed response rates
CS
ITI
Pokes/min (log scale)
Trial
Parsed response rates
CS
ITI
Pokes/min (log scale)
Trial

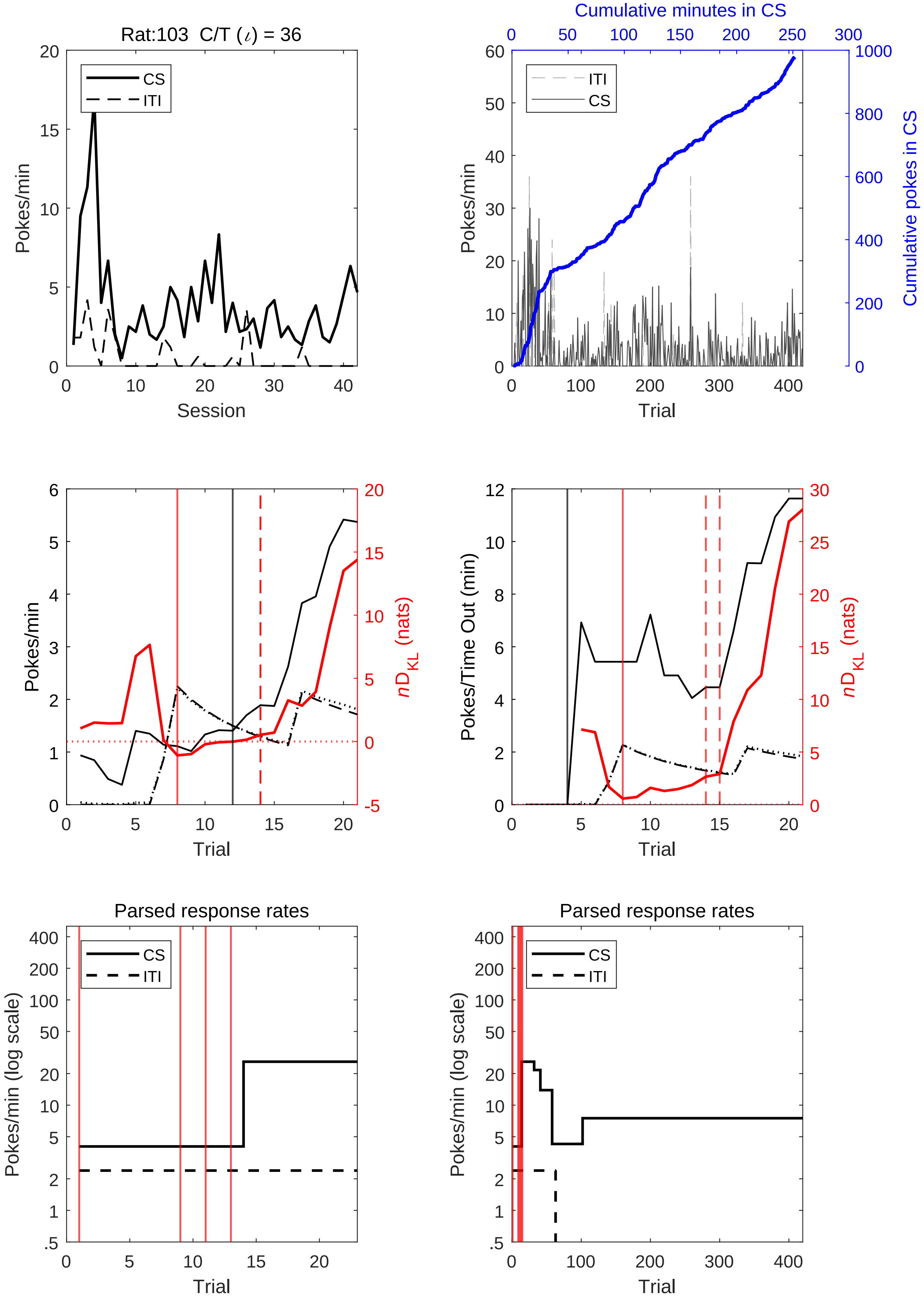
Rat:103 C/T (ι) = 36
CS
ITI
Pokes/min
Session
Cumulative minutes in CS
Cumulative pokes in CS
Trial
nD_KL (nats)
Pokes/Time Out (min)
Parsed response rates
Pokes/min (log scale)

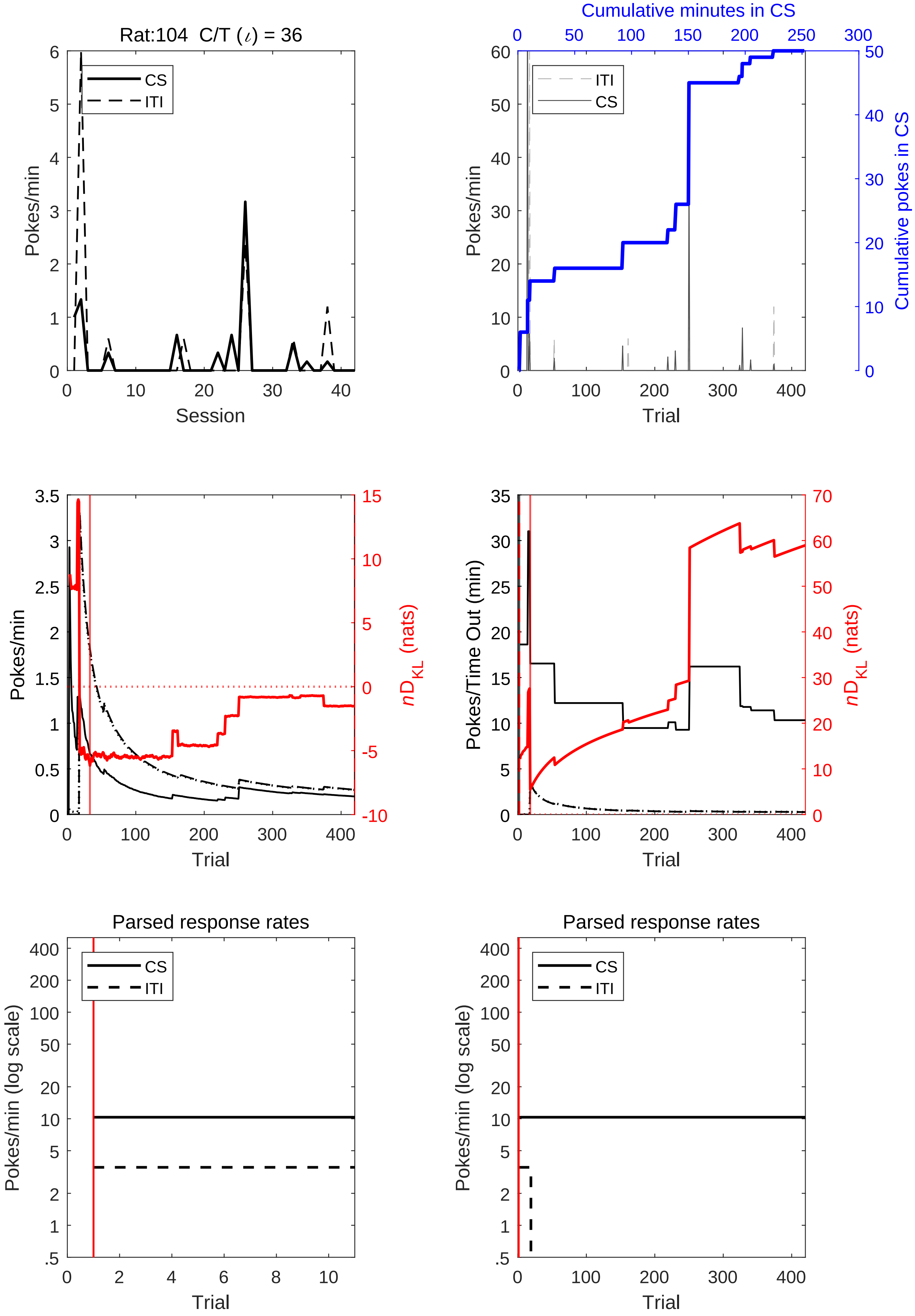

Rat:104 C/T (ι) = 36
CS
ITI
Pokes/min
Session
Cumulative minutes in CS
Cumulative pokes in CS
Trial
nD_KL (nats)
Pokes/Time Out (min)
Parsed response rates
Pokes/min (log scale)

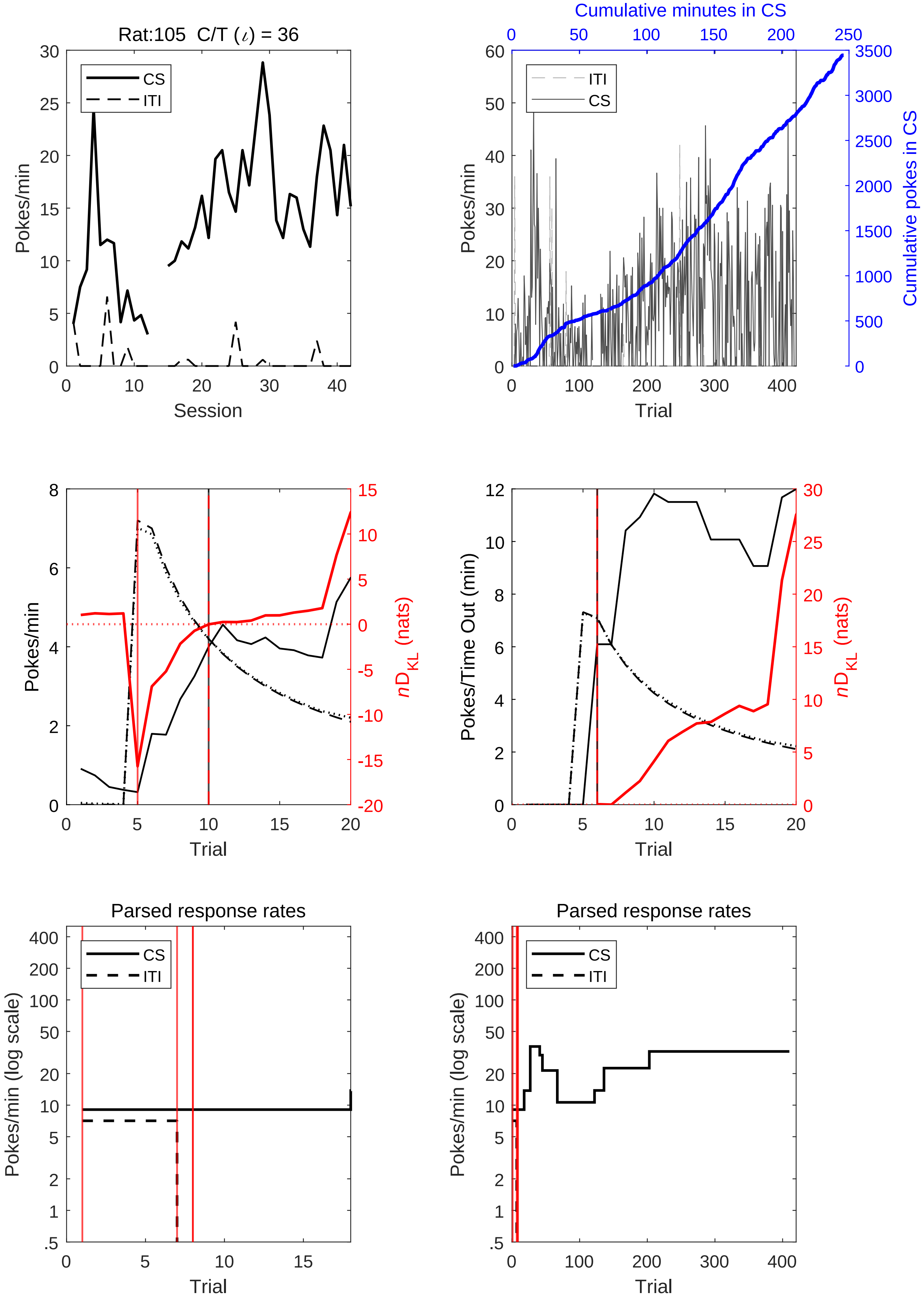
Rat:105 C/T (ι) = 36
CS
ITI
Pokes/min
Session
Cumulative minutes in CS
Cumulative pokes in CS
Trial
$nD_{KL}$ (nats)
Pokes/Time Out (min)
Parsed response rates
Pokes/min (log scale)

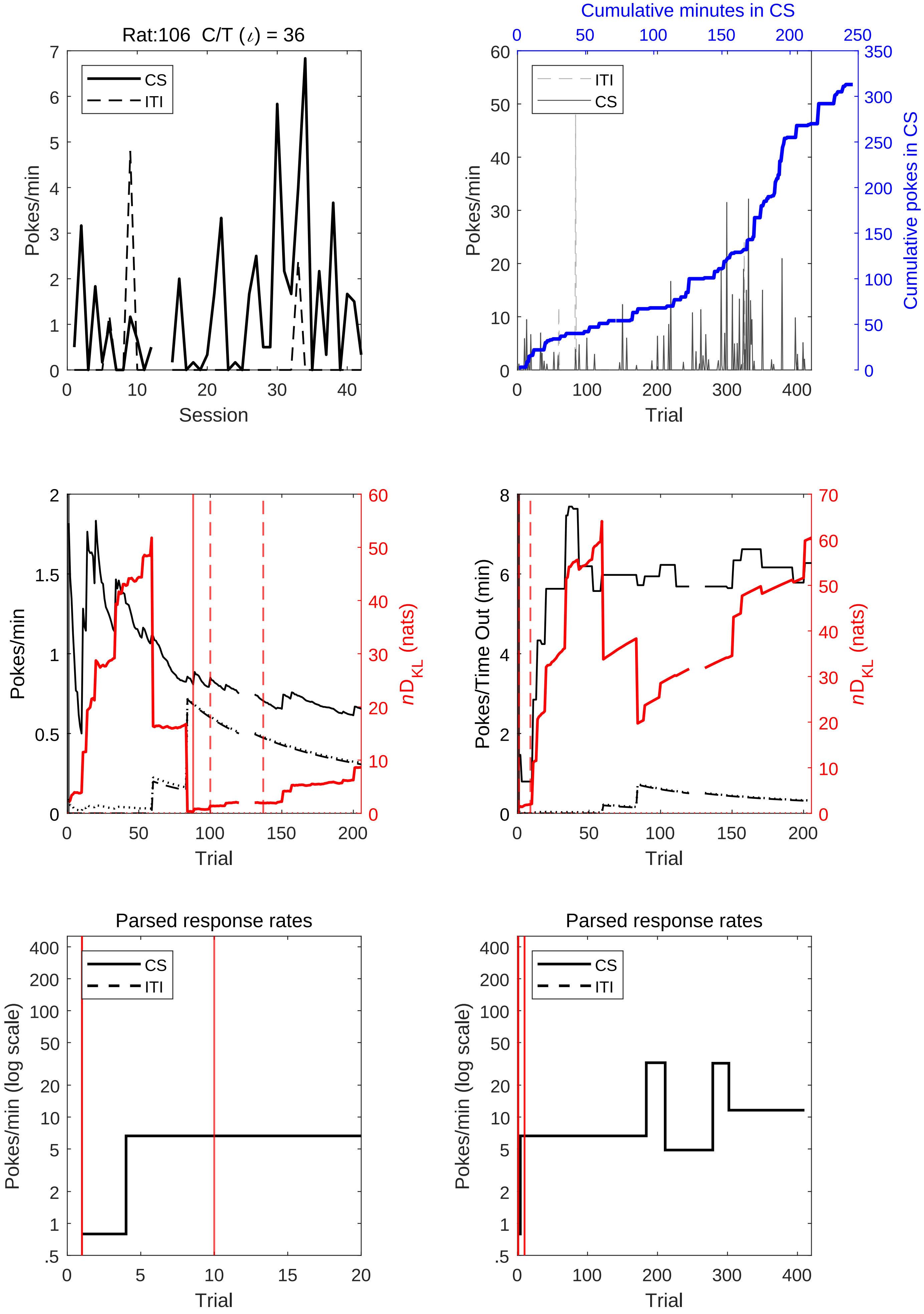
Rat:106 C/T (ι) = 36
CS
ITI
Pokes/min
Session
Cumulative minutes in CS
Cumulative pokes in CS
Trial
nD_KL (nats)
Pokes/Time Out (min)
Parsed response rates
Pokes/min (log scale)

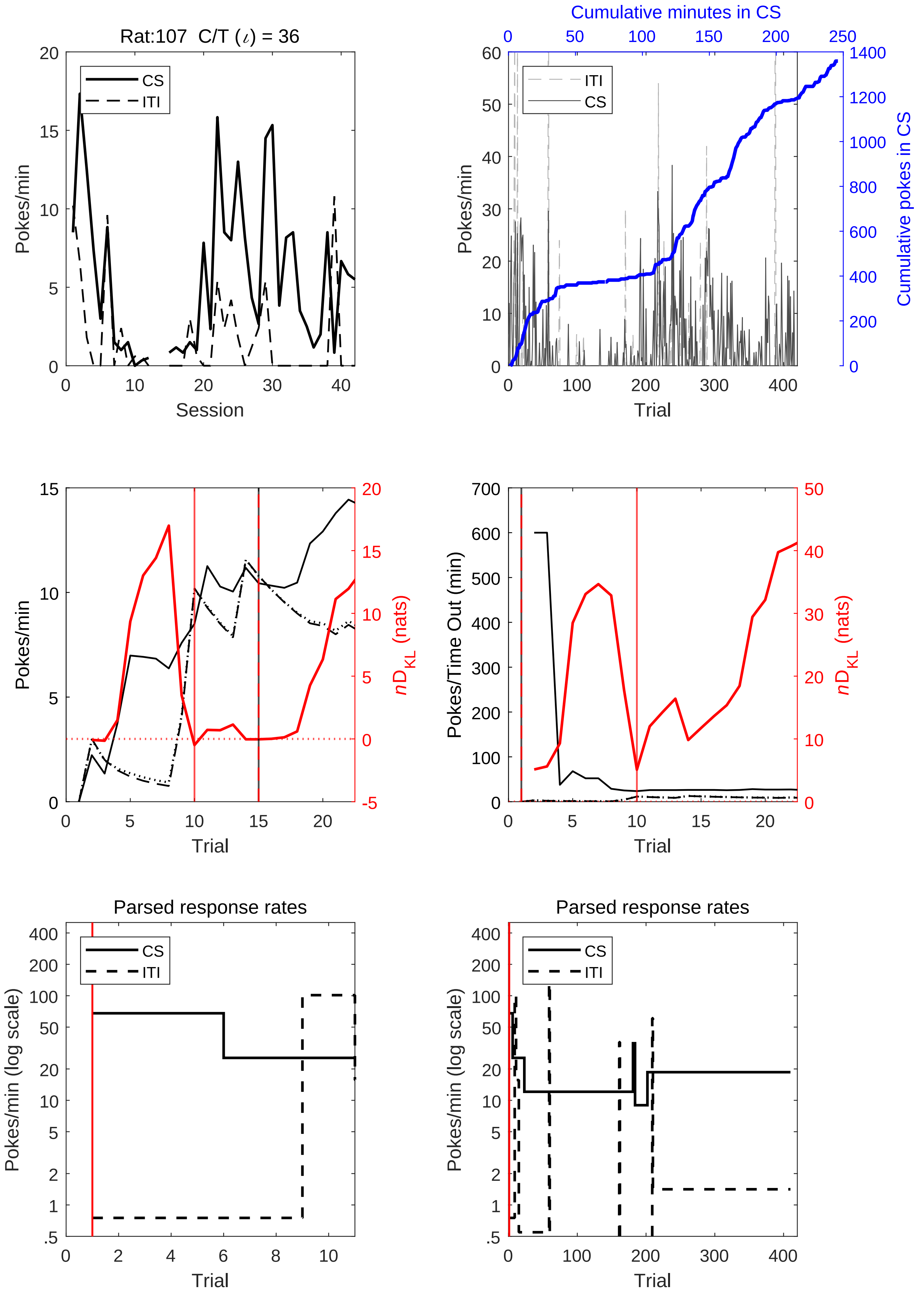

Rat:107 C/T (ι) = 36
CS
ITI
Pokes/min
Session
Cumulative minutes in CS
Cumulative pokes in CS
Trial
$nD_{KL}$ (nats)
Pokes/Time Out (min)
Parsed response rates
Pokes/min (log scale)

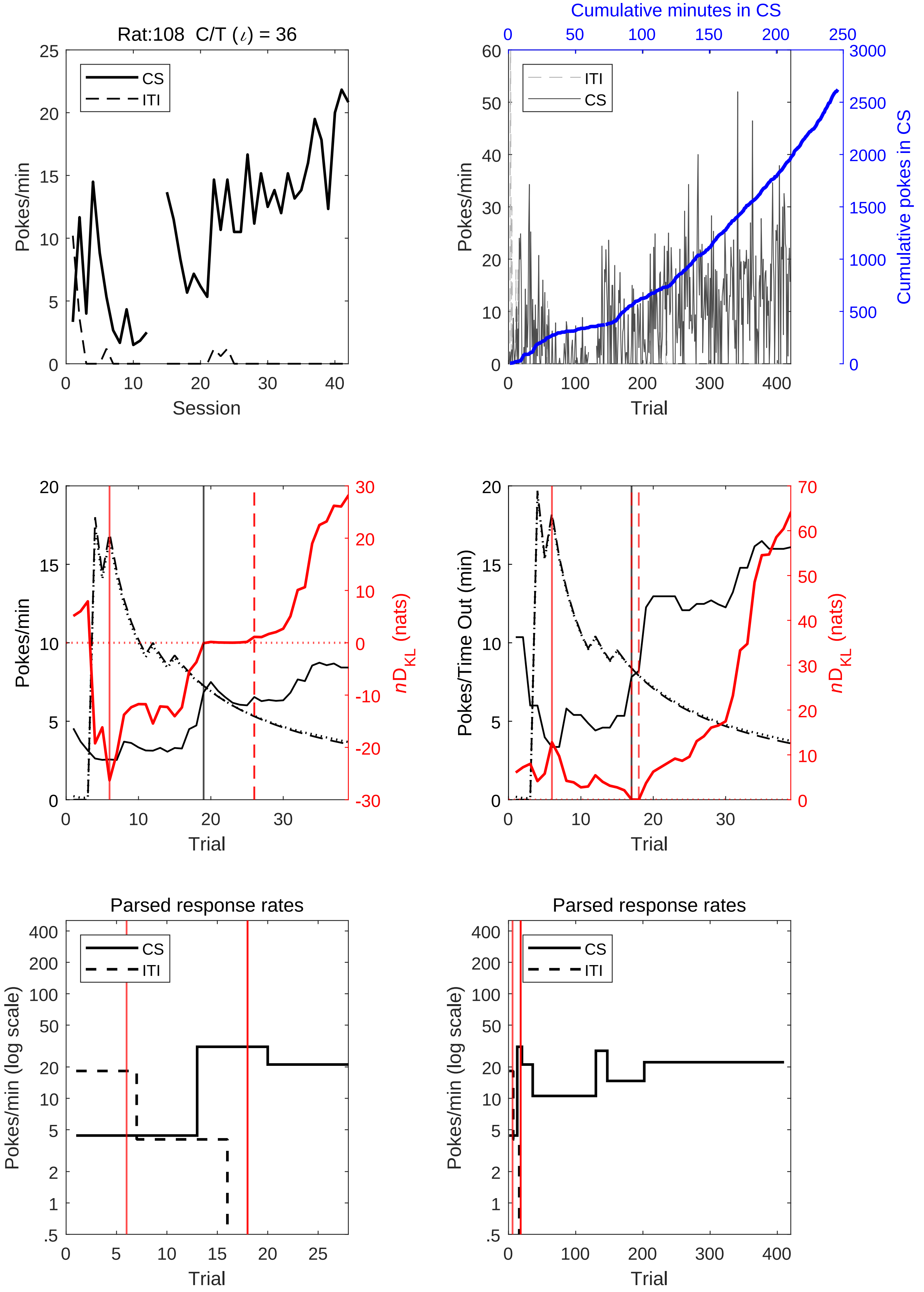
Rat:108 C/T (ι) = 36
CS
ITI
Pokes/min
Session
Cumulative minutes in CS
Cumulative pokes in CS
Trial
nD_KL (nats)
Pokes/Time Out (min)
Parsed response rates
Pokes/min (log scale)

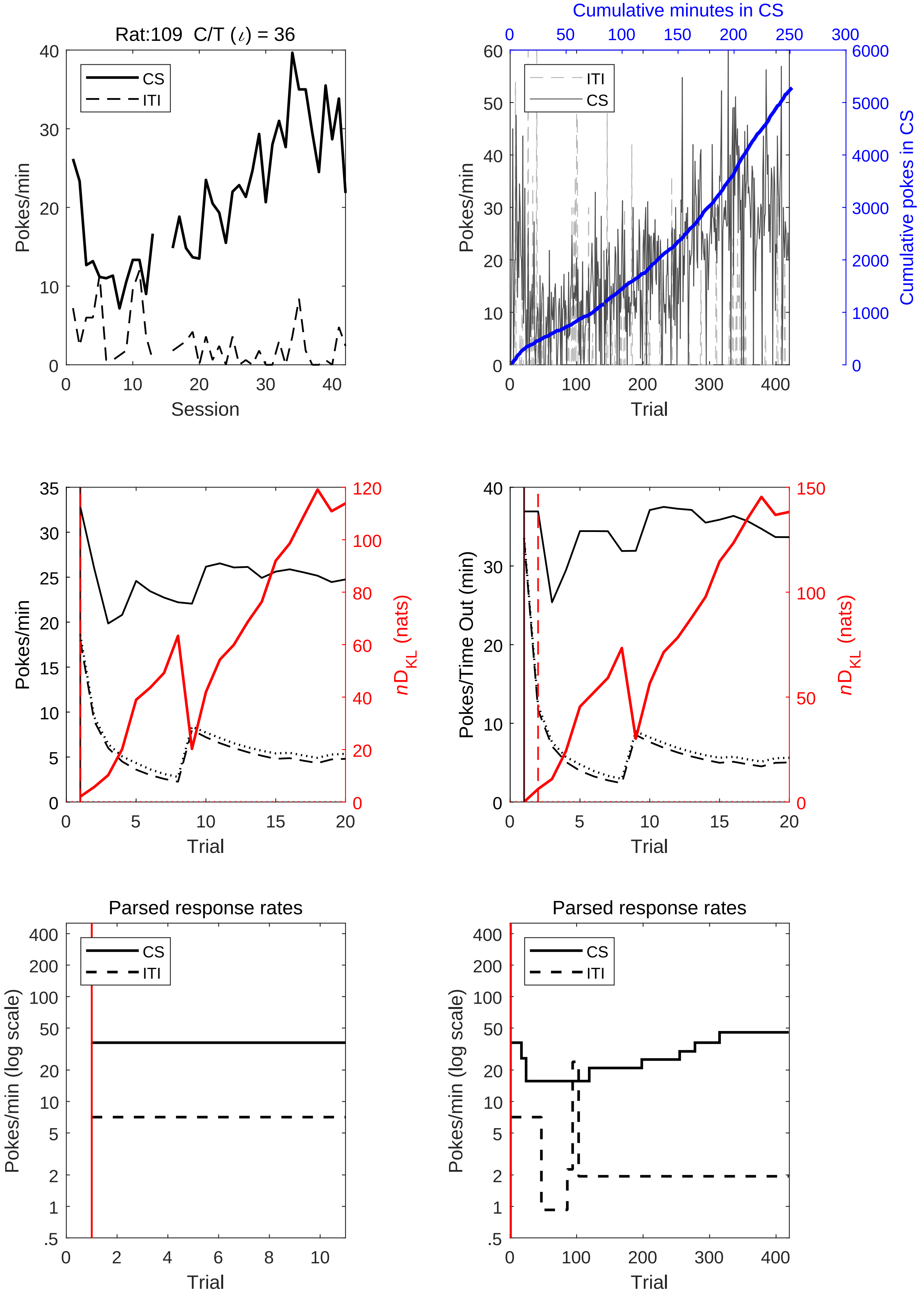
Rat:109 C/T (ι) = 36
CS
ITI
Pokes/min
Session
Cumulative minutes in CS
ITI
CS
Pokes/min
Cumulative pokes in CS
Trial
Pokes/min
$nD_{KL}$ (nats)
Trial
Pokes/Time Out (min)
$nD_{KL}$ (nats)
Trial
Parsed response rates
CS
ITI
Pokes/min (log scale)
Trial
Parsed response rates
CS
ITI
Pokes/min (log scale)
Trial

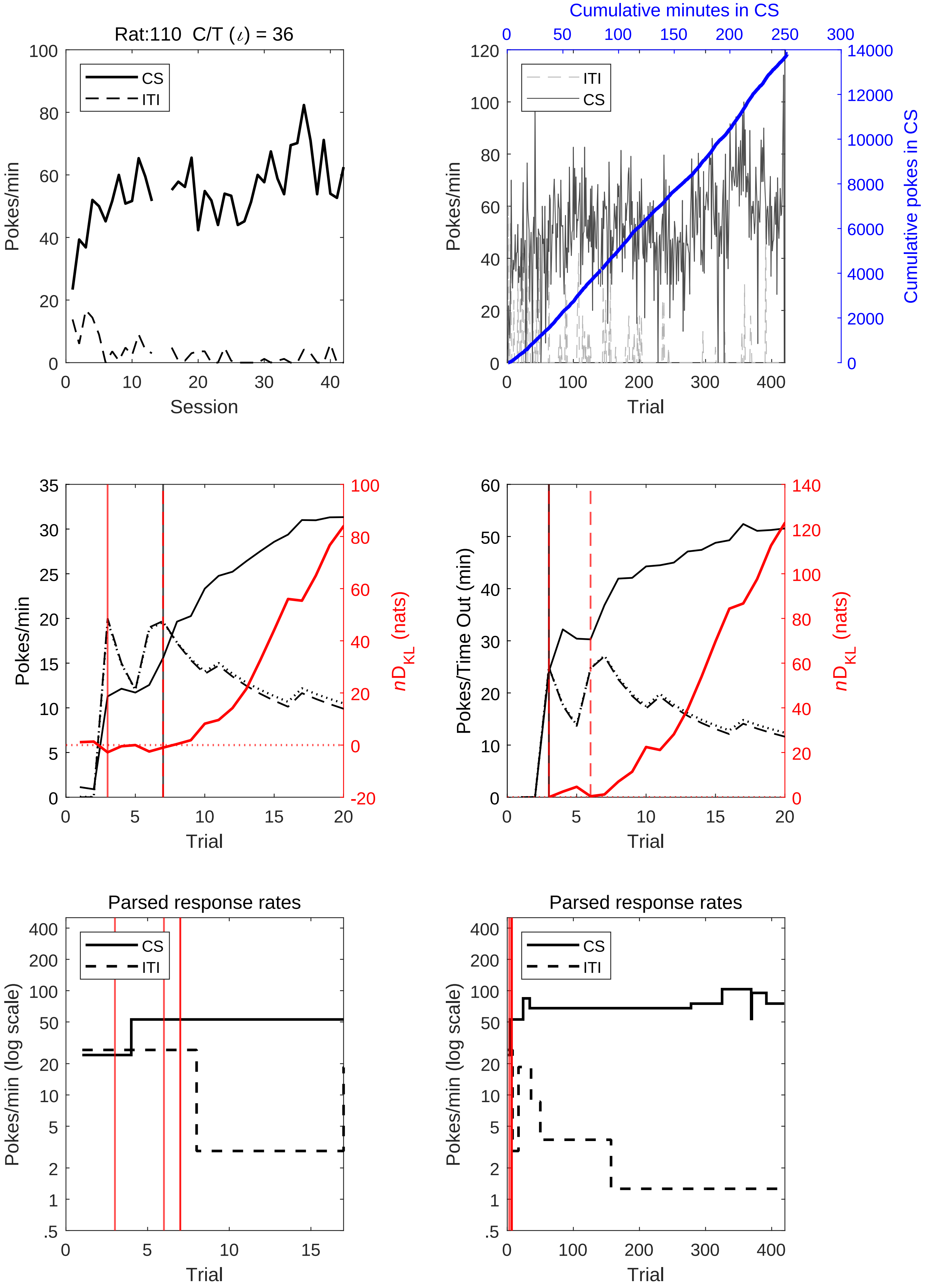
Rat:110 C/T (ι) = 36
CS
ITI
Pokes/min
Session
Cumulative minutes in CS
Cumulative pokes in CS
Trial
Pokes/Time Out (min)
nD_KL (nats)
Parsed response rates
Pokes/min (log scale)

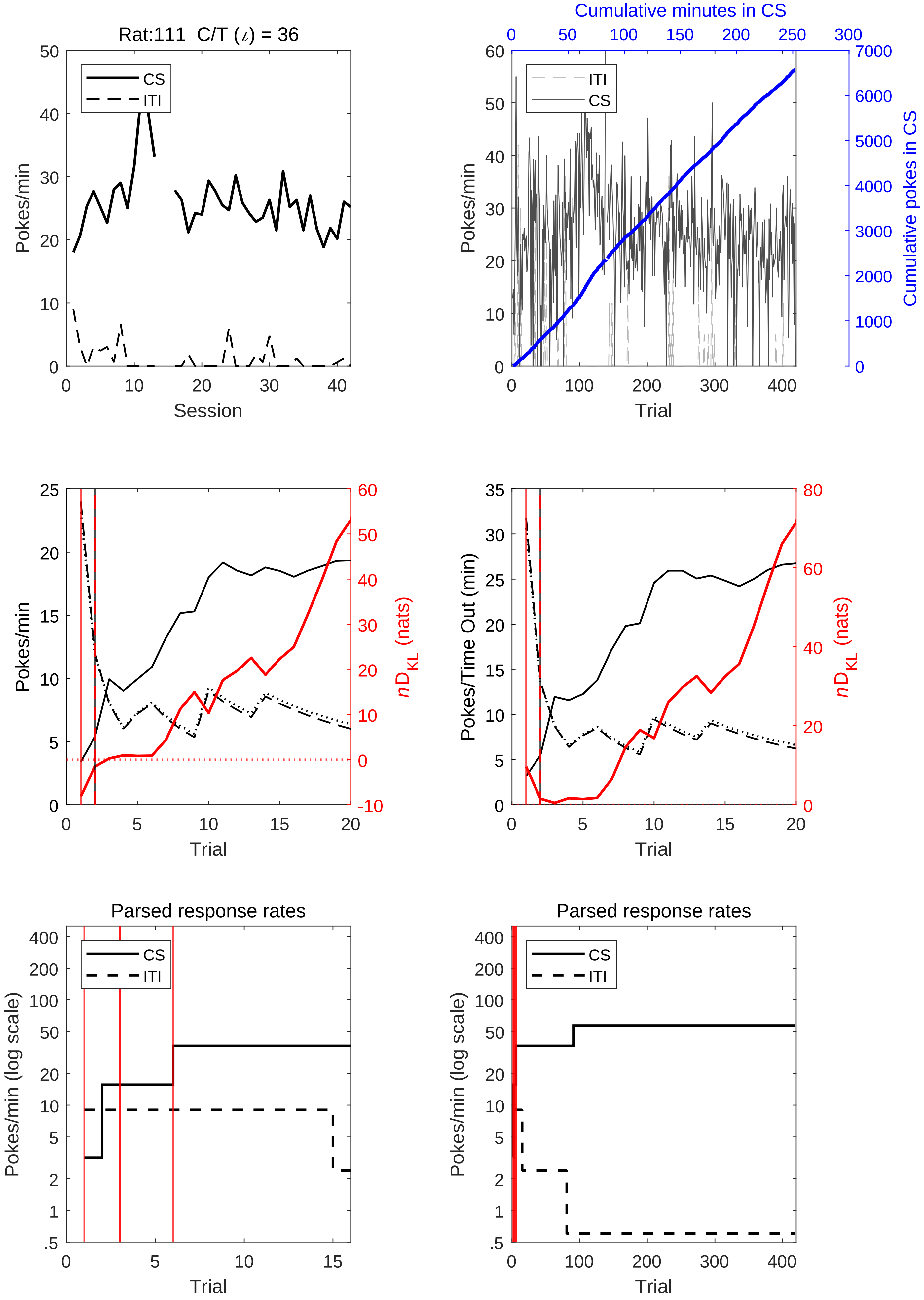
Rat:111 C/T (ι) = 36
CS
ITI
Pokes/min
Session
Cumulative minutes in CS
Cumulative pokes in CS
Trial
Pokes/Time Out (min)
nD_KL (nats)
Parsed response rates
Pokes/min (log scale)

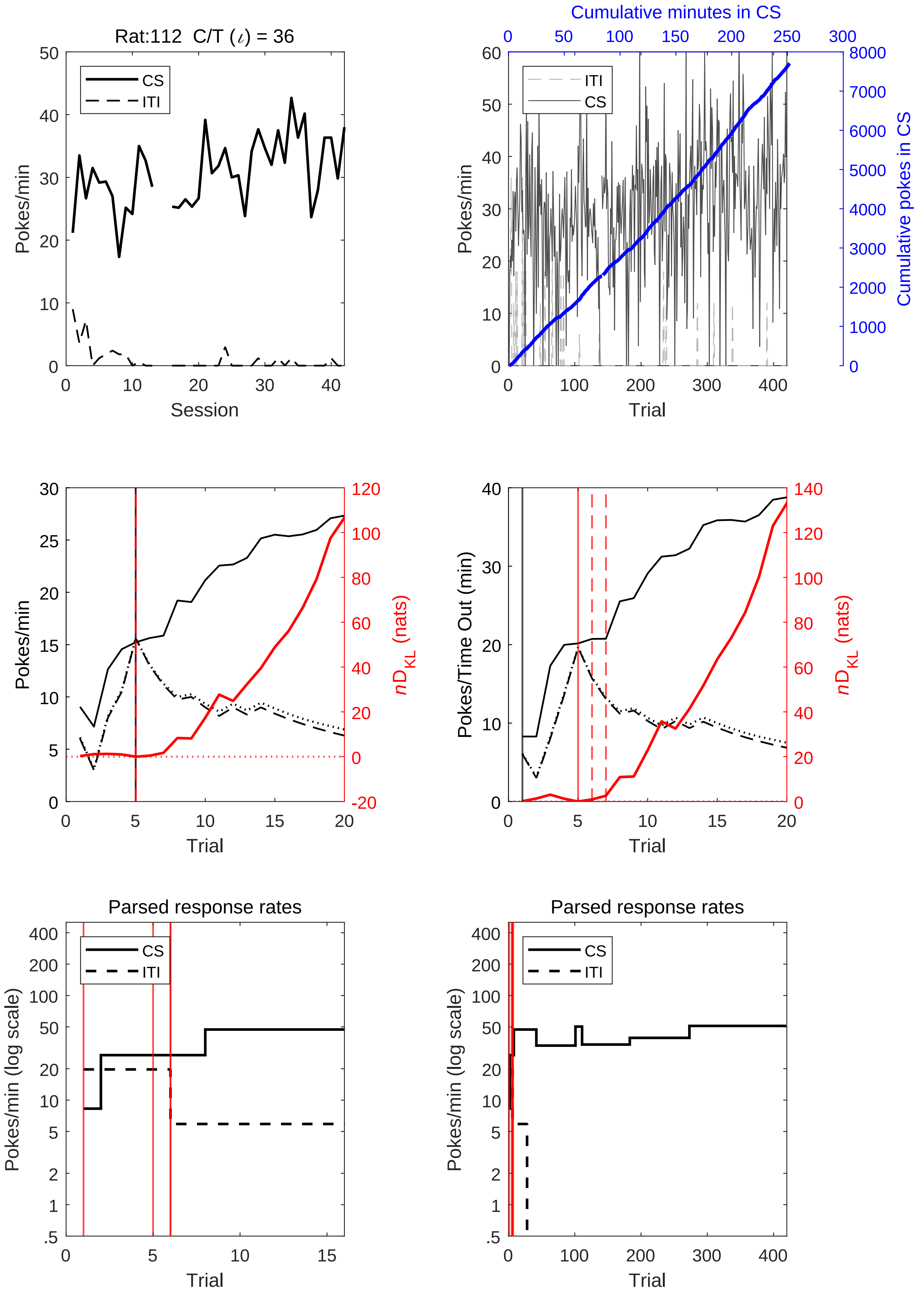
Rat:112 C/T (ι) = 36
CS
ITI
Pokes/min
Session
Cumulative minutes in CS
Cumulative pokes in CS
Trial
nD_KL (nats)
Pokes/Time Out (min)
Parsed response rates
Pokes/min (log scale)

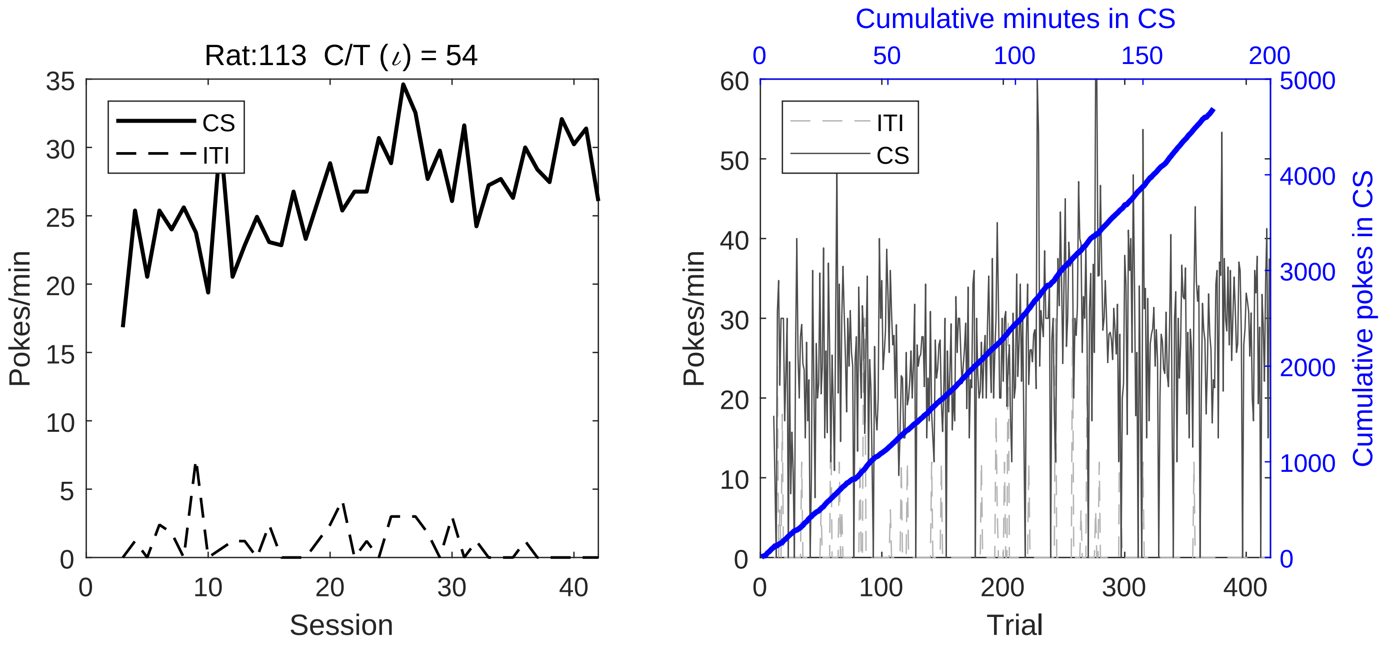
Rat:113  C/T (ι) = 54
CS
ITI
Pokes/min
Session
Cumulative minutes in CS
Trial
Cumulative pokes in CS

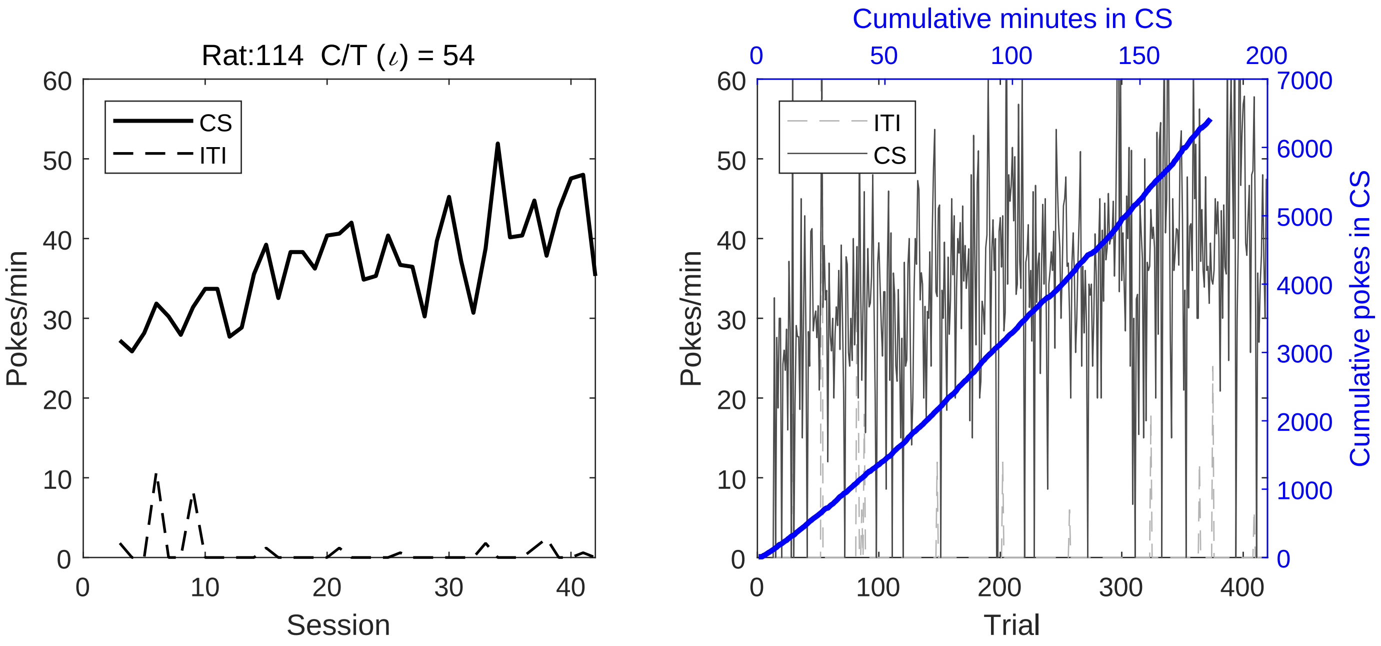

Rat:114 C/T ($\iota$) = 54
CS
ITI
Pokes/min
Session
Cumulative minutes in CS
ITI
CS
Pokes/min
Trial
Cumulative pokes in CS

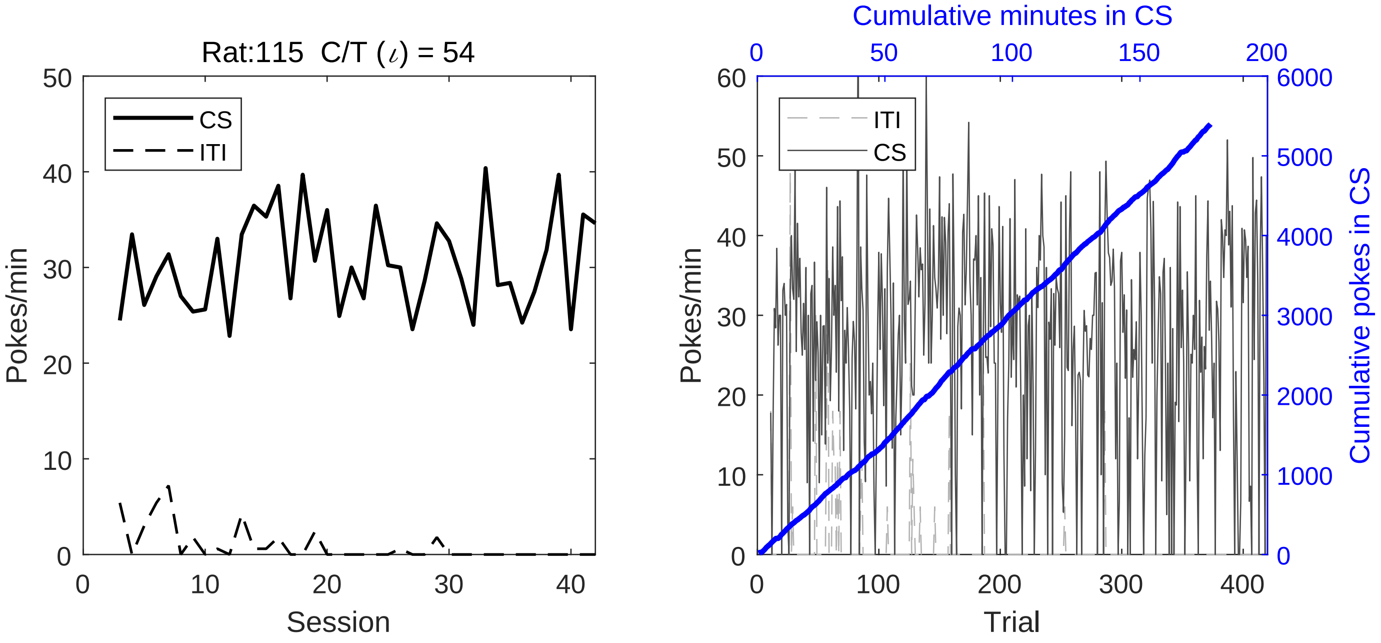
Rat:115 C/T ($\iota$) = 54
CS
ITI
Pokes/min
Session
Cumulative minutes in CS
ITI
CS
Pokes/min
Cumulative pokes in CS
Trial

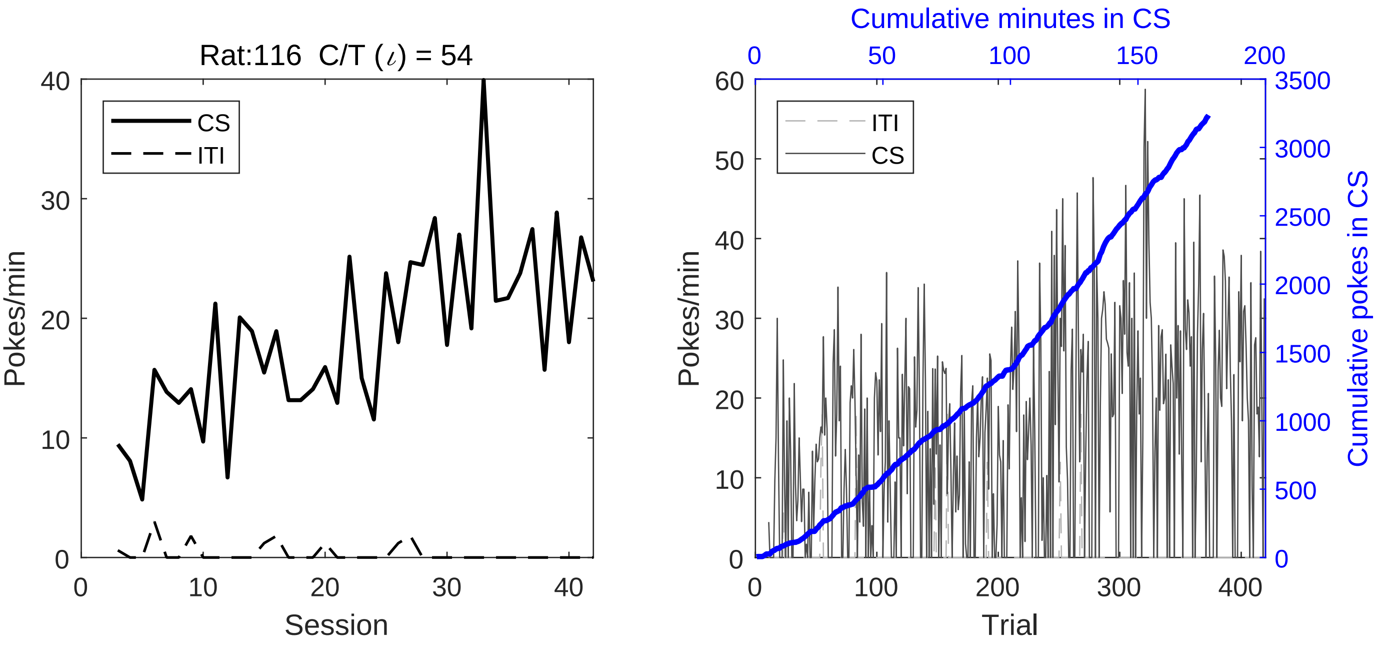
Rat:116 C/T (ι) = 54
CS
ITI
Pokes/min
Session
Cumulative minutes in CS
ITI
CS
Pokes/min
Trial
Cumulative pokes in CS

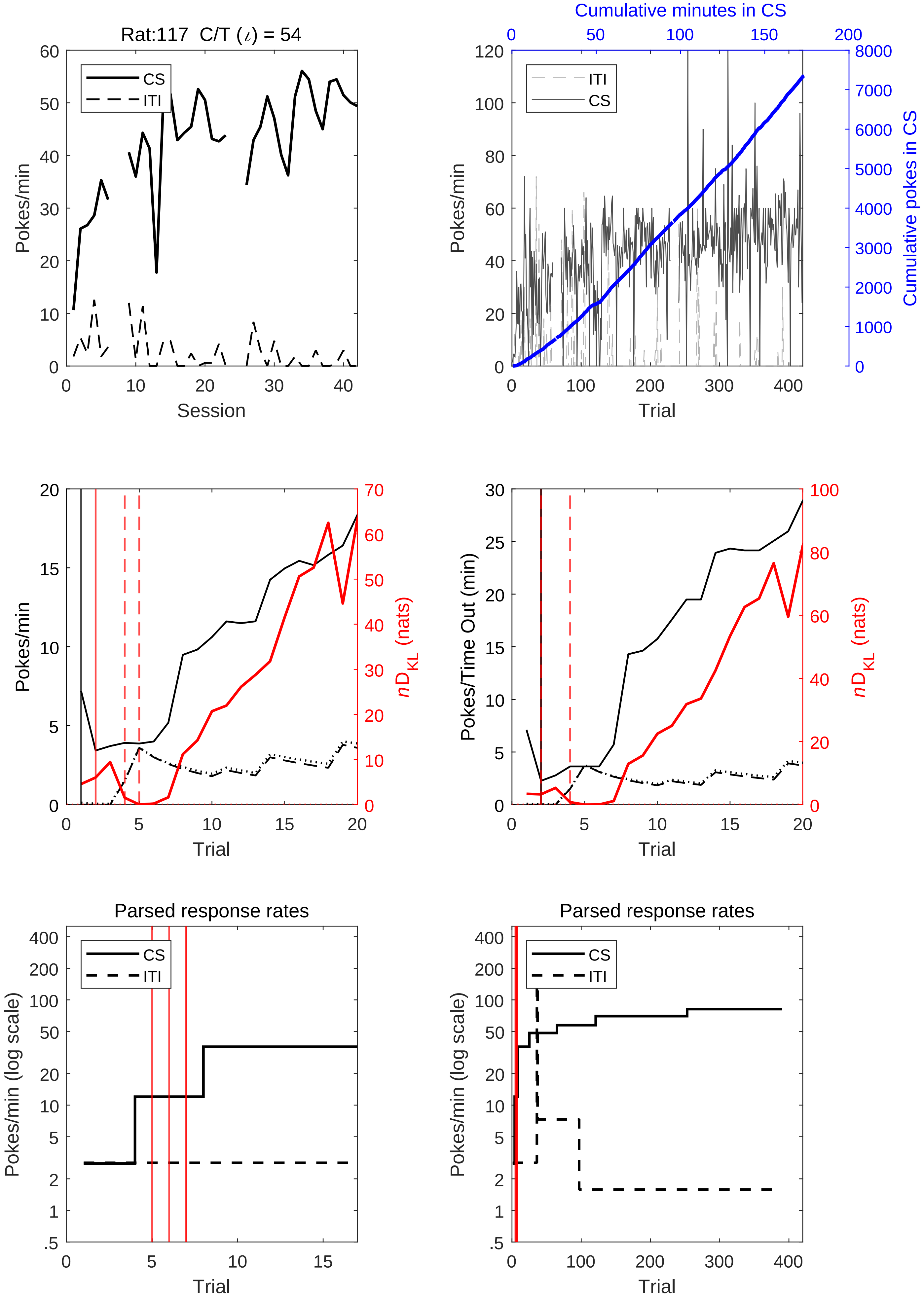
Rat:117 C/T (ι) = 54
CS
ITI
Pokes/min
Session
Cumulative minutes in CS
Cumulative pokes in CS
Trial
Pokes/Time Out (min)
nD_KL (nats)
Parsed response rates
Pokes/min (log scale)

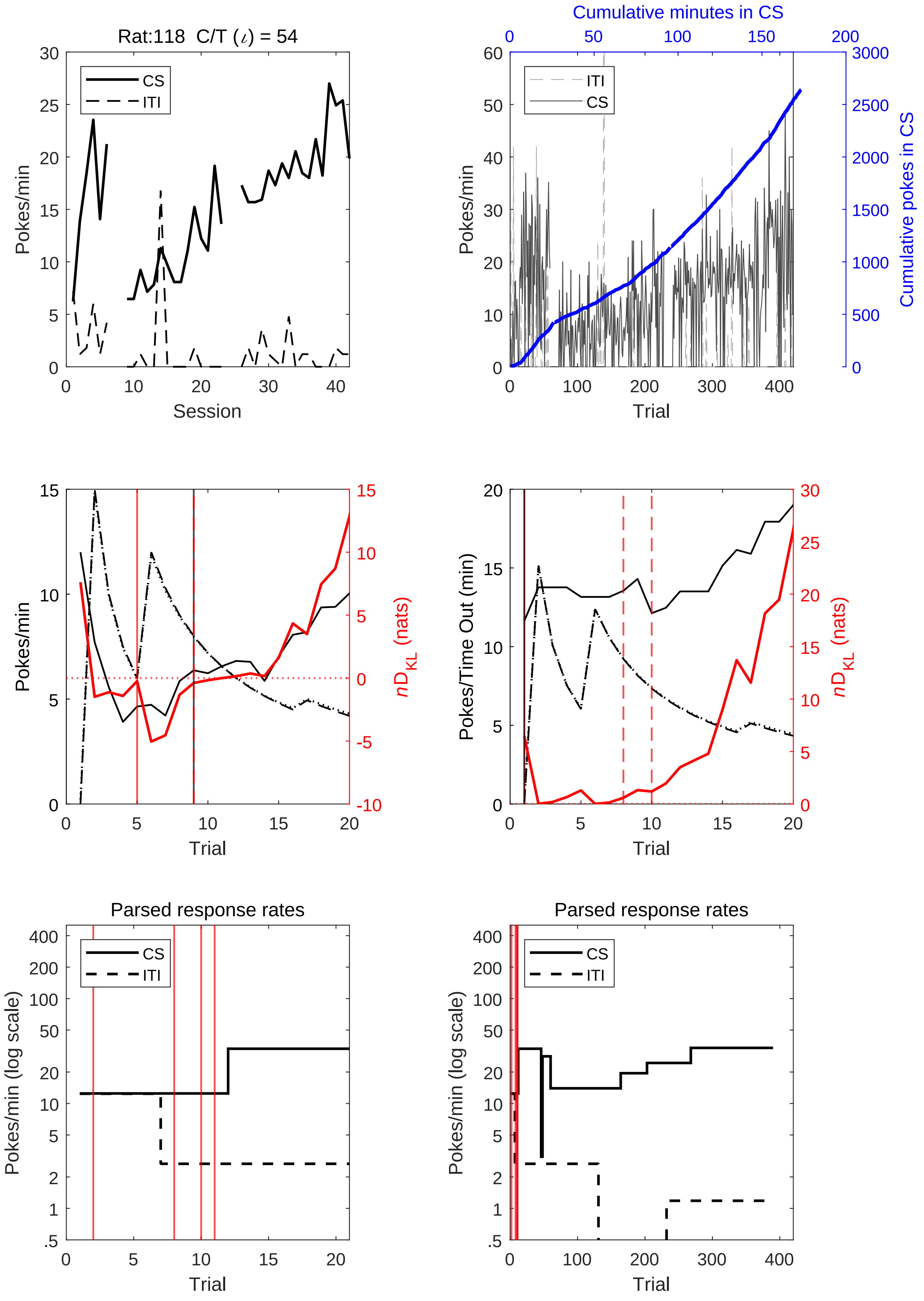

Rat:118 C/T (ι) = 54
CS
ITI
Pokes/min
Session
Cumulative minutes in CS
Cumulative pokes in CS
Trial
Pokes/Time Out (min)
nD_KL (nats)
Parsed response rates
Pokes/min (log scale)

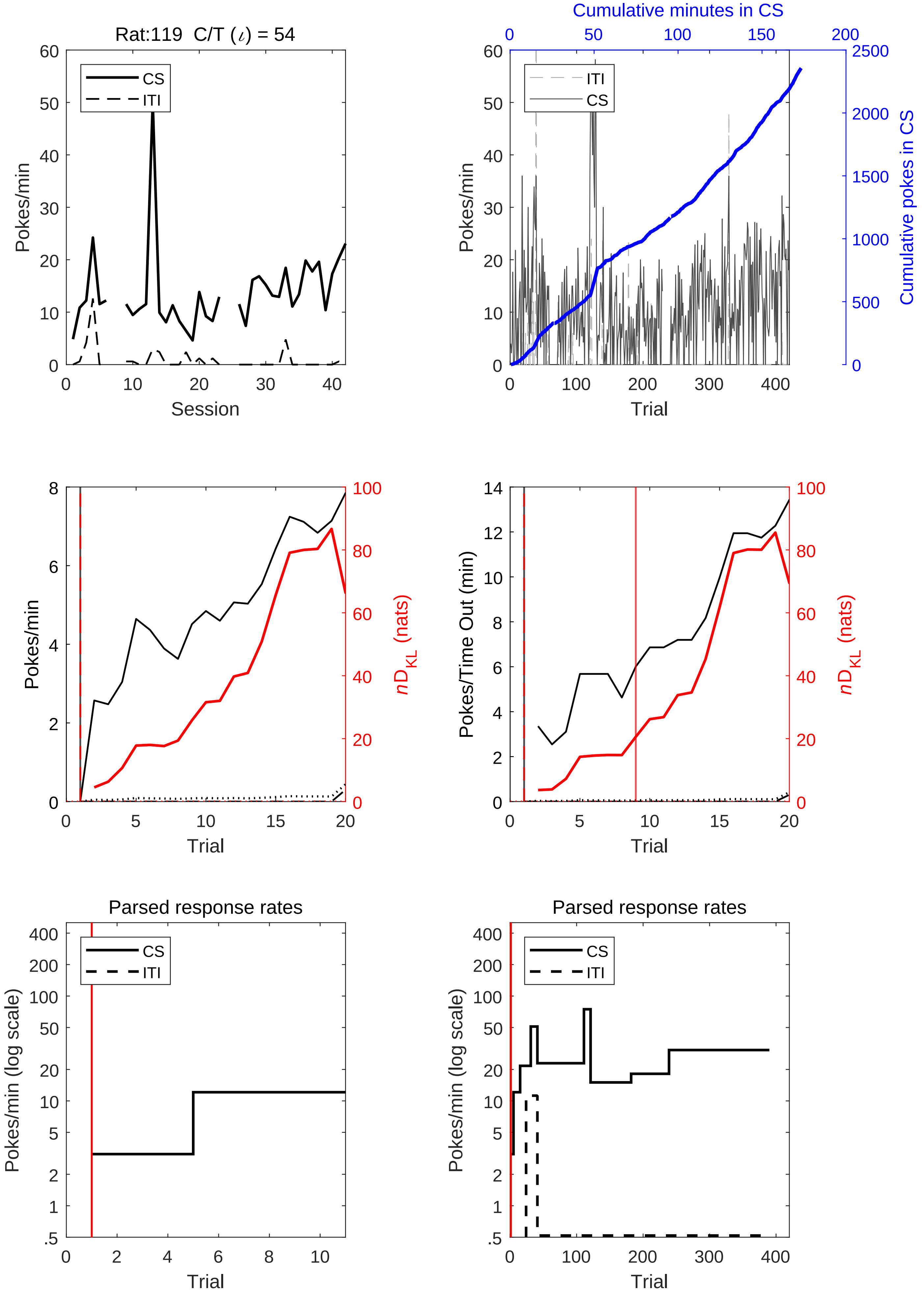
Rat:119 C/T ($\iota$) = 54
CS
ITI
Pokes/min
Session
Cumulative minutes in CS
ITI
CS
Pokes/min
Cumulative pokes in CS
Trial
Pokes/min
$nD_{KL}$ (nats)
Trial
Pokes/Time Out (min)
$nD_{KL}$ (nats)
Trial
Parsed response rates
CS
ITI
Pokes/min (log scale)
Trial
Parsed response rates
CS
ITI
Pokes/min (log scale)
Trial

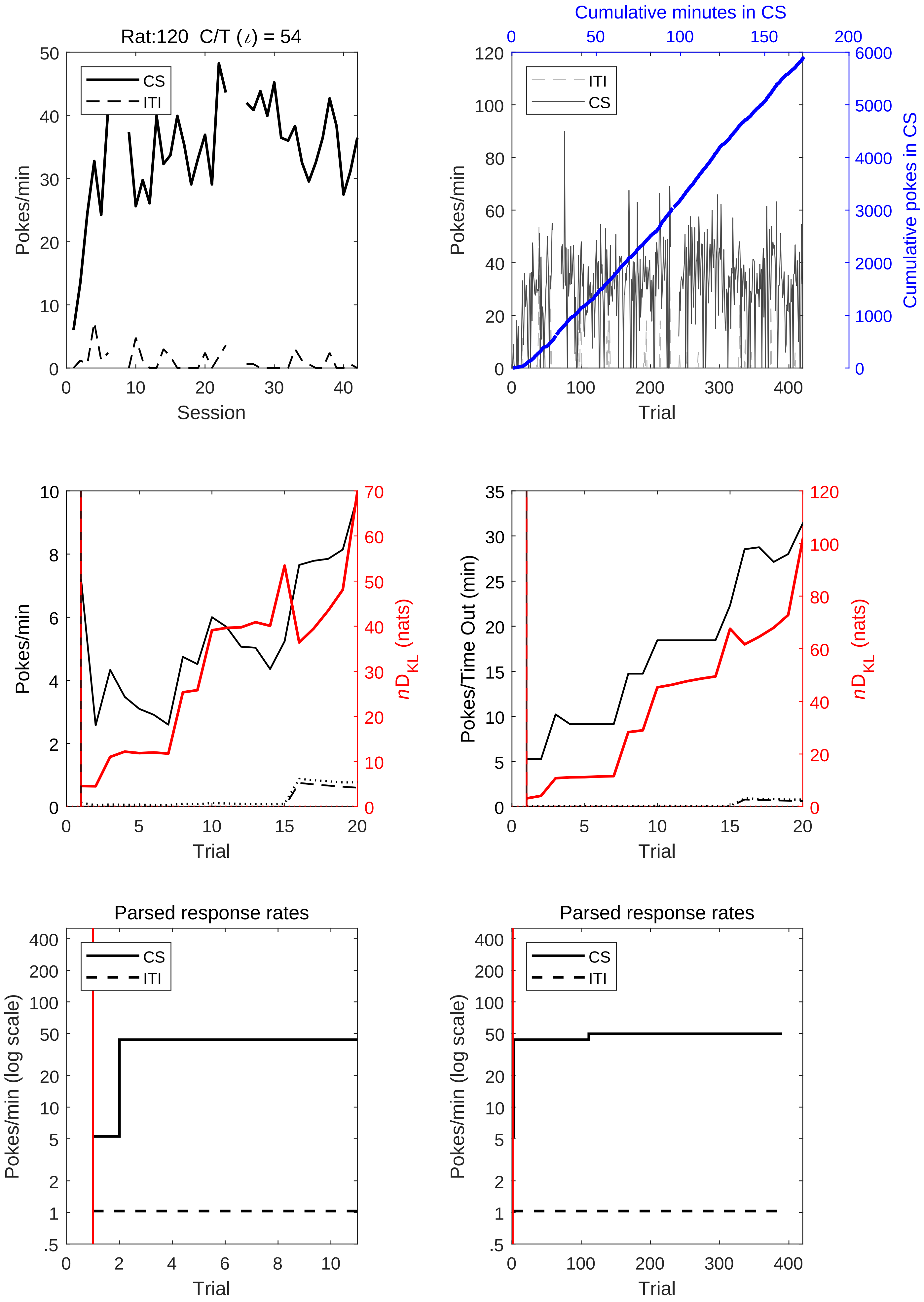
Rat:120 C/T (ι) = 54
CS
ITI
Pokes/min
Session
Cumulative minutes in CS
ITI
CS
Pokes/min
Trial
Cumulative pokes in CS
Pokes/min
Trial
nD_KL (nats)
Pokes/Time Out (min)
Trial
nD_KL (nats)
Parsed response rates
CS
ITI
Pokes/min (log scale)
Trial
Parsed response rates
CS
ITI
Pokes/min (log scale)
Trial

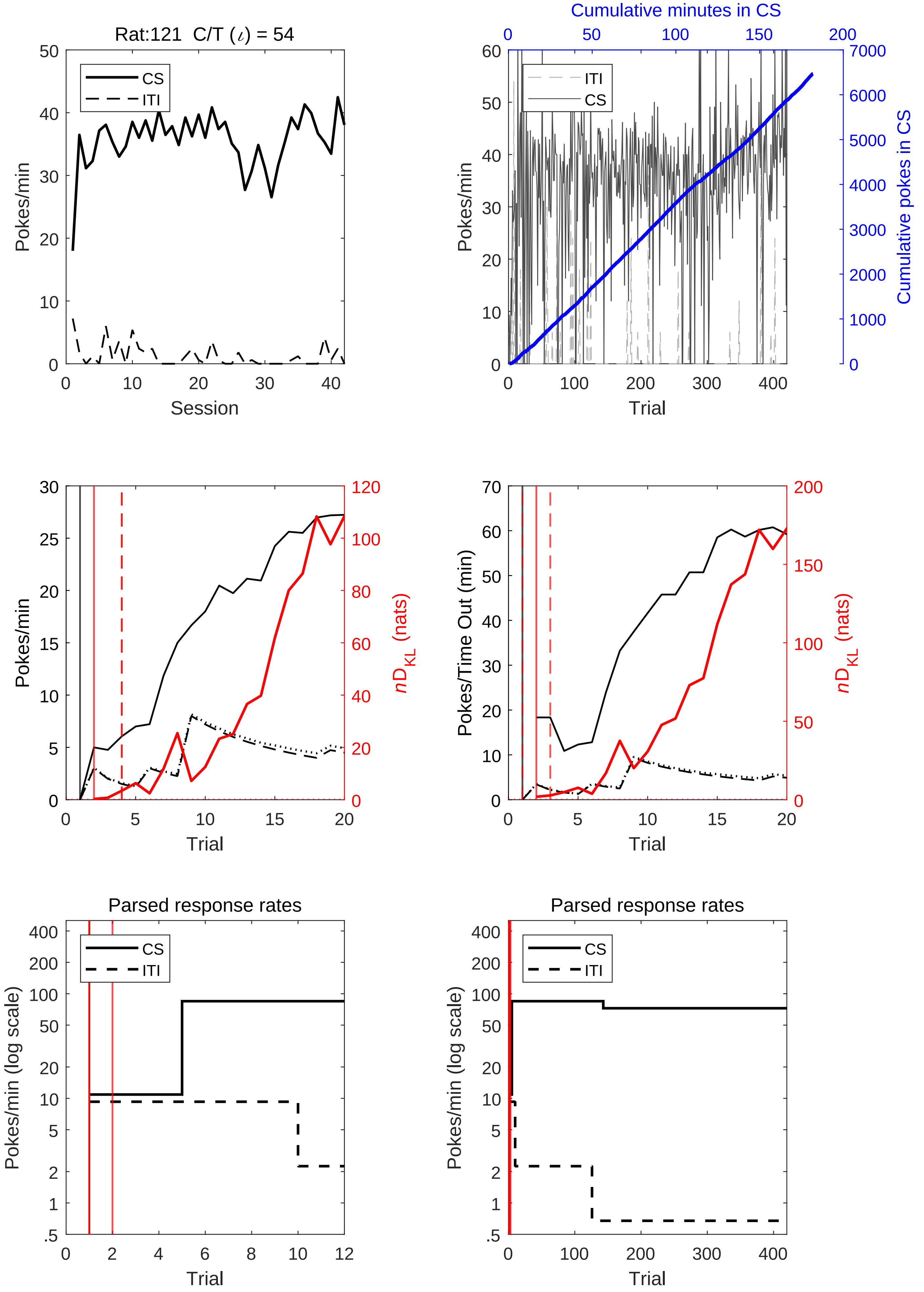

Rat:121 C/T (ι) = 54
CS
ITI
Pokes/min
Session
Cumulative minutes in CS
Cumulative pokes in CS
Trial
nD_KL (nats)
Pokes/Time Out (min)
Parsed response rates
Pokes/min (log scale)

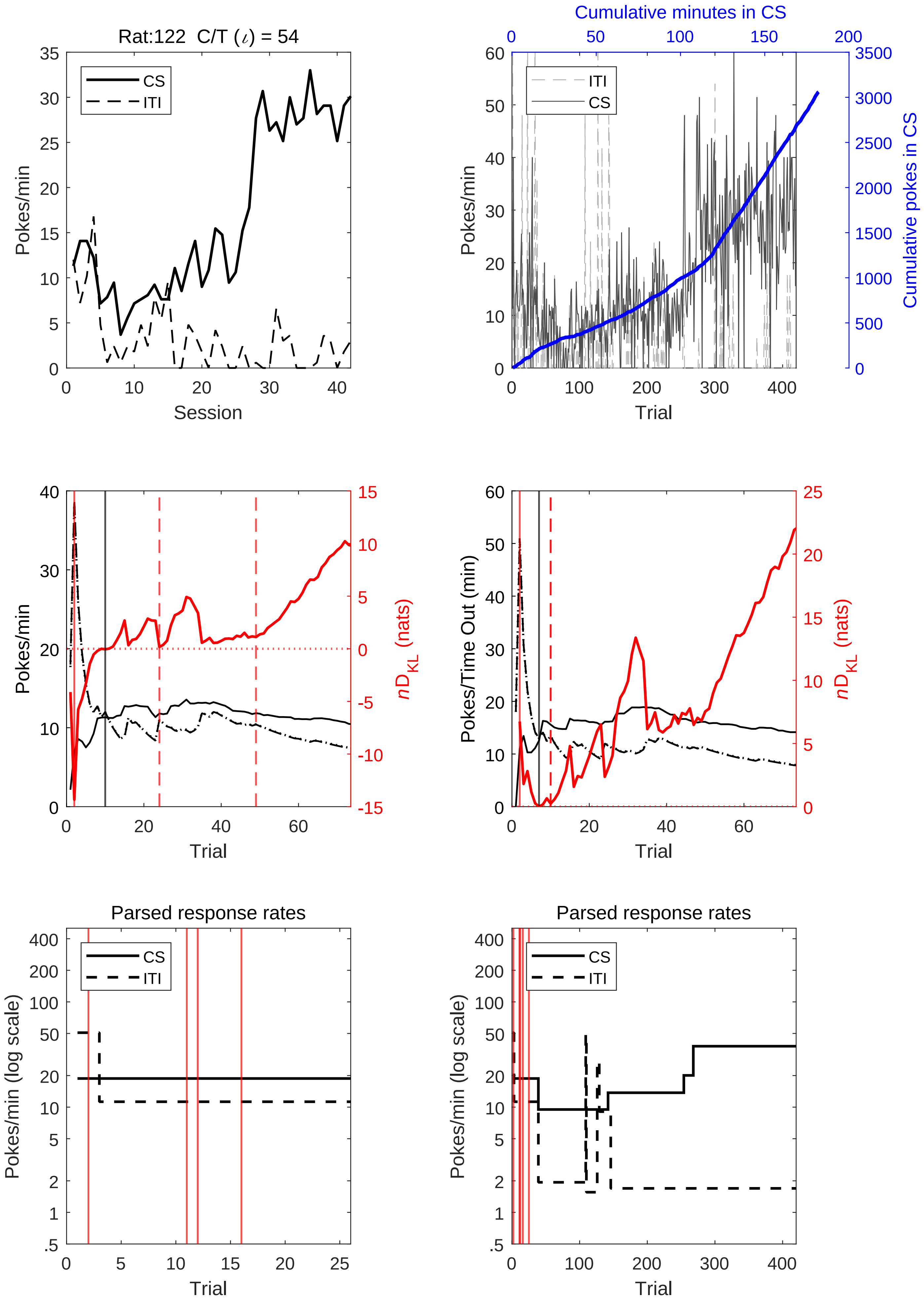
Rat:122 C/T (ι) = 54
CS
ITI
Pokes/min
Session
Cumulative minutes in CS
Cumulative pokes in CS
Trial
nD_KL (nats)
Pokes/Time Out (min)
Parsed response rates
Pokes/min (log scale)

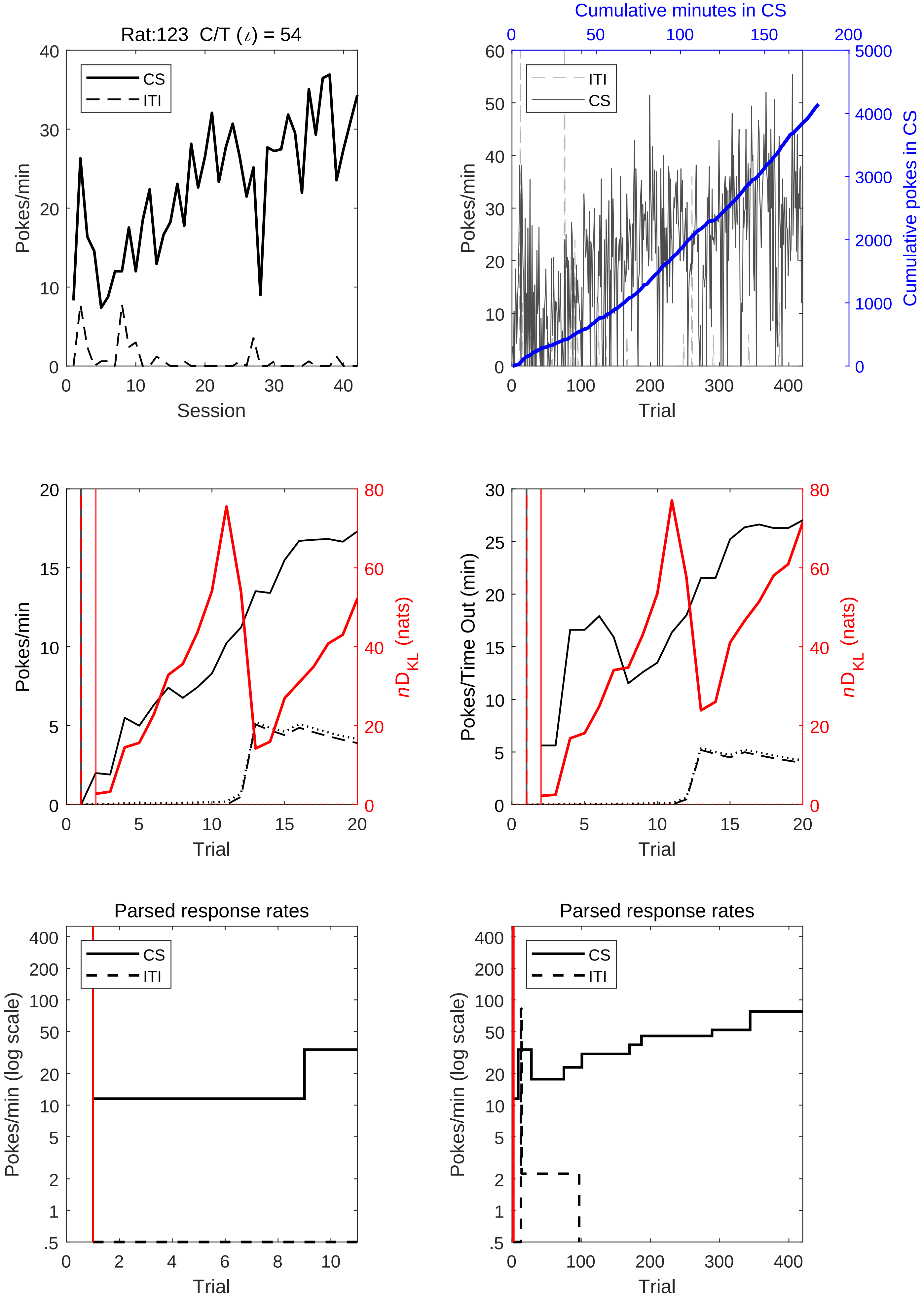

Rat:123 C/T (ι) = 54
CS
ITI
Pokes/min
Session
Cumulative minutes in CS
Cumulative pokes in CS
Trial
nD_KL (nats)
Pokes/Time Out (min)
Parsed response rates
Pokes/min (log scale)

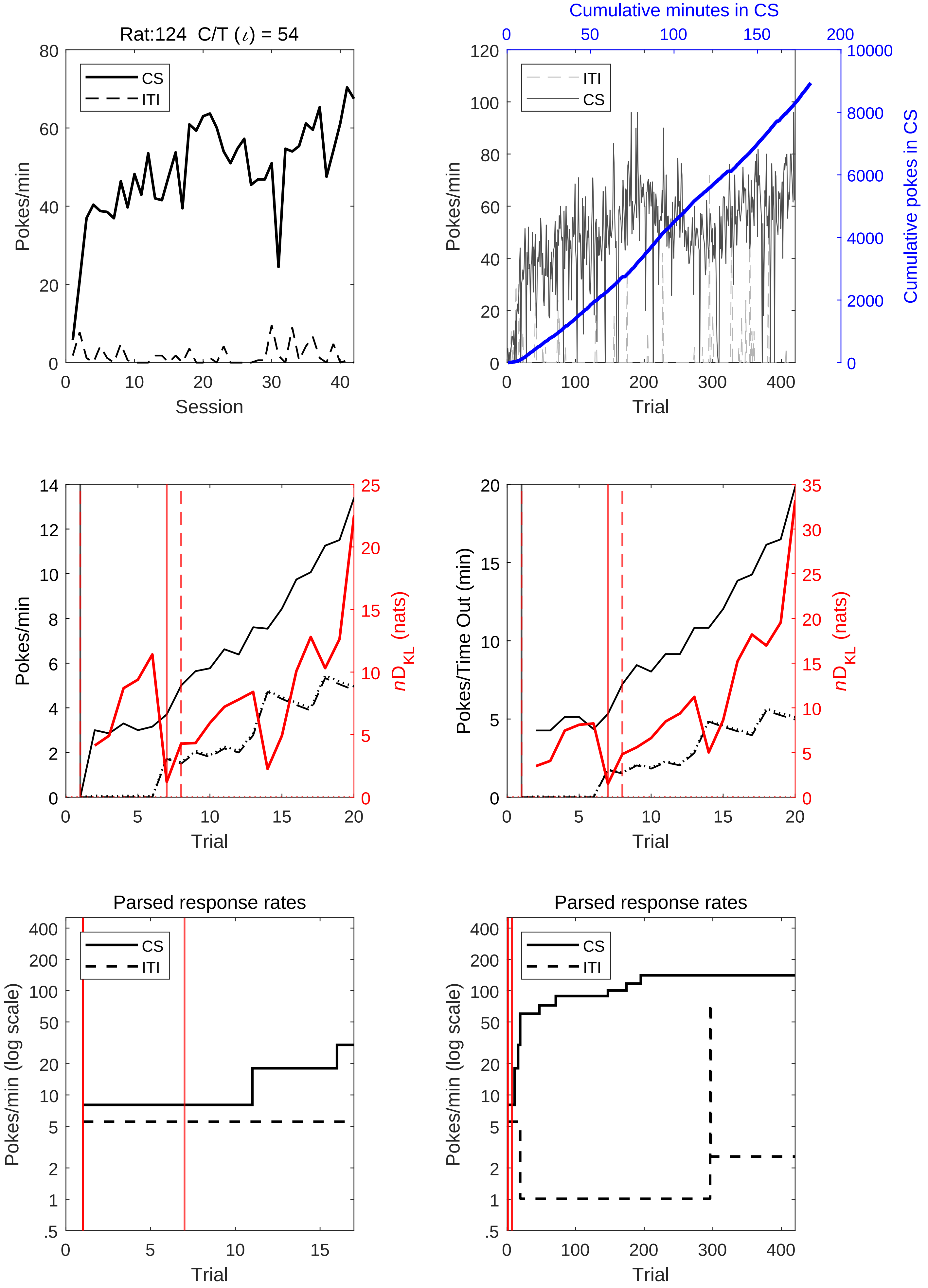
Rat:124 C/T (ι) = 54
CS
ITI
Pokes/min
Session
Cumulative minutes in CS
Cumulative pokes in CS
Trial
nD_KL (nats)
Pokes/Time Out (min)
Parsed response rates
Pokes/min (log scale)

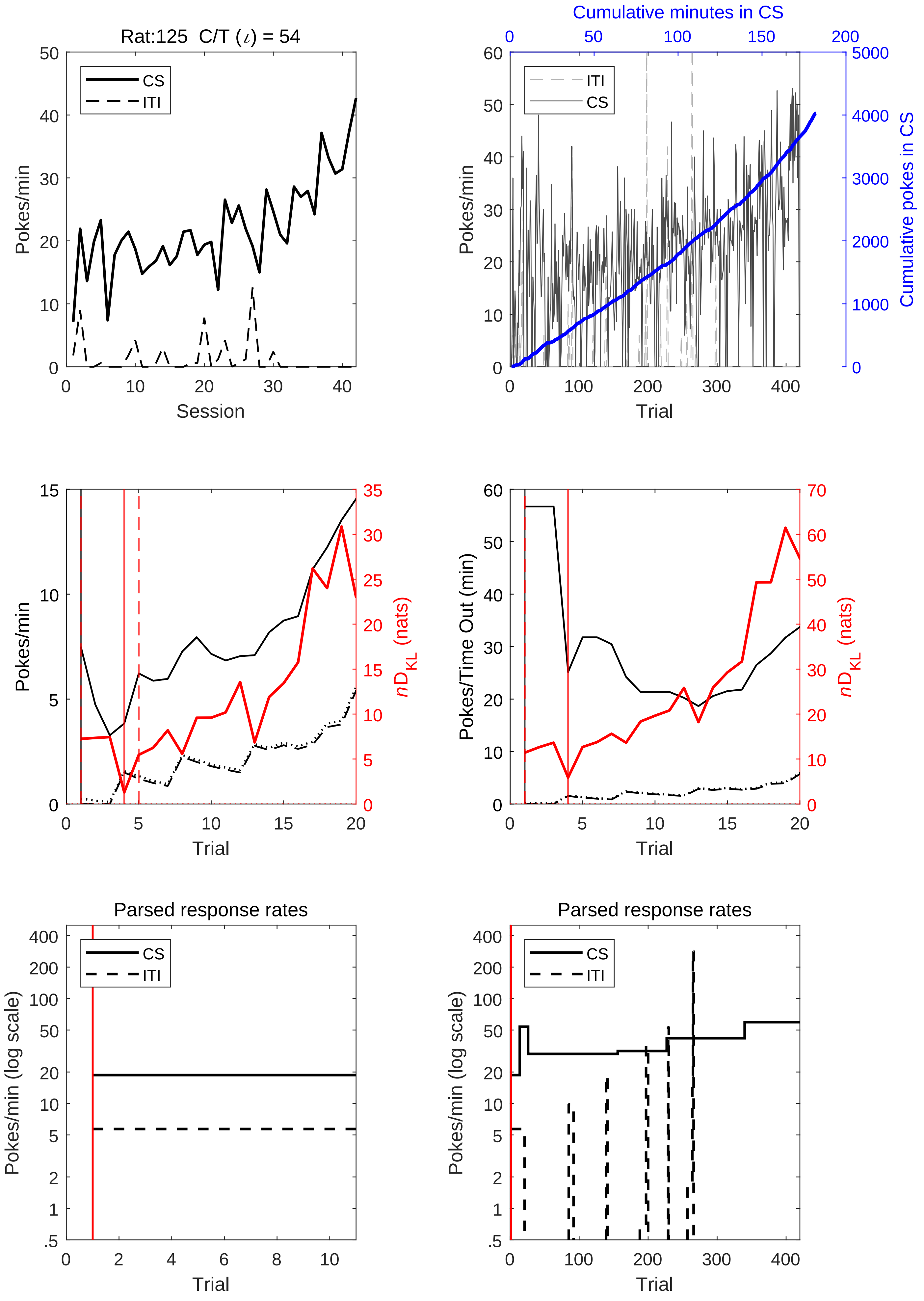
Rat:125 C/T (ι) = 54
CS
ITI
Pokes/min
Session
Cumulative minutes in CS
Cumulative pokes in CS
Trial
nD_KL (nats)
Pokes/Time Out (min)
Parsed response rates
Pokes/min (log scale)

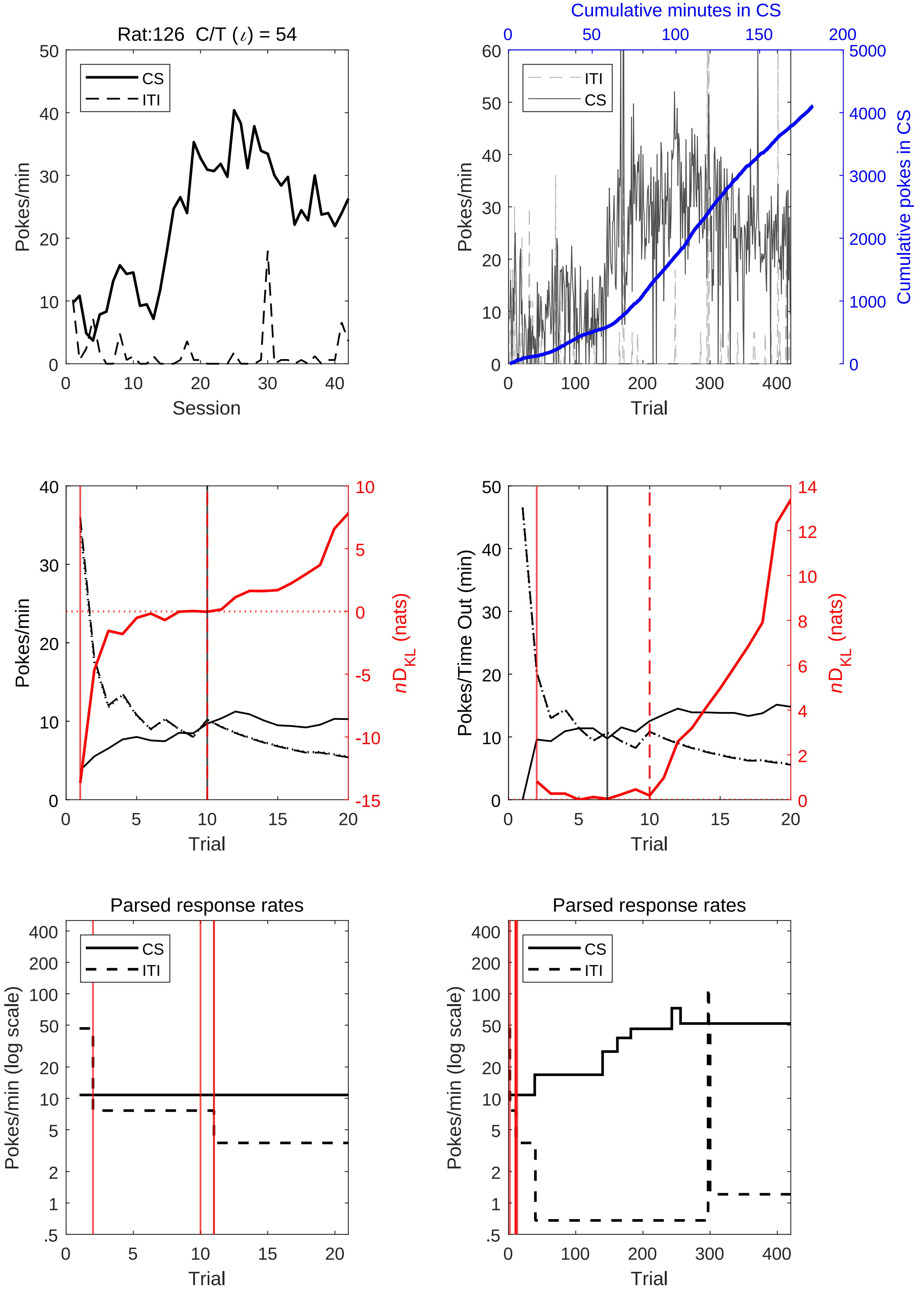
Rat:126 C/T (ι) = 54
CS
ITI
Pokes/min
Session
Cumulative minutes in CS
Cumulative pokes in CS
Trial
nD_KL (nats)
Pokes/Time Out (min)
Parsed response rates
Pokes/min (log scale)

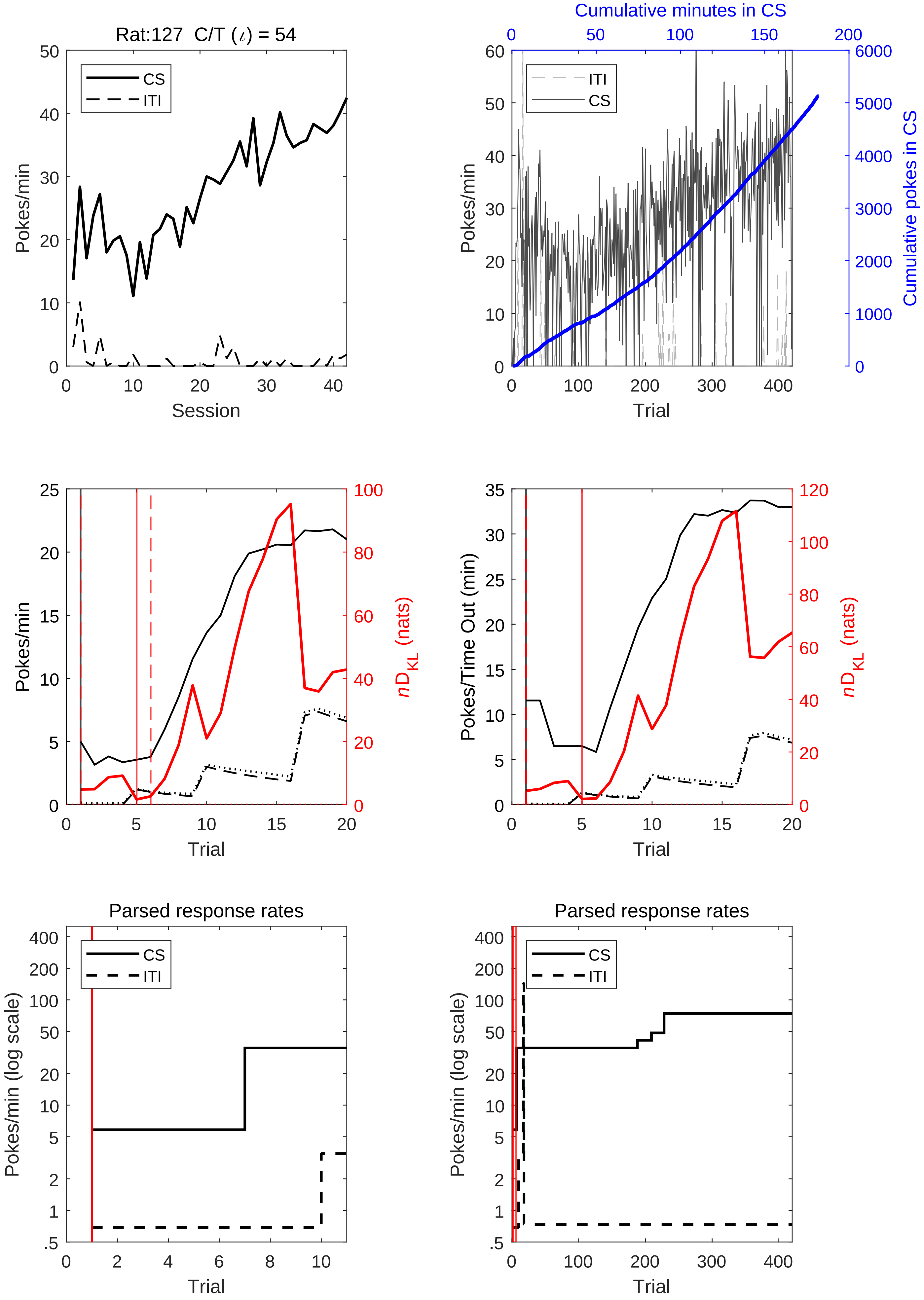
Rat:127 C/T (ι) = 54
CS
ITI
Pokes/min
Session
Cumulative minutes in CS
Cumulative pokes in CS
Trial
Pokes/Time Out (min)
nD_KL (nats)
Parsed response rates
Pokes/min (log scale)

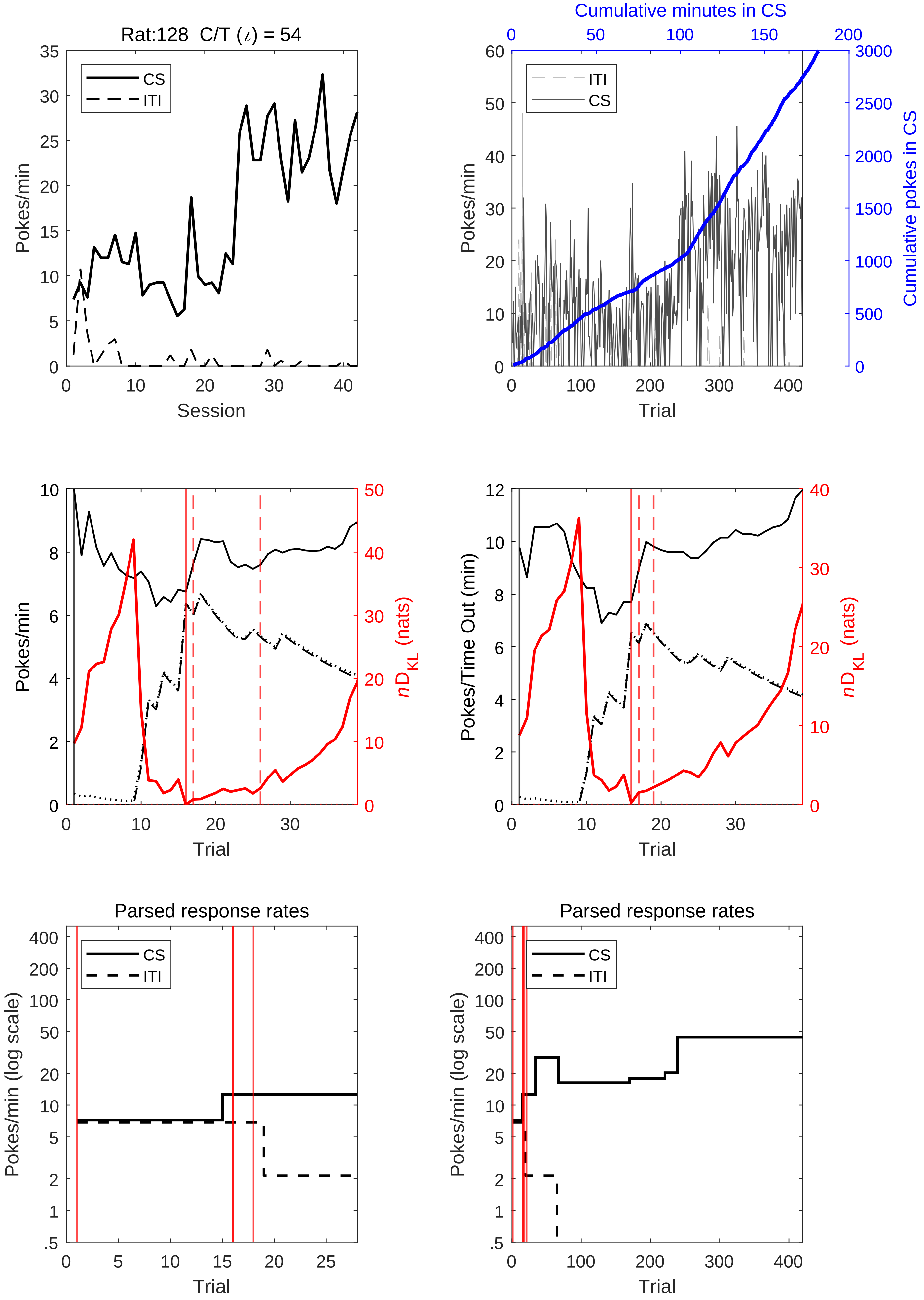

Rat:128 C/T (ι) = 54
CS
ITI
Pokes/min
Session
Cumulative minutes in CS
Cumulative pokes in CS
Pokes/min
Trial
Pokes/min
$n$D$_{KL}$ (nats)
Trial
Pokes/Time Out (min)
$n$D$_{KL}$ (nats)
Trial
Parsed response rates
Pokes/min (log scale)
Trial
Parsed response rates
Pokes/min (log scale)
Trial

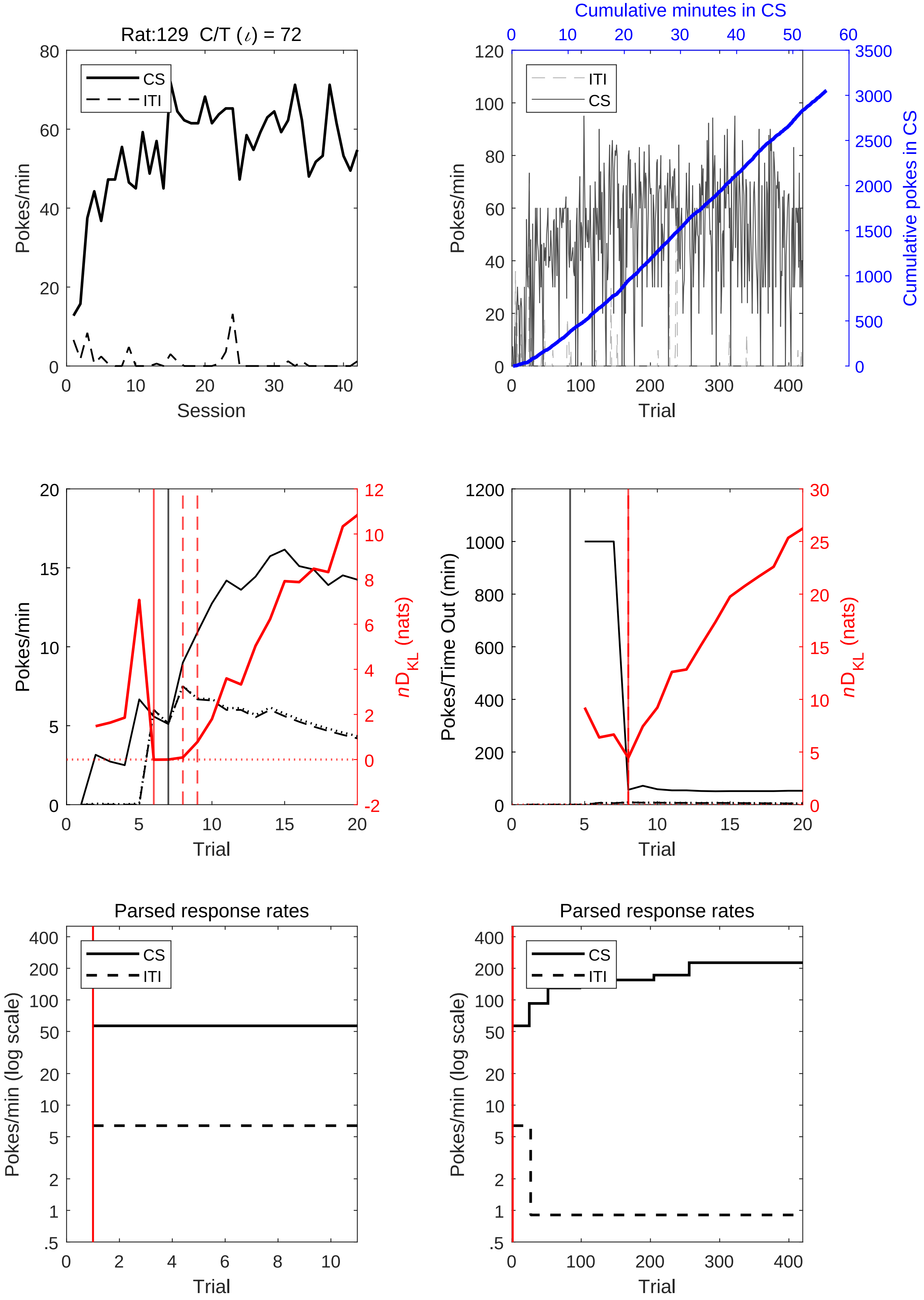
Rat:129 C/T (ι) = 72
Pokes/min
Session
CS
ITI
Cumulative minutes in CS
Cumulative pokes in CS
Trial
Pokes/Time Out (min)
nD_KL (nats)
Parsed response rates
Pokes/min (log scale)

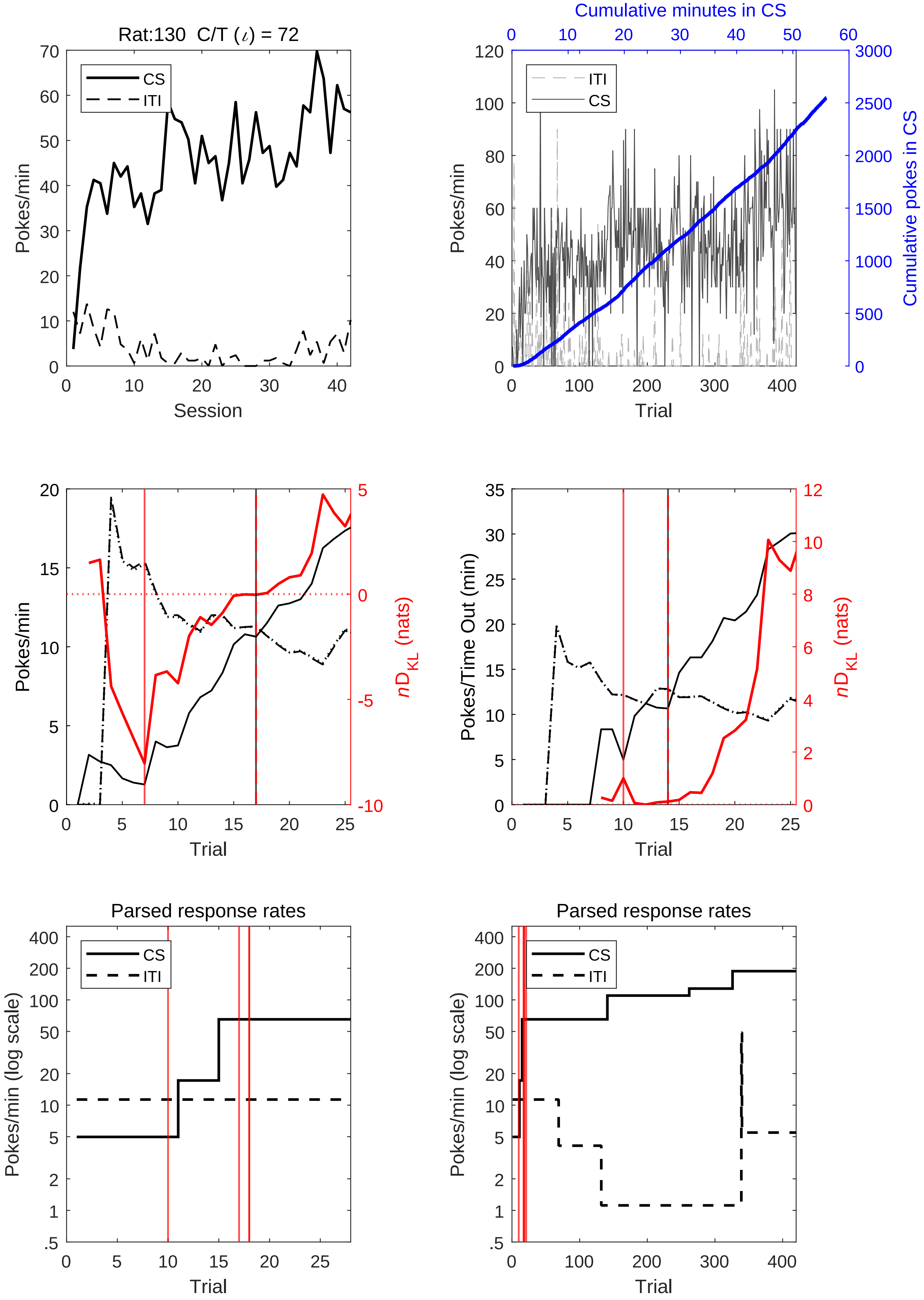

Rat:130 C/T (ι) = 72
CS
ITI
Pokes/min
Session
Cumulative minutes in CS
Cumulative pokes in CS
Trial
nD_KL (nats)
Pokes/Time Out (min)
Parsed response rates
Pokes/min (log scale)

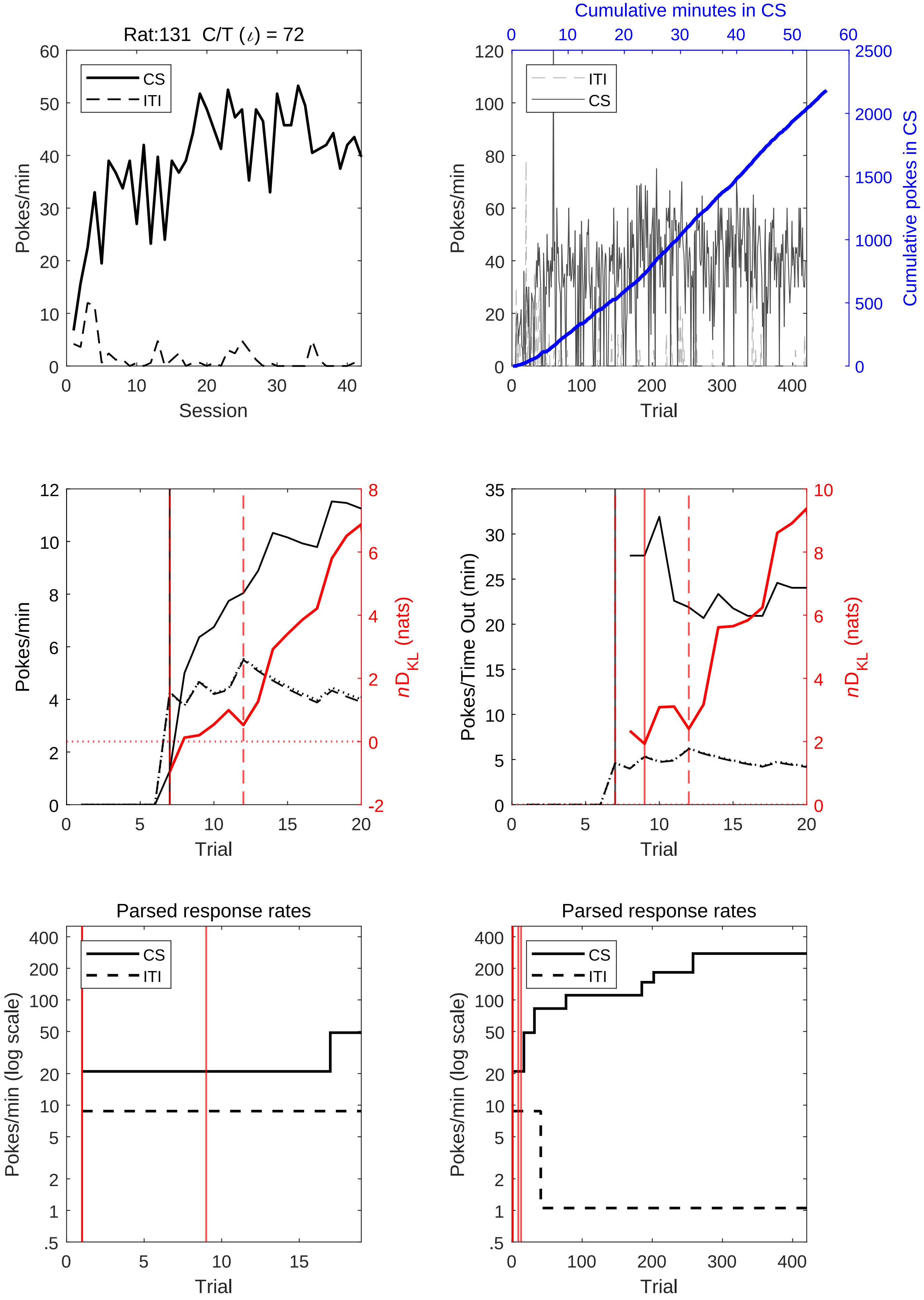
Rat:131 C/T ($\iota$) = 72
60
50
40
30
20
10
0
Pokes/min
CS
ITI
0
10
20
30
40
Session
Cumulative minutes in CS
0
10
20
30
40
50
60
120
100
80
60
40
20
0
Pokes/min
ITI
CS
2500
2000
1500
1000
500
0
Cumulative pokes in CS
0
100
200
300
400
Trial
12
10
8
6
4
2
0
Pokes/min
8
6
4
2
0
-2
$nD_{KL}$ (nats)
0
5
10
15
20
Trial
35
30
25
20
15
10
5
0
Pokes/Time Out (min)
10
8
6
4
2
0
$nD_{KL}$ (nats)
0
5
10
15
20
Trial
Parsed response rates
400
200
100
50
20
10
5
2
1
.5
Pokes/min (log scale)
CS
ITI
0
5
10
15
Trial
Parsed response rates
400
200
100
50
20
10
5
2
1
.5
Pokes/min (log scale)
CS
ITI
0
100
200
300
400
Trial

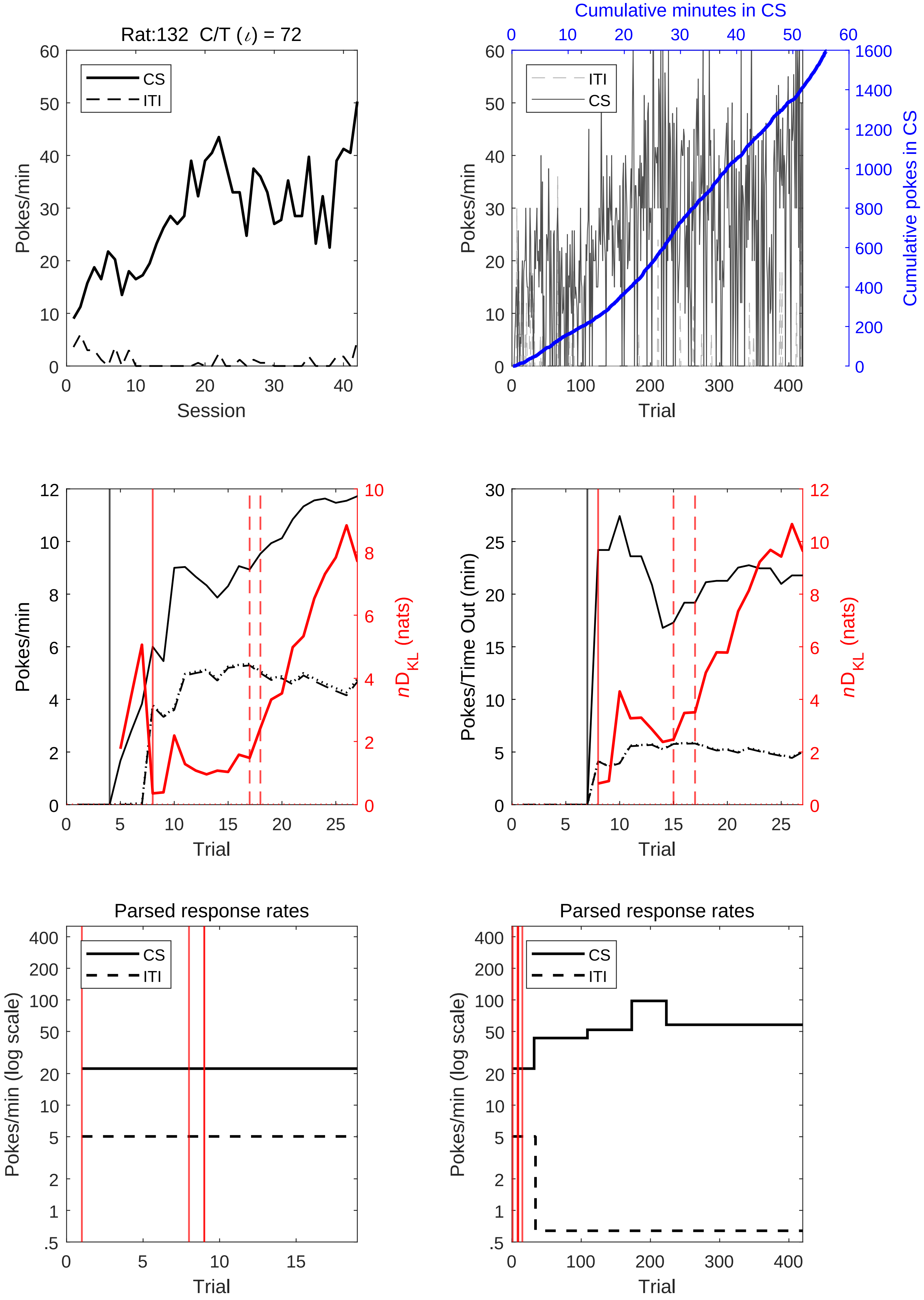

Rat:132 C/T (ι) = 72
CS
ITI
Pokes/min
Session
Cumulative minutes in CS
Cumulative pokes in CS
Trial
Pokes/Time Out (min)
$nD_{KL}$ (nats)
Parsed response rates
Pokes/min (log scale)

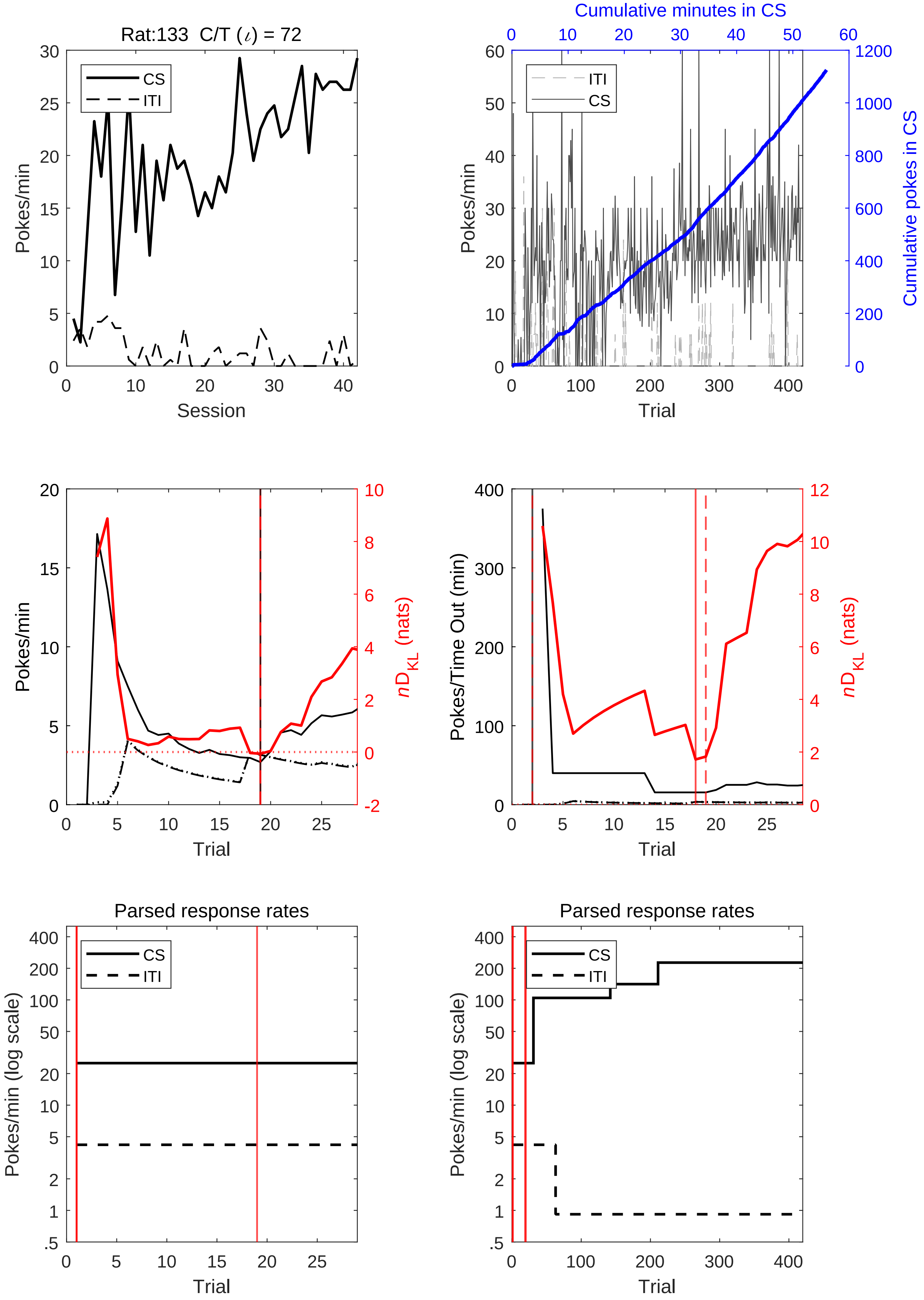
Rat:133 C/T ($\iota$) = 72
CS
ITI
Pokes/min
Session
Cumulative minutes in CS
Cumulative pokes in CS
Trial
$nD_{KL}$ (nats)
Pokes/Time Out (min)
Parsed response rates
Pokes/min (log scale)

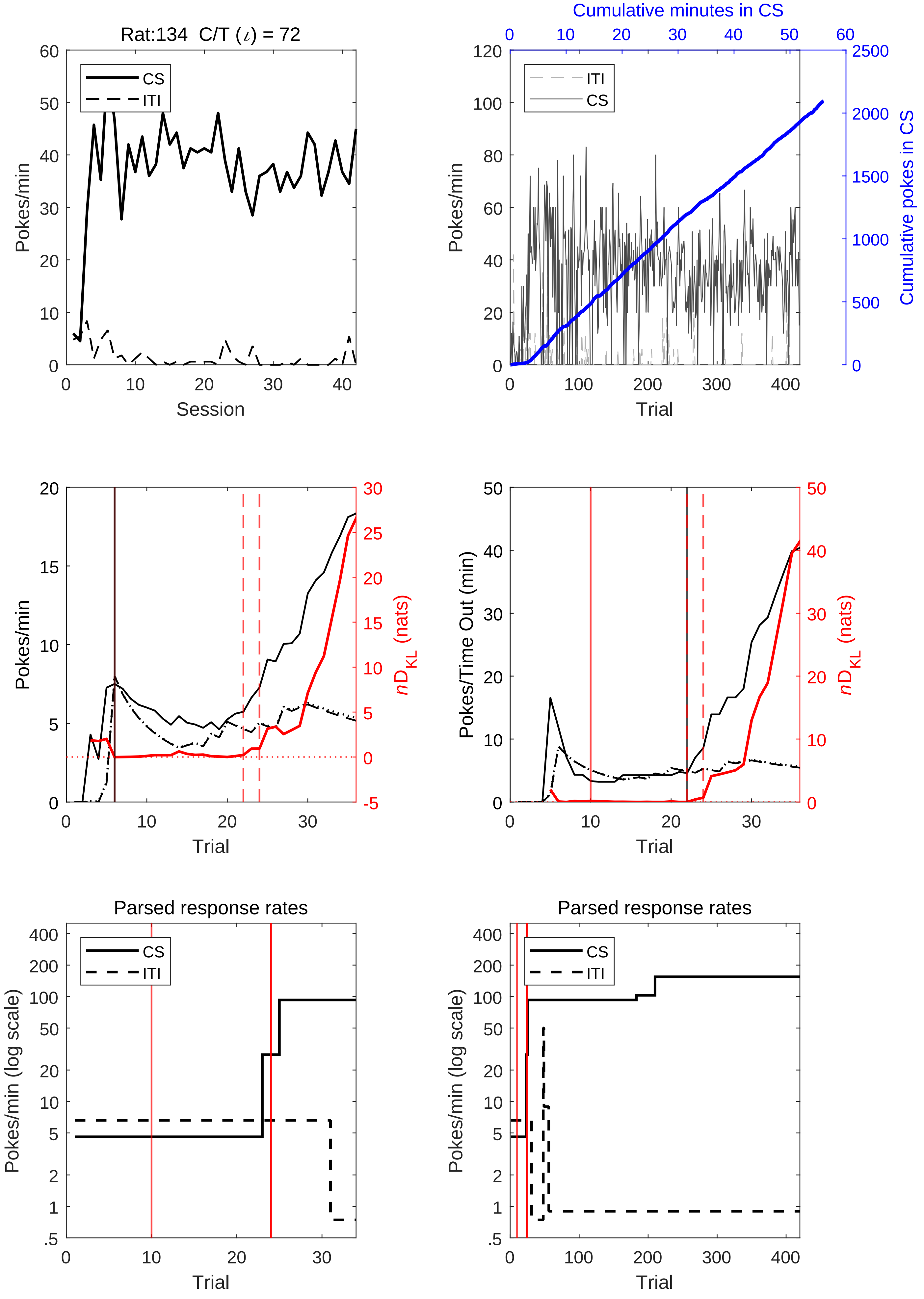
Rat:134 C/T (ι) = 72
CS
ITI
Pokes/min
Session
Cumulative minutes in CS
ITI
CS
Pokes/min
Cumulative pokes in CS
Trial
Pokes/min
$nD_{KL}$ (nats)
Trial
Pokes/Time Out (min)
$nD_{KL}$ (nats)
Trial
Parsed response rates
CS
ITI
Pokes/min (log scale)
Trial
Parsed response rates
CS
ITI
Pokes/min (log scale)
Trial

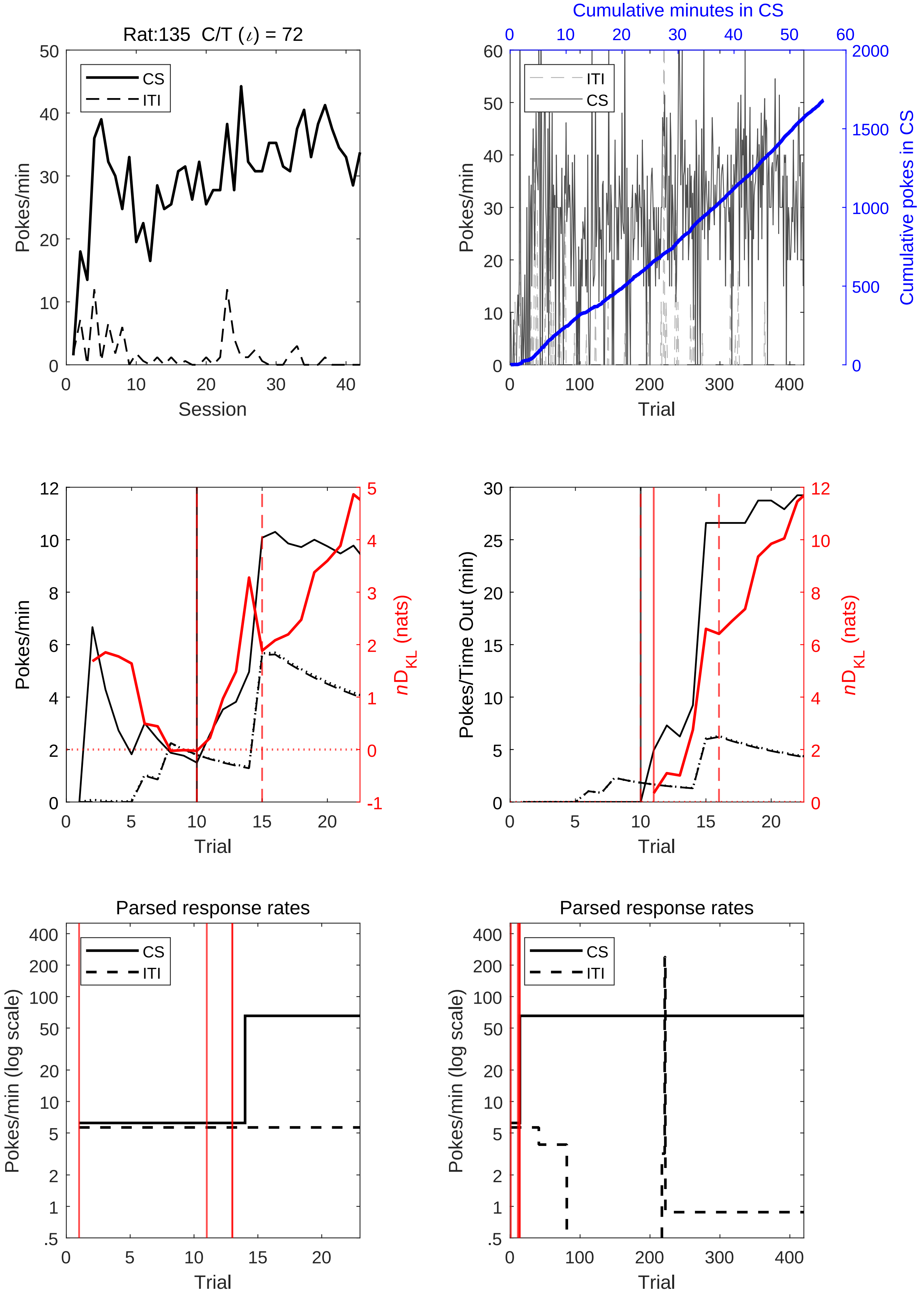
Rat:135 C/T (ι) = 72
CS
ITI
Pokes/min
Session
Cumulative minutes in CS
Cumulative pokes in CS
Trial
nD_KL (nats)
Pokes/Time Out (min)
Parsed response rates
Pokes/min (log scale)

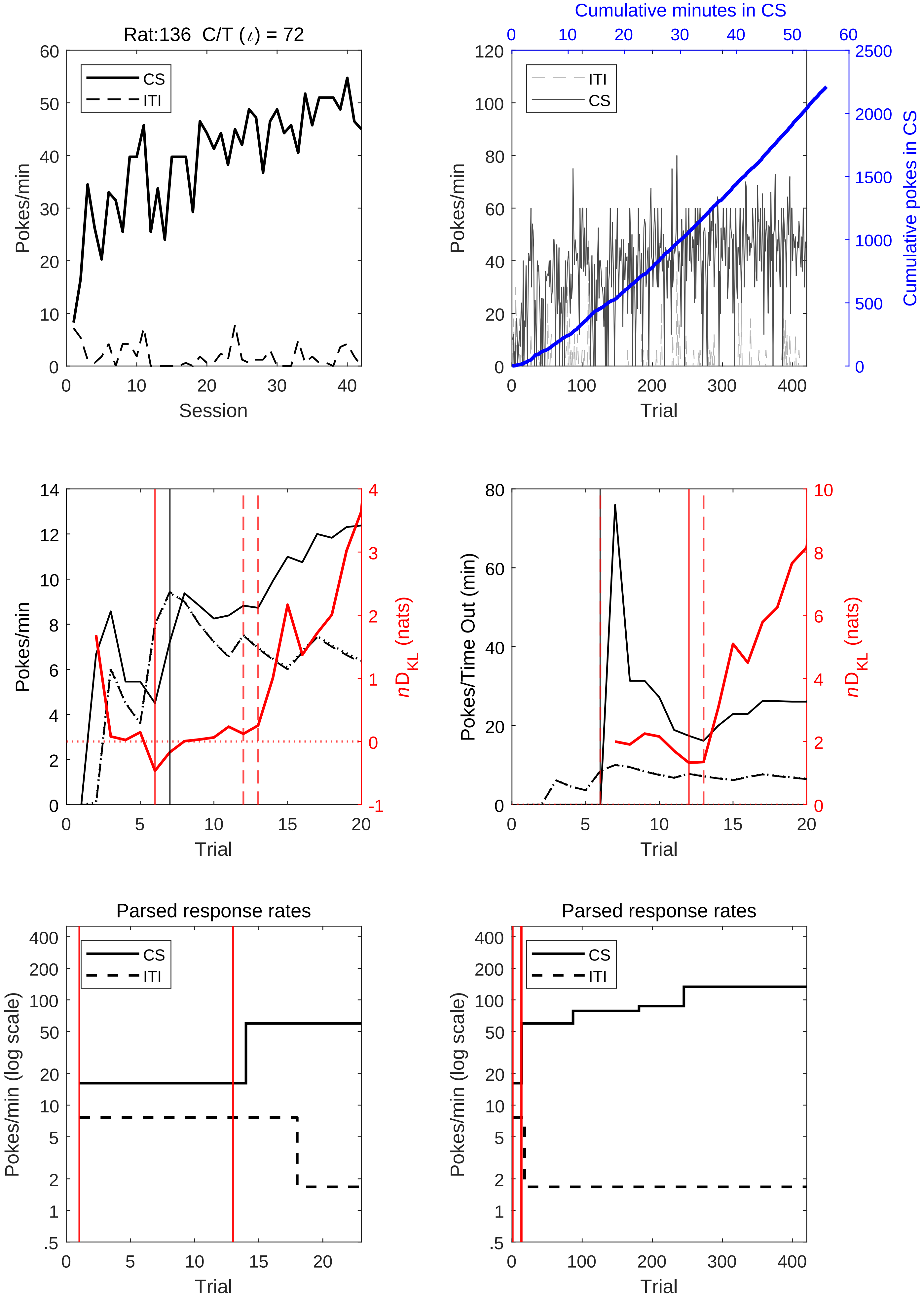

Rat:136 C/T (ι) = 72
CS
ITI
Pokes/min
Session
Cumulative minutes in CS
Cumulative pokes in CS
Trial
nD_KL (nats)
Pokes/Time Out (min)
Parsed response rates
Pokes/min (log scale)

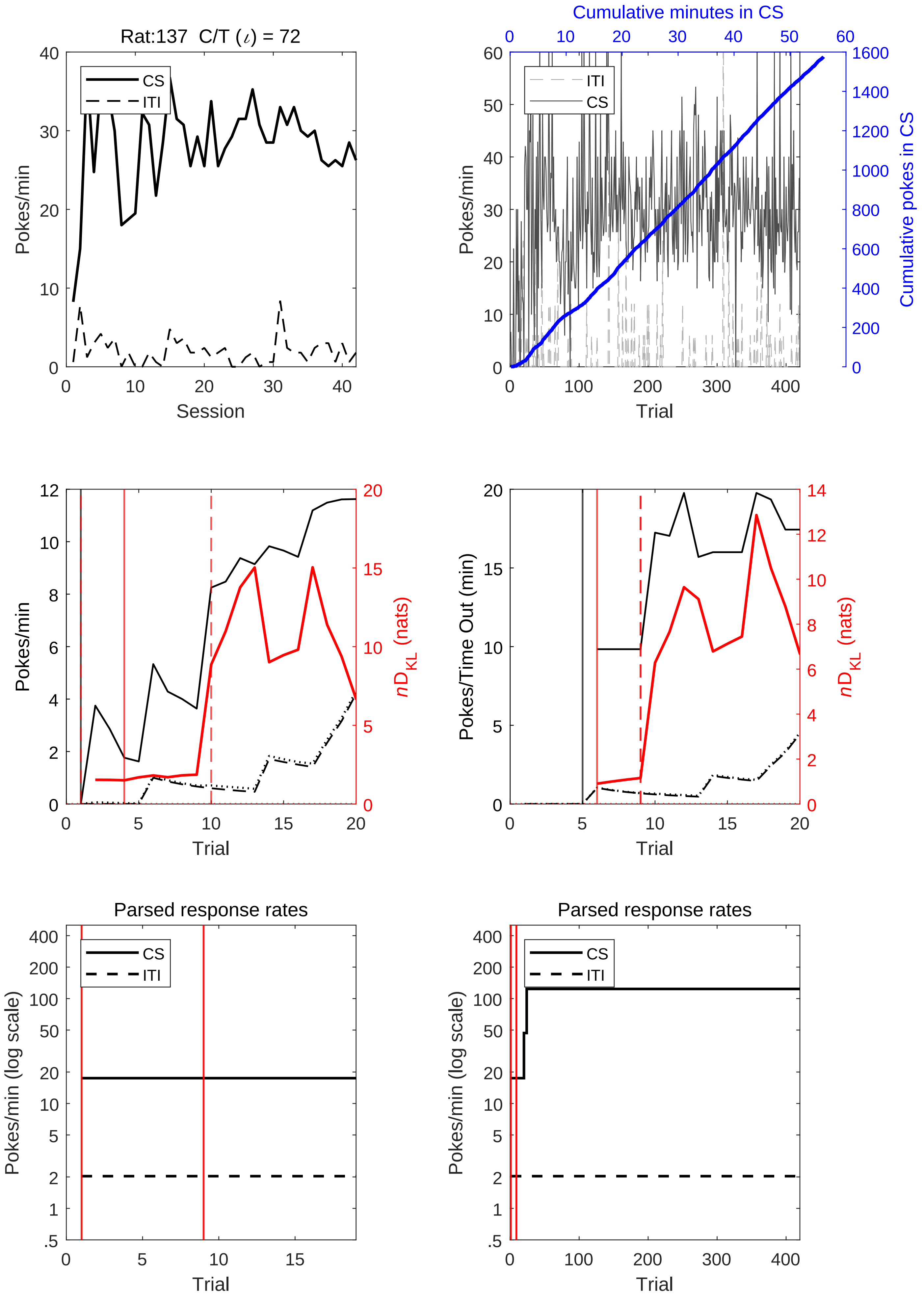
Rat:137 C/T (ι) = 72
CS
ITI
Pokes/min
Session
Cumulative minutes in CS
Cumulative pokes in CS
Trial
Pokes/Time Out (min)
nD_KL (nats)
Parsed response rates
Pokes/min (log scale)

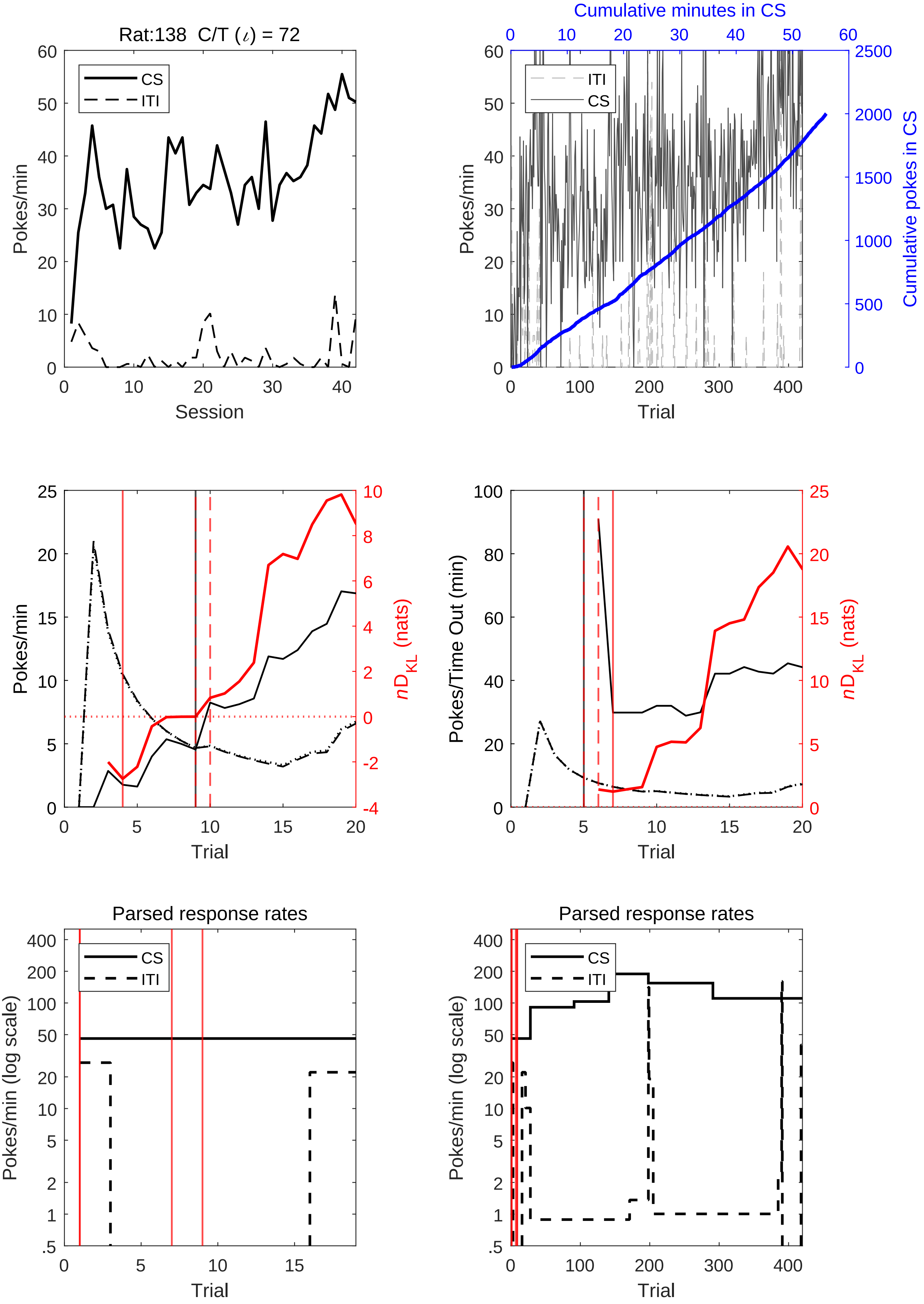
Rat:138 C/T (ι) = 72
Pokes/min
Session
CS
ITI
Cumulative minutes in CS
Cumulative pokes in CS
Trial
Pokes/Time Out (min)
nD_KL (nats)
Parsed response rates
Pokes/min (log scale)

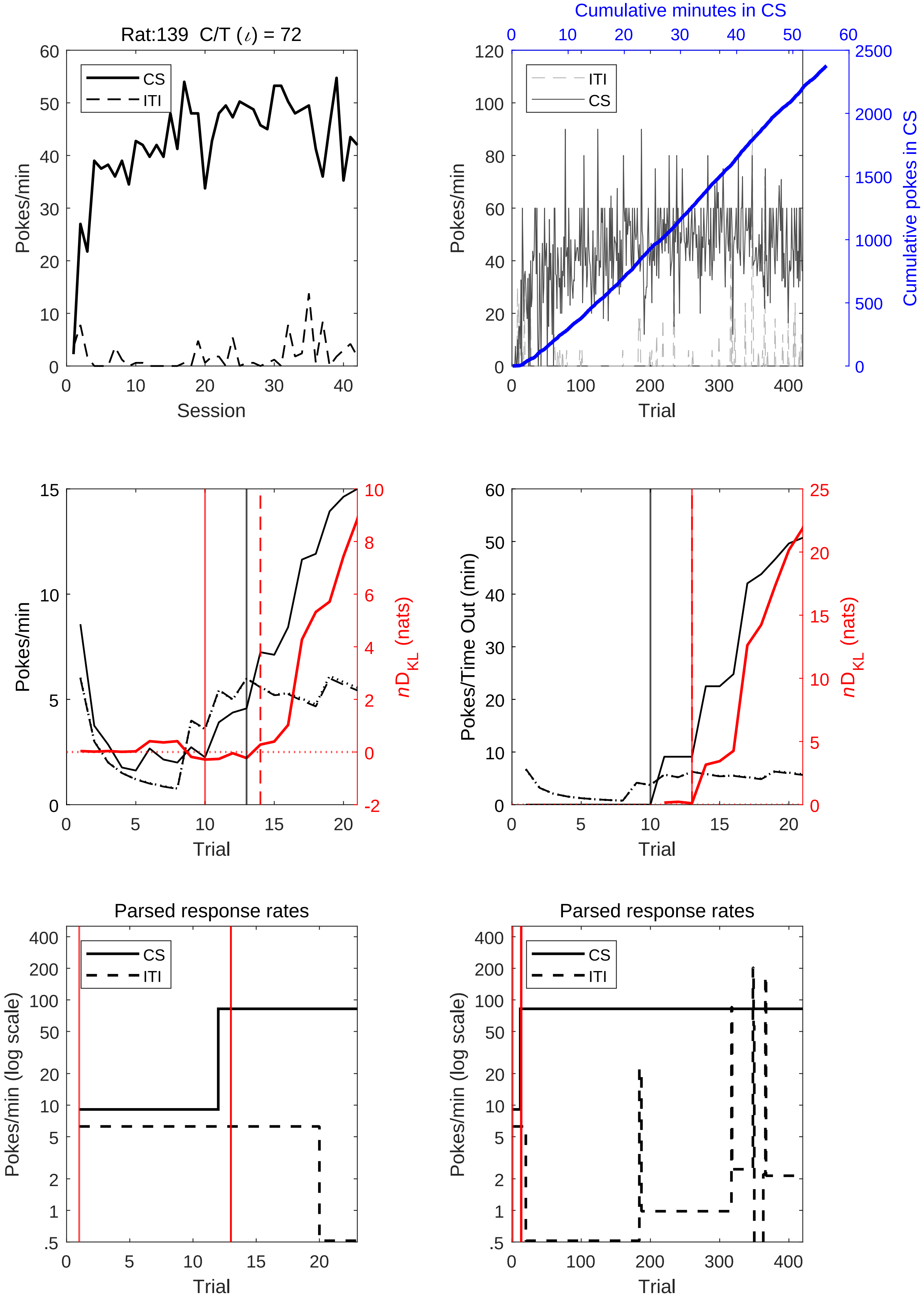
Rat:139 C/T (ι) = 72
CS
ITI
Pokes/min
Session
Cumulative minutes in CS
Cumulative pokes in CS
Trial
nD_KL (nats)
Pokes/Time Out (min)
Parsed response rates
Pokes/min (log scale)

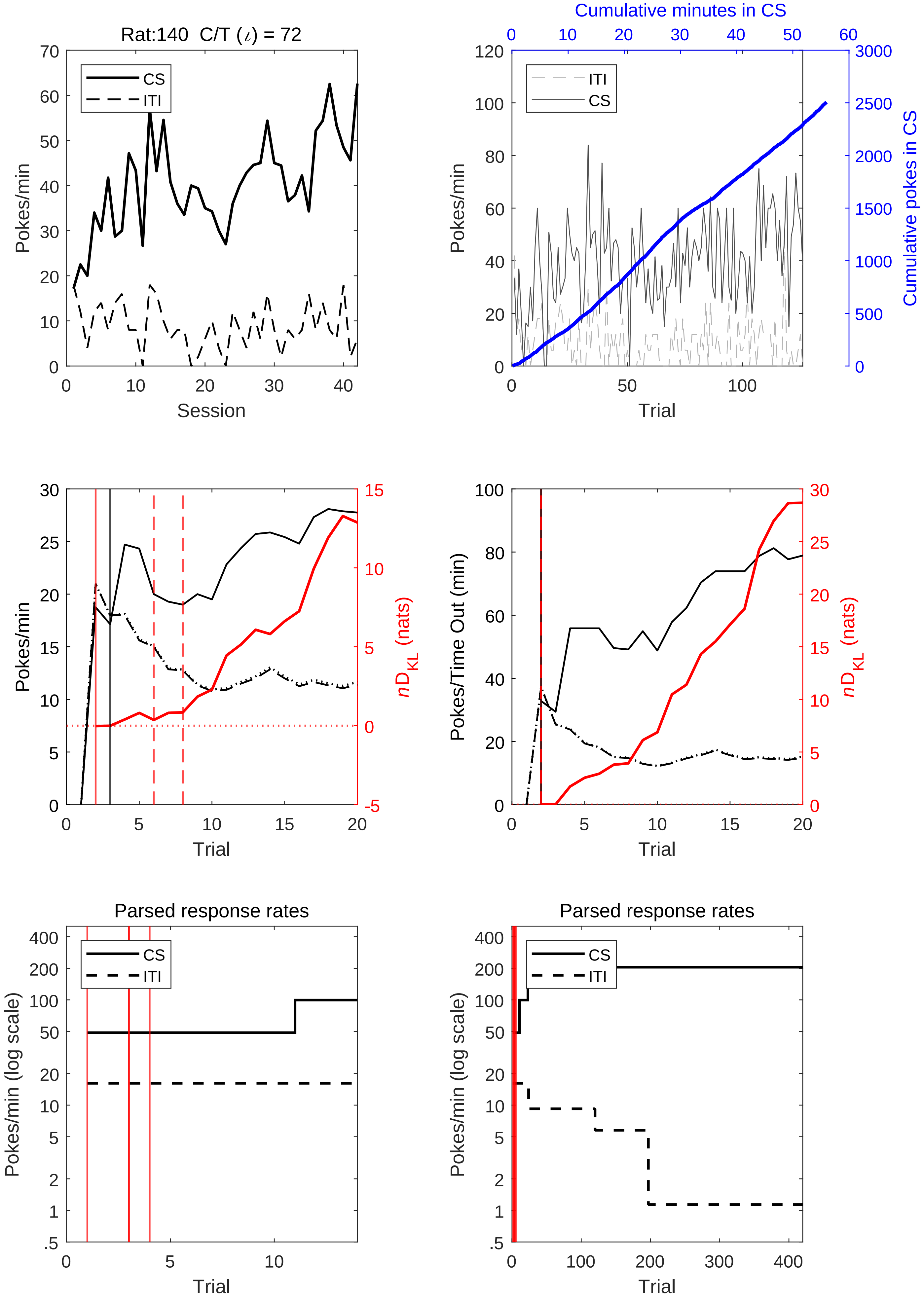
Rat:140 C/T (ι) = 72
CS
ITI
Pokes/min
Session
Cumulative minutes in CS
Cumulative pokes in CS
Trial
Pokes/Time Out (min)
nD_KL (nats)
Parsed response rates
Pokes/min (log scale)

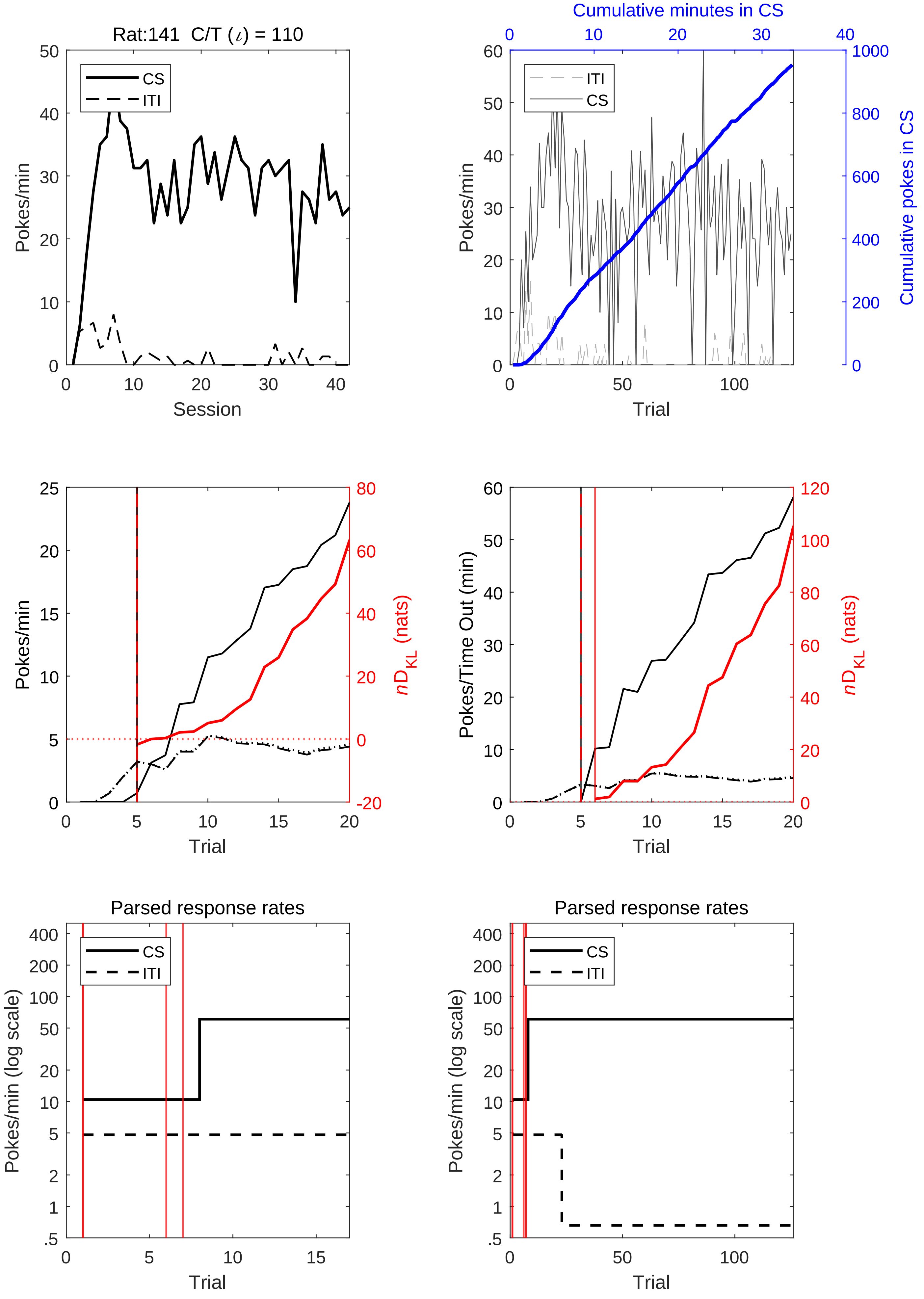
Rat:141 C/T (ι) = 110
Pokes/min
Session
CS
ITI
Cumulative minutes in CS
Pokes/min
Cumulative pokes in CS
Trial
Pokes/min
$nD_{KL}$ (nats)
Trial
Pokes/Time Out (min)
$nD_{KL}$ (nats)
Trial
Parsed response rates
Pokes/min (log scale)
Trial
Parsed response rates
Pokes/min (log scale)
Trial

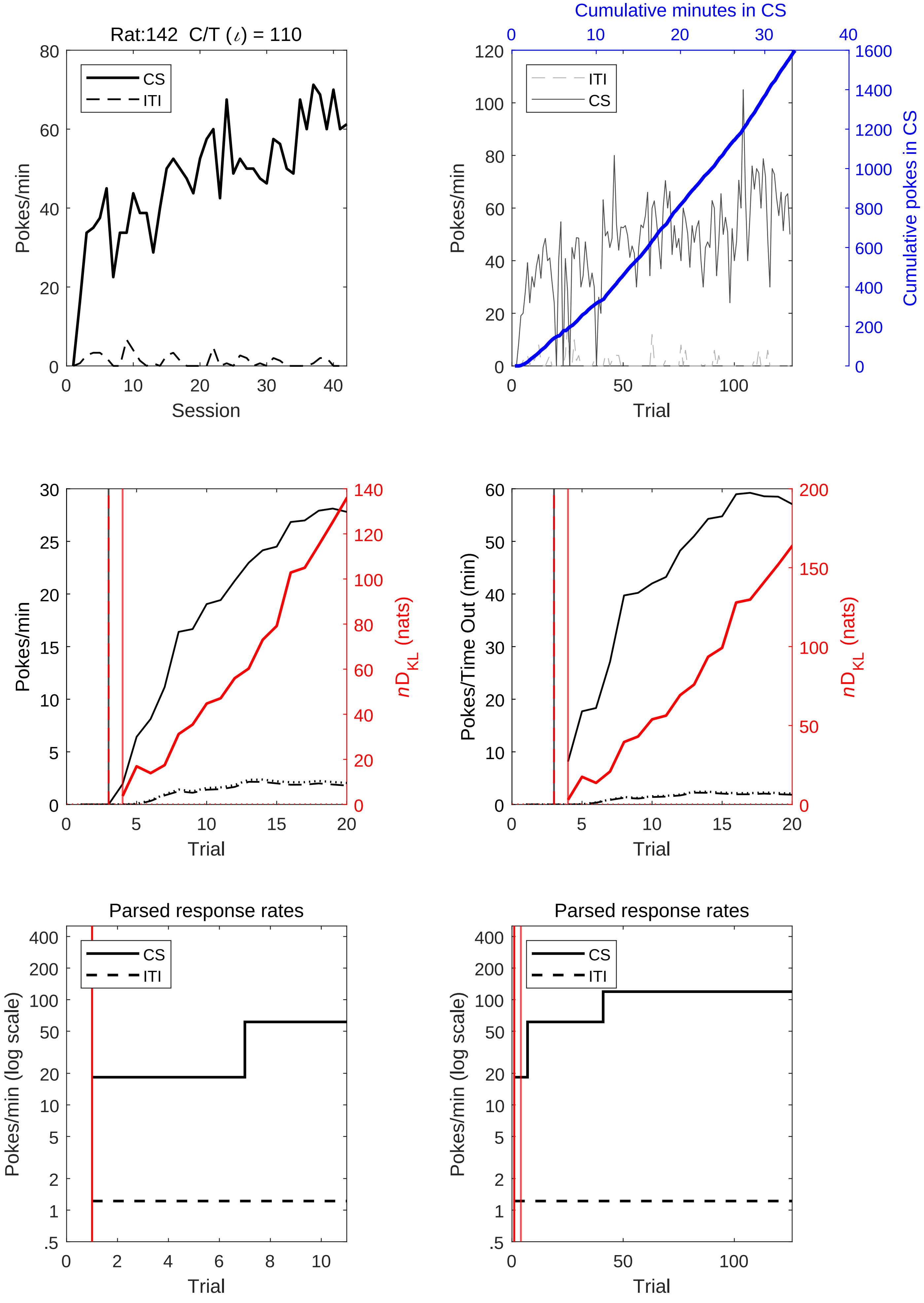

Rat:142 C/T ($\iota$) = 110
CS
ITI
Pokes/min
Session
Cumulative minutes in CS
Cumulative pokes in CS
Trial
$nD_{KL}$ (nats)
Pokes/Time Out (min)
Parsed response rates
Pokes/min (log scale)

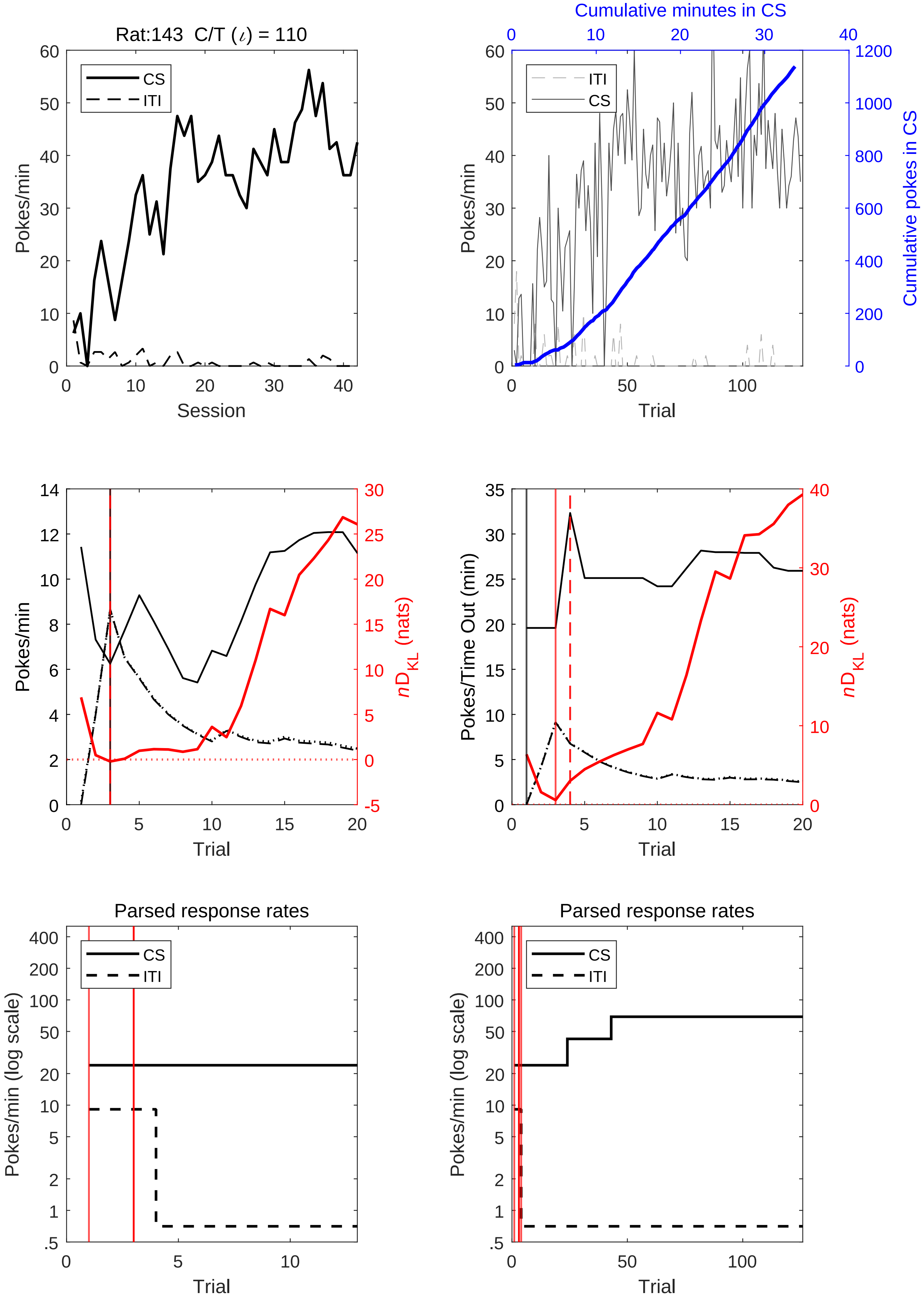

Rat:143 C/T (ι) = 110
CS
ITI
Pokes/min
Session
Cumulative minutes in CS
Cumulative pokes in CS
Trial
Pokes/Time Out (min)
nD_KL (nats)
Parsed response rates
Pokes/min (log scale)

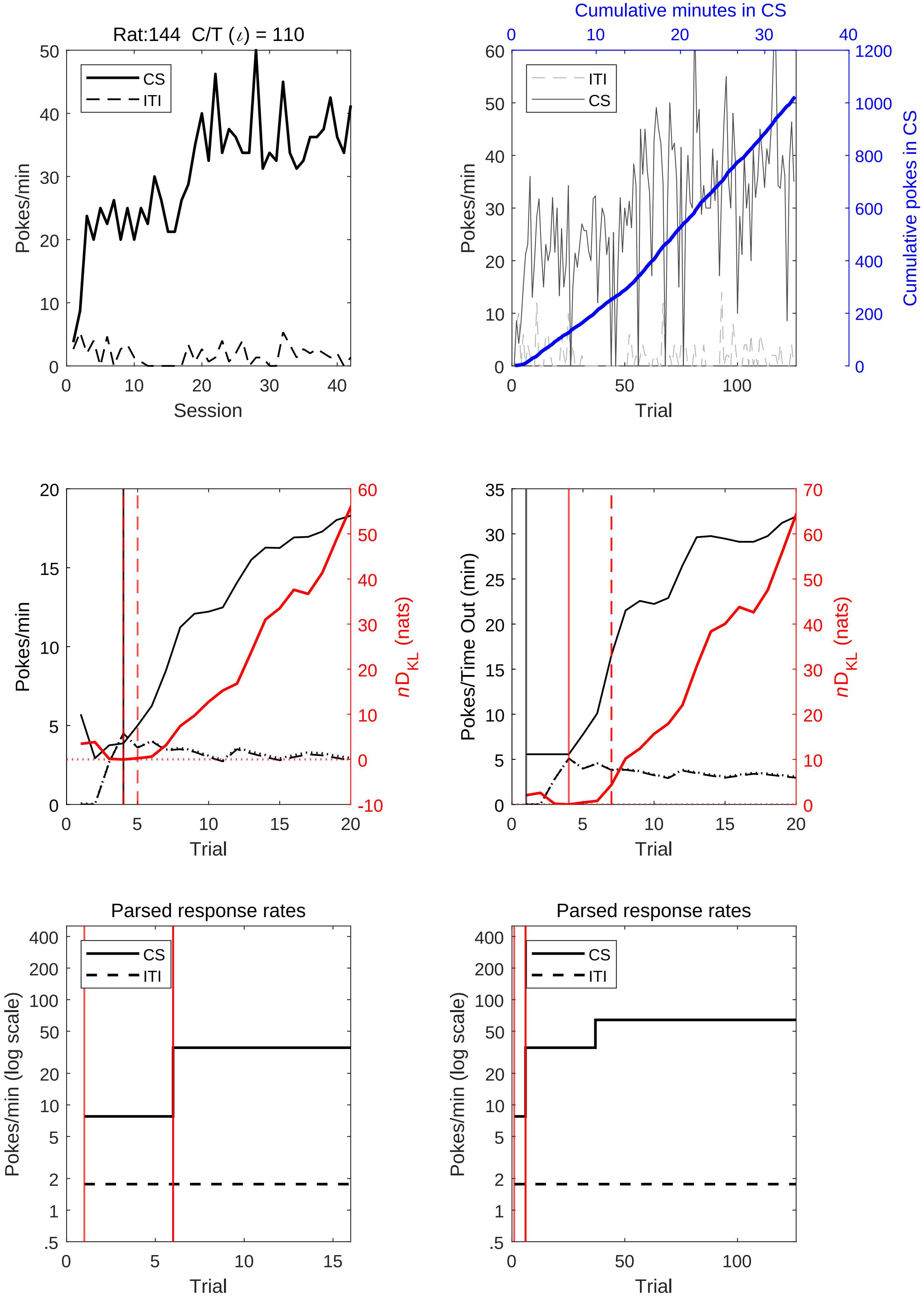
Rat:144 C/T (ι) = 110
CS
ITI
Pokes/min
Session
Cumulative minutes in CS
Cumulative pokes in CS
Trial
Pokes/Time Out (min)
nD_KL (nats)
Parsed response rates
Pokes/min (log scale)

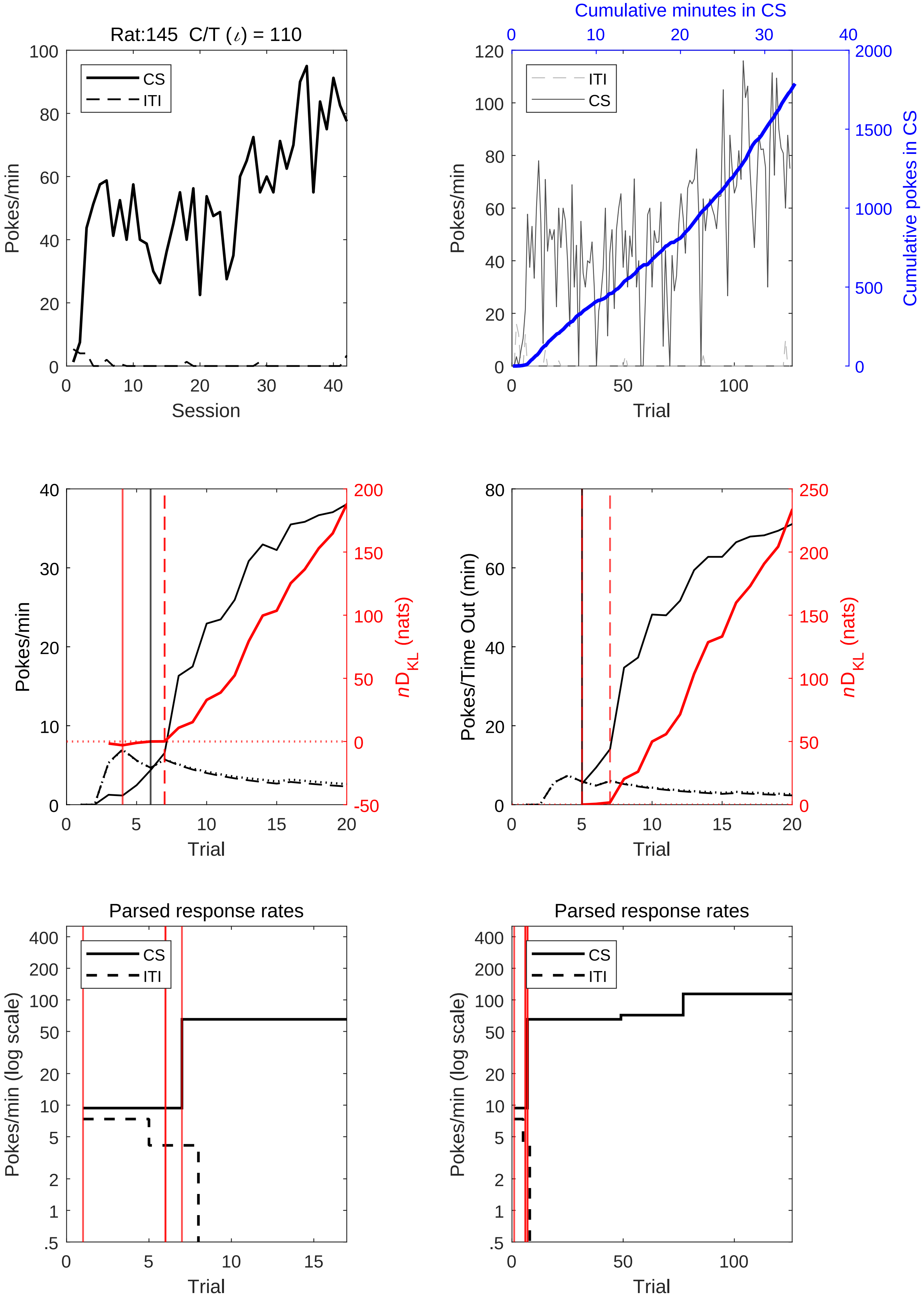

Rat:145 C/T (ι) = 110
CS
ITI
Pokes/min
Session
Cumulative minutes in CS
Cumulative pokes in CS
Trial
Pokes/Time Out (min)
nD_KL (nats)
Parsed response rates
Pokes/min (log scale)

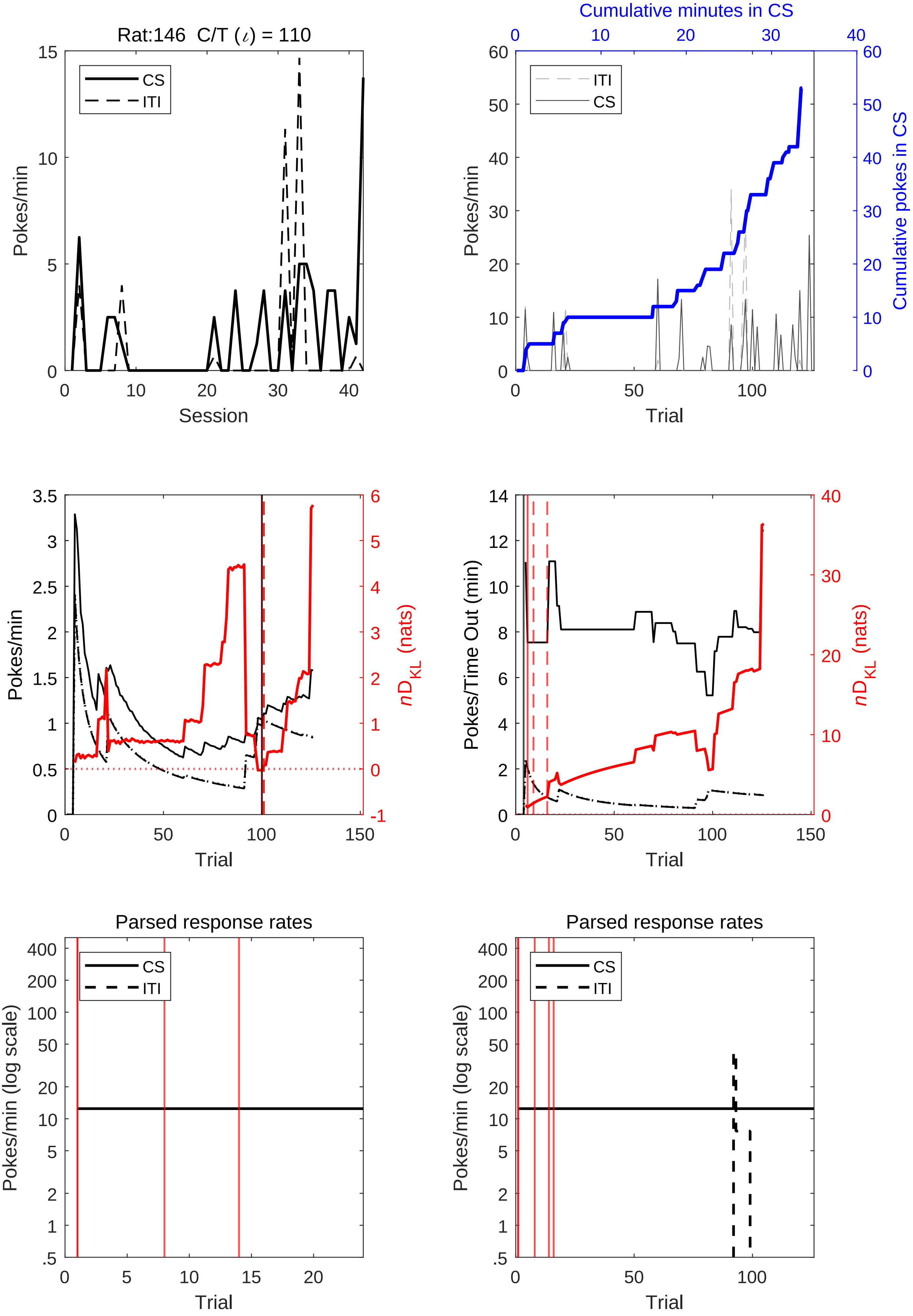
Rat:146 C/T (ι) = 110
CS
ITI
Pokes/min
Session
Cumulative minutes in CS
Cumulative pokes in CS
Trial
nD_KL (nats)
Pokes/Time Out (min)
Parsed response rates
Pokes/min (log scale)

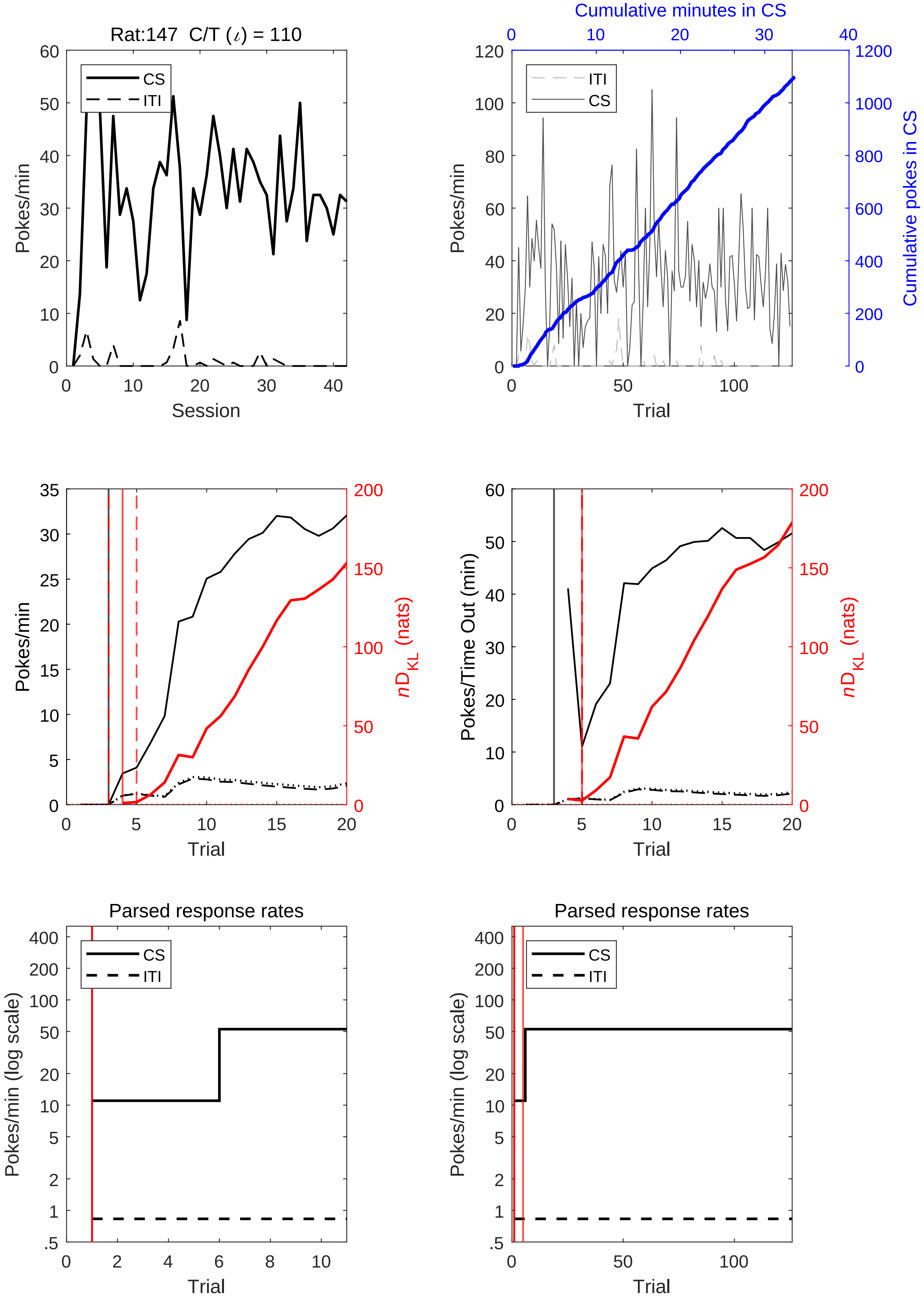

Rat:147 C/T (ι) = 110
CS
ITI
Pokes/min
Session
Cumulative minutes in CS
Cumulative pokes in CS
Trial
Pokes/Time Out (min)
nD_KL (nats)
Parsed response rates
Pokes/min (log scale)

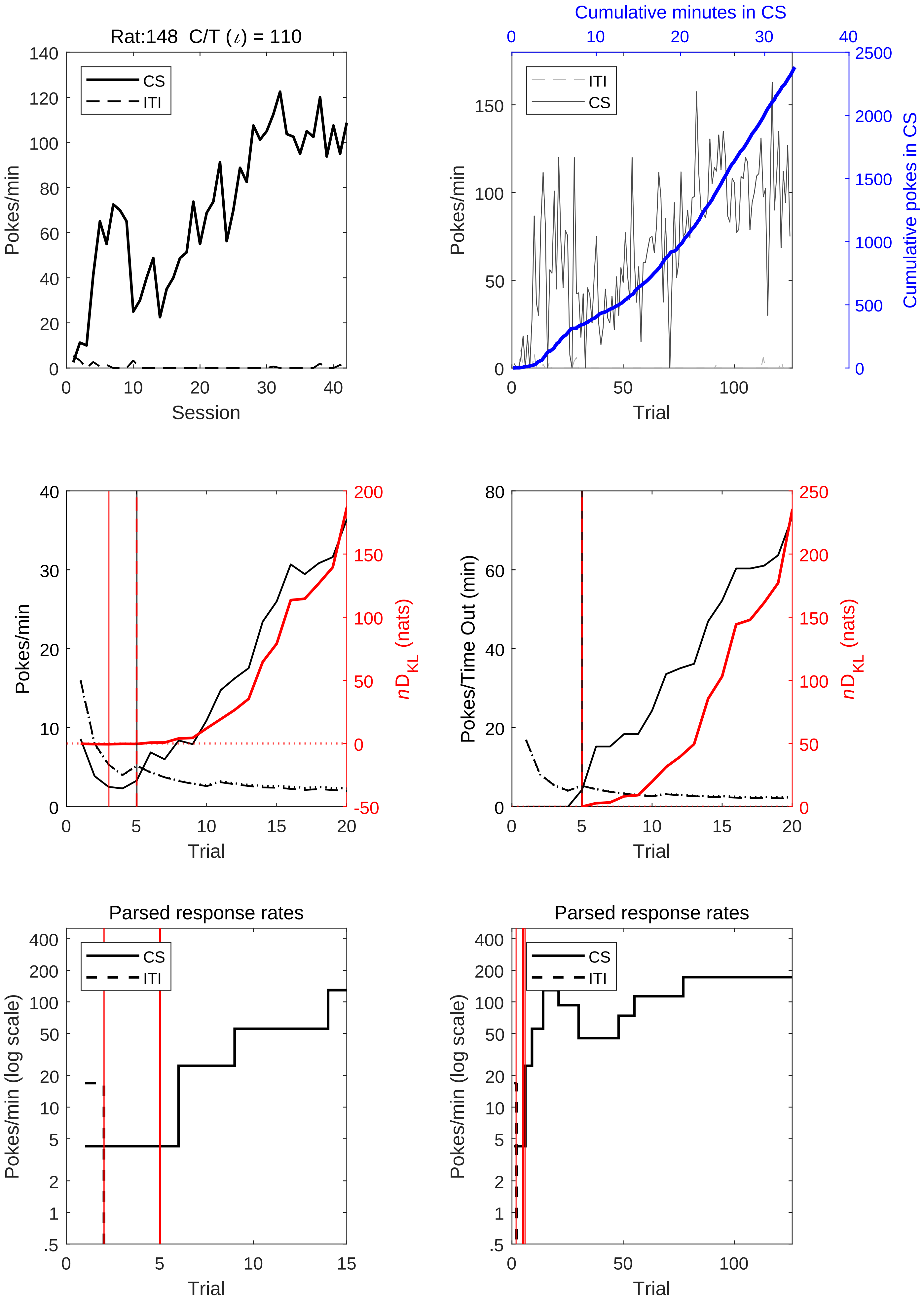
Rat:148 C/T (ι) = 110
CS
ITI
Pokes/min
Session
Cumulative minutes in CS
Cumulative pokes in CS
Trial
Pokes/min
nD_KL (nats)
Pokes/Time Out (min)
Parsed response rates
Pokes/min (log scale)

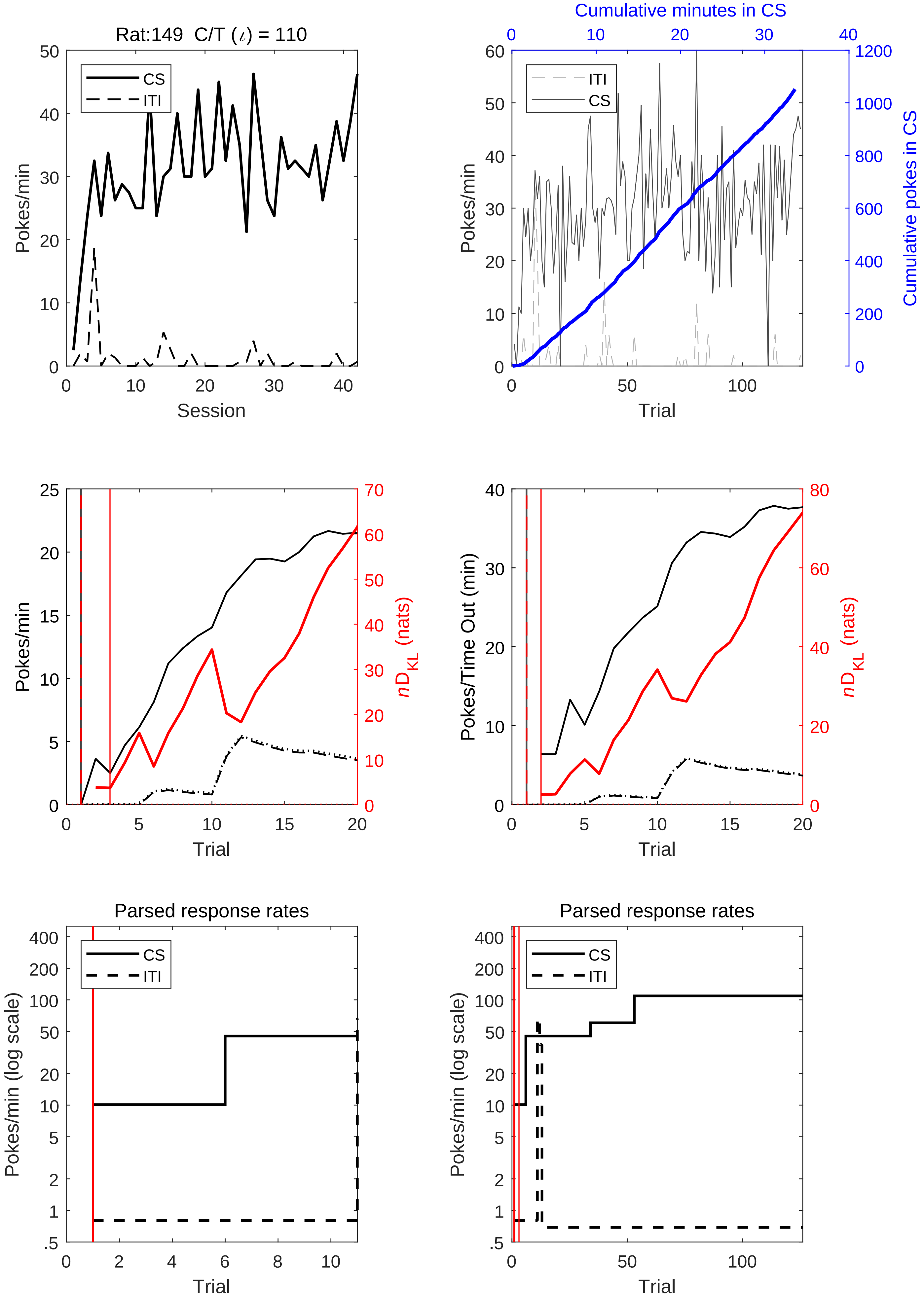
Rat:149 C/T (ι) = 110
CS
ITI
Pokes/min
Session
Cumulative minutes in CS
Cumulative pokes in CS
Trial
Pokes/Time Out (min)
$nD_{KL}$ (nats)
Parsed response rates
Pokes/min (log scale)

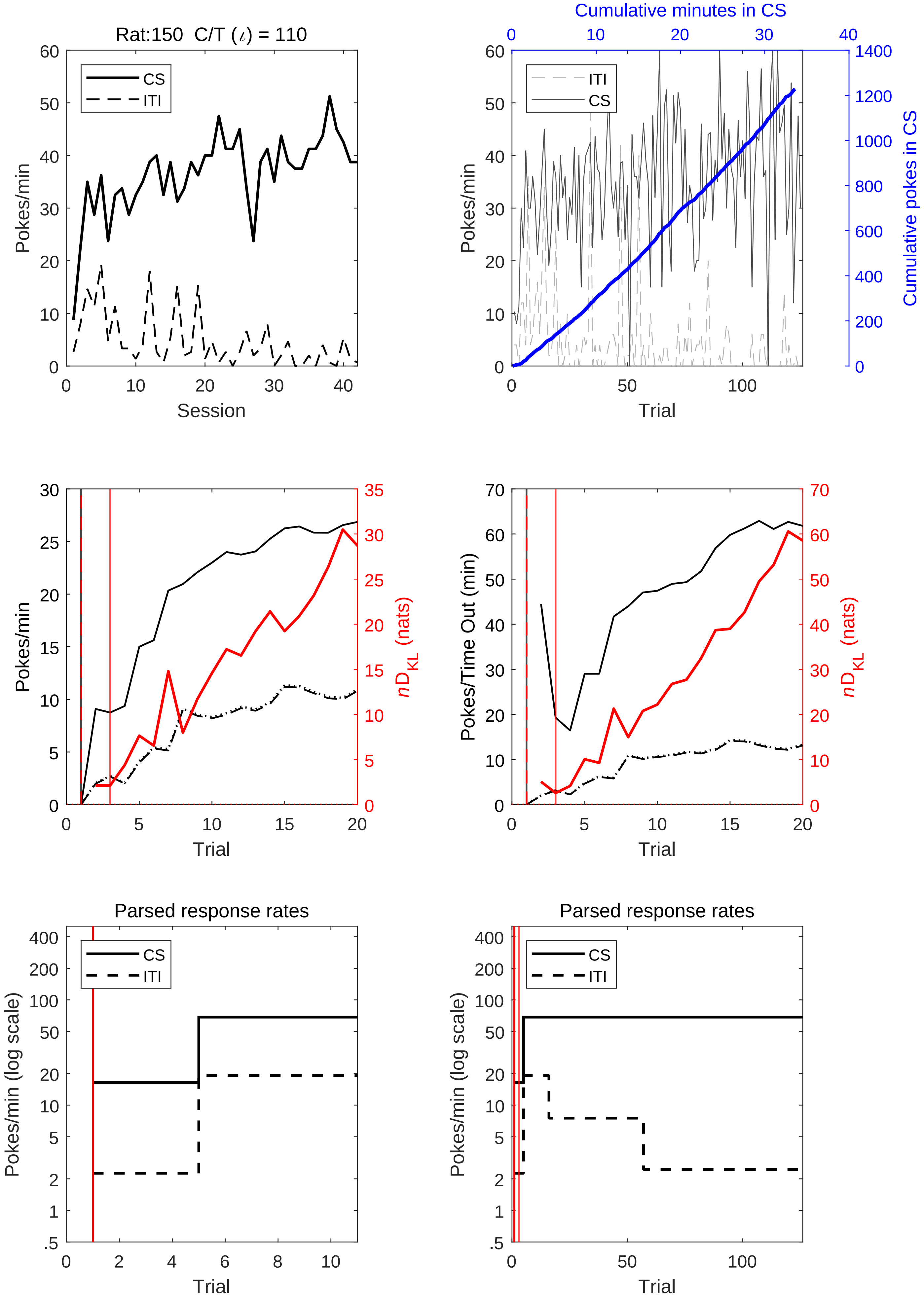

Rat:150 C/T (ι) = 110
CS
ITI
Pokes/min
Session
Cumulative minutes in CS
ITI
CS
Pokes/min
Trial
Cumulative pokes in CS
Pokes/min
Trial
nD_KL (nats)
Pokes/Time Out (min)
Trial
nD_KL (nats)
Parsed response rates
CS
ITI
Pokes/min (log scale)
Trial
Parsed response rates
CS
ITI
Pokes/min (log scale)
Trial

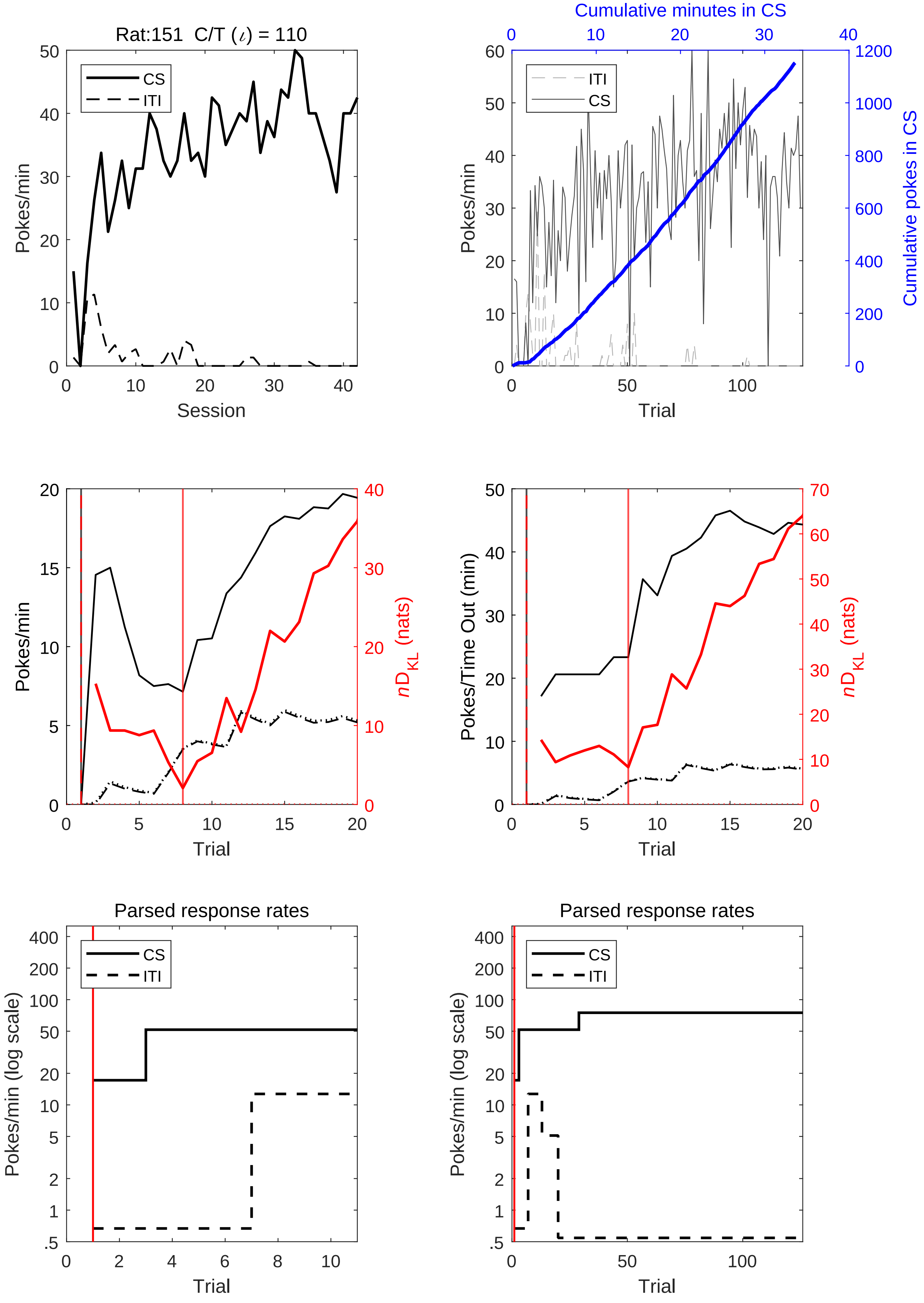
Rat:151 C/T ($\iota$) = 110
CS
ITI
Pokes/min
Session
Cumulative minutes in CS
ITI
CS
Pokes/min
Cumulative pokes in CS
Trial
Pokes/min
$n$D$_{KL}$ (nats)
Trial
Pokes/Time Out (min)
$n$D$_{KL}$ (nats)
Trial
Parsed response rates
CS
ITI
Pokes/min (log scale)
Trial
Parsed response rates
CS
ITI
Pokes/min (log scale)
Trial

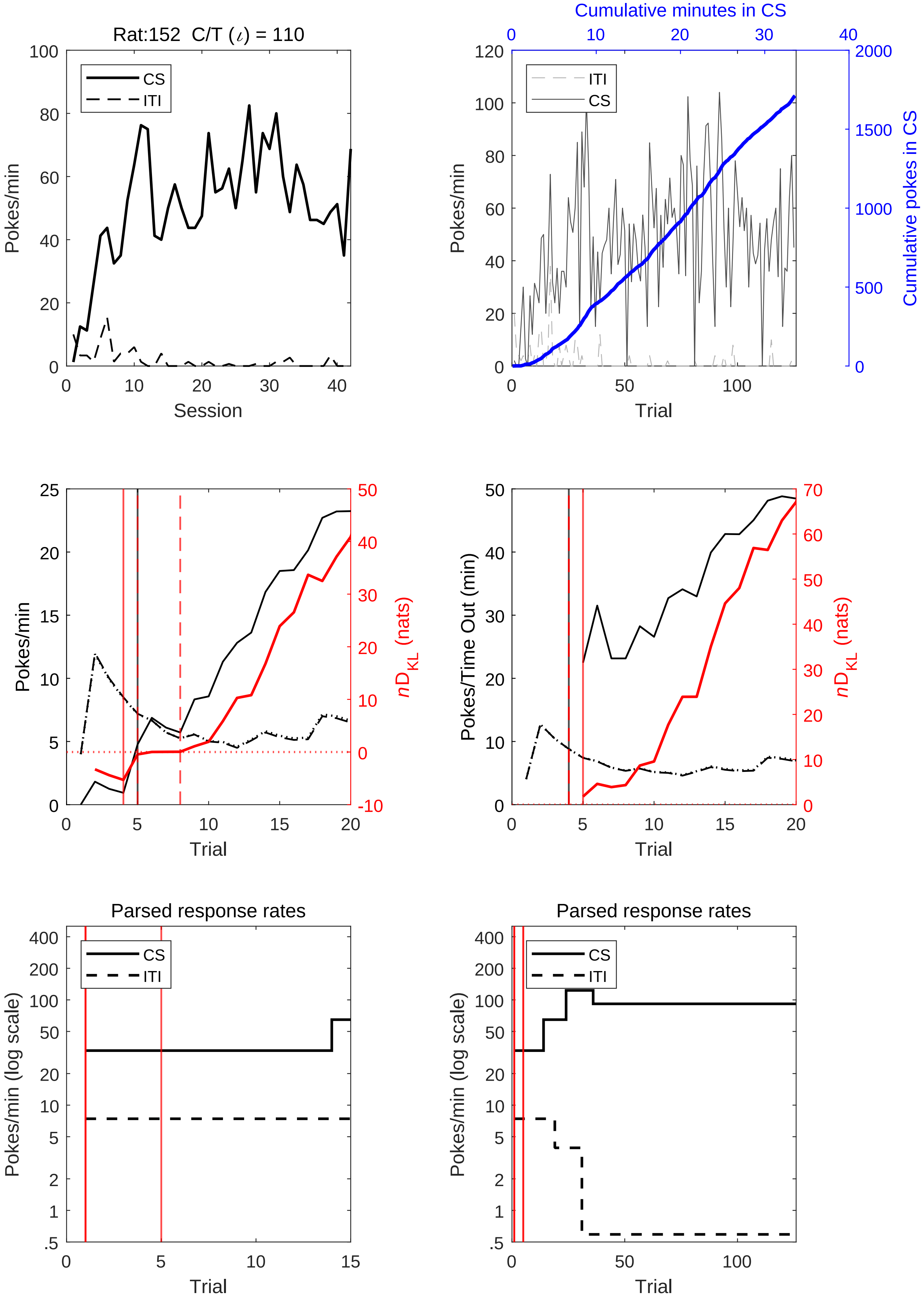

Rat:152 C/T (ι) = 110
CS
ITI
Pokes/min
Session
Cumulative minutes in CS
Cumulative pokes in CS
Trial
Pokes/Time Out (min)
nD_KL (nats)
Parsed response rates
Pokes/min (log scale)

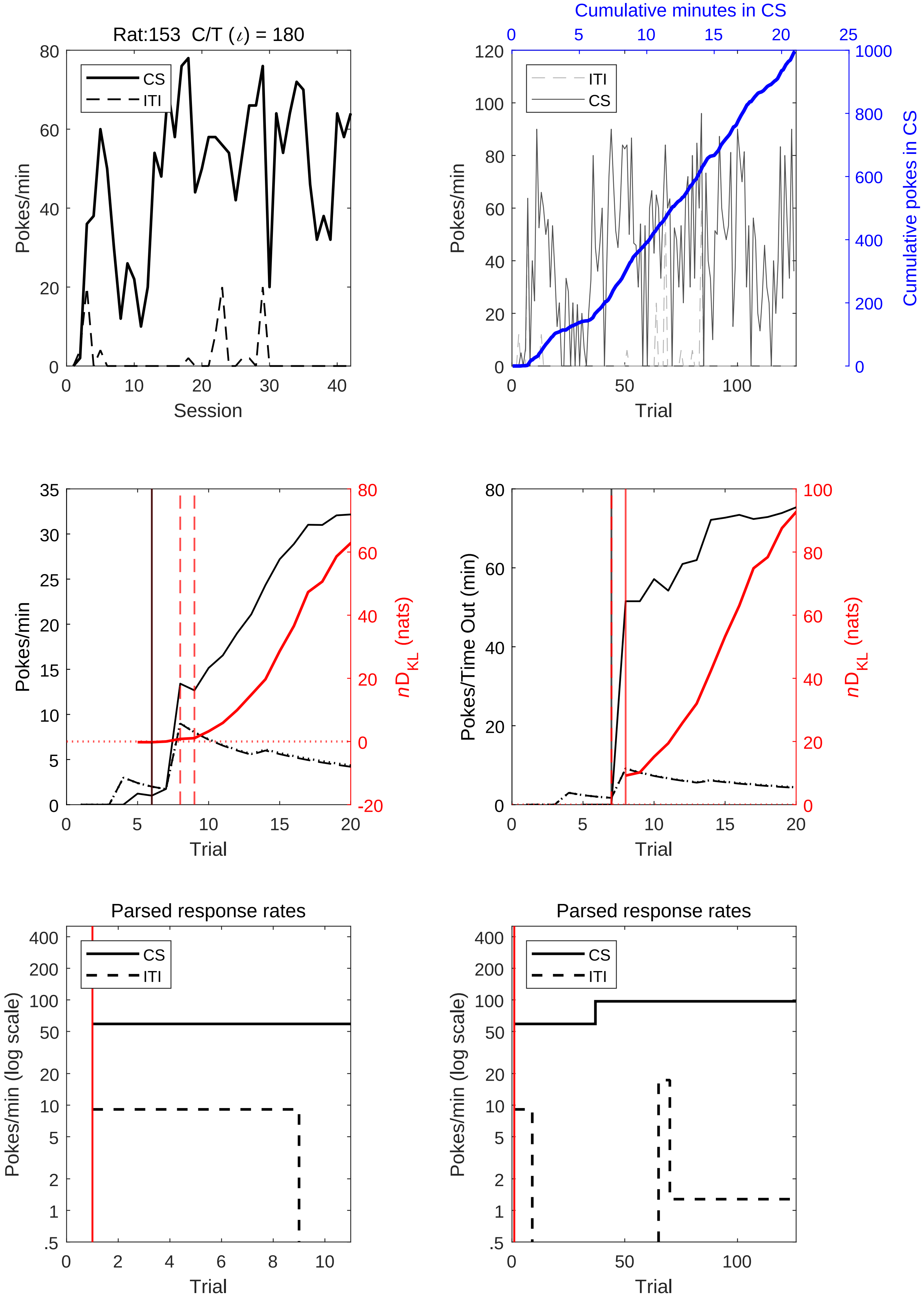

Rat:153 C/T (ι) = 180
CS
ITI
Pokes/min
Session
Cumulative minutes in CS
Cumulative pokes in CS
Trial
$nD_{KL}$ (nats)
Pokes/Time Out (min)
Parsed response rates
Pokes/min (log scale)

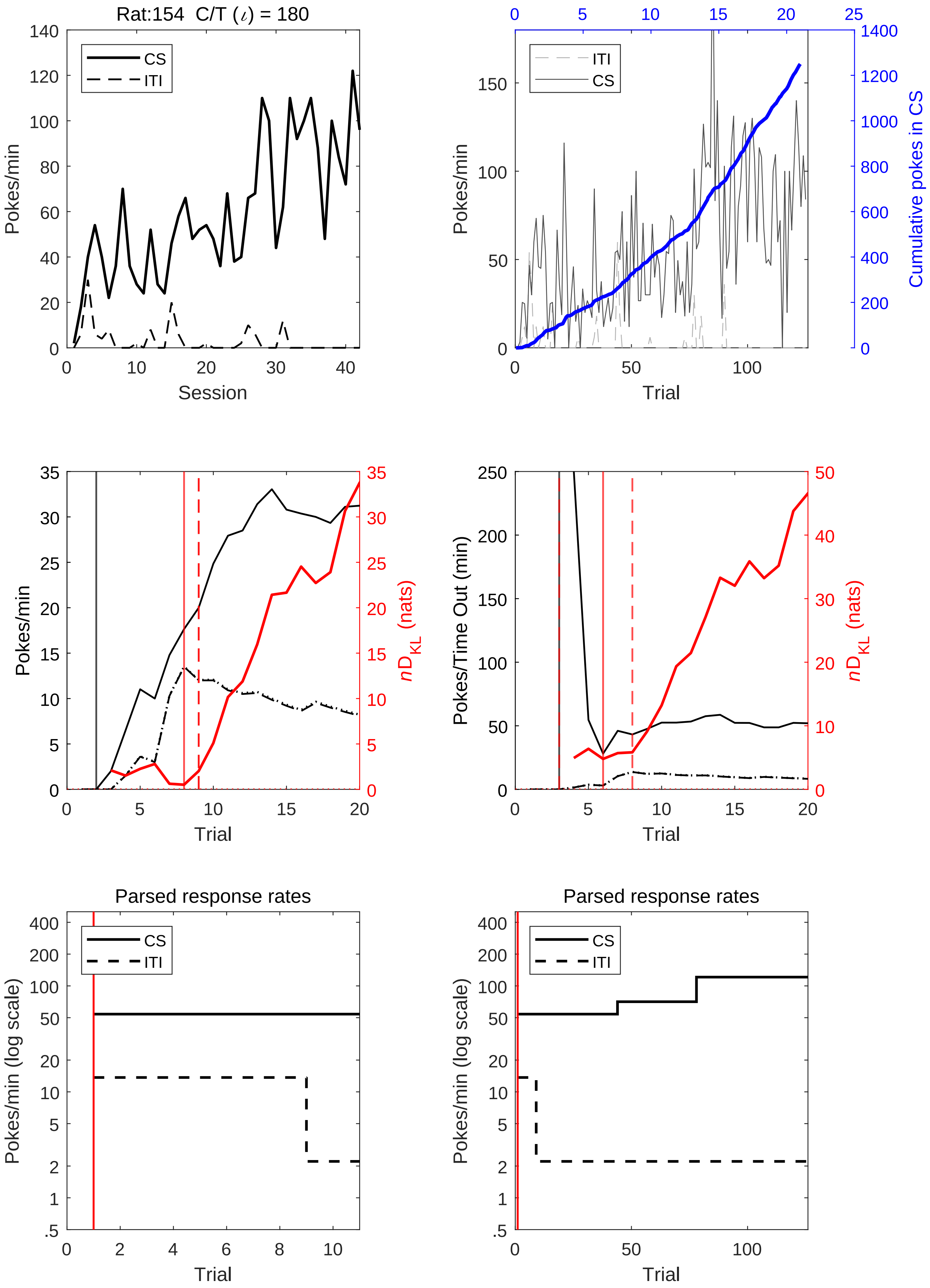
Rat:154 C/T (ι) = 180
CS
ITI
Pokes/min
Session
Cumulative minutes in CS
Cumulative pokes in CS
Trial
Pokes/Time Out (min)
nD_KL (nats)
Parsed response rates
Pokes/min (log scale)

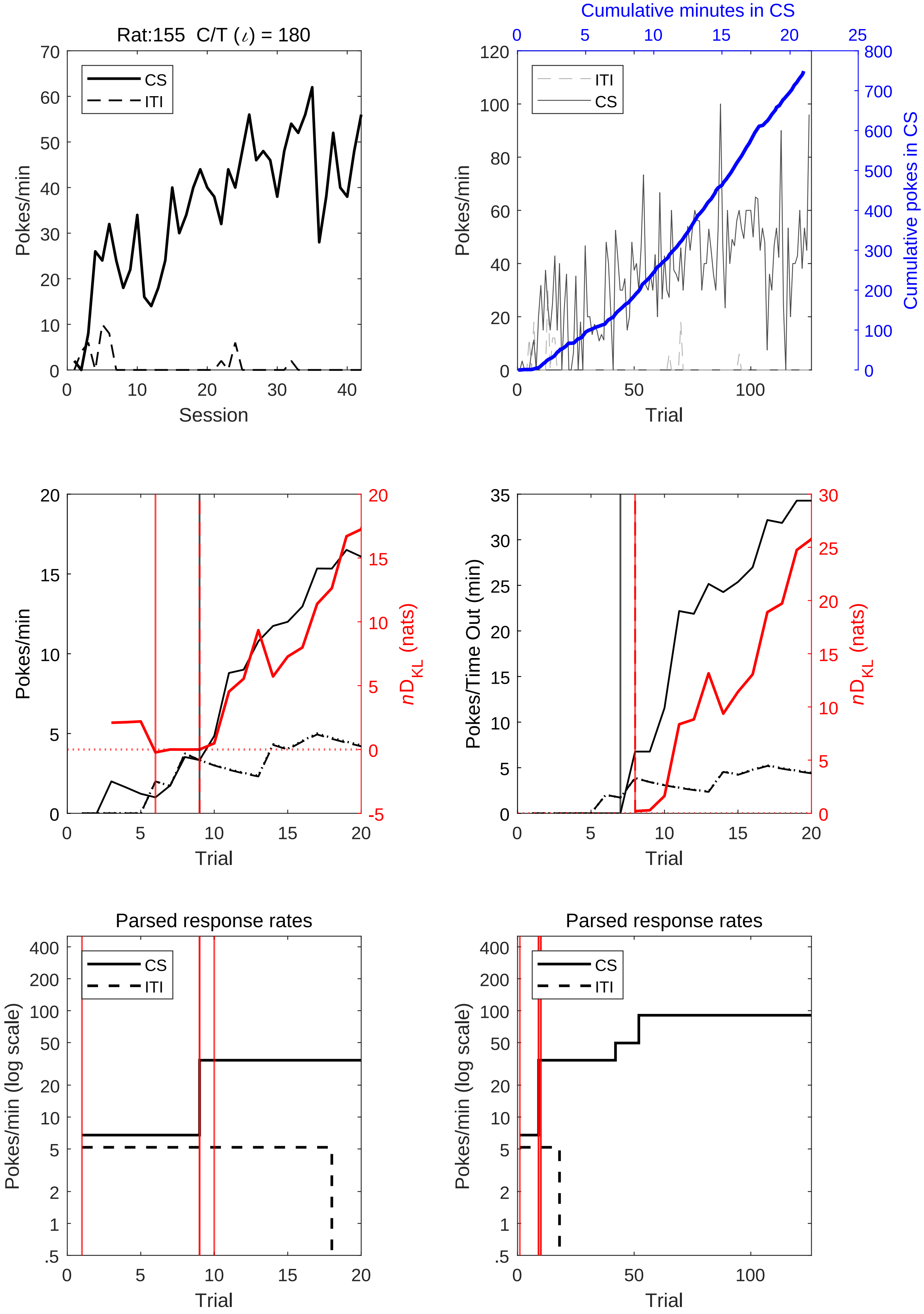

Rat:155 C/T (ι) = 180
CS
ITI
Pokes/min
Session
Cumulative minutes in CS
ITI
CS
Pokes/min
Cumulative pokes in CS
Trial
Pokes/min
nD_KL (nats)
Trial
Pokes/Time Out (min)
nD_KL (nats)
Trial
Parsed response rates
CS
ITI
Pokes/min (log scale)
Trial
Parsed response rates
CS
ITI
Pokes/min (log scale)
Trial

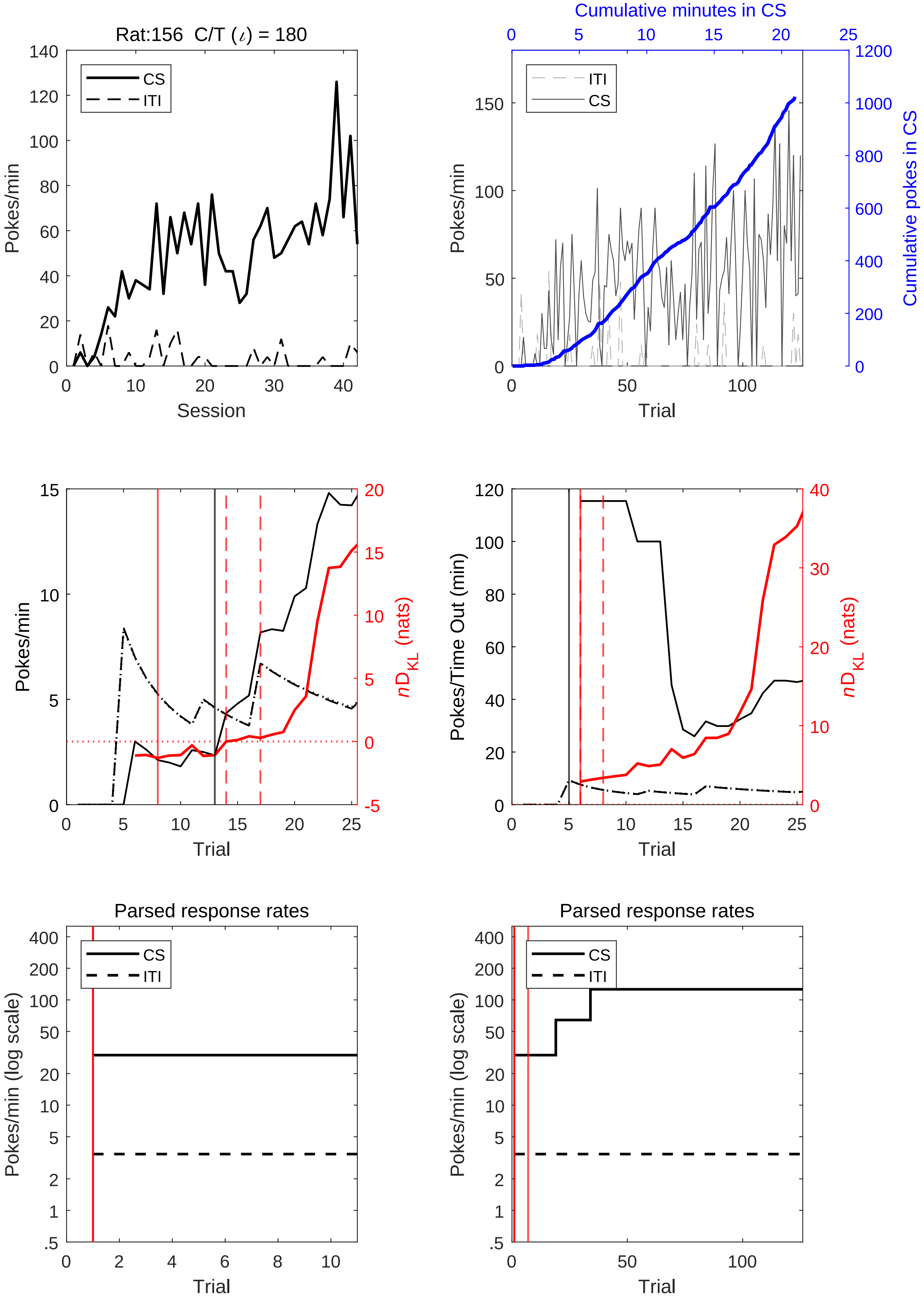
Rat:156 C/T (ι) = 180
CS
ITI
Pokes/min
Session
Cumulative minutes in CS
Cumulative pokes in CS
Trial
$nD_{KL}$ (nats)
Pokes/Time Out (min)
Parsed response rates
Pokes/min (log scale)

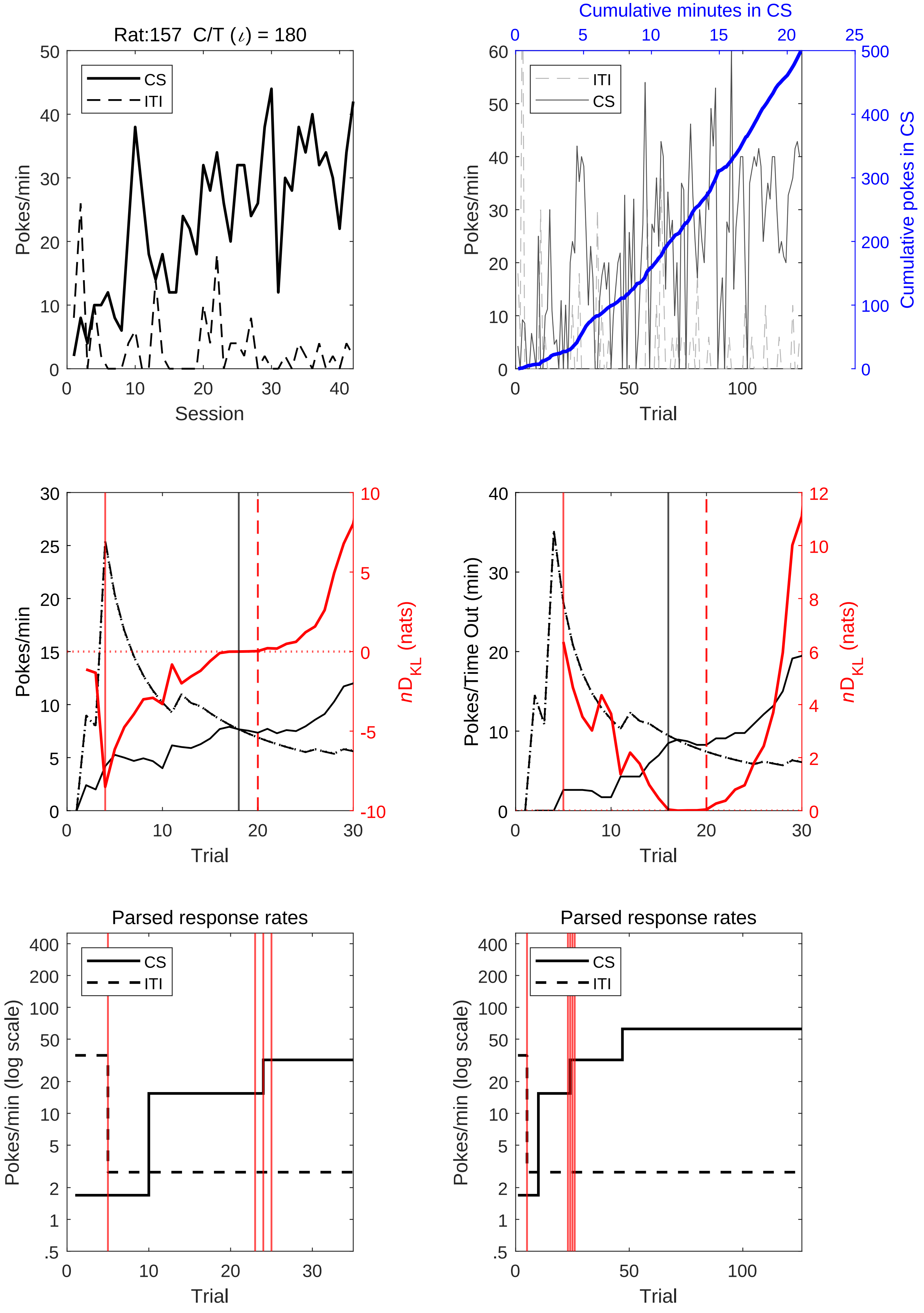
Rat:157 C/T (ι) = 180
CS
ITI
Pokes/min
Session
Cumulative minutes in CS
Cumulative pokes in CS
Trial
nD_KL (nats)
Pokes/Time Out (min)
Parsed response rates
Pokes/min (log scale)

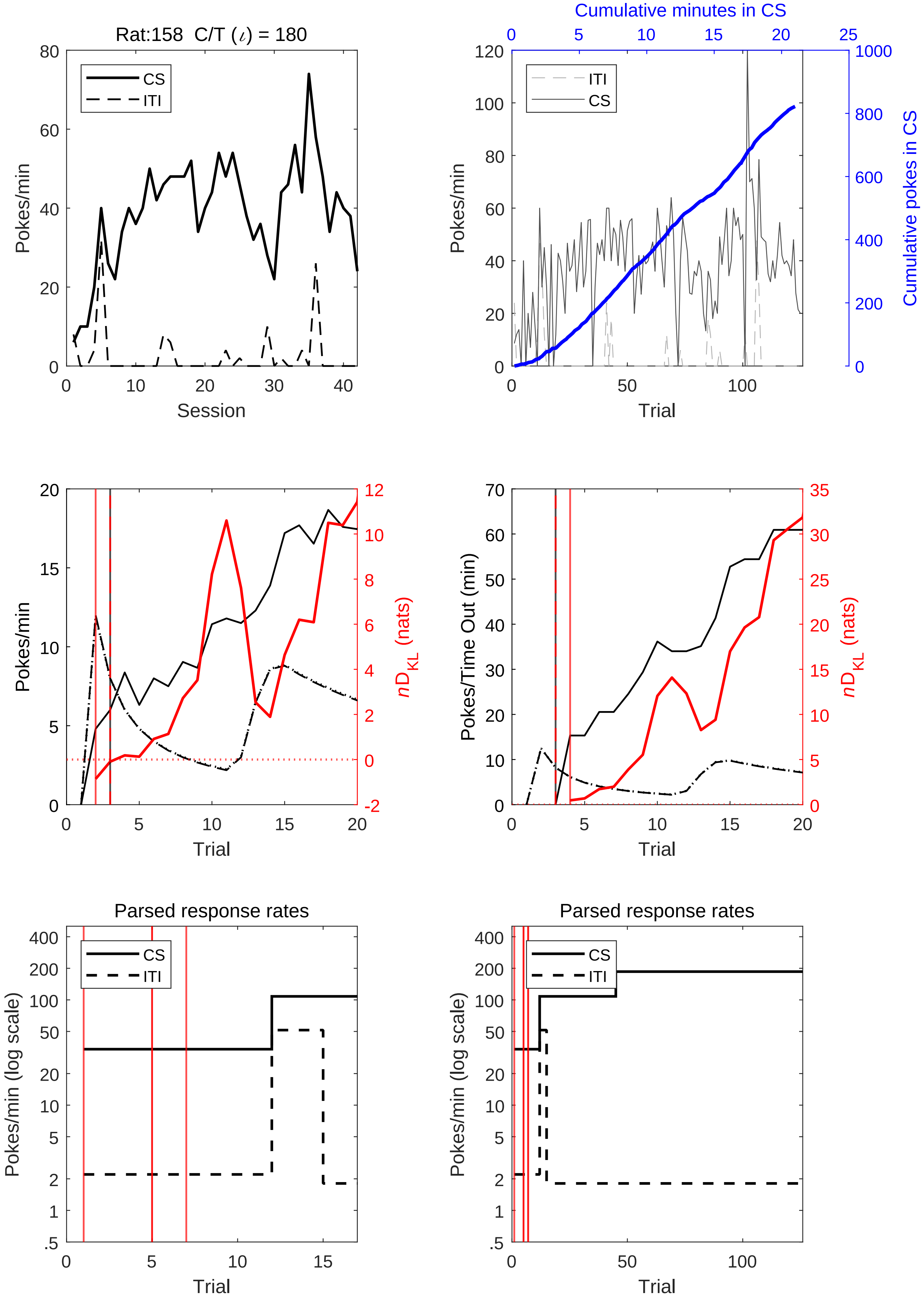
Rat:158 C/T (ι) = 180
Pokes/min
Session
CS
ITI
Cumulative minutes in CS
Cumulative pokes in CS
Trial
Pokes/Time Out (min)
nD_KL (nats)
Parsed response rates
Pokes/min (log scale)

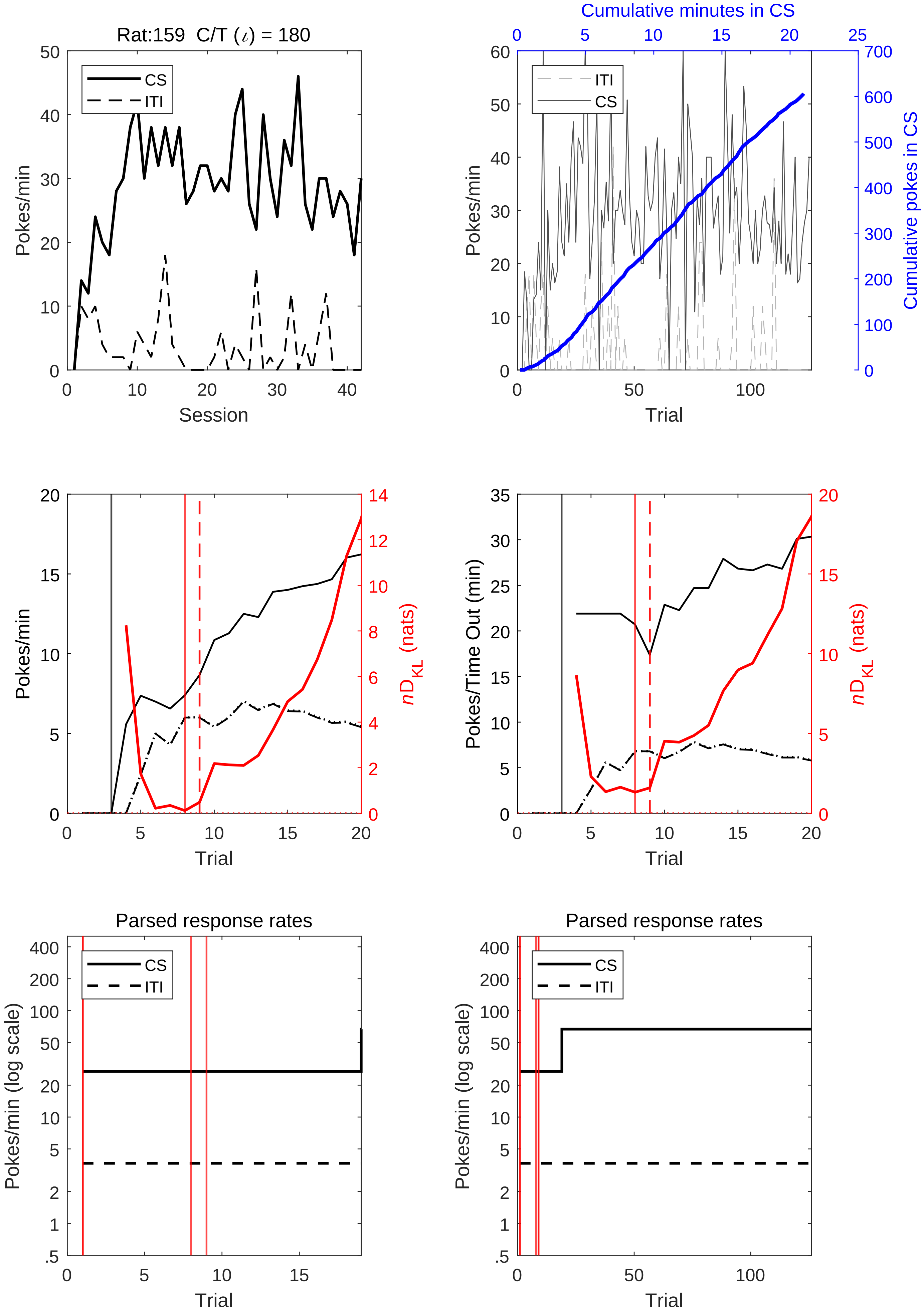
Rat:159 C/T (ι) = 180
CS
ITI
Pokes/min
Session
Cumulative minutes in CS
Cumulative pokes in CS
Trial
nD_KL (nats)
Pokes/Time Out (min)
Parsed response rates
Pokes/min (log scale)

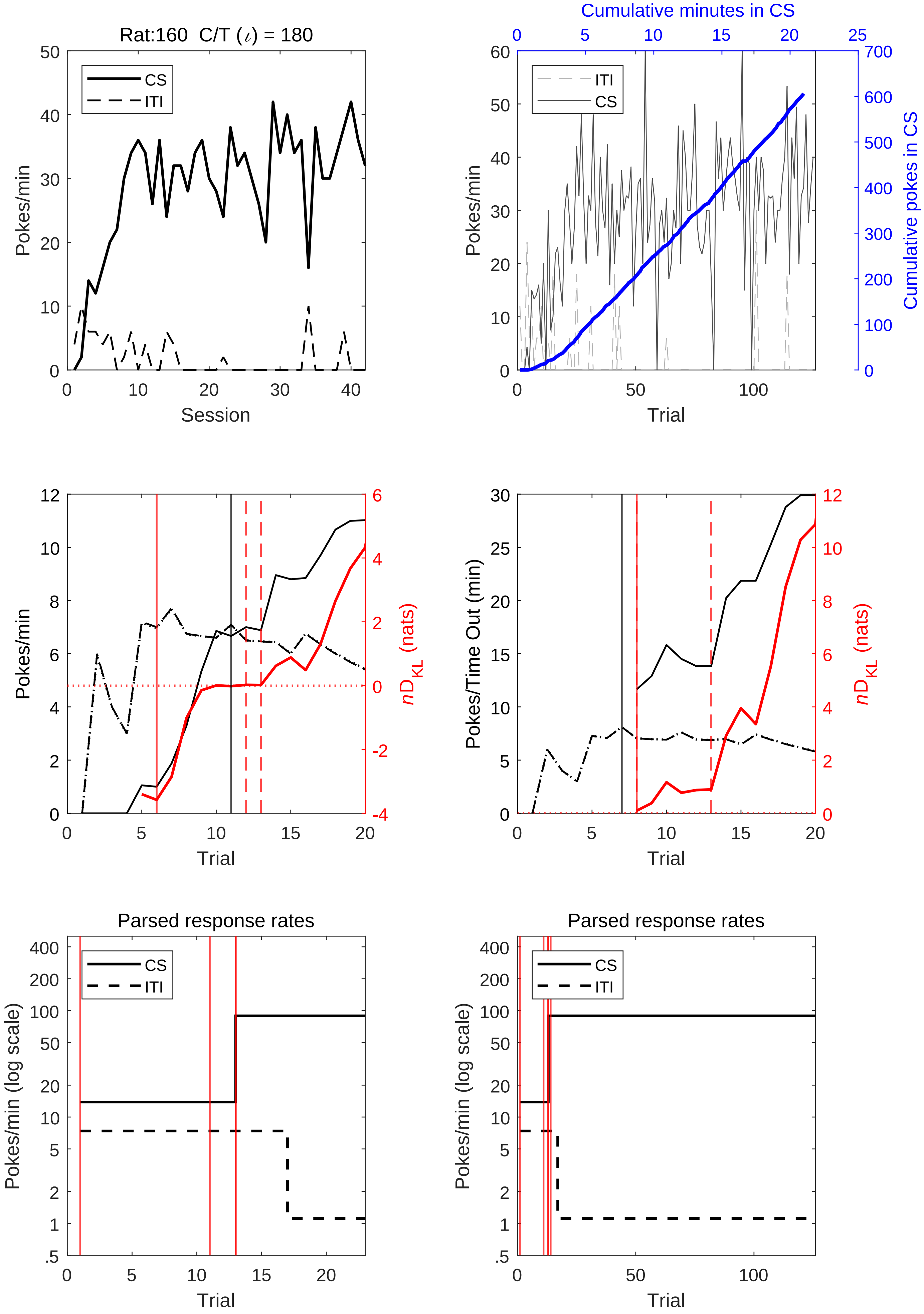
Rat:160 C/T ($\iota$) = 180
CS
ITI
Pokes/min
Session
Cumulative minutes in CS
Cumulative pokes in CS
Trial
Pokes/Time Out (min)
$nD_{KL}$ (nats)
Parsed response rates
Pokes/min (log scale)

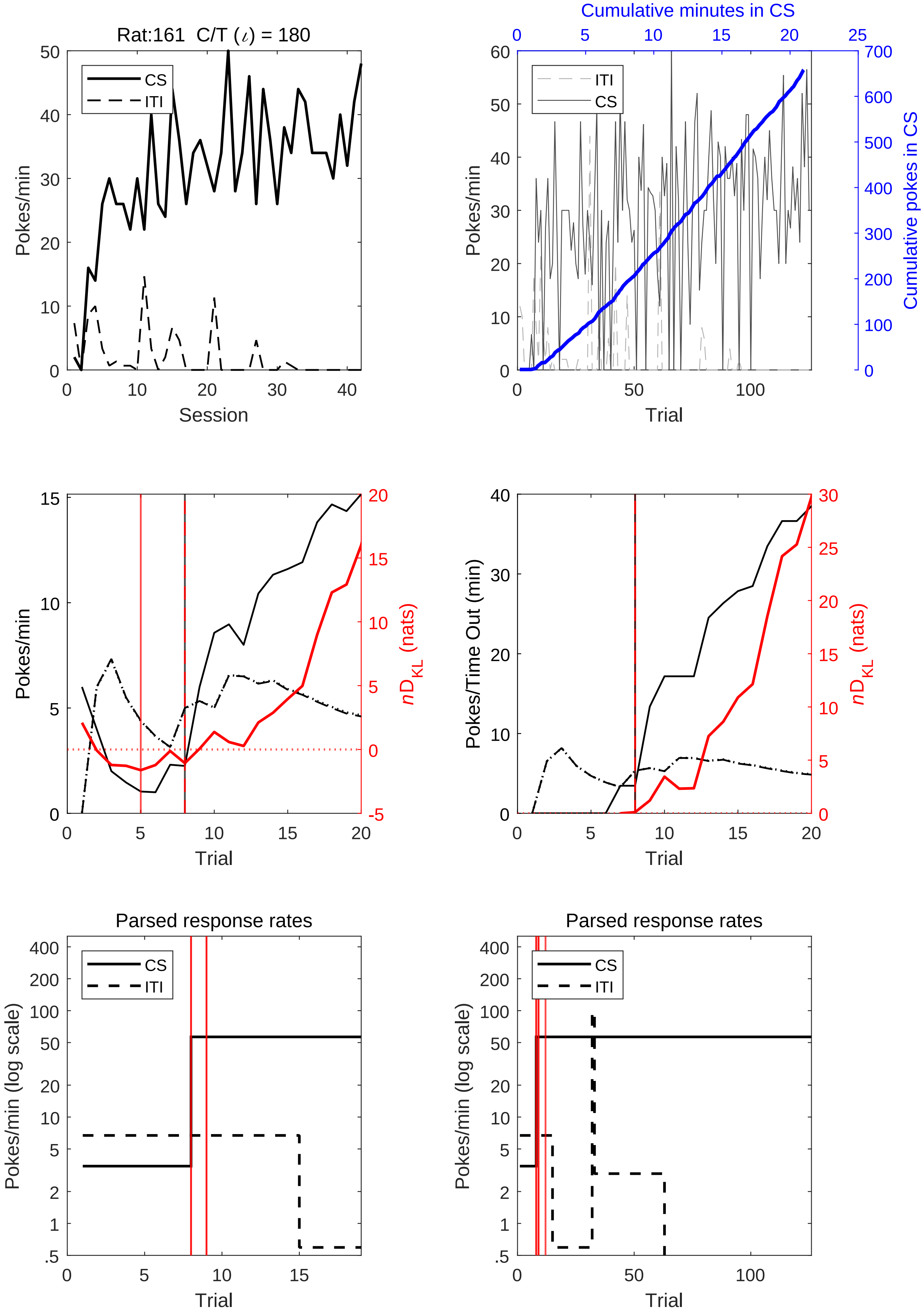

Rat:161 C/T (ι) = 180
CS
ITI
Pokes/min
Session
Cumulative minutes in CS
Cumulative pokes in CS
Trial
Pokes/Time Out (min)
$nD_{KL}$ (nats)
Parsed response rates
Pokes/min (log scale)

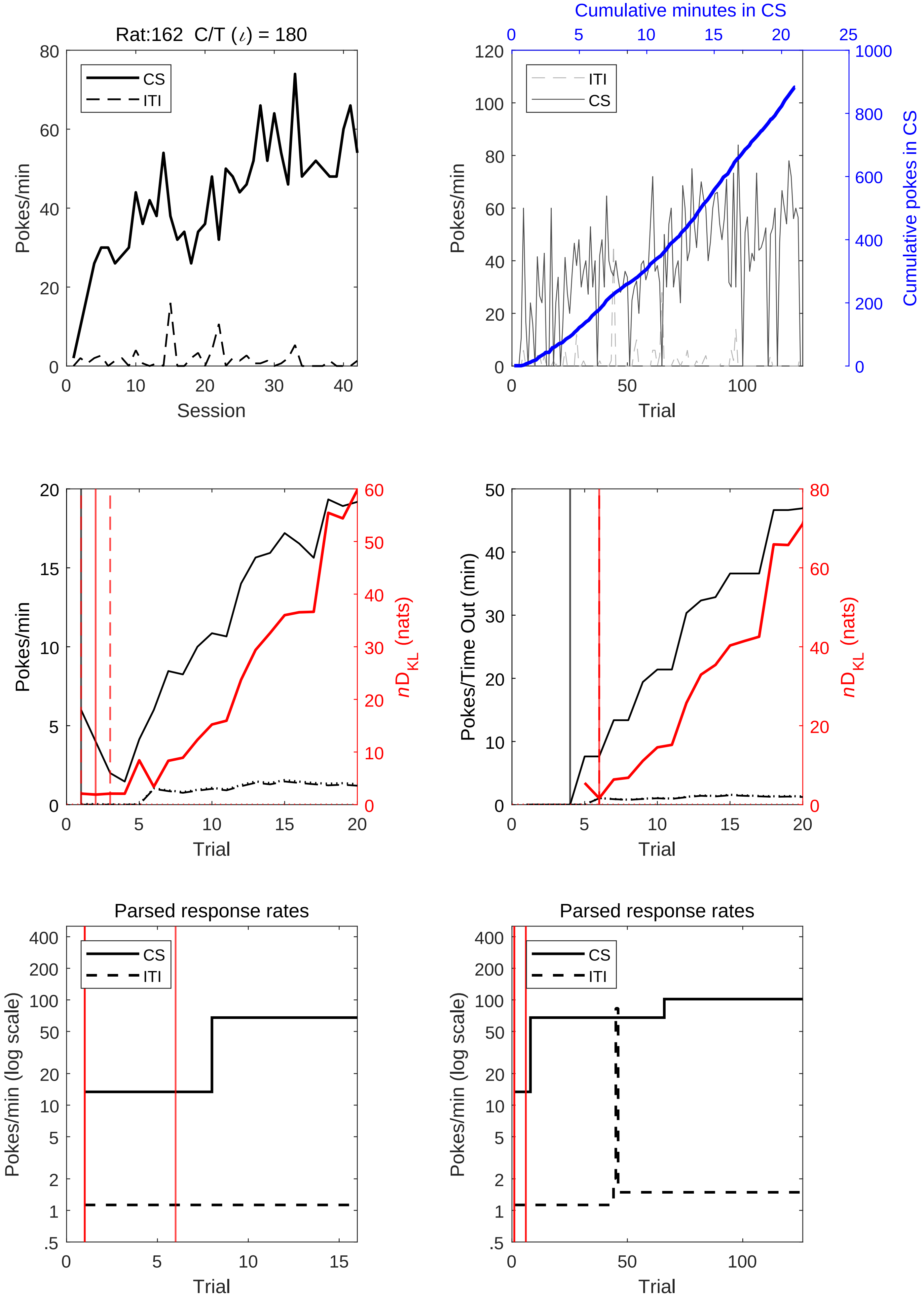
Rat:162 C/T (ι) = 180
CS
ITI
Pokes/min
Session
Cumulative minutes in CS
Cumulative pokes in CS
Trial
Pokes/min
Trial
nD_KL (nats)
Pokes/Time Out (min)
Trial
Parsed response rates
Pokes/min (log scale)
Trial
Parsed response rates
Pokes/min (log scale)
Trial

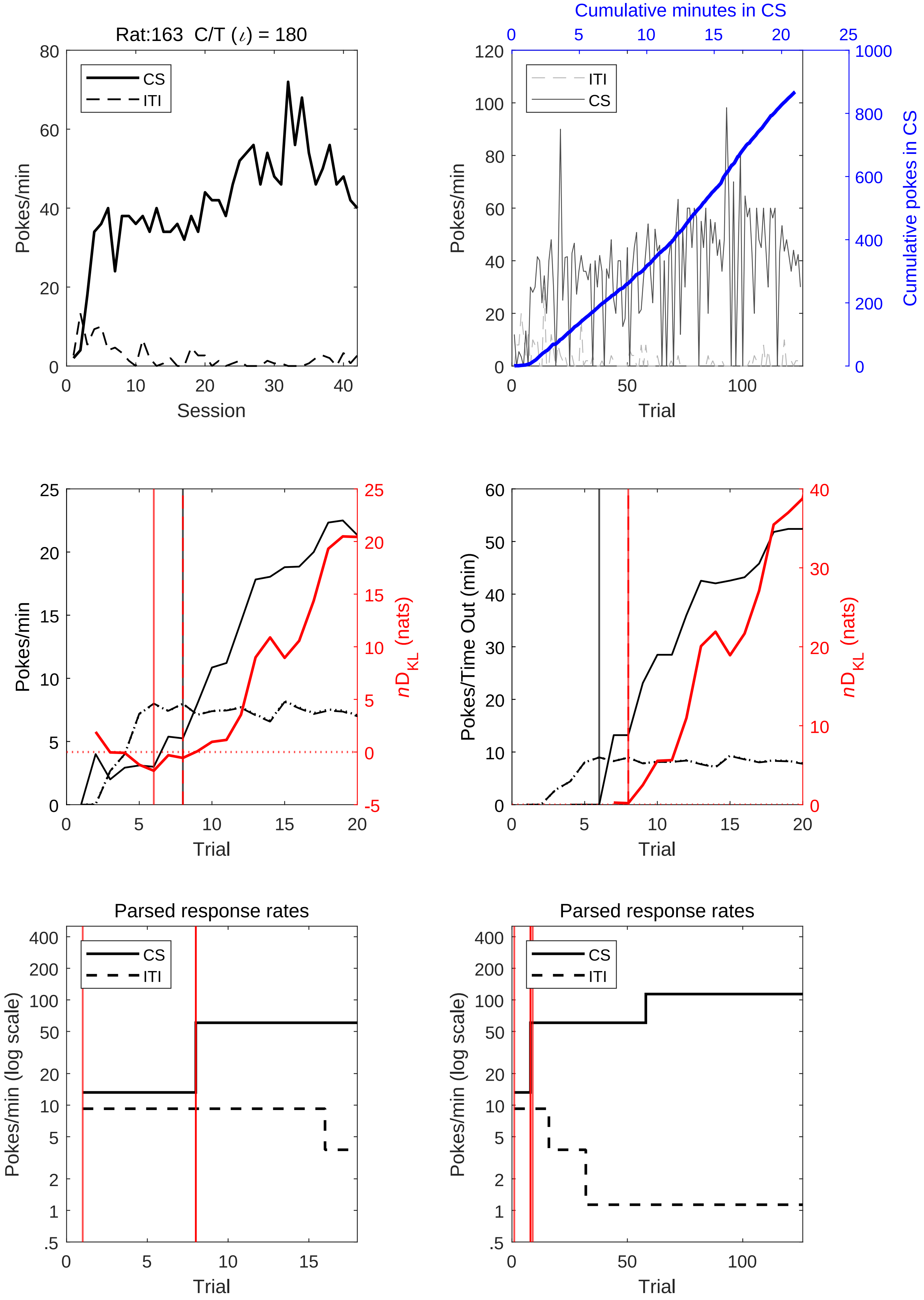
Rat:163 C/T (ι) = 180
CS
ITI
Pokes/min
Session
Cumulative minutes in CS
Cumulative pokes in CS
Trial
Pokes/Time Out (min)
nD_KL (nats)
Parsed response rates
Pokes/min (log scale)

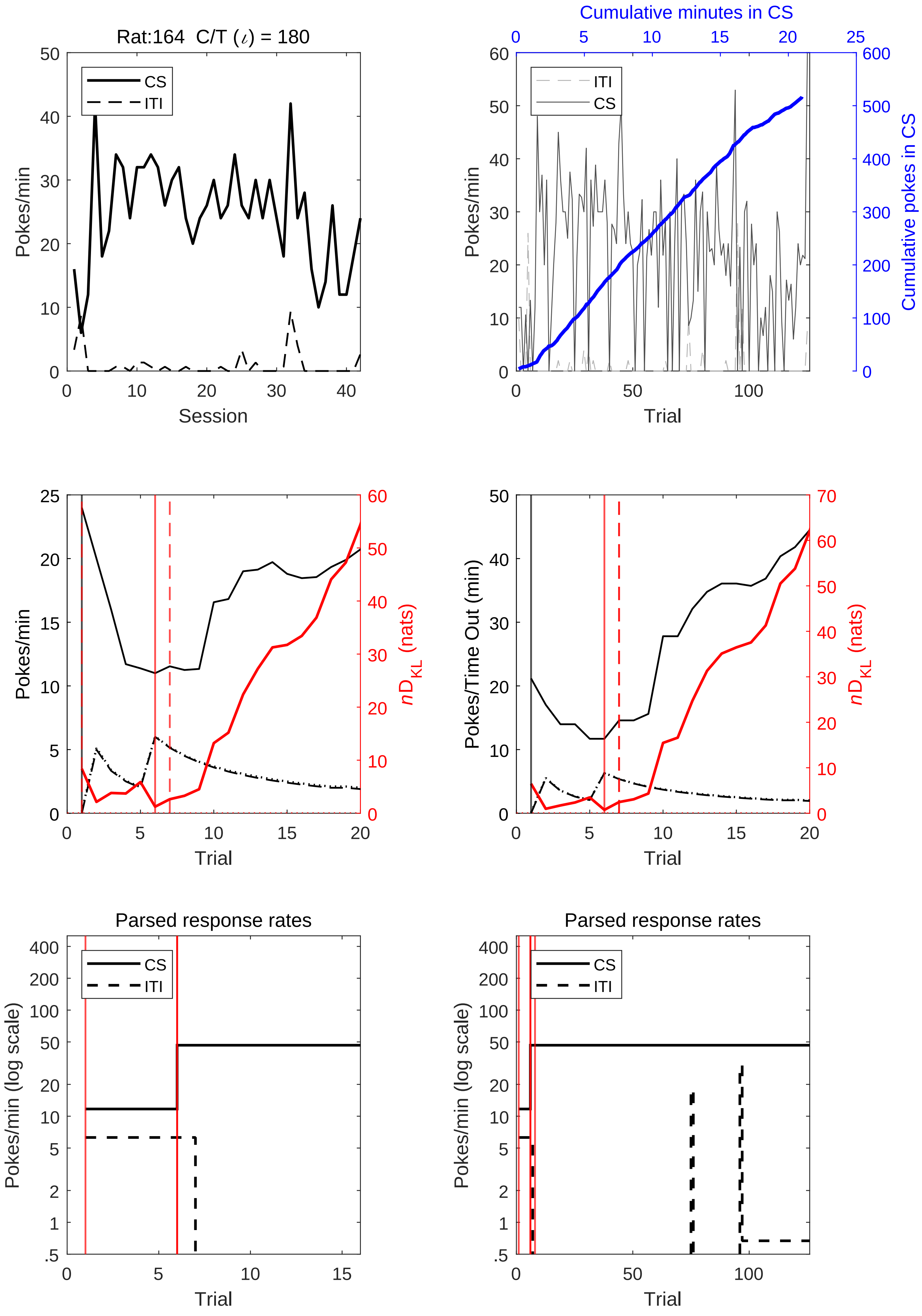
Rat:164 C/T (ι) = 180
CS
ITI
Pokes/min
Session
Cumulative minutes in CS
Cumulative pokes in CS
Trial
Pokes/Time Out (min)
nD_KL (nats)
Parsed response rates
Pokes/min (log scale)

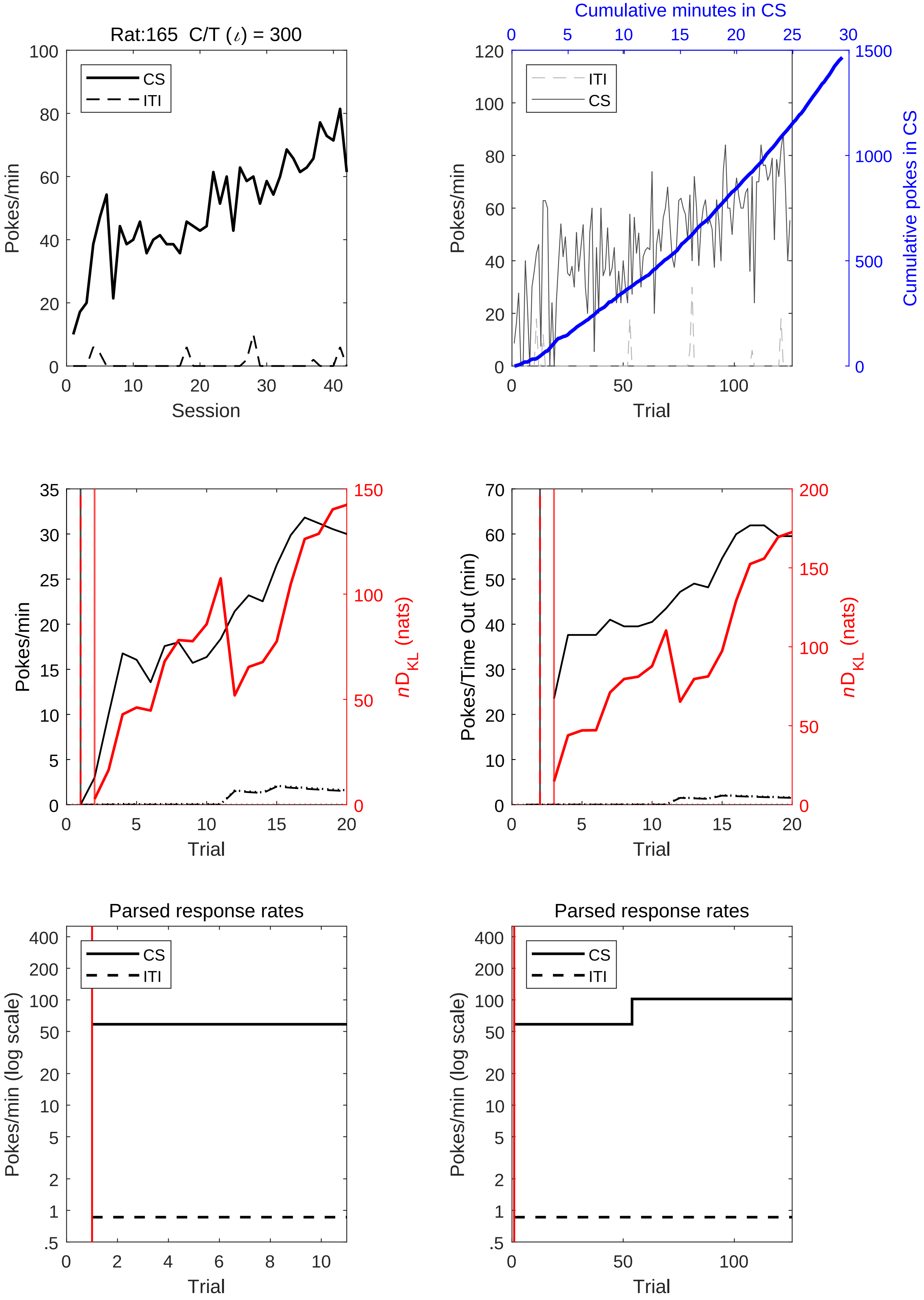
Rat:165 C/T (ι) = 300
Pokes/min
Session
CS
ITI
Cumulative minutes in CS
Cumulative pokes in CS
Pokes/min
Trial
ITI
CS
Pokes/min
Trial
nD_KL (nats)
Pokes/Time Out (min)
Trial
nD_KL (nats)
Parsed response rates
Pokes/min (log scale)
Trial
CS
ITI
Parsed response rates
Pokes/min (log scale)
Trial
CS
ITI

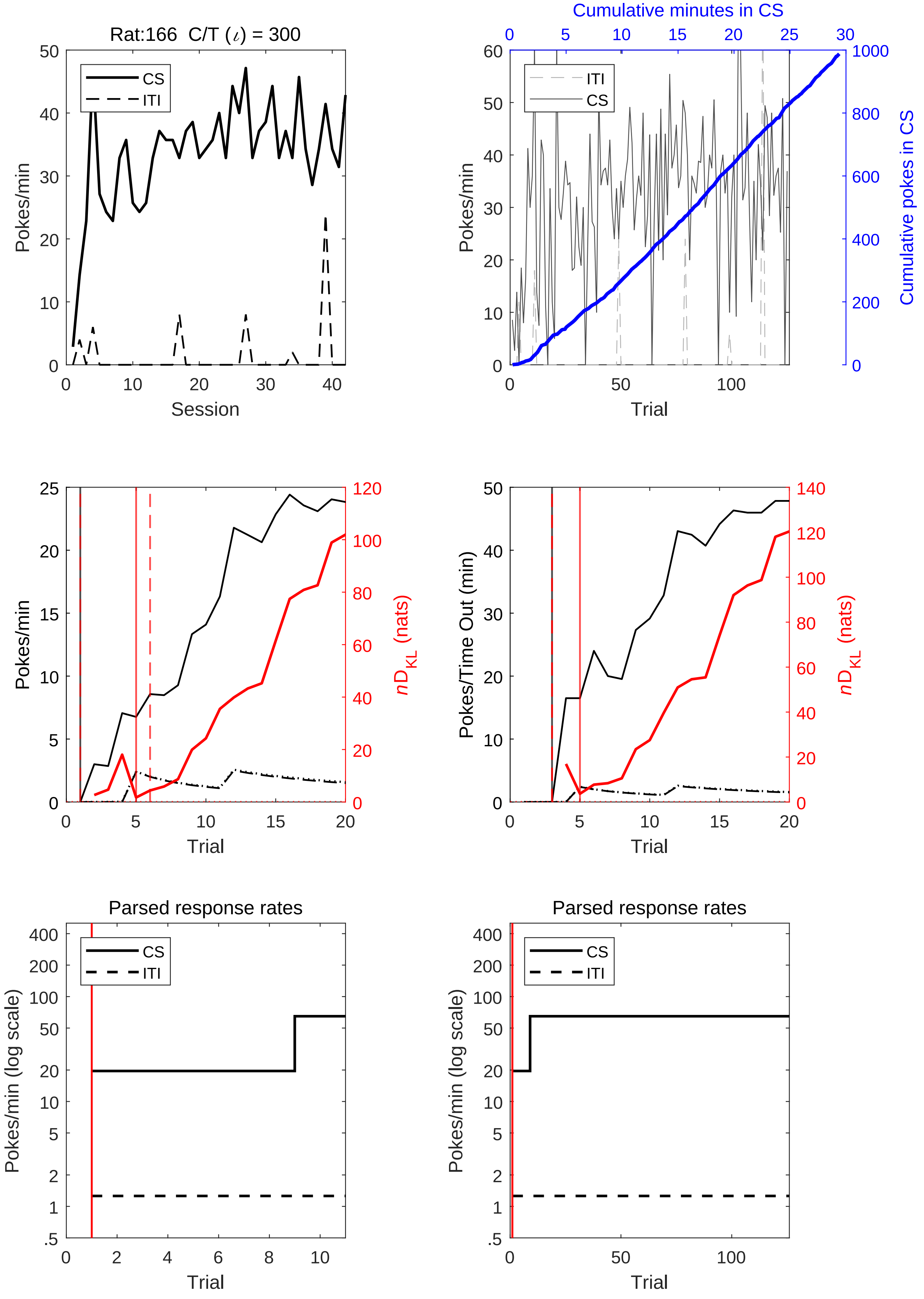

Rat:166 C/T ($\iota$) = 300
CS
ITI
Pokes/min
Session
Cumulative minutes in CS
ITI
CS
Pokes/min
Cumulative pokes in CS
Trial
Pokes/min
$nD_{KL}$ (nats)
Trial
Pokes/Time Out (min)
$nD_{KL}$ (nats)
Trial
Parsed response rates
CS
ITI
Pokes/min (log scale)
Trial
Parsed response rates
CS
ITI
Pokes/min (log scale)
Trial

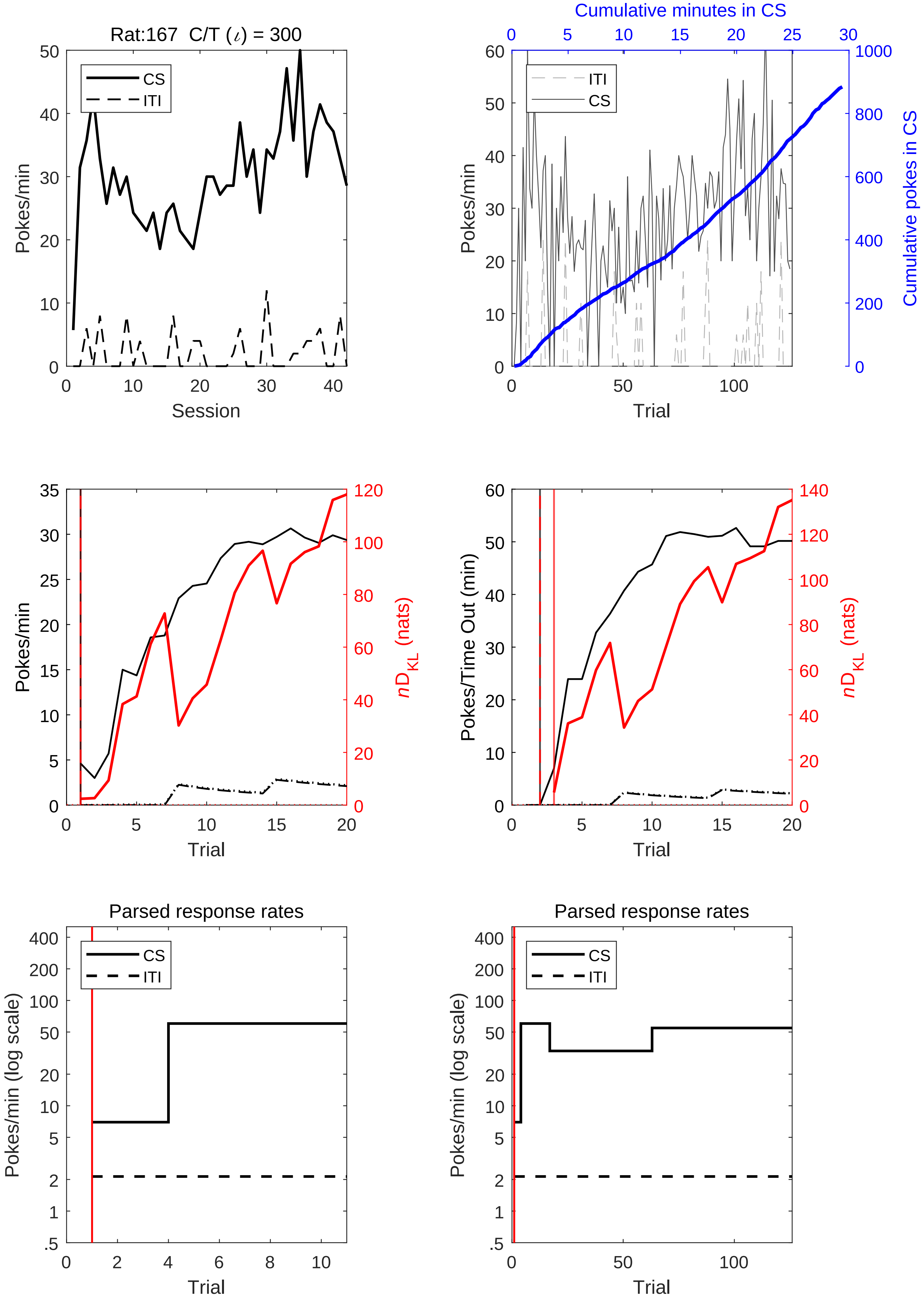
Rat:167 C/T (ι) = 300
CS
ITI
Pokes/min
Session
Cumulative minutes in CS
Pokes/min
Cumulative pokes in CS
Trial
Pokes/min
nD_KL (nats)
Trial
Pokes/Time Out (min)
nD_KL (nats)
Trial
Parsed response rates
Pokes/min (log scale)
Trial
Parsed response rates
Pokes/min (log scale)
Trial

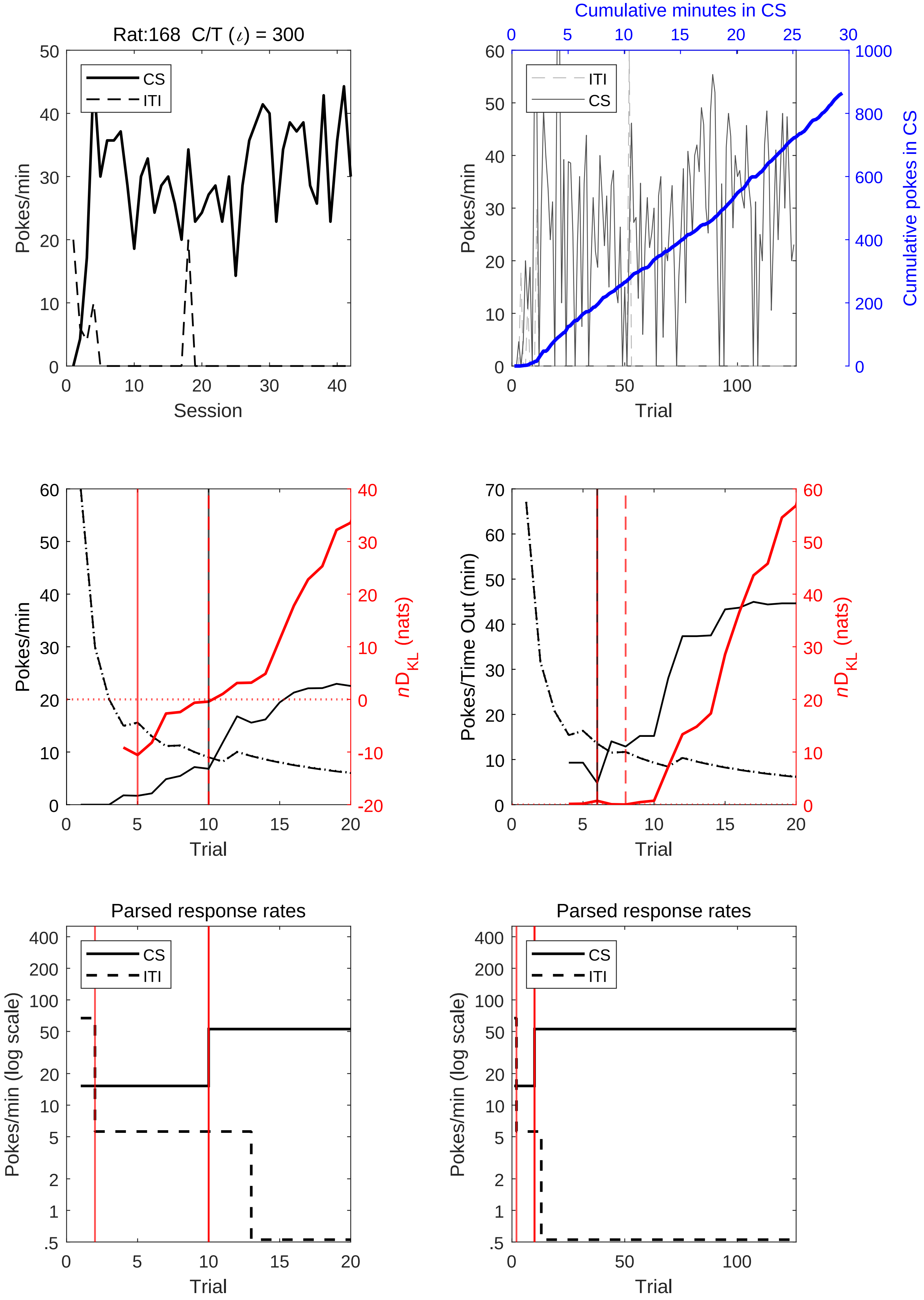
Rat:168 C/T (ι) = 300
CS
ITI
Pokes/min
Session
Cumulative minutes in CS
Cumulative pokes in CS
Trial
nD_KL (nats)
Pokes/Time Out (min)
Parsed response rates
Pokes/min (log scale)

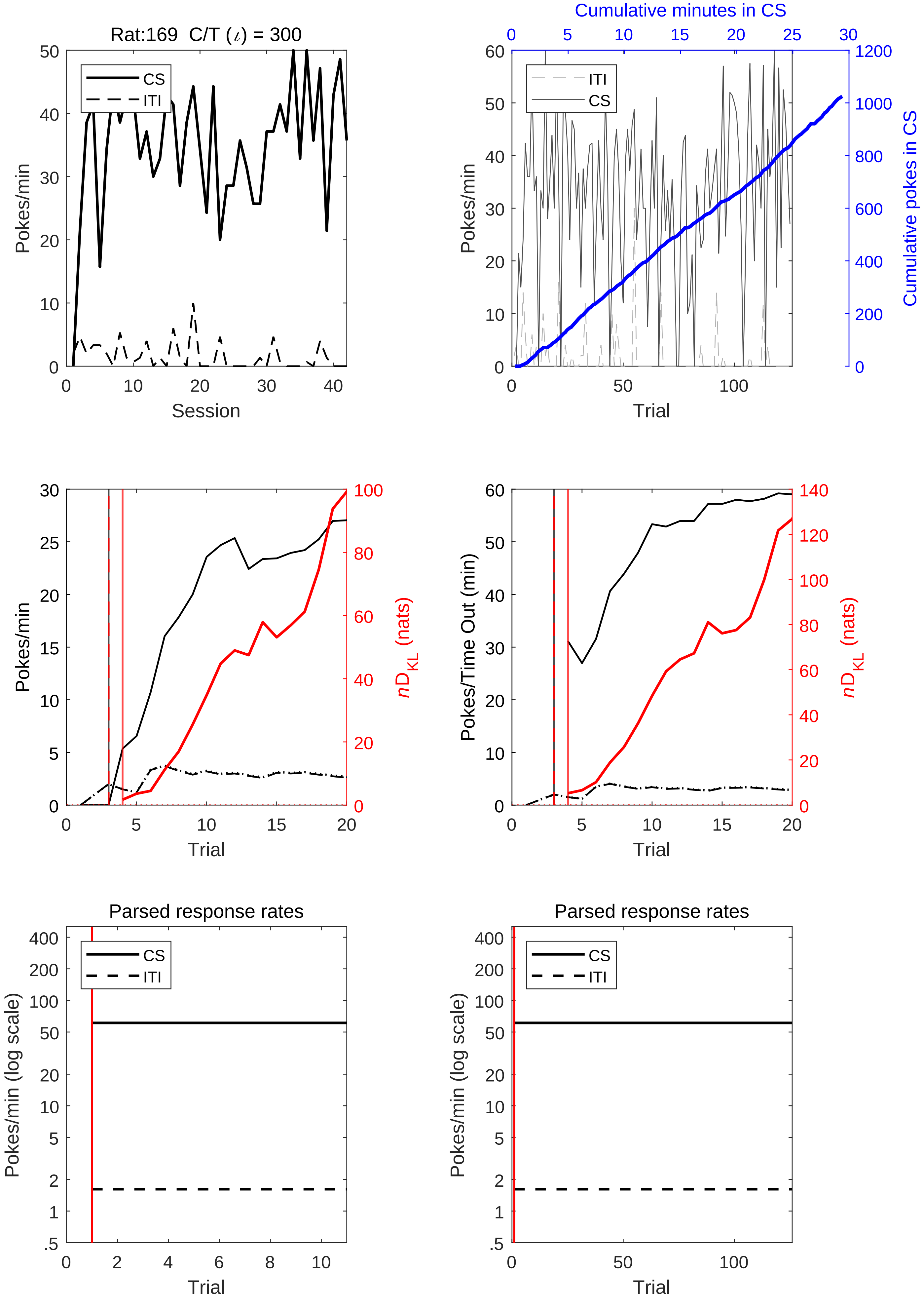

Rat:169 C/T (ι) = 300
CS
ITI
Pokes/min
Session
Cumulative minutes in CS
Cumulative pokes in CS
Trial
nD_KL (nats)
Pokes/Time Out (min)
Parsed response rates
Pokes/min (log scale)

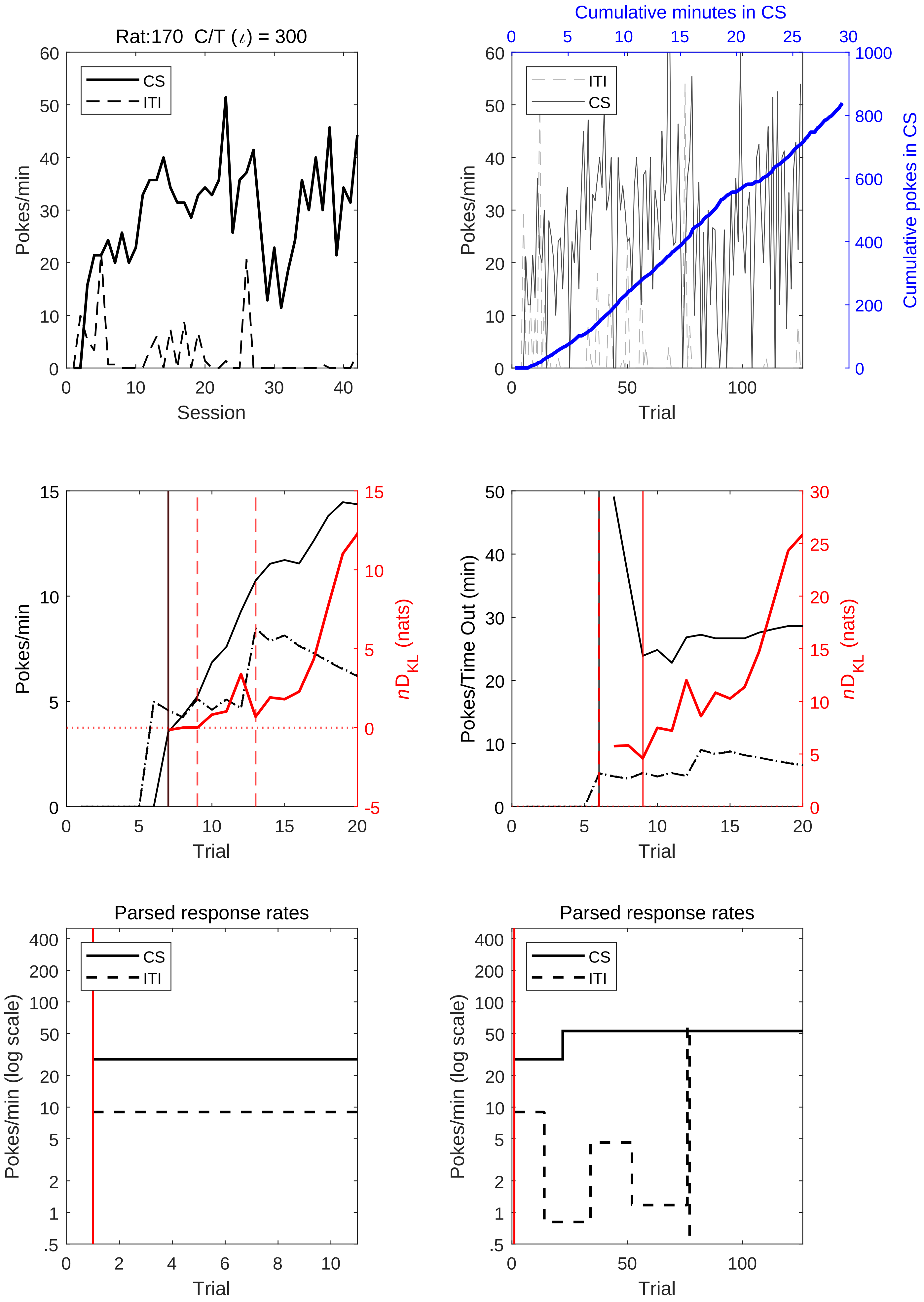
Rat:170  C/T (ι) = 300
CS
ITI
Pokes/min
Session
Cumulative minutes in CS
ITI
CS
Pokes/min
Cumulative pokes in CS
Trial
Pokes/min
nD_KL (nats)
Trial
Pokes/Time Out (min)
nD_KL (nats)
Trial
Parsed response rates
CS
ITI
Pokes/min (log scale)
Trial
Parsed response rates
CS
ITI
Pokes/min (log scale)
Trial

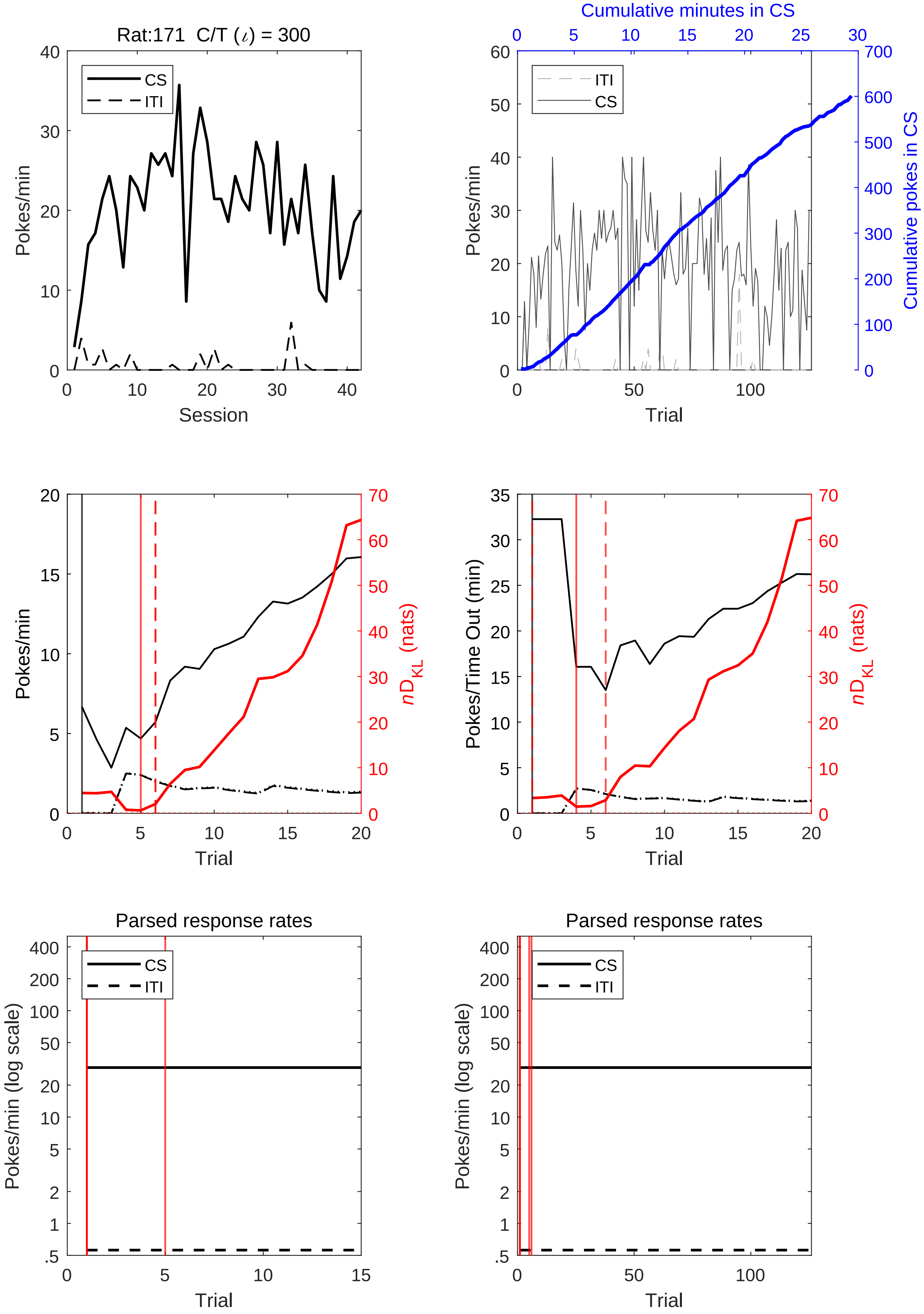
Rat:171 C/T (ι) = 300
CS
ITI
Pokes/min
Session
Cumulative minutes in CS
ITI
CS
Pokes/min
Cumulative pokes in CS
Trial
Pokes/min
nD_KL (nats)
Trial
Pokes/Time Out (min)
nD_KL (nats)
Trial
Parsed response rates
CS
ITI
Pokes/min (log scale)
Trial
Parsed response rates
CS
ITI
Pokes/min (log scale)
Trial

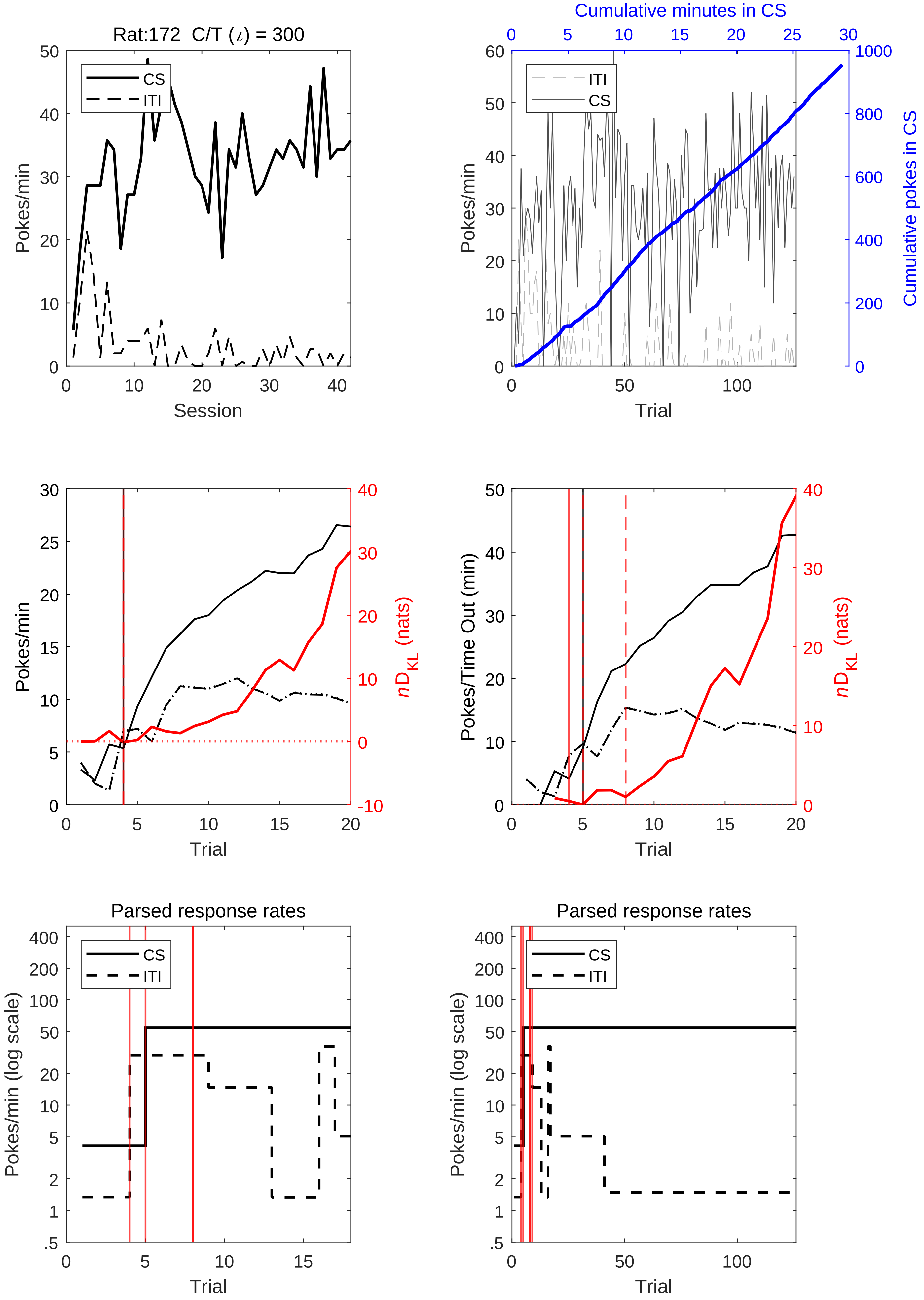

Rat:172 C/T (ι) = 300
CS
ITI
Pokes/min
Session
Cumulative minutes in CS
Cumulative pokes in CS
Trial
Pokes/Time Out (min)
$nD_{KL}$ (nats)
Parsed response rates
Pokes/min (log scale)

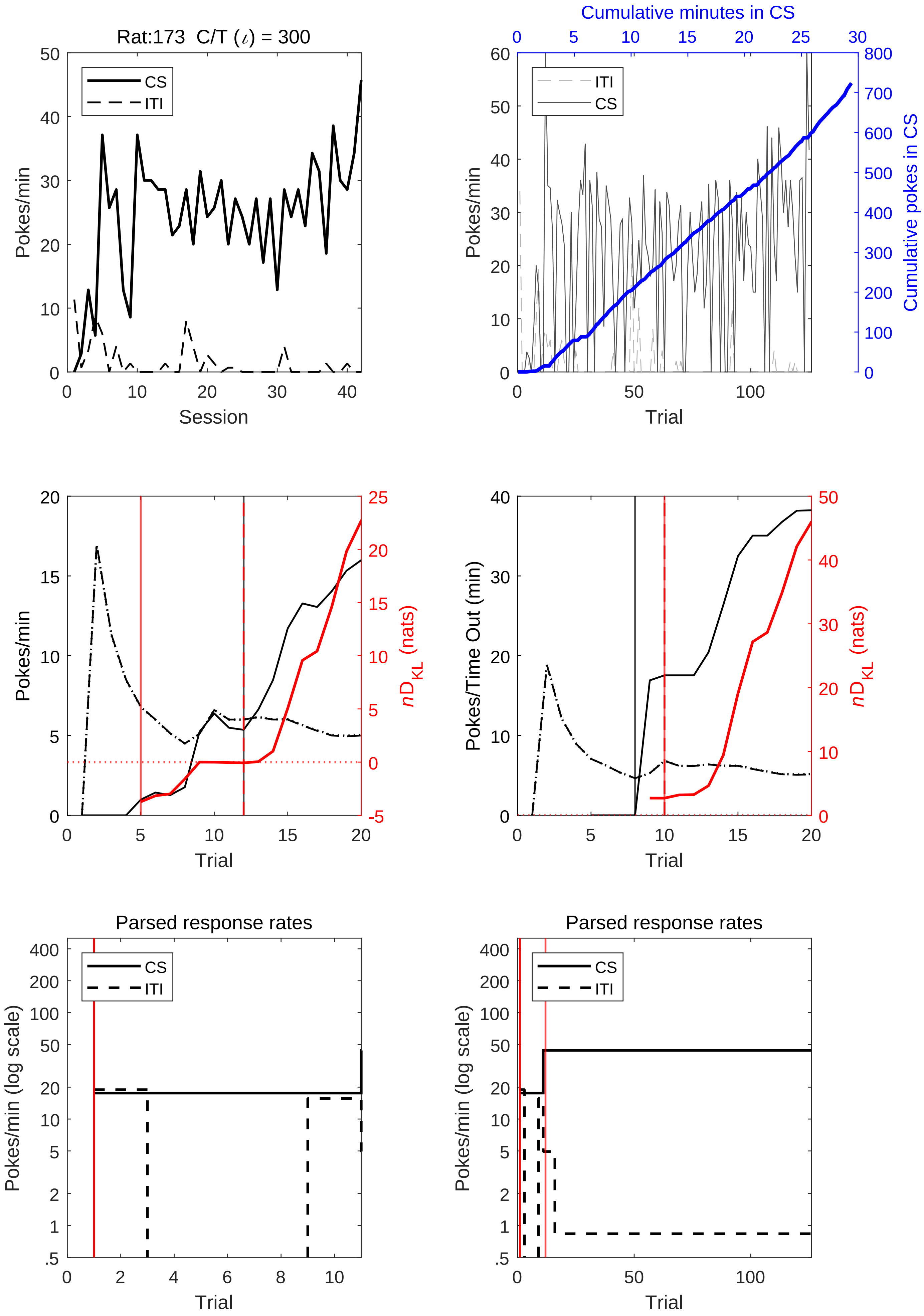
Rat:173  C/T (ι) = 300
CS
ITI
Pokes/min
Session
Cumulative minutes in CS
Pokes/min
Cumulative pokes in CS
Trial
Pokes/min
nD_KL (nats)
Trial
Pokes/Time Out (min)
nD_KL (nats)
Trial
Parsed response rates
Pokes/min (log scale)
Trial
Parsed response rates
Pokes/min (log scale)
Trial

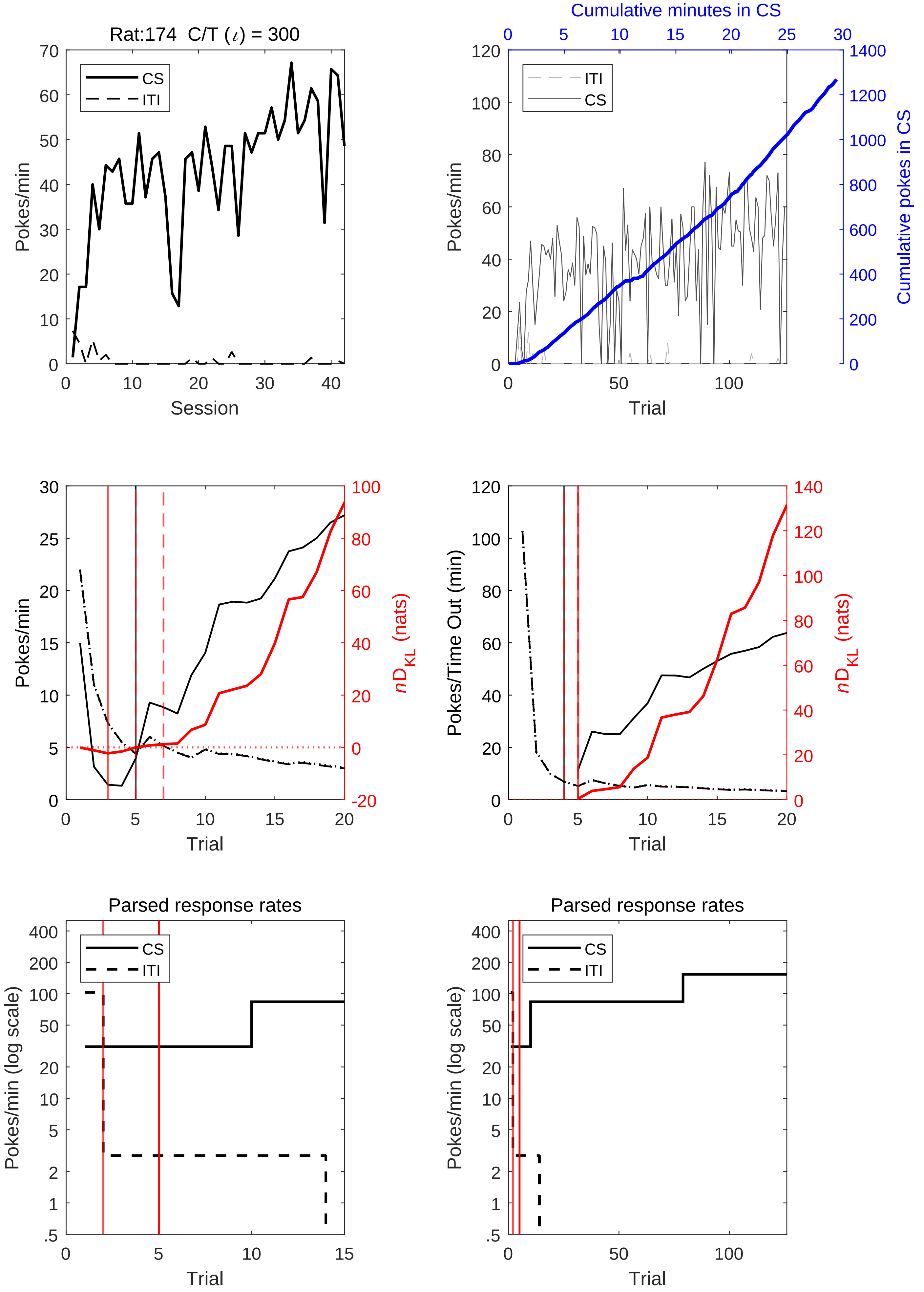

Rat:174 C/T (ι) = 300
CS
ITI
Pokes/min
Session
Cumulative minutes in CS
Cumulative pokes in CS
Trial
nD_KL (nats)
Pokes/Time Out (min)
Parsed response rates
Pokes/min (log scale)

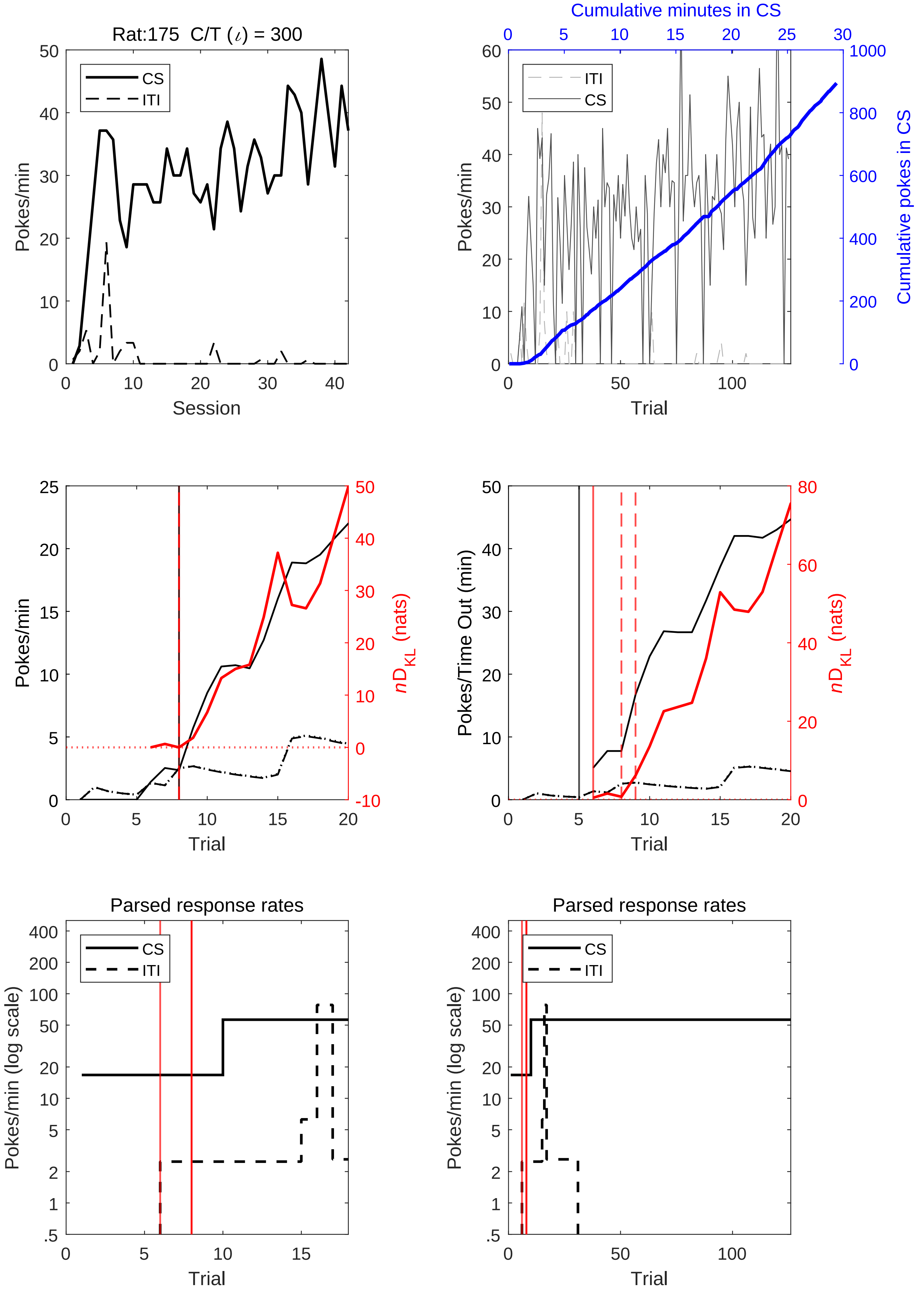
Rat:175 C/T (ι) = 300
CS
ITI
Pokes/min
Session
Cumulative minutes in CS
Cumulative pokes in CS
Trial
nD_KL (nats)
Pokes/Time Out (min)
Parsed response rates
Pokes/min (log scale)

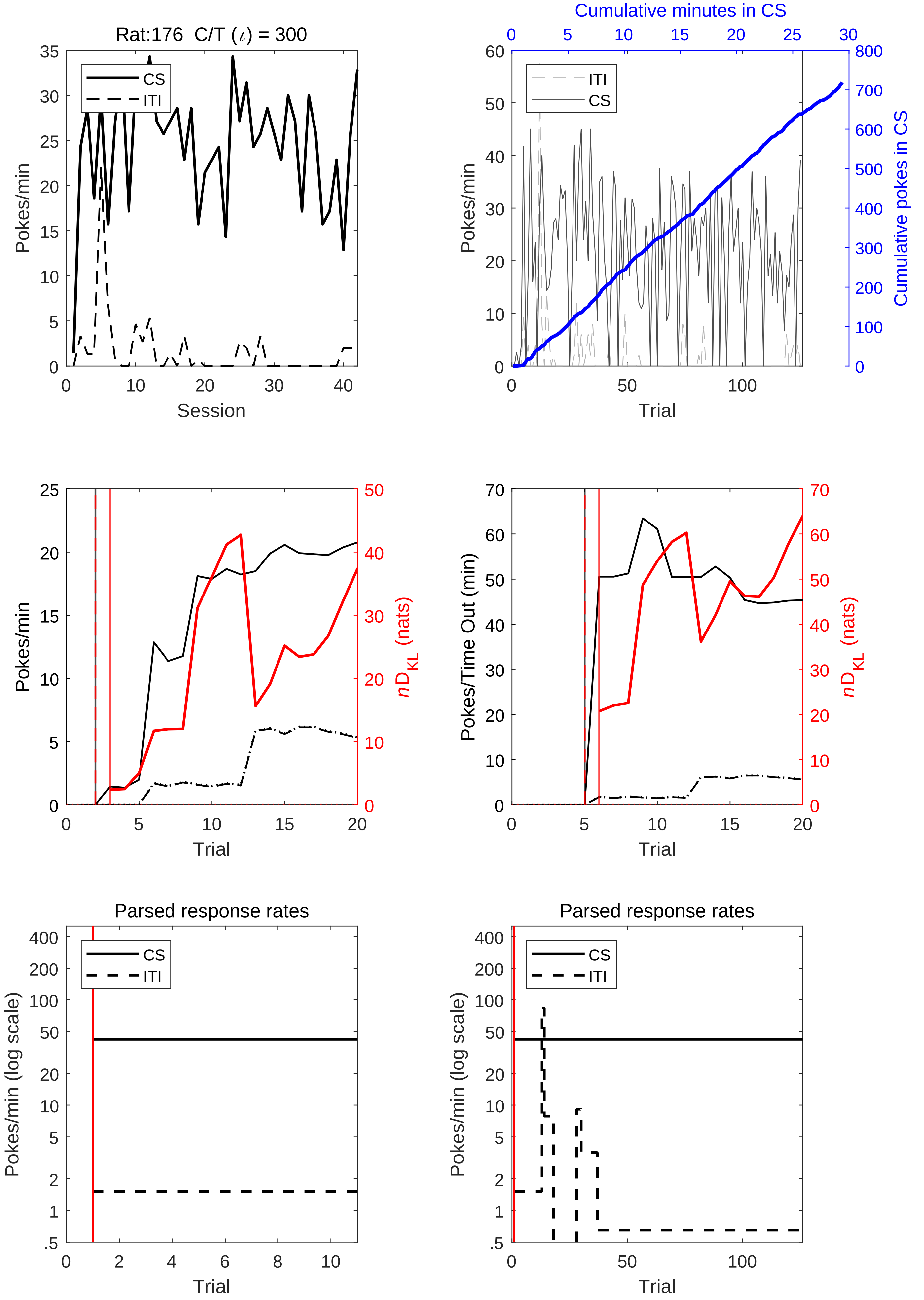

Rat:176 C/T ($\iota$) = 300
Pokes/min
Session
CS
ITI
Cumulative minutes in CS
Cumulative pokes in CS
Trial
Pokes/Time Out (min)
$nD_{KL}$ (nats)
Parsed response rates
Pokes/min (log scale)
